# Target-based discovery of a broad spectrum flukicide

**DOI:** 10.1101/2023.09.22.559026

**Authors:** Daniel J. Sprague, Sang-Kyu Park, Svenja Gramberg, Lisa Bauer, Claudia M. Rohr, Evgeny G. Chulkov, Emery Smith, Louis Scampavia, Timothy P. Spicer, Simone Haeberlein, Jonathan S. Marchant

**Author notes:** Supplementary Information (Supplementary Figures x5, Supplementary Tables x3, Supplementary Videos x4, Synthetic Procedures and Characterization Data).

## Abstract

**Diseases caused by parasitic flatworms impart a considerable healthcare burden worldwide. Many of these diseases – for example, the parasitic blood fluke infection, schistosomiasis – are treated with the drug praziquantel (PZQ). However, PZQ is ineffective against disease caused by liver flukes from the genus *Fasciola*. This is due to a single amino acid change within the target of PZQ, a transient receptor potential ion channel (TRPM_PZQ_), in *Fasciola* species. Here we identify benzamidoquinazolinone analogs that are active against *Fasciola* TRPM_PZQ_. Structure-activity studies define an optimized ligand (BZQ) that caused protracted paralysis and damage to the protective tegument of these liver flukes. BZQ also retained activity against *Schistosoma mansoni* comparable to PZQ and was active against TRPM_PZQ_ orthologs in all profiled species of parasitic fluke. This broad spectrum activity was manifest as BZQ adopts a pose within the binding pocket of TRPM_PZQ_ dependent on a ubiquitously conserved residue. BZQ therefore acts as a universal activator of trematode TRPM_PZQ_ and a first-in-class, broad spectrum flukicide.**

## Introduction

Trematodes (parasitic flukes) cause various diseases in humans. Blood flukes from the genus *Schistosoma* cause schistosomiasis, a disease that afflicts over 200 million people worldwide. Liver fluke, lung flukes, as well as intestinal flukes cause various food-borne trematodiases that add to the global neglected tropical disease burden. These parasitic flatworm infections are treated using a drug called praziquantel (PZQ), which has been the mainstay of clinical therapy for over 40 years [1-3].

However, PZQ is not effective against all types of flukes. Notably, PZQ lacks efficacy against liver flukes from the genus *Fasciola* (for example, *Fasciola hepatica* and *Fasciola gigantica*) [4, 5]. These parasites cause fasciolosis, a food-borne infection and zoonosis afflicting both humans and livestock. Fasciolosis-related complications in agriculture underpin considerable financial losses [6]. Fasciolosis is currently treated with triclabendazole (TCBZ), however widespread agricultural exposure has led to TCBZ resistance in Europe, South America, and Oceania [7-9]. Therefore, from a perspective of both food security and efficacy in the clinic, there is an unmet need to develop new drugs effective against these liver flukes [10, 11].

A logical strategy would be to understand why PZQ is ineffective against *Fasciola* spp., and then iterate solutions for broadening the spectrum of PZQ action. Such an approach has been made feasible by the recent discovery of the parasite target of PZQ [12], a transient receptor potential ion channel in the melastatin family called TRPM_PZQ_ [12-14]. PZQ activates TRPM_PZQ_ by engaging a binding pocket within the voltage-sensor like domain (transmembrane helices S1-S4) of the ion channel [15]. Critically, TRPM_PZQ_ in *Fasciola spp.* exhibits a natural amino acid variant (a threonine in the S1 helix of the binding pocket compared to an asparagine found in all other flukes [15, 16]). This difference likely removes a critical interaction needed for PZQ-evoked channel activation, rendering PZQ inactive at *Fasciola* TRPM_PZQ_ [15, 16] and thereby ineffective as a treatment for fasciolosis.

Based on this recent insight, we reasoned that a ligand that activates TRPM_PZQ_ but is tolerant of this naturally occurring binding pocket variation would act as a broad spectrum flukicide. This study reports the identification and characterization of such a chemotype overcomingthe insensitivity of *Fasciola spp.* to PZQ, and outperforming TCBZ, the current gold-standard therapy.

## Results

Target-based screening was used to identify TRPM_PZQ_ activators [17]. Using a fluorometric Ca^2+^ reporter assay, HEK293 cells that inducibly expressed either *Schistosoma mansoni* TRPM_PZQ_ (*Sm*.TRPM_PZQ_) or *Fasciola hepatica* TRPM_PZQ_ (*Fh*.TRPM_PZQ_) were treated with increasing concentrations of different ligands. The workflow prioritized identification of *Sm*.TRPM_PZQ_ agonists, given praziquantel (**(±)-PZQ**) provided a known positive control for *Sm*.TRPM_PZQ_ activation. Racemic PZQ (**(±)-PZQ**) activated *Sm*.TRPM_PZQ_ (**Figure 1A**) but not *Fh*.TRPM_PZQ_ (**Figure 1B**), acting as a potent agonist of *Sm*.TRPM_PZQ_ (EC_50_ of 0.18 ± 0.02 µM, **Figure 1C**)). These data were consistent with prior results [15, 16], and the known lack of effectiveness of PZQ in treating fasciolosis. Using this approach, a larger pool of *Sm*.TRPM_PZQ_ activators identified from in-house screens were counter-screened against *Fh*.TRPM_PZQ_. This led to the discovery of compound **1**, an *N*-benzamidoquinazolinone that activated both *Sm*.TRPM_PZQ_ and *Fh*.TRPM_PZQ_ (**Figure 1D & E**). Compound **1** was an agonist at both *Sm*.TRPM_PZQ_ (EC_50_ = 1.15 ± 0.11 µM) and *Fh*.TRPM_PZQ_ (EC_50_ = 3.0 ± 0.57 µM) active in the low micromolar range (**Figure 1F**).

**Figure 1.**
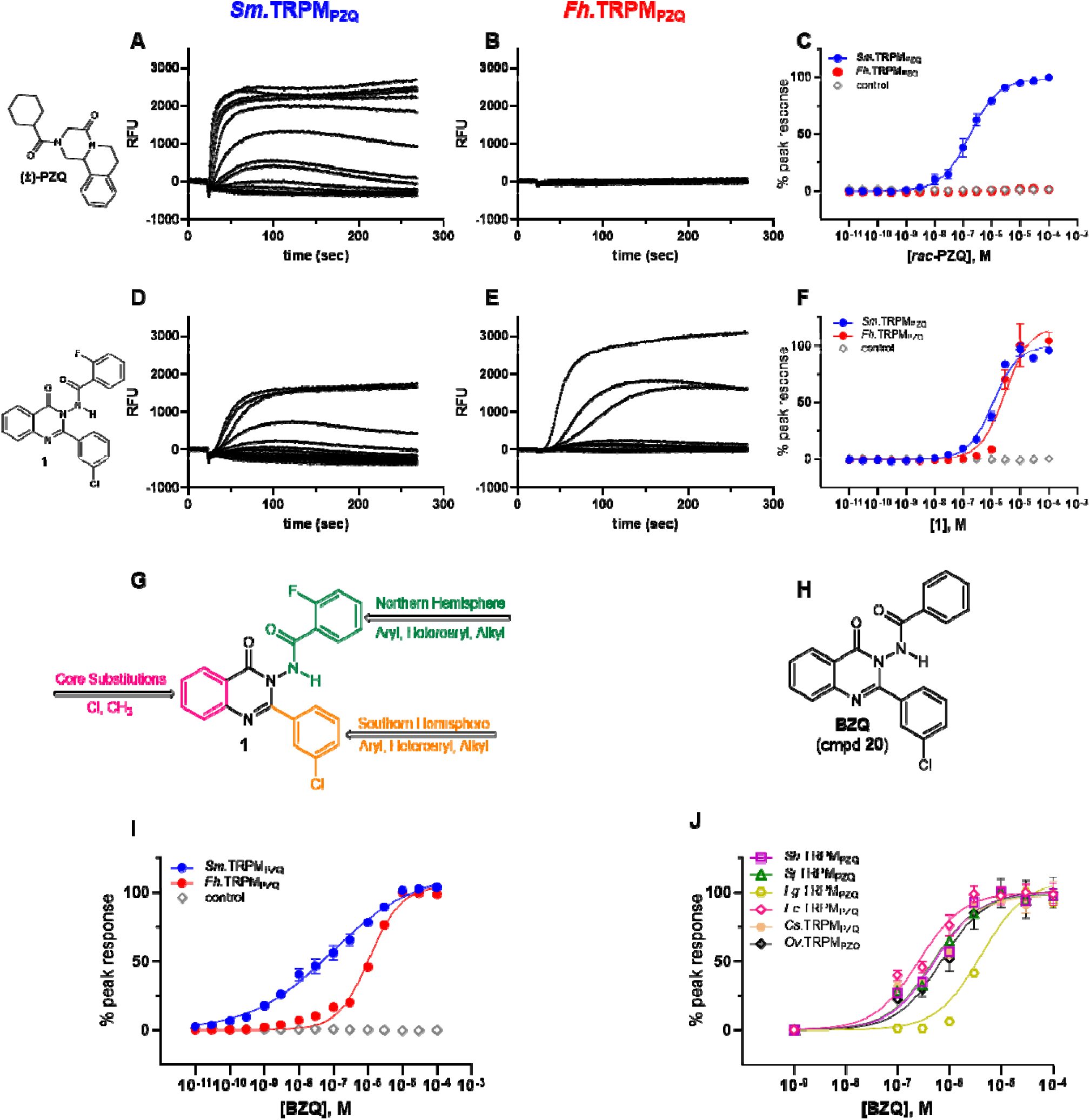
Functional profiling of TRPM_PZQ_ orthologs. Representative Ca^2+^ flux traces depicting the effect of (**A, B**) **(±)-PZQ** or (**D, E**) compound **1** in HEK293 cells stably expressing (A,D) *Sm*.TRPM_PZQ_ or (B,E) *Fh*.TRPM_PZQ_. Cells were treated with increasing concentrations (0-100 µM) of each drug added after ∼20 s of sampling the baseline fluorescence emission. (**C&F**) Concentration-response curves resulting from activation of *Sm*.TRPM_PZQ_ (blue circles) or *Fh*.TRPM_PZQ_ (red circles) by (C) **(±)-PZQ** or (F) compound **1**. Control responses in HEK293 cells lacking TRPM_PZQ_ are shown (grey diamonds). (**G**) Schematic of modifiable regions on compound **1**. Three regions on the *N-*benzamidoquinazolinone core were targeted for modification. The Northern Hemisphere (green), Southern Hemisphere (orange), and the aromatic core (pink). (**H**) Chemical structure of the optimized benzamidoquinazolinone, **BZQ**, after SAR studies. (**I**) Concentration-response curves for **BZQ** in HEK293 cells stably expressing *Sm*.TRPM_PZQ_ (blue circles) or *Fh*.TRPM_PZQ_ (red circles), compared to control responses (grey diamonds). (**J**) Concentration-response curves for **BZQ** in cells transiently expressing various TRPM_PZQ_ orthologs. These were: *Schistosoma mansoni* (*Sm*.TRPM_PZQ_, closed blue circles), *Fasciola hepatica* (*Fh*.TRPM_PZQ_, closed red circles), *Schistosoma haematobium* (*Sh*.TRPM_PZQ_, open purple squares), *Schistosoma japonicum* (*Sj*.TRPM_PZQ_, open green triangles), *Fasciola gigantica* (*Fg*.TRPM_PZQ_, open gold hexagons), *Echinostoma caproni* (*Ec*.TRPM_PZQ_, open grey diamonds), *Clonorchis sinensis* (*Cs*.TRPM_PZQ_), and *Opisthorchis viverrini* (*Ov*.TRPM_PZQ_, open black diamonds). Concentration-response curves were normalized to the maximum response at each channel and represent the mean ± SE of n ≥ 3 independent experiments, each comprised of technical duplicates.

A structure-activity relationship (SAR) analysis was then performed to optimize the activity of the *N*-benzamidoquinazolinone scaffold of (**1**) against TRPM_PZQ_. Three sterically and electronically modifiable regions of (**1**) were interrogated: the Northern Hemisphere, Southern Hemisphere, and the aromatic core (**Figure 1G**).

Beginning with Northern Hemisphere analogs (**Table 1**), alterations at the 2-position of the 2-fluorophenyl ring were evaluated. Replacement of the fluorine with chlorine (**2**), trifluoromethyl (**3**), and nitro (**4**) groups resulted in inactive molecules at both *Sm*.TRPM_PZQ_ and *Fh*.TRPM_PZQ_ (**Table 1**, entries 2-4). A methyl substituent at the 2-position (**5**) was active at both channels, but less potent than (**1**) (Table 1, entry 5). Substitutions at the 3-position of the phenyl ring were however tolerated. A 3-fluorophenyl (**6)**, 3-methylphenyl (**7**) and 3-methoxyphenyl (**8**) analog retained activity at both TRPM_PZQ_ channels, whereas nitration of the 3-position (**9**) ablated activity (Table 1, entries 6-9). Substitution at the 4-position of the aryl ring (4-fluorophenyl (**10**), 4-methylphenyl (**11**), 4-nitrophenyl (**12**), and 4-methoxyphenyl (**13**) analogs) resulted in inactive molecules (Table 1, entries 10-13). An aromatic ring was necessary as cyclohexyl (**14**) and methyl (**15**) analogs were inactive at both TRPM_PZQ_ channels (Table 1, entries 14-15). Replacing the 2-fluorophenyl ring with 2-thiophene (**16**) and 3-thiophene (**17**) analogs also produced molecules with decreased potency compared with **1** (Table 1, entries 16-17), and 2-furyl (**18**) and 4-pyridyl (**19**) analogs were either inactive or much less potent (Table 1, entries 18-19). Collectively, these results underscored stringent structural requirements for efficacy at TRPM_PZQ_.

**Table 1.** SAR of the Northern Hemisphere.

| Entry | Compound | R' | Sm.TRPM<sub>PZQ</sub> (μM) | Fh.TRPM<sub>PZQ</sub> (μM) |
| --- | --- | --- | --- | --- |
| 1 | 1 | 2-F-Ph | 1.15 ± 0.11 | 3.0 ± 0.57 |
| 2 | 2 | 2-Cl-Ph | inactive | inactive |
| 3 | 3 | 2-CF<sub>3</sub>-Ph | inactive | inactive |
| 4 | 4 | 2-NO<sub>2</sub>-Ph | inactive | inactive |
| 5 | 5 | 2-CH<sub>3</sub>-Ph | > 10 | > 10 |
| 6 | 6 | 3-F-Ph | 1.42 ± 0.16 | 6.82 ± 2.51 |
| 7 | 7 | 3-CH<sub>3</sub>-Ph | 4.30 ± 0.80 | 12.1 ± 0.73 |
| 8 | 8 | 3-OCH<sub>3</sub>-Ph | 1.12 ± 0.23 | 3.72 ± 0.44 |
| 9 | 9 | 3-NO<sub>2</sub>-Ph | inactive | inactive |
| 10 | 10 | 4-F-Ph | inactive | inactive |
| 11 | 11 | 4-CH<sub>3</sub>-Ph | inactive | inactive |
| 12 | 12 | 4-NO<sub>2</sub>-Ph | inactive | inactive |
| 13 | 13 | 4-OCH<sub>3</sub>-Ph | inactive | inactive |
| 14 | 14 | Cyclohexyl | inactive | inactive |
| 15 | 15 | Methyl | inactive | inactive |
| 16 | 16 | 2-Thiophene | 5.50 ± 1.00 | 5.50 ± 0.77 |
| 17 | 17 | 3-Thiophene | 2.43 ± 0.25 | >10 |
| 18 | 18 | 2-Furyl | inactive | inactive |
| 19 | 19 | 4-Pyridyl | >50 | >50 |
| 20 | 20 | Phenyl | 0.090 ± 0.02 | 1.08 ± 0.06 |
| 21 | 21 | Bn | inactive | inactive |
| 22 | 22 |  | inactive | inactive |

On the basis of this ‘tight’ SAR, we decreased the sterics around the Northern Hemisphere by removing all substituents from the phenyl ring. This produced compound **20** (**Figure 1H**), a benzamidoquinazolinone that displayed improved potency compared with (**1**) at both *Sm*.TRPM_PZQ_ (EC_50_ = 0.09 ± 0.02 µM) and *Fh*.TRPM_PZQ_ (EC_50_ = 1.08 ± 0.06 µM) (**Table 2**, entry 20 and **Figure 1I**). However, homologating the phenyl ring by one carbon (**21**) yielded an inactive molecule (Table 1, entry 21). Methylating the amide nitrogen also resulted in a molecule (**22)** inactive at both TRPM_PZQ_ channels (Table 1, entry 22), demonstrating the N-***H*** of the benzamide is necessary for channel activation. Ultimately, the analog possessing an unsubstituted phenyl ring (**20**) was the most potent agonist at both *Sm*.TRPM_PZQ_ and *Fh*.TRPM_PZQ_, such that this analog was carried forward as the optimal Northern Hemisphere. Next, SAR of the Southern Hemisphere was investigated (**Supplementary Table 1**). A total of 15 compounds were made but no improvements in potency were identified favoring retention of the 3-chlorophenyl Southern Hemisphere in (**20**). Finally, substitutions around the core ring were profiled (**Supplementary Table 2**). Again, no improvements over (**20**) were identified. A complete summary of concentration-response curves for all 42 molecules synthesized in this SAR campaign is provided in the Supplementary Results (**Supplementary Figures 1-3**).

Considering all these SAR data, compound **20**, named here as **BZQ** (*N*-**<u>b</u>**en**<u>z</u>**amido**<u>q</u>**uinazolinone), emerged as the optimized candidate (Figure 1H). BZQ was a potent activator of *Sm*.TRPM_PZQ_ and *Fh*.TRPM_PZQ_ (Figure 1I). BZQ also displayed activity against other fluke TRPM_PZQ_ orthologs, assayed after transient transfection in HEK293 cells. These TRPM_PZQ_ orthologs encompassed *Schistosoma haematobium* TRPM_PZQ_ (*Sh*.TRPM_PZQ,_ EC_50_ = 0.51 ± 0.07 µM), *Schistosoma japonicum* TRPM_PZQ_ (*Sj*.TRPM_PZQ_, EC_50_ = 0.47 ± 0.05 µM), *Fasciola gigantica* TRPM_PZQ_ (*Fg*.TRPM_PZQ,_ EC_50_ = 4.08 µM), *Echinostoma caproni* TRPM_PZQ_ (*Ec*.TRPM_PZQ_, EC_50_ = 0.25 ± 0.05 µM), *Clonorchis sinensis* TRPM_PZQ_ (*Cs*.TRPM_PZQ_, EC_50_ = 0.47 ± 0.09 µM), and *Opisthorchis viverrini* TRPM_PZQ_ (*Ov*.TRPM_PZQ_, EC_50_ = 0.71 ± 0.19 µM) (**Figure 1J**). **BZQ** therefore acted as a potent, broad spectrum TRPM_PZQ_ agonist active against every fluke TRPM_PZQ_ that was profiled.

Electrophysiological analyses were then executed as an orthogonal assay to validate the action of the benzamidoquinazolinone analogs (**Figure 2**). Analog **1** and **BZQ** were first profiled in whole cell current measurements. Each compound elicited inward currents through both *Sm*.TRPM_PZQ_ or *Fh*.TRPM_PZQ_ that were sensitive to La^3+^ blockade (e.g., *Fh*.TRPM_PZQ_ activated by **1**, **Figure 2A**). Peak currents for both analogs were similar, but **BZQ** was more potent than **1** at both *Sm*.TRPM_PZQ_ (EC_50_ for **BZQ** = 0.11 ± 0.02 µM versus EC_50_ for **1** = 2.2 ± 0.34 µM) and *Fh*.TRPM_PZQ_ (EC_50_ for **BZQ** = 0.27 ± 0.08 µM versus EC_50_ for **1** = 3.08 ± 0.58 µM) (**Figure 2B**). In recordings made from cell-attached patches, **BZQ** evoked single channel activity in cells expressing either *Fh*.TRPM_PZQ_ or *Sm*.TRPM_PZQ_, whereas PZQ only activated *Sm*.TRPM_PZQ_ (**Figure 2C**). The current-voltage relationships for responses to **BZQ** were linear, consistent with activation of a non-voltage dependent current with a conductance of 152 ± 12 pS at *Sm*.TRPM_PZQ_ and 138 ± 4 pS at *Fh*.TRPM_PZQ_ (**Figure 2D**). This compared to a conductance of 116 ± 3 pS evoked by PZQ at *Sm*.TRPM_PZQ_ (Figure 2D). The P_open_ values for *Fh*.TRPM_PZQ_ activated by **BZQ** and for *Sm*.TRPM_PZQ_ activated by either **BZQ** or (±)-**PZQ** were all similar (**Figure 2E**). **BZQ** was therefore confirmed as a potent activator of both *Sm*.TRPM_PZQ_ and *Fh*.TRPM_PZQ_ with BZQ exhibiting properties similar to the actions of (±)-**PZQ** at *Sm*.TRPM_PZQ_.

**Figure 2.**
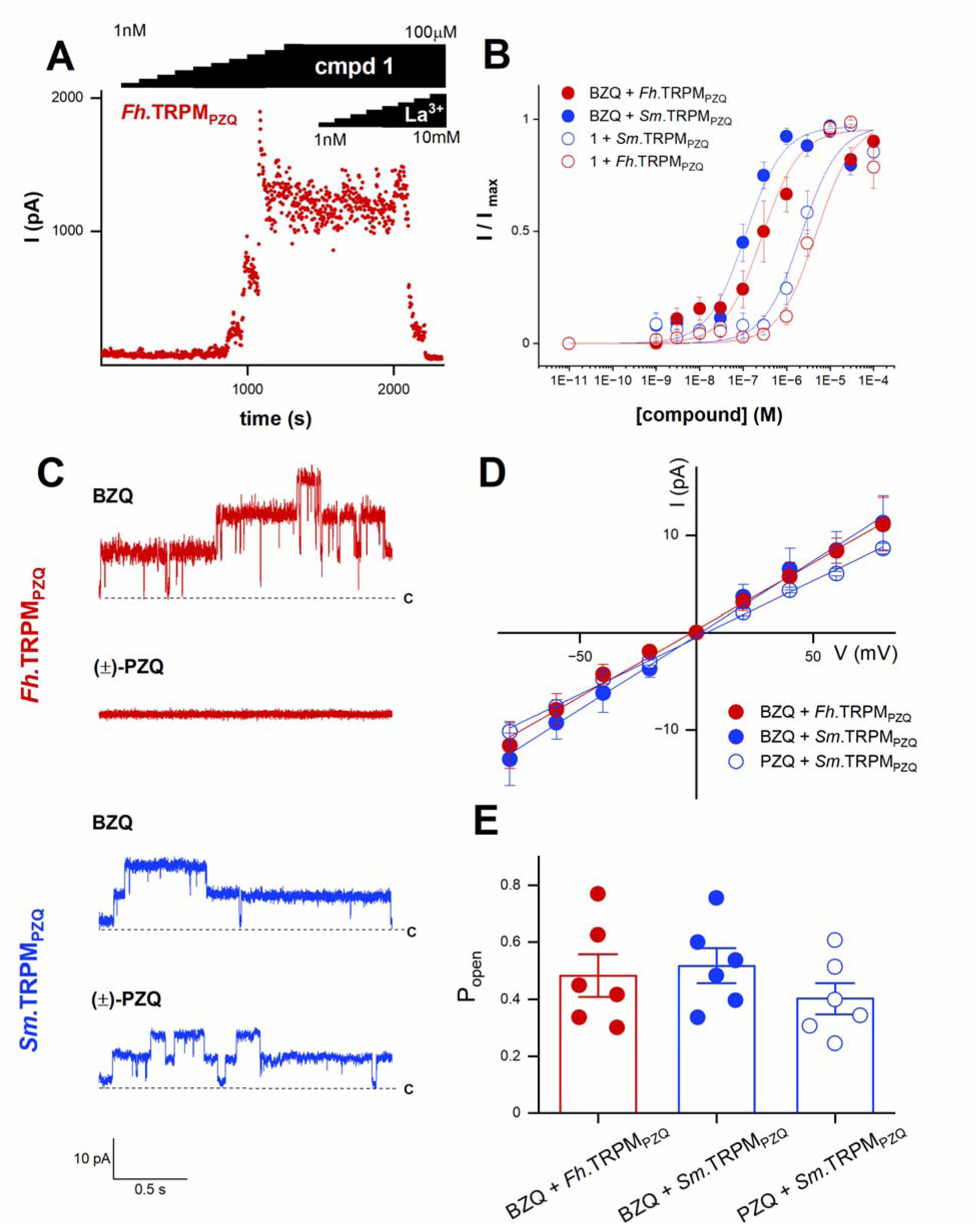
Electrophysiological analysis of BZQ action. (**A**) Whole-cell current (pA) versus time (s) plot of *Fh*.TRPM_PZQ_ expressing HEK293 cell perfused with different concentrations of compound **1** (1 nM to 100 µM) prior to addition of LaCl_3_ (1 nM to 10 mM). Extracellular solution: 140 mM NaCl, 5 mM glucose, 4 mM KCl, 2 mM CaCl*_2_*, 1 mM MgCl*_2_*, pH 7.4 with NaOH. Intracellular solution: 130 mM CsF, 10 mM CsCl, 10 mM NaCl, 10 mM EGTA, 10 mM HEPES, pH 7.4 with NaOH. (**B**) Concentration response curves for BZQ and 1 from experiments such as shown in (A) recorded from *Fh*.TRPM_PZQ_ (red) or *Sm*.TRPM_PZQ_ (blue) expressing HEK293 cells. Data are shown as mean ± SE, n≥6. (**C**) Representative cell-attached recordings from *Fh*.TRPM_PZQ_ or *Sm*.TRPM_PZQ_ expressing HEK293 cells in the presence of (±)-PZQ (10 µM) or BZQ (10 µM) in the bath solution. Bath solution: 145 mM NaCl, 10 mM HEPES, 1 mM EGTA, pH 7.4. Pipette solution: 145 mM NaCl, 10 mM HEPES, 1 mM EGTA, pH 7.4. c, closed state. Holding voltage, 60mV. (**D**) Current-voltage plot for *Sm*.TRPM_PZQ_ (blue) activated by BZQ (closed circle, 10 µM) or (±)-PZQ (open circle, 10 µM)) and *Fh*.TRPM_PZQ_ activated by BZQ (red, 10 µM). Data are shown as mean ± SE, n≥3. (**E**) Single channel open probability (P_open_) of *Fh*.TRPM_PZQ_ (red) or *Sm*.TRPM_PZQ_ (blue) activated by BZQ or (±)-PZQ (each at 10 µM) in the bath solution. Data are shown as mean ± SE, n≥6.

The action of **(±)-PZQ** and **BZQ** was then profiled against *Schistosoma mansoni* and *Fasciola hepatica* flukes *ex vivo*. PZQ is known to cause a rapid, spastic paralysis of schistosomes accompanied with widespread damage to the tegument [13]. Consistent with prior work, **(±)-PZQ** (500 nM) induced a rapid contraction of adult *S. mansoni* worms when compared with vehicle-treated worms (**Figure 3A**). Administration of **BZQ** (500 nM) to *S. mansoni* also caused a rapid, spastic paralysis (Figure 3A). However, treatment of adult *F. hepatica* with **(±)-PZQ** failed to evoke contraction (**Figure 3B**, left), resembling vehicle-treated worms even when applied at high concentration (50 µM). In contrast, **BZQ** (6.25 µM) caused contraction and paralysis of the liver fluke, (Figure 3B), and flukes exposed to **BZQ** did not respond to mechanical stimulation.

**Figure 3.**
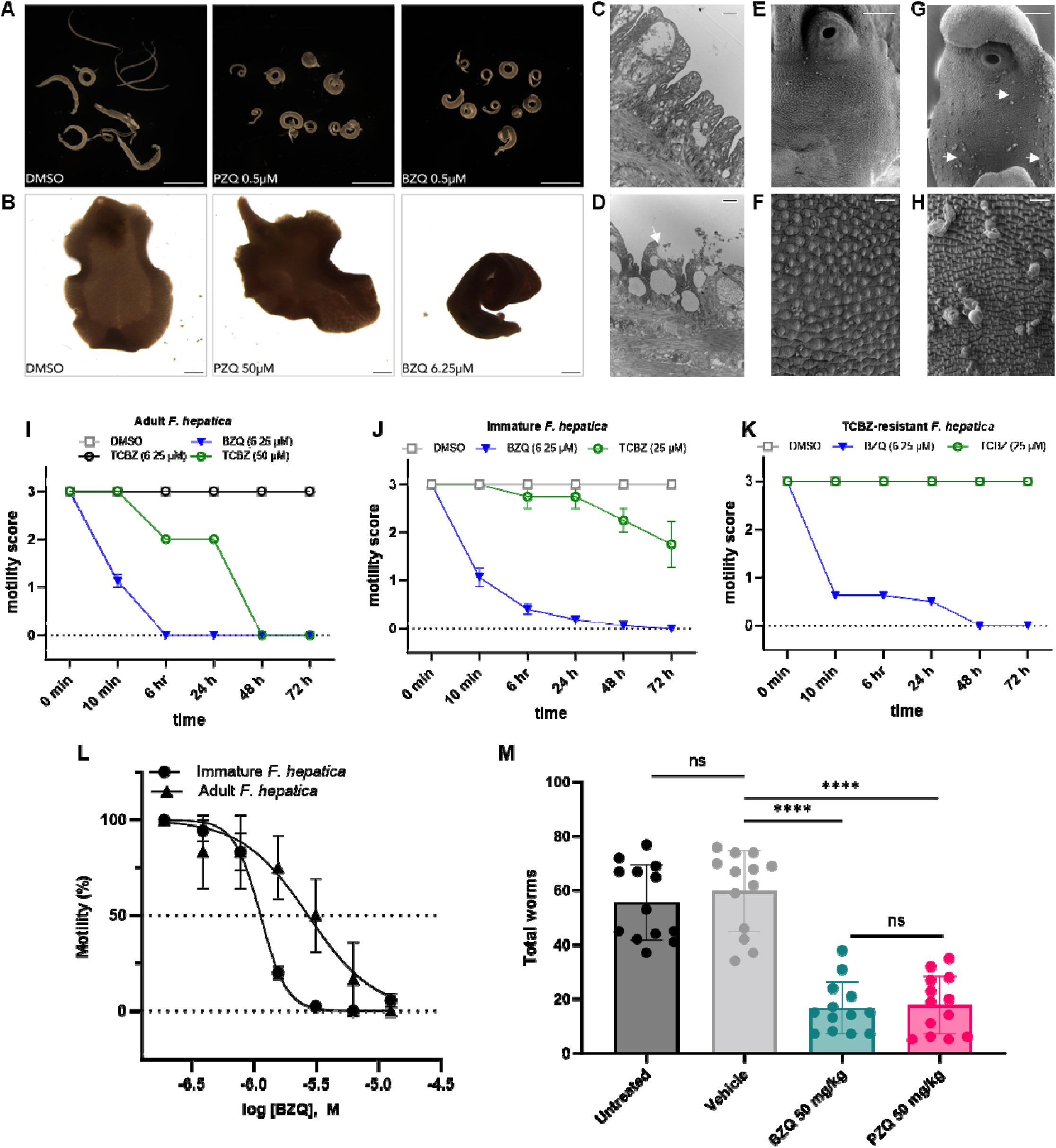
Effects of BZQ and (±)-PZQ on parasitic flukes. (**A&B**) Exposure of *S. mansoni* and *F. hepatica* to **(±)-PZQ** or **BZQ** compared with DMSO (1-1.25%, control). A rapid contraction of schistosomes to **(±)-PZQ** (0.5 µM) or **BZQ** (0.5 µM) was apparent. BZQ (6.25 µM), but not **(±)-**PZQ (50 µM), caused spastic paralysis of adult liver flukes. (**C-H**) Studies of the ultrastructure of **BZQ**-treated flukes. Transmission electron microscopy of drug-induced damage to *S. mansoni* tegument (**C)** without treatment or (**D**) after treatment with **BZQ** (1 μM). Scanning electron microscopy of drug-induced damage to immature *F. hepatica* tegument after treatment with: (**E&F**) DMSO (1.25%, control) or (**G&H**) **BZQ** (6.25 μM, 24 h exposure). BZQ caused blebs to occur on the fluke surface (arrows). (**I-K**) Motility of (**I**) adult, (**J**) triclabendazole (TCBZ)-sensitive immature, and (**K**) TCBZ-resistant immature *F. hepatica* after treatment with **BZQ** (blue triangles) or triclabendazole (black/green circles) compared with application of DMSO (1.25%, control, grey squares). Motility scores are reported as the mean ± SE of n ≥ 3 independent experiments. (**L**) Dose-response curve for motility of adult (triangle) and immature (circle) *F. hepatica* treated with BZQ. **(M) BZQ** activity in a murine model of schistosomiasis. Mice, infected with schistosomes, were treated at 7 weeks post-infection with either **BZQ** or **(±)-PZQ**. Mice were dosed once daily with each drug for three sequential days (50 mg/kg, intraperitoneally), and worm burden was evaluated on the fourth day after initiation of treatment. Worm burden was reduced by treatment with **BZQ** or **(±)-PZQ** compared with either untreated or vehicle-treated mice as described in the methods. N = 13 mice per group; data are shown as mean ± SD and analyzed using the Mann-Whitney test. **** = p ≤ 0.0001, ns = not significant. Scale bars for (**A**) = 250 µm, (**B)** = 1 mm; (**C&D**) = 1 µM, (**E&G**) = 500 µm, (**F&H**) = 10 µm.

Fluke surface ultrastructure was examined after drug exposure. In *S. mansoni*, the integrity of the tegument (**Figure 3C**) was disrupted by **BZQ** (**Figure 3D**). The normal tegumental appearance of *F. hepatica* (**Figures 3E&F**) was also disrupted by exposure to **BZQ** (6.25 µM, 24 h), with widespread bleb formation (**Figures 3G&H**). After 72 hours, these small blebs had fused to form large collapsed blisters on the fluke surface (**Supplementary Figure 4**). Therefore **BZQ,** unlike **(±)-PZQ**, caused muscle contraction and surface damage in *F. hepatica*, mimicking the phenotypes caused by **(±)-PZQ** in schistosomes.

**BZQ** was considerably more potent against liver flukes than the reference drug TCBZ. Parasite motility drastically decreased within 10 min of **BZQ** exposure, while the effects of TCBZ on motility were slower to manifest and required higher drug concentrations (**Figure 3I**). The effects of these treatments on adult *F. hepatica* motility are shown in the Supplementary Videos (**Supplementary Videos 1-4**). In assays using either TCBZ-sensitive (**Figure 3J**) or TCBZ-resistant immature liver flukes (**Figure 3K**), **BZQ** also caused a rapid, protracted paralysis. The IC_50_ of **BZQ** on motility of immature and adult *F. hepatica* was 1.12 ± 0.11 µM and 2.72 ± 0.21 µM respectively (**Figure 3L**). These values were consistent with the potency of **BZQ** at *Fh*.TRPM_PZQ_ (EC_50_ = 1.08 ± 0.06 µM, Figure 1H). Thus, BZQ is highly potent against different *Fasciola* strains and life stages, unlike most other currently available drugs [18]. Similar effects (inhibition of motility, tegument damage) were also seen with other benzamidoquinazolinones shown to be active at *Fh*.TRPM_PZQ_ (**Supplementary Figure 5**).

The activity of **BZQ** was compared with **(±)-PZQ** in a murine model of schistosomiasis. Both molecules were administered at equivalent doses (50 mg/kg, 1x/day, 3 days), and both proved equally effective at reducing worm burden *in vivo* (**Figure 3M**). Untreated mice had a worm burden of 56 ± 14 worms and mice treated with vehicle had a worm burden of 60 ± 15 worms. In contrast, **(±)-PZQ** treated mice had a worm burden of 18 ± 11 worms (a 68% reduction), and **BZQ** treated mice had a worm burden of 17 ± 10 worms (a 70% reduction). These residual worms recovered from **BZQ** injected mice were contracted and immobile. Thus, the activity of **BZQ** was apparent at the target (Figures 1&2), and against worms assayed *ex vivo* (Figures 3A&B) or *in vivo* (Figure 3M) validating **BZQ** as a highly promising anthelmintic candidate.

How does **BZQ** activate both *Sm*.TRPM_PZQ_ and *Fh*.TRPM_PZQ_? PZQ activates *Sm*.TRPM_PZQ_ via engagement of a binding pocket at the base of the voltage sensing-like domain (VSLD) of the ion channel [15]. This binding pocket is framed by transmembrane helices S1-S4, and the cytoplasmic TRP helix (**Figure 4A**). Three groups of interactions have been shown to be essential for PZQ activity. First, an interaction between the S1 helix and the internal carbonyl of PZQ (**Figure 4B**). Second, an interaction between the S4 helix (R1514 in S4) and the external carbonyl of PZQ (Figure 4B). Finally, interactions between PZQ and the TRP domain (Y1678 and R1681 form additional hydrogen bonds with the internal carbonyl of PZQ, Figure 4B).

**Figure 4.**
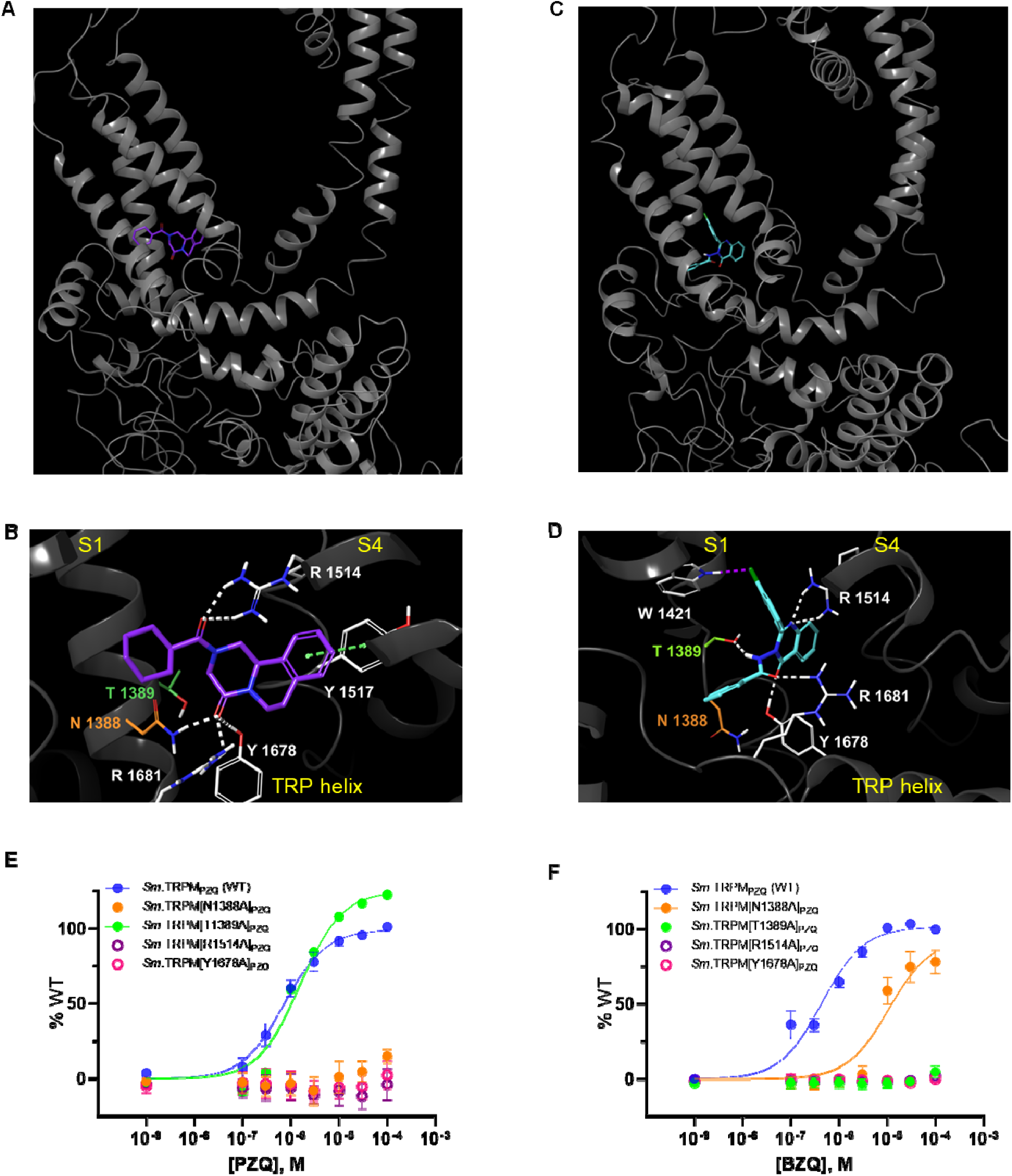
TRPM_PZQ_ engagement by BZQ. *In silico* binding pose for (**A&B**) **(*R*)-PZQ** and (**C&D**) **BZQ** in *Sm*.TRPM_PZQ_. Concentration-response curves for (**E**) **(±)-PZQ** and (**F**) **BZQ** in specified *Sm*.TRPM_PZQ_ mutants. WT = blue circles, *Sm*.TRPM[N1388]A_PZQ_ = orange circles, *Sm*.TRPM[T1389A]_PZQ_ = green circles, *Sm*.TRPM[R1514A]_PZQ_ = open purple circles, *Sm*.TRPM[Y1678A]_PZQ_ = open pink circles. Data are presented as mean ± SEM of biological triplicates performed in technical duplicate.

Computational modeling was applied to understand how **BZQ** engages *Fh*.TRPM_PZQ_. Induced-fit docking (IFD) resulted in a pose of **BZQ** within the VSLD of *Sm*.TRPM_PZQ_ (**Figure 4C**). The VSLD binding pocket of *Fh*.TRPM_PZQ_ was well conserved compared with *Sm*.TRPM_PZQ_. Of the 23 amino acids that lie within 5Å of PZQ binding pose in *Sm*.TRPM_PZQ_, 22 of them were conserved in *Fh*.TRPM_PZQ_ [15]. The exception was a threonine residue (T1270, S1) in *Fh*.TRPM_PZQ_, in place of an asparagine residue (N1388, S1) that was predicted to form a critical hydrogen bond between PZQ and *Sm*.TRPM_PZQ_. This natural variation underpins the inactivity of PZQ at *Fh*.TRPM_PZQ_ [15, 16]. The majority of key interactions seen in the **PZQ** binding pose (R1514, Y1678, and R1681) were retained for **BZQ** in *Sm*.TRPM_PZQ_ (**Figure 4D**). The difference was that **BZQ** was not predicted to interact with N1388, the variant S1 residue between the *Sm*.TRPM_PZQ_ and *Fh*.TRPM_PZQ_ VSLD binding pockets. Instead, the amide N-***H*** of **BZQ** was predicted to form a hydrogen bond with the oxygen of the adjacent threonine (T1389), one residue further along the S1 helix (Figure 4D).

To experimentally interrogate the **BZQ** binding pose prediction, point mutations were generated and profiled in Ca^2+^ reporter assays. Whereas PZQ activation of *Sm*.TRPM_PZQ_ was tolerant of mutation of this threonine residue (*Sm*.TRPM[T1389A], **Figure 4E**), BZQ activation was not (*Sm*.TRPM[T1389A], **Figure 4F**). In contrast, PZQ activation of *Sm*.TRPM_PZQ_ required the S1 asparagine (*Sm*.TRPM[N1388A]_PZQ_, Figure 4E), whereas BZQ did not (*Sm*.TRPM[N1388A]_PZQ_, Figure 4F). However, for both ligands, R1514A (S4) and Y1678A (TRP) mutations abolished activity (Figure 4E&F), consistent with the shared interactions predicted in the computational model (Figure 4B&D). Therefore, **BZQ** reproduced the same interactions as **(*R*)-PZQ** with the S4 helix and the TRP domain of TRPM_PZQ_ but unlike **(*R*)-PZQ** the interaction with S1 was predicted to utilize a conserved threonine residue present in both *Sm*.TRPM_PZQ_ and *Fh*.TRPM_PZQ_. These **BZQ**-interacting residues are retained across all fluke TRPM_PZQ_ orthologs explaining the broad-spectrum activity of **BZQ**.

## Discussion

Development of new leads to counter neglected tropical diseases is an urgent priority, as current therapeutic portfolios are limited and, for many diseases, have remained unchanged for decades. Fasciolosis provides an example of an infectious disease where new drugs would be valuable from both a clinical and veterinary perspective, given growing resistance to triclabendazole and the ineffectiveness of praziquantel. Here, we have identified the benzamidoquinazolinone core and optimized a ligand, **BZQ**, that displays efficacy against *Fasciola hepatica* comparable with the action of PZQ against other trematodes.

**BZQ** was discovered by target-based screening of TRPM_PZQ_, the parasitic flatworm target of **(±)-PZQ** [12]. BZQ activated *Sm*.TRPM_PZQ_, *Fh*.TRPM_PZQ_, as well as TRPM_PZQ_ orthologs from all other flukes tested (Figure 1). Electrophysiology studies validated BZQ activity in an orthogonal assay (Figure 2), and BZQ mimicked the action of PZQ on parasitic flatworms (Figure 3). That a molecule discovered from target-based screening phenocopied PZQ activity on *Fasciola spp.* provide further support for correct validation of TRPM_PZQ_ as the relevant target of PZQ.

Discovery of **BZQ** also validates TRPM_PZQ_ as a druggable target, although the SAR tolerated within the VSLD cavity of TRPM_PZQ_ remained stringent. The majority of derivatives of **BZQ** (Table 1, Supplementary Tables 1&2) were inactive or poorly active, matching conclusions from prior SAR studies with praziquantel [15]. **(±)-PZQ** and **BZQ** retain broadly similar properties (**Supplementary Table 3**). Both ligands are tetracyclic bis(amides), with their hydrophobic extremities linked by a polar midriff. Both ligands are similarly hydrophobic, and both present a similar polar surface area relative to their molecular weight. This stringency likely reflects requirements for complementarity within the hydrophobic VSLD binding pocket. Quinazolinones have broadly reported biological activities, including antiparasitic [19] and antiviral activities [20]. Quinazolinone-type benzamides have been studied from a synthetic standpoint on account of their chirality and chiroptic properties [21, 22]. However, **BZQ**, with the *N-*benzamidoquinazolinone scaffold harboring both 2- and 4-(N-N bond)substitutions, is a novel molecule that has not been previously characterized.

Activity against *Fasciola spp.* was a result of **BZQ** binding to *Fh.*TRPM_PZQ_ in a manner tolerant of the sequence variation present within the VSLD cavity of these particular liver flukes, that renders PZQ ineffective [15, 16]. Comparison of the binding pose of **(*R*)-PZQ** and **BZQ** in *Sm*.TRPM_PZQ_ highlights the general principles underpinning TRPM_PZQ_ activation. Both chemotypes exhibit interactions with S4 (R1514) and the TRP helix (R1681, Y1678) that surround the VSLD binding pocket (**Figure 4**). These interactions are with residues conserved with the human TRPM8 binding pocket (R1514 ∼R842 in hTRPM8, R1681 ∼R1008 in hTRPM8, Y1678 ∼Y1005 in hTRPM8) that are known to display mobility in their sidechain configuration permitting engagement of different hTRPM8 chemotypes [23]. While these S4 and TRP domain residue interactions are identical for both **(*R*)-PZQ** and **BZQ**, the site of S1 engagement differed – such that **BZQ** bypassed the need to interact with the variant residue (**Figure 4**). Instead, **BZQ** is predicted to interact with a conserved S1 threonine residue that is present in all fluke TRPM_PZQ_ orthologs. The interaction of the amide nitrogen of **BZQ** with the threonine oxygen in both orthologs is consistent with the SAR analysis where methylation of the amide resulted in **22**, a molecule that failed to activate *Sm*.TRPM_PZQ_ or *Fh*.TRPM_PZQ_ (**Table 1**). **BZQ** therefore not only overcomes the lack of efficacy of PZQ against *Fasciola spp.*, but provides higher efficacy than the current gold standard TCBZ against both TCBZ-susceptible and TCBZ-resistant parasite strains. BZQ also displays *in vivo* antischistosomal activity comparable to PZQ. This establishes **BZQ** as a broad-spectrum flukicide and a promising anthelmintic lead for further development.

## Materials and Methods

### Materials & Reagents

(±)-PZQ was purchased from Sigma. All cell culture reagents were from ThermoFisher. Synthetic procedures and characterization data are detailed in the Supplementary Information.

### Cell culture

HEK-293 cell lines were sourced from ATCC (CRL-1573) and authenticated by STR profiling (ATCC). Cells were screened negative for mycoplasma contamination by monthly scheduled testing (LookOut^®^ Mycoplasma PCR Detection Kit, Sigma). Stable cell lines expressing *Sm*.TRPM_PZQ_ or *Fh*.TRPM_PZQ_ were generated using the Flp-In T-REX core kit (Invitrogen) as follows. Flp-In T-REx 293 cells were co-transfected with pOG44 (Invitrogen) and a TRPM_PZQ_ expression plasmid (*Sm*.TRPM_PZQ_ or *Fh*.TRPM_PZQ_) containing a Flp recombination target (FRT) site (pcDNA5/FRT) at a 3:1 ratio using Lipofectamine 2000 (Invitrogen). Two days following transfection, cells were trypsinized and seeded into 100 mm dishes and selection was initiated with 10 μg/ml blasticidin (Invivogen) and 200 μg/ml hygromycin B (Invitrogen) for a period of 7-10 days. Single colonies were isolated, and the expression of *Sm*.TRPM_PZQ_ or *Fh*.TRPM_PZQ_ compared using the FLIPR Ca^2+^ assay. Clones exhibiting an optimal signal to noise were prioritized for experiments. Prior to assays, stable cells expressing *Sm*.TRPM_PZQ_ or *Fh*.TRPM_PZQ_ were induced by addition of tetracycline (2 μg/ml, 24 h). Transient transfections were performed using Lipofectamine 2000 as reported previously [15].

### FLIPR Ca^2+^ assay

The Fluorescence Imaging Plate Reader (FLIPR) Ca^2+^ reporter assay was performed in black-walled, clear-bottomed 384-well plates coated with poly-D-lysine (Greiner Bio-One, Germany). Briefly, non-transfected or transfected HEK293 cells were seeded (20,000 cells/well) in DMEM growth media containing 10% FBS. After 24 hours, medium was removed, and replaced with 20 µl of Fluo-4 NW dye loading solution (Molecular Devices), previously reconstituted in assay buffer (Hanks’ balanced salt solution with Ca^2+^, Mg^2+^, 20 mM HEPES and 2.5 mM probenecid). Cells were incubated for 30 min at 37°C (5% CO_2_) followed by an additional 30 min incubation at room temperature. Drug dilutions were prepared in assay buffer, but without probenecid and fluorescent dye, in 384-well plates (Greiner Bio-One). Using a FLIPR^TETRA^ (Molecular Devices), basal fluorescence (filter settings λ_ex_=470-495 nm, λ_em_=515-575 nm) from each well was monitored for 20 s, then 5 μl of drug or vehicle solution was added (25 μl total volume) and the signal was recorded over 250 s. Changes in fluorescence were represented as relative fluorescence units after subtracting the average basal fluorescence (average basal fluorescence over 20 s) from the recorded values. Concentration-response analysis was performed using sigmoidal curve fitting functions in Prism using data from n ≥ 3 independent transfections, with n ≥ 3 technical replicates per assay.

### Electrophysiology

For whole cell current measurements, assays were performed using the Patchliner automated patch-clamp system (Nanion, Germany). HEK293 cells stably expressing *Sm*.TRPM_PZQ_ or *Fh*.TRPM_PZQ_ were grown in a T25 flask and harvested at 60-80% confluency using Accutase (ThermoFisher) (∼0.5 ml per T25 flask, treated for 5 mins). The suspension was diluted with 3 ml of extracellular buffer (140 mM NaCl, 5 mM glucose, 4 mM KCl, 2 mM CaCl_2_, 1 mM MgCl_2_, pH 7.4) and the resulting suspension used for assays. Cells were trapped on the assay chip (NPC–16, medium resistance) using negative pressure, and tight contact was achieved using a seal enhancement solution (130 mM NaCl, 5 mM glucose, 4 mM KCl, 10 mM CaCl_2_, 1 mM MgCl_2_, pH 7.4). After a gigaohm seal was obtained, a vacuum pulse was sent to the chip to obtain a whole-cell patch. Recordings were made using an intracellular buffer of 130 mM CsF, 10 mM CsCl, 10 mM NaCl, 10 mM EGTA, 10 mM HEPES, pH 7.4. Compounds were perfused through the microfluidic system at various concentrations prior to perfusion with La^3+^ (1 nM to 10 mM) to follow current inactivation at the end of each assay. For single-channel recordings, HEK293 cells expressing *Sm*.TRPM_PZQ_ or *Fh*.TRPM_PZQ_ were placed in symmetrical buffer (145 mM NaCl, 10 mM HEPES, 1 mM EGTA, pH 7.4) and treated with compound **1** (10 µM), **BZQ** (10 µM), or **(±)-PZQ** (10 µM).

### Computational Procedures

Modeling was performed in the Schrodinger Computational Suite (v2022-4 or v2023-1), using the Maestro GUI (v13.1). All modeling was performed with default settings unless otherwise noted. Generation of the homology model of *Sm*.TRPM_PZQ_, along with the complex with (*R*)-PZQ, has been previously described and validated [15, 16]. To prepare BZQ for modeling, the molecule was drawn in ChemDraw Professional (v21.0.0), imported into the Maestro GUI, and minimized using the LigPrep tool in the OPLS4 force field at pH=7.4. The output structure was used for subsequent studies. Induced-Fit Docking (IFD), a model that employs flexibility of both ligand and protein in the docking procedure, was performed in an iterative fashion. With a grid generated around (*R*)-PZQ in the *Sm.*TRPM_PZQ_ homology model, IFD of BZQ was performed with both the channel and ligand van der Waals scaling set to 0.30. Standard precision (SP) settings were used for initial redocking, and residues were optimized within 8.0 Å of the poses. Poses were manually examined, and the highest-ranking pose that displayed interactions consistent with functional data was prioritized. From this pose, model refinement was performed with IFD with default scaling settings and using the XP protocol, a more precise algorithm, for glide redocking. This resulted in the reproducibly stable poses depicted in Figure 4.

### Analysis of the effects of drugs on parasitic flukes

For *ex vivo* drug screening experiments using *F. hepatica*, liver flukes were obtained from male Wistar rats RjHan:WI (Janvier, France) experimentally infected with 20-25 metacercariae of an Italian strain (Ridgeway Research, UK). Immature flukes were collected from livers at 4 weeks post-infection (p.i.), and adult flukes from bile ducts at 12 weeks p.i. Animal experiments were in accordance with Directive 2010/63/EU on the protection of animals used for scientific purposes and the German Animal Welfare Act. The experiments were approved by the Regional Council (Regierungspraesidium) Giessen (V54-19c20 15 h 02 GI 18/10 Nr. A16/2018). Anthelminthic activity of benzamidoquinazolinone analogs (0.19-12.5 µM) was assessed *in vitro* by culturing worms in RPMI medium (supplemented with 5% chicken serum, 1% ABAM-solution (10,000 units penicillin, 10 mg streptomycin and 25 mg amphotericin B per ml), all from Gibco) for up to 72 h at 37°C in a 5% CO_2_ atmosphere. Triclabendazole (25-50 µM) was used as the positive control, and DMSO as the solvent control. Medium and compounds were refreshed every 24 h, and inhibitor-induced effects on worm motility assessed using a stereo microscope at 10x magnification (M125 C, Leica, Germany) using the following scores: 3 (normal motility), 2 (reduced motility), 1 (weak and sporadic movements), 0.5 (minimal movement only upon mechanical stimulation) and 0 (dead).

For *ex vivo* drug screening against *S. mansoni*, adult worms were isolated as described [24]. Harvested schistosomes were washed in DMEM high glucose medium supplemented with HEPES (25mM), pyruvate and 5% heat inactivated FBS (Gibco) and penicillin-streptomycin (100 units/mL) and incubated overnight (37°C/5% CO_2_) in vented petri dishes (100×25mm). For movement analysis, assays were performed using 3 male worms per well in a six well dish. Video recordings were captured using a Zeiss Discovery v20 stereomicroscope with a QiCAM 12-bit cooled color CCD camera controlled by Image-Pro imaging software (*v.* 11). Recordings (60 seconds) of worm motility (1 image every 4 secs), before and after addition of different drugs were analyzed as described previously [15]. For *in vivo* drug screening, female Swiss Webster mice (infected with *S. mansoni* cercariae (NMRI strain) at between 4-6 weeks old) were obtained from the Schistosomiasis Resource Center at the Biomedical Research Institute (Rockville, MD) under contract HHSN272201000005I for distribution via BEI Resources. At 7 weeks post-infection, mice were randomly sorted into 4 groups of 13 individuals for drug efficacy assays. One group was left untreated as an infected control. Experimental groups were treated with either vehicle, praziquantel (50 mg/kg, intraperitoneal, 1x daily), or BZQ (50 mg/kg, intraperitoneal, 1x daily) for 3 sequential days. These drug solutions were prepared fresh on every day of treatment. In the following order, the solid was solubilized in 100% DMSO (15 µL), diluted with PEG_400_ (70 µL), and vortexed for 1 min. Phosphate-buffered saline (50 µL, containing 5% w/v Trappsol^®^) was added 10 µL at a time, with extensive vortexing between each addition to ensure solubilization. These drug solutions were then used within 4 hours of preparation. If any precipitation was observed upon standing, gentle warming in a heating block resolubilized the compound. On the day after the third dose, mice were euthanized by intraperitoneal injection of a pentobarbital solution, and the liver and mesenteric vasculature was dissected and perfused to score worm burden. All animal experiments for schistosome harvest followed ethical regulations approved by the MCW IACUC committee.

### Electron microscopy

Schistosomes were fixed in a mixture of glutaraldehyde (2.5%) and paraformaldehyde (2%) in sodium cacodylate buffer (100 mM) for 3 h at ambient temperature [25, 26]. The fixed worms were then washed with cacodylate buffer to remove fixing solution and then post-fixed in aqueous osmium tetroxide (1%) for 1 h on ice. Schistosomes were processed through a graded methanol series to 100% methanol and then acetonitrile before being infiltrated with epoxy resin overnight at 4°C followed by polymerization at 70°C for 8 h [26]. Polymerized resin blocks were sectioned (60 nm thickness, RMC PTXL ultramicrotome). Sections were stained with uranyl acetate and lead citrate and viewed under a JEOL 1400 Flash TEM. Images were captured on Hamamatsu digital camera running AMT imaging software (v 7.0.1.422).

Immature *F. hepatica* flukes were prepared for scanning electron microscopy by fixation in 100 mM cacodylate buffer containing 2.5% (v/v) glutaraldehyde and 1% (v/v) formaldehyde for 24 h at 4°C. Fixed samples were stored in 0.1% (v/v) formaldehyde in cacodylate buffer before postfixation in 1% (v/v) osmium tetroxide in 100 mM cacodylate buffer for 1 h at ambient temperature. After rinsing twice with ultra-pure water, samples were dehydrated through a graded ethanol series on ice to 100% ethanol. Samples were dried in a CPD030 critical point dryer (BAL-TEC AG; Balzers, Liechtenstein) and coated with gold transferred in a SCD004 sputter system (BAL-TEC AG). Images were captured with a Gemini DSM 982 (Carl Zeiss Microscopy; Oberkochen, Germany), operated at 3 kV.

## Supporting information

Supplementary Video 1. Vehicle-treated (DMSO, 1.25%) adult F. hepatica.

## Acknowledgements.

Funding was provided by the National Institutes of Health (R01 AI155405 to JSM), a Catalyst Award from the Falk Medical Research Trust (to JSM), the German Research Foundation (HA 6963/2-1 to SH) and the Hessen State Ministry of Higher Education, Research, and the Arts (HMWK) (DRUID-C6 to SH). DJS acknowledges support from the NIH (T32 HL134643) and the MCW Cardiovascular Center’s A.O. Smith Fellowship Scholars Program. Screening utilized equipment procured through a NIH equipment grant (1S10OD032248-01 to TPS & LS). We acknowledge the Research Computing Center at the Medical College of Wisconsin, Paul Kerber and Dr. Francis Peterson for maintaining the MCW NMR facilities, the Indiana University Mass Spectrometry Center for HRMS analysis, the MCW Electron Microscopy Core, and the Imaging Unit of the Biomedical Research Center Seltersberg.

## Competing Interests

JSM, DJS, LS, and TPS have pending patent applications for the compounds described in this study.

## Notes

### Competing Interest Statement

The authors have declared no competing interest.

