## Supplementary Information for "Target-based discovery of a broad spectrum flukicide"

**SI-X**

### Supplementary Results

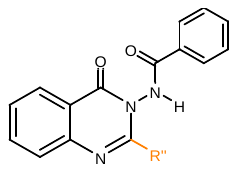

| **Entry** | **Compound** | **R’’** | ***Sm*.TRPM_PZQ_ (µM)** | ***Fh*.TRPM_PZQ_ (µM)** |
| --- | --- | --- | --- | --- |
| **1** | **20** | 3-Cl-Ph | 0.090 ± 0.02 | 1.08 ± 0.06 |
| **2** | **23** | 4-Cl-Ph | *inactive* | *inactive* |
| **3** | **24** | 4-OCH_3_-Ph | *inactive* | *inactive* |
| **4** | **25** | 2-CF_3_-Ph | *inactive* | *inactive* |
| **5** | **26** | 3-OCH_3_-Ph | *inactive* | *inactive* |
| **6** | **27** | 3-OCF_3_-Ph | *inactive* | *inactive* |
| **7** | **28** | 3-NO_2_-Ph | *inactive* | *inactive* |
| **8** | **29** | 3-CF_3_-Ph | *inactive* | *inactive* |
| **9** | **30** | 3-CH_3_-Ph | 0.16 ± 0.03 | 4.39 ± 0.93 |
| **10** | **31** | Cyclohexyl | *inactive* | *inactive* |
| **11^a^** | **32** | Methyl | *inactive* | *inactive* |
| **12** | **33** | 3-Cl-Bn | *inactive* | *inactive* |
| **13** | **34** | 2-Thiophene | 0.90 ± 0.08 | >50 |
| **14** | **35** | 3-Thiophene | 1.03 ± 0.11 | 8.5 ± 0.68 |
| **15** | **36** | 2-Furyl | *inactive* | *inactive* |
| **16** | **37** | Ph | 3.20 ± 0.53 | 26.7 ± 5.3 |

Supplementary Table 1. SAR of the Southern Hemisphere. ^a^This molecule was synthesized with the 2-fluorophenyl Northern Hemisphere rather than the phenyl Northern Hemisphere, and thus can be directly compared with compound **1**.

**Supplementary Table 2**

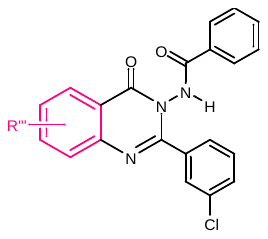

| **Entry** | **Compound** | **R’’’** | ***Sm*.TRPM_PZQ_ (µM)** | ***Fh*.TRPM_PZQ_ (µM)** |
| --- | --- | --- | --- | --- |
| **1** | **38** | 5-CH_3_ | *inactive* | *inactive* |
| **2** | **39** | 6-Cl | *inactive* | *inactive* |
| **3** | **40** | 6-CH_3_ | *inactive* | *inactive* |
| **4** | **41** | 8-Cl | >100 | >100 |
| **5** | **42** | 8-CH_3_ | >100 | >100 |

### Supplementary Table 2. **SAR of the Aromatic Core**.

**Supplementary Figure 1**

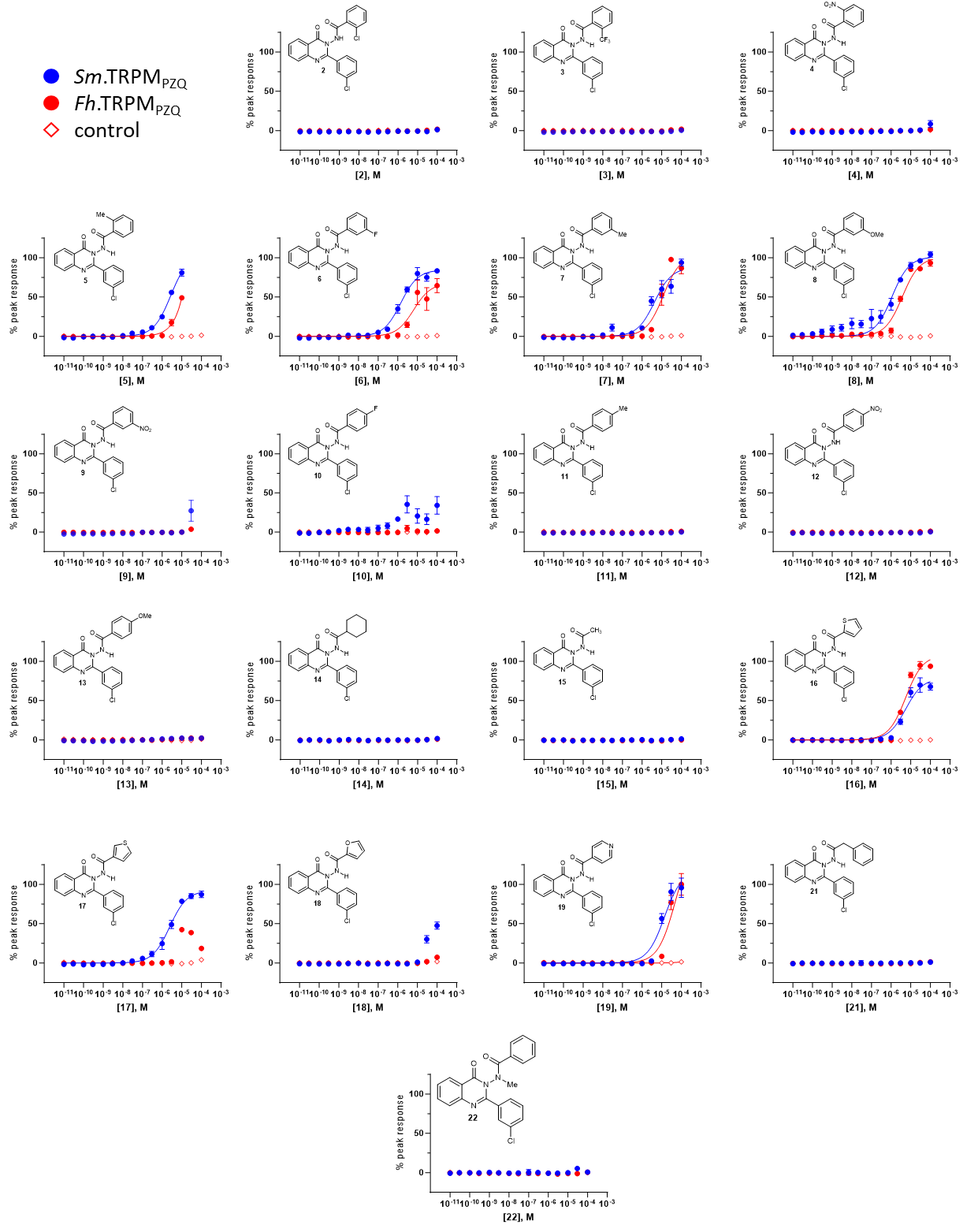

Supplementary Figure 1. Dose-response curves of Different Northern Hemisphere Analogs**.** Responses of *Sm*.TRPM_PZQ_ to the molecules are shown in blue circles, *Fh*.TRPM_PZQ_ in red circles, and control (untransfected cells) red diamonds.

**Supplementary Figure 2**

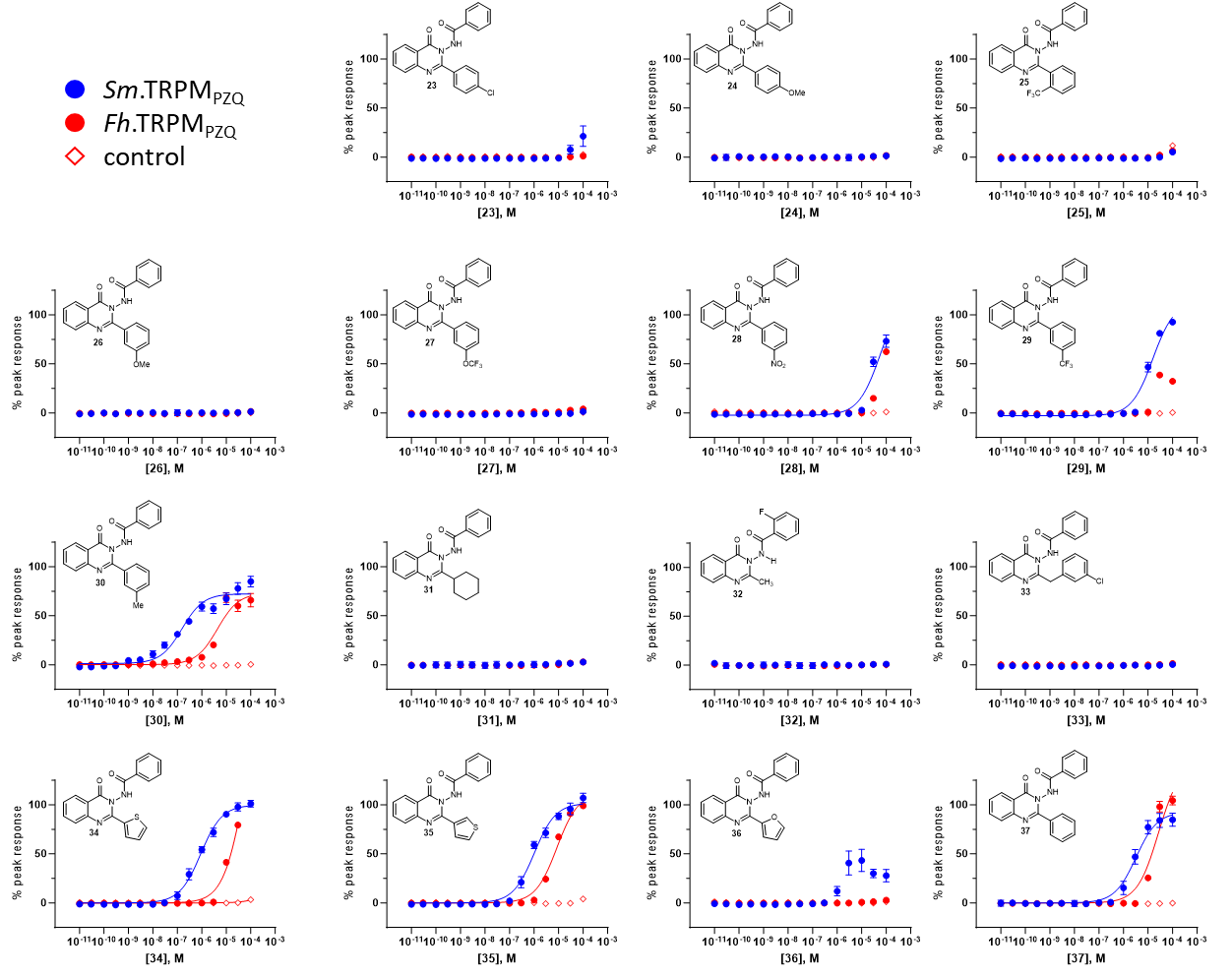

Supplementary Figure 2. Dose-response curves of Different Southern Hemisphere Analogs. Responses of *Sm*.TRPM_PZQ_ to the molecules are shown in blue circles, *Fh*.TRPM_PZQ_ in red circles, and control (untransfected cells) red diamonds.

**Supplementary Figure 3**

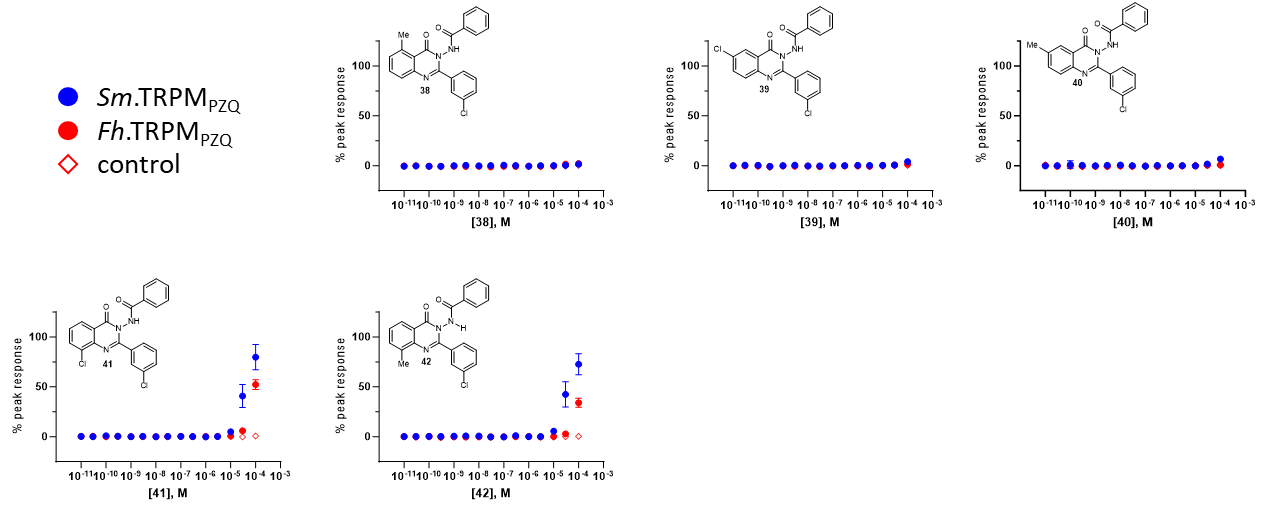

Supplementary Figure 3. Dose-response curves of Different Aromatic Core Analogs. Responses of *Sm*.TRPM_PZQ_ to the drugs are shown in blue circles, *Fh*.TRPM_PZQ_ in red circles, and control (untransfected cells) red diamonds.

**Supplementary Figure 4**

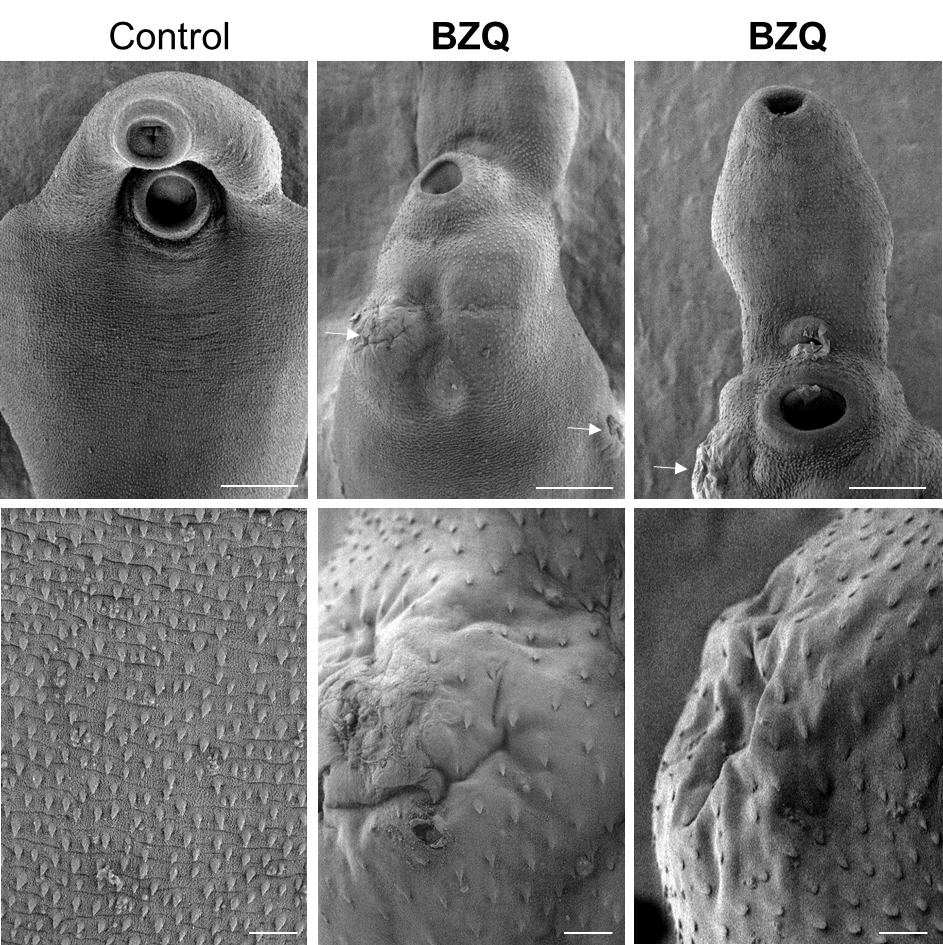

Supplementary Figure 4. Effects of BZQ on *F. hepatica* after prolonged culture. Tegumental damage in *F. hepatica* after extended treatment with **BZQ**. Immature liver flukes were treated for 72 h with **BZQ** (12.5 µM) or DMSO (0.5%) as control and ventral surfaces imaged by scanning electron microscopy. Large collapsed blisters (arrows) were found after exposure (two identical replicates are shown). Scale bars = 500 µm (top row) and 10 µm (bottom row).

**Supplementary Figure 5**

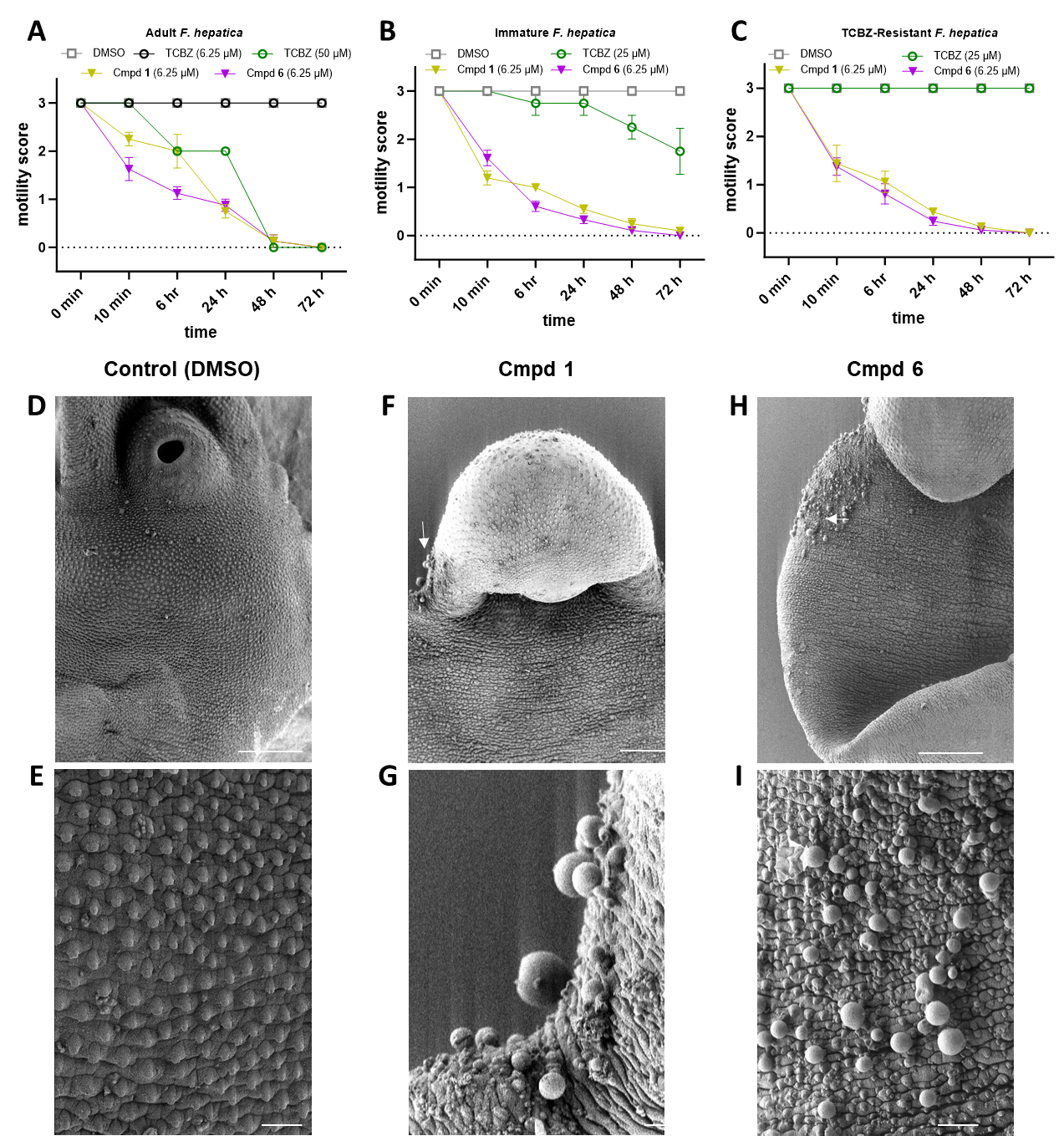

Supplementary Figure 5. Effects of various benzamidoquinazolinones on *F. hepatica*. (**A-C**) Motility of (**A**) adult, (**B**) triclabendazole (TCBZ)-sensitive immature, and (**C**) TCBZ-resistant immature *F. hepatica* after treatment with compound **1** (gold triangles) and compound **6** (purple triangles) compared with application of TCBZ (open circles) or DMSO (1.25%, open grey squares). (**D-I**) Tegumental damage in *F. hepatica*. Immature liver flukes were treated for 24 h with DMSO (control, **D&E**), Cmpd **1** (6.25 µM**,** **F&G**) or compound **6** (6.25 µM, **H&I**). **1** and **6** caused bleb formation (arrows) after exposure. Images are representative of three replicates. Scale bars = 500 µm (top row) and 10 µm (bottom row).

**Supplementary Table 3**

|  | 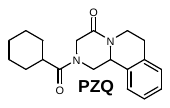 | 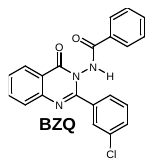 |
| --- | --- | --- |
| **Formula** | C_19_H_24_N_2_O_2_ | C_21_H_14_ClN_3_O_2_ |
| **MW** | 312.41 | 375.81 |
| **cLogP** | 3.36 | 3.77 |
| **TPSA** | 40.6 Å | 64.0 Å |
| **EC_50_ *(Sm*.TRPM_PZQ_)** | 180 ± 20 nM | 90 ± 20 nM |
| **EC_50_ (*Fh*.TRPM_PZQ_)** | no activity | 1080 ± 60 nM |

### Supplementary Table 3. Comparison of TRPM_PZQ_ agonists.

### **Supplementary Video Descriptions**

### **Supplementary Video 1. Vehicle-treated (DMSO, 1.25%) adult *F. hepatica***.

### **Supplementary Video 2. (±)-PZQ-treated (50 µM) adult *F. hepatica***.

### **Supplementary Video 3. BZQ-treated (6.25 µM) adult *F. hepatica***.

### **Supplementary Video 4. TCBZ-treated (50 µM) adult *F. hepatica***.

### Synthetic Chemistry Procedures

### General Remarks

All reagents and solvents were commercial grade and purified prior to use if necessary. Thin layer chromatography (TLC) was performed using glass-backed silica gel (250 μm) plates or glass-backed Biotage^®^ KP-NH plates. UV light, and/or the use of potassium iodoplatinate, potassium permanganate and ninhydrin stains were used to visualize products. MPLC was performed on a Biotage Isolera in conjunction with a Biotage Dalton 2000 using the conditions indicated. Nuclear magnetic resonance spectra (NMR) were acquired on a Bruker AV-III-500 (500 MHz) spectrometer equipped with a TCI cryoprobe. Chemical shifts were measured relative to residual solvent peaks as an internal standard set to δ 7.26 and δ 77.0 (CDCl_3_) or 2.50 and 39.5 (DMSO-*d_6_*). All reported compounds were >95% pure by ^1^H NMR analysis. Low-resolution mass spectra (LRMS)^[1]^ were recorded on a Biotage Dalton 2000 or an Advion Expression Compact Mass Spectrometer using the indicated ionization method. High-resolution mass spectra (HRMS)^[2]^ were recorded at the Indiana University Mass Spectrometry Facility on a Thermo Scientific Orbitrap XL spectrometer by use of the indicated ionization method. A post-acquisition gain correction was applied using reserpine as a lock mass. Compound **24** was previously reported.^[3]^

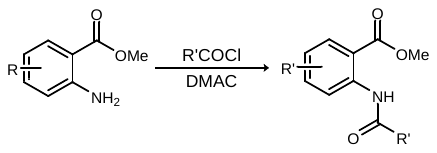

General Procedure A**.** To a solution of the anthranilate in *N,N*-dimethylacetamide at ambient temperature was added the acid chloride dropwise, and the reaction was stirred at ambient temperature for 16 h. The reaction was poured into H_2_O, and the precipitated solid was collected by filtration, washed with water, and dried *in vacuo* to afford the product.

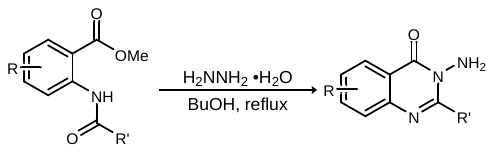

General Procedure B**.** A suspension of the ester in H_2_NNH_2_•H_2_O and 1-butanol was refluxed in a heating mantle for 24 h. After cooling to ambient temperature, the reaction was poured into H_2_O and stirred vigorously for 30 minutes. The precipitate was filtered and dried *in vacuo* to afford the product.

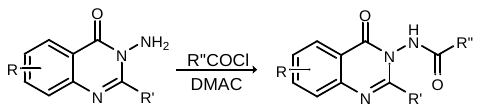

General Procedure C**.** To a solution of the amine in *N,N*-dimethylacetamide at ambient temperature was added the acid chloride dropwise, and the reaction was stirred at ambient temperature for 16 h. The reaction was poured into H_2_O, and the precipitated solid was collected by filtration, washed with water, and dried *in vacuo* to afford the product.

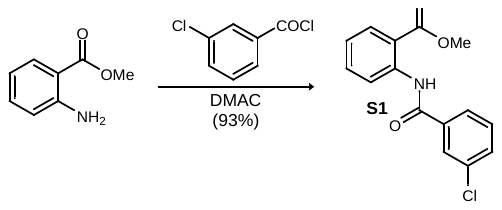

Methyl 2-(3-chlorobenzamido)benzoate (S1)**.** Following General Procedure A, the amine (8.56 mL, 66.1 mmol) and acid chloride (10.2 mL, 79.7 mmol) in *N*-dimethylacetamide (660 mL), after filtration, afforded the amide as a fluffy, colorless solid (17.8 g, 93% yield). ^1^H NMR (500 MHz, DMSO-*d_6_*) δ 11.49 (s, 1H), 8.43 (d, *J* = 8.3 Hz, 1H), 8.00 (d, *J* = 7.9, 1.3 Hz, 1H), 7.96 (br s, 1H), 7.91 (d, *J* = 7.8 Hz, 1H), 7.76-7.60 (series of m, 3H), 7.27 (t, *J* = 7.4 Hz, 1H), 3.88 (s, 3H); ^13^C NMR (125 MHz, DMSO-*d_6_*) 167.8, 163.4, 139.6, 136.4, 134.2, 133.7, 132.0, 131.0, 130.7, 127.1, 125.7, 123.8, 121.4, 118.1, 52.6; HRMS (ESI) m/z: [M+Na]^+^ calcd for C_15_H_12_ClN_3_NaO_3_ 312.0398 found 312.0399.

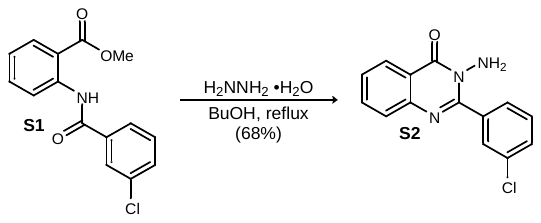

3-Amino-2-(3-chlorophenyl)quinazolin-4(3*H*)-one (S2)**.** Following General Procedure B, the ester (15.5 g, 53.5 mmol) in H_2_NNH_2_•H_2_O (110 mL) and 1-butanol (110 mL), after filtration, afforded the product as a free-flowing, colorless solid (9.85 g, 68% yield). ^1^H NMR (500 MHz, DMSO-*d*_6_) δ 8.19 (d, *J* = 7.4 Hz, 1H), 7.87 (br s, 1H), 7.84 (br dd, *J* = 7.6, 1.2 Hz, 1H), 7.76 (d, *J* = 7.6 Hz, 1H), 7.73 (d, *J* = 8.1 Hz, 1H), 7.61-7.55 (br m, 2H), 7.51 (dd, *J* = 7.8 Hz, 1H), 5.65 (s, 2H); ^13^C NMR (125 MHz, DMSO-*d*_6_) ppm 162.5, 155.9, 147.9, 138.1, 135.7, 133.4, 130.8, 130.74, 130.70, 129.6, 128.8, 128.4, 127.4, 121.6; HRMS (ESI) m/z: [M+H]^+^ calcd for C_14_H_11_ClN_3_O 272.0585; found 272.0587.

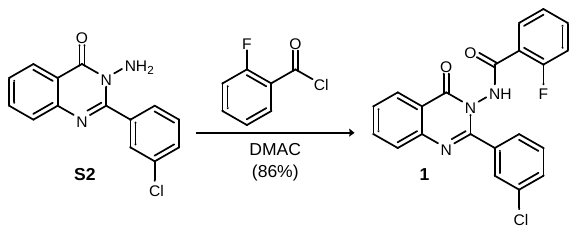

*N*-(2-(3-Chlorophenyl)-4-oxoquinazolin-3(4*H*)-yl)-2-fluorobenzamide (1)**.** Following General Procedure C, the amine (1.00 g, 3.68 mmol) and acid chloride (528 µL, 4.42 mmol) in *N,N*-dimethylacetamide (40 mL), after filtration, afforded the amide as a colorless solid (1.25 g, 86% yield). ^1^H NMR (500 MHz, CDCl_3_) δ 9.07 (d, *J* = 11.7 Hz, 1H), 8.29 (d. *J* = 7.7 Hz, 1H), 7.96 (ddd, *J* = 7.7, 7.7, 1.4 Hz, 1H), 7.86-7.74 (series of br m, 3H), 7.64 (br d, *J* = 7.7 Hz, 1H), 7.58-7.48 (series of br m, 2H), 7.42 (d, *J* = 8.5 Hz, 1H), 7.35 (dd, *J* = 7.8, 7.8 Hz, 1H), 7.24 (dd, *J* = 7.6 Hz, 1H), 7.10 (dd, *J* = 11.6, 8.5 Hz, 1H); ^13^C NMR (125 MHz, CDCl_3_) ppm 163.5 (d, ^3^*J*_C-F_ = 3.8 Hz), 160.8 (d, ^1^*J*_C-F_ = 249 Hz), 160.2, 154.4, 146.8, 135.3, 135.0, 134.9 (d, ^3^*J*_C-F_ = 9.3 Hz), 134.2, 132.2 (d, ^4^*J*_C-F_ = 1.9 Hz), 130.5, 129.5, 128.7, 128.1, 127.6, 127.2, 126.6, 125.0 (d, ^3^*J*_C-F_ = 3.2 Hz), 121.0, 117.9 (d, ^2^*J*_C-F_ = 12.5 Hz), 116.3 (d, ^2^*J*_C-F_ = 24.0 Hz); ^19^F NMR (470 MHz, CDCl_3_) δ -110.0; HRMS (ESI) m/z: [M+H]^+^ calcd for C_21_H_14_ClFN_3_O_2_ 395.0753; found 395.0756.

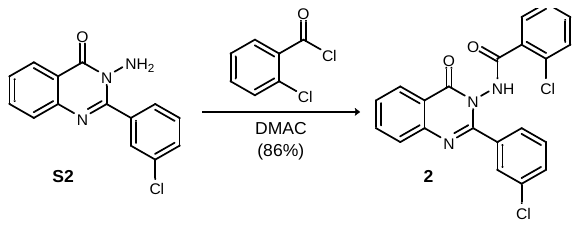

2-Chloro-*N*-(2-(3-chlorophenyl)-4-oxoquinazolin-3(4*H*)-yl)benzamide (2). Following General Procedure C, the amine (75.0 mg, 276 µmol) and acid chloride (42.0 µL, 331 µmol) in dimethylacetamide (2.8 mL), after filtration, afforded the amide as a colorless solid (97.2 mg, 86% yield). ^1^H NMR (500 MHz, DMSO-*d_6_*) δ 11.78 (s, 1H), 8.26 (dd, *J* = 7.9, 1.0 Hz, 1H), 7.97 (ddd, *J* = 8.3, 8.3, 1.4 Hz, 1H), 7.82 (d, *J* = 8.0 Hz, 1H), 7.73 (br s, 1H), 7.70-7.61 (series of br m, 3H), 7.59-7.50 (series of br m, 3H), 7.49-7.41 (m, 1H), 7.31 (d, *J* = 7.5 Hz, 1H); ^13^C NMR (125 MHz, DMSO-*d_6_*) ppm 165.2, 159.1, 154.7, 146.4, 135.5, 135.1, 133.0, 132.7, 132.3, 130.4, 130.2, 130.1, 129.0, 128.2, 128.0, 127.9, 127.32, 127.25, 126.7, 120.9; HRMS (ESI) m/z: [M+H]^+^ calcd for C_21_H_14_Cl_2_N_3_O_2_ 410.0458; found 410.0460.

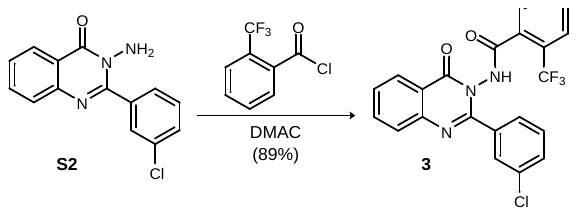

*N*-(2-(3-Chlorophenyl)-4-oxoquinazolin-3(4*H*)-yl)-2-(trifluoromethyl)benzamide (3)**.** Following General Procedure C, the amine (75 mg, 276 µmol) and acid chloride (45 µL, 304 µmol) in dimethylacetamide (3.0 mL) after filtration afforded the product as a colorless solid (109 mg, 89% yield). ^1^H NMR (500 MHz, DMSO-*d*_6_) δ 11.90 (s, 1H), 8.26 (d, *J* = 7.4 Hz, 1H), 7.97 (br dd, *J* = 7.1, 7.1 Hz, 1H), 7.89-7.60 (series of br m, 8H), 7.56 (dd, *J* = 7.9 Hz, 1H), 7.45 (d, *J* = 7.4 Hz, 1H); ^13^C NMR (125 MHz, DMSO-*d*_6_) ppm 165.7, 159.3, 154.7, 146.4, 135.5, 135.0, 132.63, 132.58, 132.1, 131.4, 130.1, 130.0, 128.9, 128.2, 127.89, 127.86, 127.3, 126.8 (q, ^3^*J*_C-F_ = 3.9 Hz), 126.7, 126.5, 123.0 (q, ^1^*J*_C-F_ = 272 Hz), 120.9; ^19^F NMR (470 MHz, DMSO-*d*_6_) δ -58.2; HRMS (ESI) m/z: [M+H]^+^ calcd for C_22_H_14_ClF_3_N_3_O_2_ 444.0721; found 444.0723.

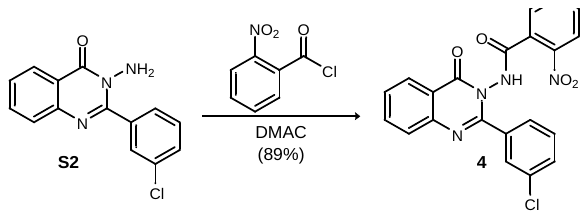

*N*-(2-(3-Chlorophenyl)-4-oxoquinazolin-3(4*H*)-yl)-2-nitrobenzamide (4)**.** Following General Procedure C, the amine (75.0 mg, 276 µmol) and acid chloride (61.4 mg, 331 µmol) in dimethylacetamide (2.8 mL), after filtration, afforded the amide as a colorless solid (103 mg, 89% yield). ^1^H NMR (500 MHz, DMSO-*d_6_*) δ 12.09 (s, 1H), 8.26 (dd, *J* = 7.9, 1.0 Hz, 1H), 8.07 (d, *J* = 8.0 Hz, 1H), 7.97 (ddd, *J* = 8.3, 8.3, 1.4 Hz, 1H), 7.87 (ddd, *J* = 7.5, 7.5, 0.7 Hz, 1H), 7.82 (d, *J* = 8.1 Hz, 1H), 7.79 (ddd, *J* = 7.5, 7.5, 1.2 Hz, 1H), 7.76-7.72 (br m, 1H), 7.70-7.62 (series of br m, 3H), 7.57 (dd, *J* = 7.9, 7.9 Hz, 1H), 7.51 (dd, *J* = 7.5, 1.0 Hz, 1H); ^13^C NMR (125 MHz, DMSO-*d_6_*) ppm 164.4, 159.3, 154.6, 147.2, 146.4, 135.5, 135.0, 133.7, 132.7, 132.4, 130.3, 130.0, 129.1, 128.5, 128.3, 127.92, 127.89, 127.3, 126.7, 124.6, 120.9; HRMS (ESI) m/z: [M+H]^+^ calcd for C_21_H_14_ClN_4_O_4_ 421.0698; found 421.0700.

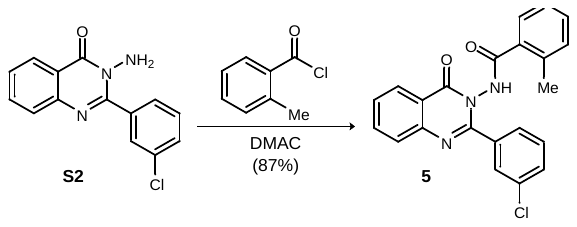

*N*-(2-(3-Chlorophenyl)-4-oxoquinazolin-3(4*H*)-yl)-2-methylbenzamide (5)**.** Following General Procedure C, the amine (100 mg, 368 µmol) and acid chloride (59.6 µL, 442 µmol) in dimethylacetamide (3.7 mL) after filtration afforded the product as a colorless solid (125 mg, 87% yield). ^1^H NMR (500 MHz, DMSO-*d*_6_) δ 11.71 (s, 1H), 8.23 (d, *J* = 7.9 Hz, 1H), 7.96 (dt, *J* = 7.7, 1.2 Hz, 1H), 7.83 (d, *J* = 8.1 Hz, 1H), 7.79 (s, 1H), 7.71-7.63 (series of m, 2H), 7.60-7.46 (series of m, 4H), 7.46-7.35 (series of m, 2H), 2.35 (s, 3H); ^13^C NMR (125 MHz, DMSO-*d*_6_) ppm 165.6, 159.4, 154.8, 146.5, 138.2, 135.5, 135.2, 133.3, 132.5, 131.1, 130.2, 130.0, 128.6, 128.3, 128.0, 127.89, 127.88, 127.2, 126.6, 124.5, 120.9, 20.9; HRMS (ESI) m/z: [M+H]^+^ calcd for C_22_H_17_ClN_3_O_2_ 390.1004; found 390.1006.

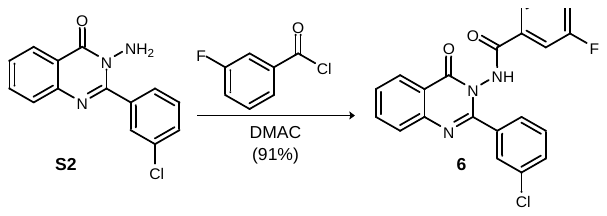

2-Chloro-*N*-(2-(3-chlorophenyl)-4-oxoquinazolin-3(4*H*)-yl)benzamide (6)**.** Following General Procedure C, the amine (75.0 mg, 276 µmol) and acid chloride (40 µL, 331 µmol) in dimethylacetamide (3 mL), after filtration, afforded the amide as a colorless solid (98.8 mg, 91% yield). ^1^H NMR (500 MHz, CDCl_3_) δ 10.4 (s, 1H), 8.36 (d, *J* = 7.9 Hz, 1H), 7.91-7.76 (series of m, 3H), 7.67 (d, *J* = 7.6 Hz, 1H), 7.58 (t, *J* = 7.2 Hz, 1H), 7.47-7.33 (series of m, 3H), 7.28-7.18 (series of m, 2H), 7.11 (dt, *J* = 8.2, 2.2 Hz, 1H); ^13^C NMR (125 MHz, CDCl_3_) ppm 164.4 (d, ^4^*J*_C-F_ = 2.5 Hz), 162.0 (d, ^1^*J*_C-F_ = 247 Hz), 161.5, 153.8, 146.6, 135.3, 134.3, 133.8, 132.0 (d, ^3^*J*_C-F_ = 7.1 Hz), 130.2, 130.0 (d, ^3^*J*_C-F_ = 7.8 Hz), 129.2, 128.4, 127.8, 127.5, 126.7, 126.3, 122.2 (d, ^4^*J*_C-F_ = 3.1 Hz), 120.2, 119.4 (d, ^2^*J*_C-F_ = 21 Hz), 114.4 (d, ^2^*J*_C-F_ = 23.2 Hz); ^19^F NMR (470 MHz, CDCl_3_) δ -110.9;^[4]^ HRMS (ESI) m/z: [M+H]^+^ calcd for C_21_H_14_ClFN_3_O_2_ 394.0753 found 394.0753.

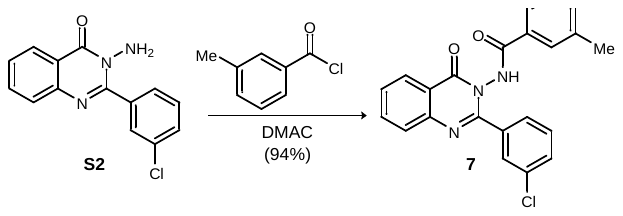

*N*-(2-(3-Chlorophenyl)-4-oxoquinazolin-3(4*H*)-yl)-3-methylbenzamide (7)**.** Following General Procedure C, the amine (75.0 mg, 276 µmol) and acid chloride (40.0 µL, 304 µmol) in dimethylacetamide (3.0 mL), after filtration, afforded the amide as a colorless solid (101 mg, 94% yield). ^1^H NMR (500 MHz, DMSO-*d_6_*) δ11.72 (s, 1H), 8.23 (d, *J* = 7.9 Hz, 1H), 7.96 (ddd, *J* = 7.6, 7.6, 1.2 Hz, 1H), 7.83 (d, *J* = 8.1 Hz, 1H), 7.79 (br s, 1H), 7.72-7.63 (series of m, 2H), 7.61-7.48 (series of m, 4H), 7.42 (dd, *J* = 7.5, 7.5 Hz, 1H), 7.39 (dd, *J* = 7.5, 7.5 Hz, 1H), 2.35 (s, 3H); ^13^C NMR (125 MHz, DMSO-*d_6_*) ppm 166.1, 159.8, 155.2, 146.9, 138.6, 135.9, 135.6, 133.7, 133.0, 131.5, 130.6, 130.4, 129.1, 128.7, 128.4, 128.32, 128.29, 127.6, 127.0, 124.9, 121.3, 21.3; HRMS (ESI) m/z: [M+H]^+^ calcd for C_22_H_17_ClN_3_O_2_ 390.1004; found 390.1006.

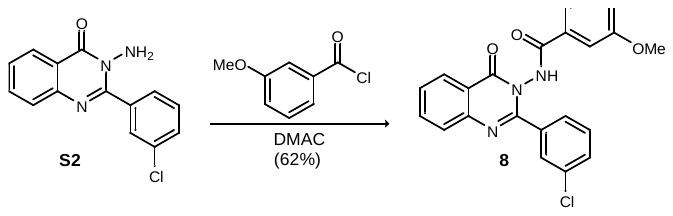

*N*-(2-(3-Chlorophenyl)-4-oxoquinazolin-3(4*H*)-yl)-3-methoxybenzamide (8)**.** Following General Procedure C, the amine (100 mg, 368 µmol) and acid chloride (62 µL, 442 µmol) in dimethylacetamide (3.7 mL) after filtration afforded the product as a colorless solid (93.1 mg, 62% yield). ^1^H NMR (500 MHz, DMSO-*d*_6_) 11.78 (s, 1H), 8.23 (d, *J* = 7.2 Hz, 1H), 7.96 (dt, *J* = 7.7, 1.2 Hz 1H), 7.83 (d, *J* = 8.1 Hz, 1H), 7.79 (br s, 1H), 7.69 (br d, *J* = 8.3 Hz, 1H), 7.66 (d, *J* = 7.3 Hz, 1H), 7.58 (br d, *J* = 8.7 Hz, 1H), 7.52 (dd, *J* = 7.8, 7.8 Hz, 1H), 7.31 (br d, *J* = 7.7 Hz, 1H), 7.24 (br s, 1H), 7.18 (dd, *J* = 8.2, 2.2 Hz, 1H), 3.80 (s, 3H); ^13^C NMR (125 MHz, DMSO-*d*_6_) ppm 165.3, 159.4, 159.3, 154.8, 146.5, 135.5, 135.1, 132.6, 132.4, 130.2, 130.01, 129.99, 128.3, 127.9, 127.2, 126.6, 120.9, 119.6, 118.6, 112.4, 55.3;^[5]^ HRMS (ESI) m/z: [M+H]^+^ calcd for C_22_H_17_ClN_3_O_3_ 406.0953; found 406.0953.

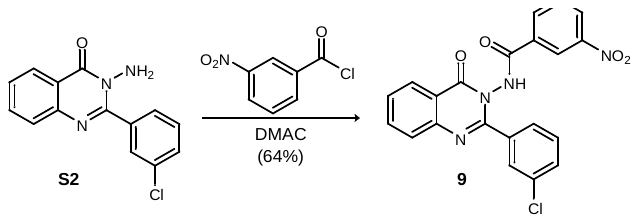

*N*-(2-(3-Chlorophenyl)-4-oxoquinazolin-3(4*H*)-yl)-3-nitrobenzamide (9)**.** Following General Procedure C, the amine (75.0 mg, 276 µmol) and acid chloride (61.4 mg, 331 µmol) in dimethylacetamide (2.8 mL), after filtration, afforded the amide as a colorless solid (74.0 mg, 64% yield). ^1^H NMR (500 MHz, DMSO-*d_6_*) δ 12.24 (s, 1H), 8.55 (br s, 1H), 8.47 (dd, *J* = 8.2, 1.5 Hz, 1H), 8.24 (d, *J* = 7.2 Hz, 1H), 8.17 (d, *J* = 7.8 Hz, 1H), 7.98 (br ddd, *J* = 7.7, 7.7, 1.2 Hz, 1H), 7.88-7.81 (series of m, 2H), 7.79 (br s, 1H), 7.71-7.61 (m, 2H), 7.58 (br d, *J* = 8.5 Hz, 1H), 7.51 (dd, *J* = 7.8, 7.8 Hz, 1H); ^13^C NMR (125 MHz, DMSO-*d_6_*) ppm 164.2, 159.7, 154.9, 148.3, 146.8, 136.0, 135.4, 134.2, 133.1, 132.6, 131.3, 130.7, 130.5, 128.7, 128.42, 128.35, 127.8, 127.5, 127.1, 122.5, 121.3; HRMS (ESI) m/z: [M+H]^+^ calcd for C_21_H_14_ClN_4_O_4_ 421.0698; found 421.0699.

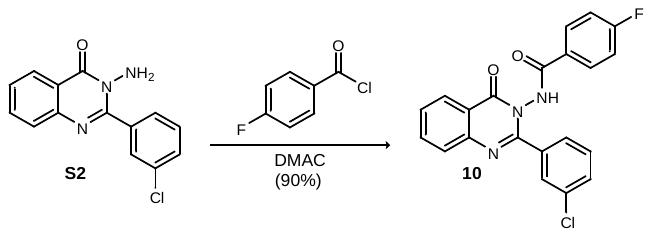

*N*-(2-(3-Chlorophenyl)-4-oxoquinazolin-3(4*H*)-yl)-4-fluorobenzamide (10)**.** Following General Procedure C, the amine (125 mg, 460 µmol) and acid chloride (65 µL, 552 µmol) in dimethylacetamide (4.6 mL) after filtration afforded the product as a colorless solid (163 mg, 90% yield). ^1^H NMR (500 MHz, DMSO-*d*_6_) δ11.84 (s, 1H), 8.22 (br d, *J* = 7.2 Hz, 1H), 8.00-7.93 (br m, 1H), 7.89-7.78 (series of br m, 3H), 7.77 (br s, 1H), 7.67 (br dd, *J* = 7.8, 7.8 Hz, 2H), 7.57 (br d, *J* = 8.1 Hz, 1H), 7.51 (dd, *J* = 7.8, 7.8 Hz, 1H), 7.37 (dd, *J* = 8.8, 8.8 Hz, 2H); ^13^C NMR (125 MHz, DMSO-*d*_6_) ppm 164.7 (d, ^1^*J*_C-F_ = 249 Hz), 164.6, 159.5, 154.8, 146.5, 135.6, 135.2, 132.7, 130.3, 130.2 (d, ^2^*J*_C-F_ = 39 Hz), 128.3, 128.0 (d, ^3^*J*_C-F_ = 5.6 Hz), 127.5 (d, ^4^*J*_C-F_ = 3.0 Hz), 127.2, 126.7, 120.9, 116.1, 115.9; ^19^F NMR (470 MHz, DMSO-*d*_6_) δ -106.5;^4^ HRMS (ESI) m/z: [M+H]^+^ calcd for C_21_H_14_ClFN_3_O_2_ 394.0753; found 394.0754.

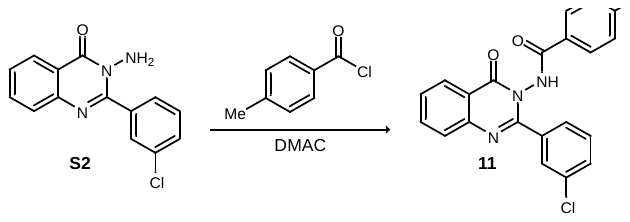

*N*-(2-(3-Chlorophenyl)-4-oxoquinazolin-3(4*H*)-yl)-4-methylbenzamide (11)**.** Following General Procedure C, the amine (75.0 mg, 276 µmol) and acid chloride (40 µL, 304 µmol) in dimethylacetamide (3.0 mL), after filtration, afforded the amide as a colorless solid (77 mg).^[6]^ ^1^H NMR (500 MHz, DMSO-*d_6_*) δ 11.69 (s, 1H), 8.23 (d, *J* = 7.8 Hz, 1H), 7.96 (br t, *J* = 7.2 Hz, 1H), 7.86-7.74 (series of br m, 4H), 7.71-7.67 (series of br m, 4H), 7.56 (br d, *J* = 8.3 Hz, 1H), 7.50 (t, *J* = 7.8 Hz, 1H), 2.36 (s, 3H); HRMS (ESI) m/z: [M+H]^+^ calcd for C_22_H_17_ClN_3_O_2_ 390.1004; found 390.1004.

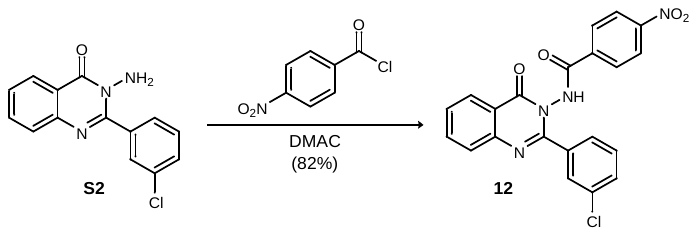

*N*-(2-(3-Chlorophenyl)-4-oxoquinazolin-3(4*H*)-yl)-4-nitrobenzamide (12)**.** Following General Procedure C, the amine (75.0 mg, 276 µmol) and acid chloride (56.4 mg, 304 µmol) in dimethylacetamide (3.0 mL), after filtration, afforded the amide as a colorless solid (95.6 mg, 82% yield). ^1^H NMR (500 MHz, DMSO-*d_6_*) δ 12.20 (s, 1H), 8.37 (d, *J* = 8.8 Hz, 2H), 8.24 (br d, *J* = 7.0 Hz, 1H), 7.98 (ddd, *J* = 8.4, 8.4, 1.2 Hz, 1H), 7.94 (d, *J* = 8.8 Hz, 2H), 7.84 (d, *J* = 8.1 Hz, 1H), 7.77 (br s, 1H), 7.71-7.65 (series of br m, 2H), 7.59 (br d, *J* = 8.8 Hz, 1H), 7.52 (dd, *J* = 7.9, 7.9 Hz, 1H); ^13^C NMR (125 MHz, DMSO-*d_6_*) ppm 164.2, 159.3, 154.5, 149.9, 146.4, 136.4, 135.6, 135.0, 132.7, 130.3, 130.0, 129.0, 128.2, 128.0, 127.9, 127.1, 126.7, 124.1, 120.8; HRMS (ESI) m/z: [M+H]^+^ calcd for C_21_H_14_ClN_4_O_4_ 421.0698; found 421.0699.

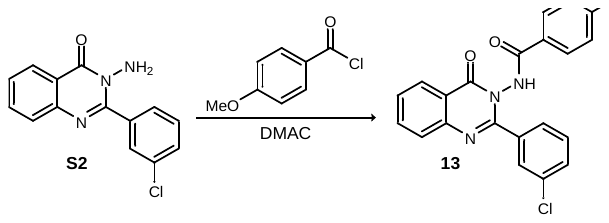

*N*-(2-(3-Chlorophenyl)-4-oxoquinazolin-3(4*H*)-yl)-4-methoxybenzamide (13)**.** Following General Procedure C, the amine (100 mg, 368 µmol) and acid chloride (60 µL, 442 µmol) in dimethylacetamide (3.7 mL), after filtration, afforded the amide as a colorless solid (245 mg).^[7]^ ^1^H NMR (500 MHz, DMSO-*d_6_*) δ 11.60 (s, 1H), 8.22 (d, *J* = 7.5 Hz, 1H), 7.96 (br t, *J* = 7.1 Hz, 1H), 7.82 (d, *J* = 8.1 Hz, 1H), 7.77 (br s, 1H), 7.74 (d, *J* = 8.8 Hz, 1H), 7.70-7.62 (series of m, 2H), 7.56 (br d, *J* = 8.6 Hz, 1H), 7.49 (t, *J* = 7.8 Hz, 1H), 7.05 (d, *J* = 8.8 Hz, 2H), 3.80 (s, 3H); HRMS (ESI) m/z: [M+H]^+^ calcd for C_22_H_17_ClN_3_O_3_ 406.0953; found 406.0954.

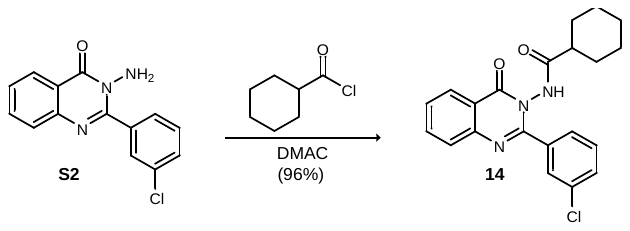

*N*-(2-(3-Chlorophenyl)-4-oxoquinazolin-3(4*H*)-yl)cyclohexanecarboxamide (14)**.** Following General Procedure C, the amine (75 mg, 276 µmol) and acid chloride (43. µL, 331 µmol) in dimethylacetamide (3.0 mL) after filtration afforded the product as a colorless solid (101 mg, 96% yield). ^1^H NMR (500 MHz, DMSO-*d_6_*) δ 8.77-8.62 (series of br m, 2H), 7.94 (br d, *J* = 7.8 Hz, 1H), 7.52 (br dd, *J* = 7.7, 5.0 Hz, 1H), 7.30-6.68 (series of br m, 4H), 5.04 (br dd, *J* = 10.1, 3.3 Hz, 1H), 4.63-4.50 (br m, 1H), 4.49-4.14 (br m, 2H), 4.08 (br d, *J* = 17.2 Hz, 1H), 3.45-3.24 (br m, 1H), 2.97-2.76 (series of br m, 3H); ^13^C NMR (125 MHz, DMSO-*d_6_*) ppm 167.0, 163.8, 150.9, 147.9, 135.2, 134.9, 133.0, 129.0, 127.1, 126.6, 125.2, 123.5, 54.1, 38.5, 28.2; HRMS (ESI) m/z: [M+H]^+^ calcd for C_21_H_21_ClN_3_O_2_ 382.1317; found 382.1319.

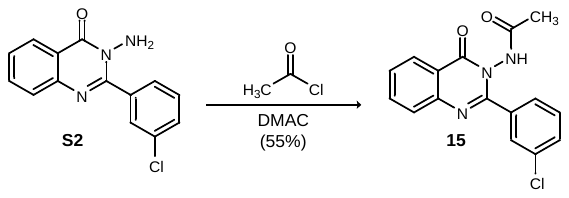

*N*-(2-(3-Chlorophenyl)-4-oxoquinazolin-3(4*H*)-yl)acetamide (15)**.** To a solution of the amine (75 mg, 276 µmol) in dimethylacetamide (3.0 mL) was added acetyl chloride (1.0 in CH_2_Cl_2_, 359 µL, 359 µmol), and the reaction was stirred at ambient temperature for 24 h. The reaction was poured into H_2_O and allowed to stand at ambient temperature for 24 h. Tan crystals formed and were collected by filtration, washed with H_2_O, and dried *in vacuo* to afford the product as a tan solid (47.7 mg, 55% yield). ^1^H NMR (500 MHz, DMSO-*d*_6_) δ 11.15 (s, 1H), 8.19 (d, *J* = 7.1 Hz, 1H), 7.96-7.89 (br m, 1H), 7.78 (d, *J* = 8.1 Hz, 1H), 7.68-7.50 (series of br m, 5H), 1.85 (s, 3H); ^13^C NMR (125 MHz, DMSO-*d*_6_) ppm 168.9, 159.3, 154.9, 146.5, 135.41, 135.39, 132.6, 130.1, 130.0, 128.2, 127.82, 127.79, 127.1, 126.6, 121.0, 20.3; HRMS (ESI) m/z: [M+H]^+^ calcd for C_16_H_13_ClN_3_O_2_ 314.0691; found 314.0693.

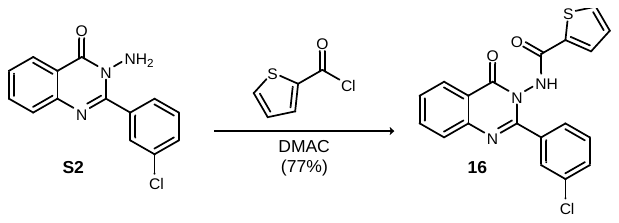

*N*-(2-(3-Chlorophenyl)-4-oxoquinazolin-3(4*H*)-yl)thiophene-2-carboxamide (16)**.** Following General Procedure C, the amine (75 mg, 276 µmol) and acid chloride (35.4 µL, 331 µmol) in dimethylacetamide (3.0 mL) after filtration afforded the product as a colorless solid (81.1 mg, 77% yield). ^1^H NMR (500 MHz, DMSO-*d*_6_) δ11.79 (s, 1H), 8.23 (d, *J* = 7.3 Hz, 1H), 7.97 (ddd, *J* = 7.1, 7.1, 1.1 Hz, 1H), 7.92 (d, *J* = 4.8 Hz, 1H), 7.83 (br d, *J* = 7.5 Hz, 1H), 7.81 (d, *J* = 4.0 Hz, 1H), 7.77 (br s, 1H), 7.66 (m, 2H), 7.56 (br d, *J* = 8.2 Hz, 1H), 7.50 (dd, *J* = 7.8 Hz, 1H), 7.22 (dd, *J* = 4.1 Hz, 1H); ^13^C NMR (125 MHz, DMSO-*d*_6_) ppm 160.5, 159.6, 154.8, 146.5, 135.6, 135.2, 135.0, 133.2, 132.6, 130.5, 130.3, 130.1, 128.5, 128.4, 128.01, 127.96, 127.3, 126.7, 120.9; HRMS (ESI) m/z: [M+H]^+^ calcd for C_19_H_13_ClN_3_O_2_S 382.0412; found 382.0413.

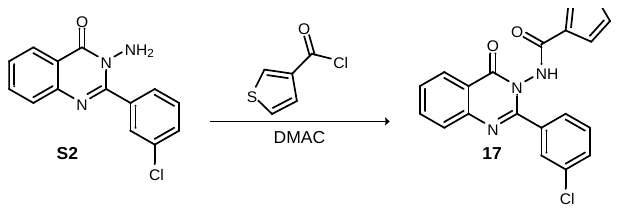

*N*-(2-(3-chlorophenyl)-4-oxoquinazolin-3(4*H*)-yl)thiophene-3-carboxamide (17)**.**

HRMS (ESI) m/z: [M+H]^+^ calcd for C_19_H_13_ClN_3_O_2_S 382.0412; found 382.0412.

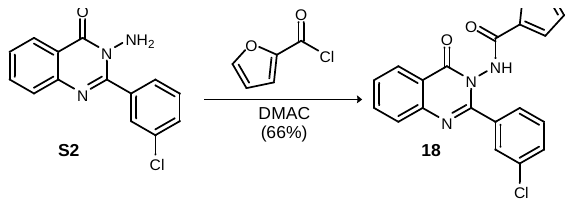

*N*-(2-(3-chlorophenyl)-4-oxoquinazolin-3(4*H*)-yl)furan-2-carboxamide (18)**.** Following General Procedure C, the amine (100 mg, 368 µmol) and acid chloride (44 µL, 442 µmol) in dimethylacetamide (3.7 mL), after filtration, afforded the amide as a colorless solid (88.8 mg, 66% yield). ^1^H NMR (500 MHz, DMSO-*d_6_*) δ 8.77-8.62 (series of br m, 2H), 7.94 (br d, *J* = 7.8 Hz, 1H), 7.52 (br dd, *J* = 7.7, 5.0 Hz, 1H), 7.30-6.68 (series of br m, 4H), 5.04 (br dd, *J* = 10.1, 3.3 Hz, 1H), 4.63-4.50 (br m, 1H), 4.49-4.14 (br m, 2H), 4.08 (br d, *J* = 17.2 Hz, 1H), 3.45-3.24 (br m, 1H), 2.97-2.76 (series of br m, 3H); ^13^C NMR (125 MHz, DMSO-*d_6_*) ppm 167.0, 163.8, 150.9, 147.9, 135.2, 134.9, 133.0, 129.0, 127.1, 126.6, 125.2, 123.5, 54.1, 38.5, 28.2; HRMS (ESI) m/z: [M+H]^+^ calcd for C_19_H_13_ClN_3_O_3_ 366.0640; found 366.0640.

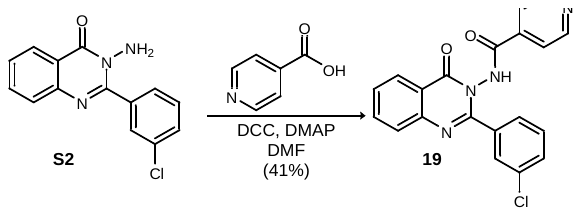

*N*-(2-(3-Chlorophenyl)-4-oxoquinazolin-3(4*H*)-yl)isonicotinamide (19)**.** To a solution of the acid (45.3 mg, 368 µmol), amine (100 mg, 368 µmol), and DMAP (9 mg, 74 µmol) in DMF (2 ML) was added DCC (1 M in CH_2_Cl_2_, 368 µL, 368 µmol), and the reaction was stirred for 24 h at ambient temperature before concentrating *in vacuo*. The residue was dissolved in CH_2_Cl_2_ and filtered through Celite^®^. The filtrate was washed with sat aq NaHCO_3_, dried, and dry-loaded onto Celite^®^. MPLC (Sfar KP-Amino Duo, 12g, 10-80% ethyl acetate in hexanes), afforded the product as a tan amorphous solid^[8]^ (57.0 mg, 41% yield). ^1^H NMR (500 MHz, DMSO-*d_6_*) δ 12.41 (s, 1H), 8.84 (d, *J* = 5.3 Hz, 2H), 8.23 (d, *J* = 7.7 Hz, 1H), 7.97 (t, *J* = 7.4 Hz, 1H), 7.84 (d, *J* = 8.1 Hz, 1H), 7.79 (s, 1H), 7.75 (d, *J* = 5.4 Hz, 2H), 7.72-7.63 (series of br m, 2H), 7.58 (br d, *J* = 8.4 Hz, 1H), 7.52 (t, *J* = 7.8 Hz, 1H); ^13^C NMR (125 MHz, DMSO-*d_6_*) ppm 163.8, 159.2, 154.4, 149.6, 146.4, 135.6, 134.9, 132.7, 130.3, 130.0, 128.3, 128.0, 127.9, 127.1, 126.7, 121.7, 120.8;^[9]^ HRMS (ESI) m/z: [M+H]^+^ calcd for C_20_H_14_ClN_4_O_2_ 377.0800; found 377.0801.

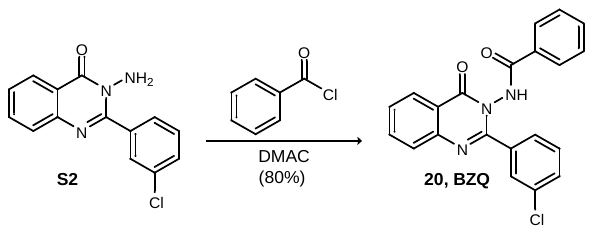

*N*-(2-(3-Chlorophenyl)-4-oxoquinazolin-3(4*H*)-yl)benzamide (20, BZQ)**.** Following General Procedure C, the amine (893 mg, 3.29 mmol) and acid chloride (459 µL, 3.95 mmol) in dimethylacetamide (33 mL) after filtration afforded **BZQ** as a colorless solid (991 mg, 80% yield). ^1^H NMR (500 MHz, DMSO-*d*_6_) δ 11.79 (s, 1H), 8.23 (d, *J* = 8.0 Hz, 1H), 8.00-7.91 (m, 1H), 7.83 (d, *J* = 8.0 Hz, 1H), 7.78 (br s, 1H), 7.73 (d, *J* = 7.3 Hz, 2H), 7.71-7.47 (series of br m, 7H); ^13^C NMR (125 MHz, DMSO-*d*_6_) ppm 165.7, 159.5, 154.9, 146.5, 135.6, 135.2, 132.8, 132.6, 131.1, 130.3, 130.0, 128.9, 128.3, 128.0, 127.9, 127.5, 127.3, 126.7, 120.9; HRMS (ESI) m/z: [M+H]^+^ calcd for C_21_H_15_ClN_3_O_2_ 376.0847; found 376.0849.

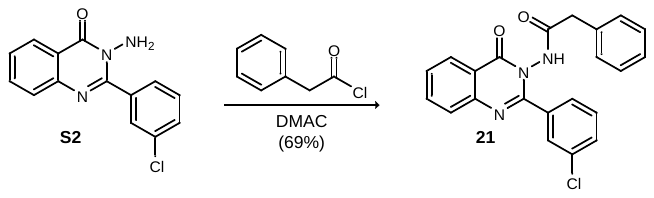

*N*-(2-(3-Chlorophenyl)-4-oxoquinazolin-3(4*H*)-yl)-2-phenylacetamide (21)**.** Following General Procedure C, the amine (75 mg, 276 µmol) and acid chloride (44 µL, 331 µmol) in dimethylacetamide (3.0 mL) after filtration afforded the product as a colorless solid (74.1 mg, 69% yield). ^1^H NMR (500 MHz, DMSO-*d*_6_) δ 11.44 (ds, 1H), 8.20 (d, *J* = 7.4 Hz, 1H), 7.96-7.89 (m, 1H), 7.77 (*J* = 8.1 Hz, 1H), 7.63 (d, *J* = 7.7 Hz, 1H), 7.61 (br s, 1H), 7.57 (br d, *J* = 7.8 Hz, 1H), 7.42 (dd, *J* = 7.8, 7.8 Hz, 1H), 7.25-7.13 (m, 3H), 6.99-6.90 (m, 2H), 3.52 (d, *J* = 14.5 Hz, 1H), 3.47 (d, *J* = 14.5 Hz, 1H); ^13^C NMR (125 MHz, DMSO-*d*_6_) ppm 169.4, 159.3, 154.9, 146.5, 135.4, 135.2, 134.4, 132.9, 130.0, 129.9, 128.7, 128.3, 128.2, 127.8, 127.1, 126.7, 126.6, 121.0;^[10]^ HRMS (ESI) m/z: [M+H]^+^ calcd for C_22_H_17_ClN_3_O_2_ 390.1004; found 390.1005.

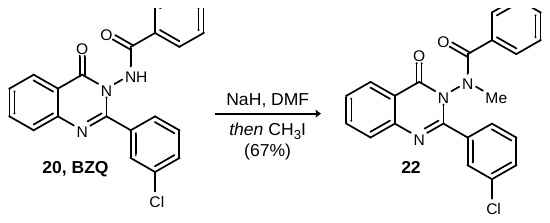

*N*-(2-(3-Chlorophenyl)-4-oxoquinazolin-3(4*H*)-yl)-*N*-methylbenzamide (22)**.** To a suspension of NaH (60% in mineral oil, 24 mg, 599 µmol) in DMF (2 mL), chilled to 0 °C in an ice bath, was added the amide (150 mg, 399 µmol) as a solution in DMF (2 mL). The reaction was stirred for 2 h, and methyl iodide (37.3 µL, 599 µmol) was added. The ice bath was removed, and the reaction was stirred at ambient temperature for 18 h. The reaction was quenched by adding sat aq NH_4_Cl and then concentrated *in vacuo*. The residue was dry loaded onto SiO_2_ and purified *via* MPLC (Sfar HC-Duo, 10 g, 10-80% ethyl acetate in hexanes) to afford the product as a free-flowing, colorless solid (104 mg, 67% yield). ^1^H NMR (500 MHz, DMSO-*d_6_*)^[11]^ δ 8.27 (d, *J* = 7.6 Hz, 1H), 7.98 (t, *J* = 7.2 Hz, 1H), 7.82 (d, *J* = 8.1 Hz, 1H), 7.75-7.60 (series of br m, 5H), 7.54 (br dd, *J* = 7.4, 7.4 Hz, 1H), 7.47 (t, *J* = 7.6 Hz, 2H), 7.12 (d, *J* = 7.2 Hz, 2H), 3.33 (s, 3H); HRMS (ESI) m/z: [M+Na]^+^ calcd for C_22_H_16_ClN_3_NaO_2_ 412.0823; found 412.0824.

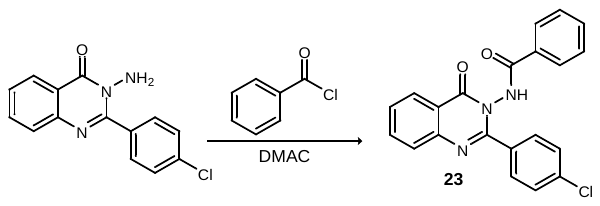

*N*-(2-(4-Chlorophenyl)-4-oxoquinazolin-3(4*H*)-yl)benzamide (23)**.** Following General Procedure C, the amine^[12]^ (100.0 mg, 368 µmol) and acid chloride (54.0 µL, 460 µmol) in dimethylacetamide (1 mL), after filtration, afforded the amide as a tan solid (61 mg)^[13]^; HRMS (ESI) m/z: [M+Na]^+^ calcd for C_21_H_15_ClN_3_O_2_ 376.0847; found 376.0849.

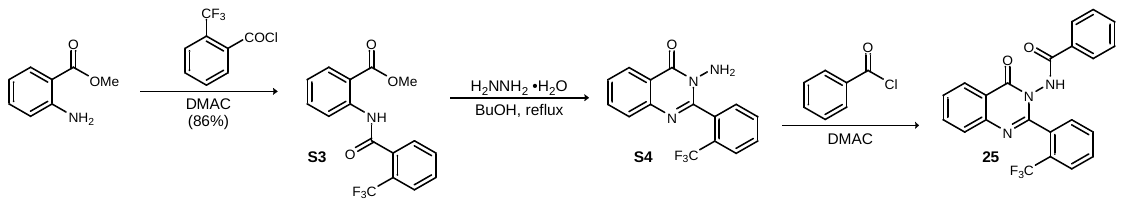

Methyl 2-(2-(trifluoromethyl)benzamido)benzoate (S3)**.** Following General Procedure A, methyl anthranilate (856 µL, 6.62 mmol) and the acid chloride (1.17 mL, 7.94 mmol) in *N,N*-dimethylacetamide (66 mL) after filtration afforded the product as a colorless solid (1.86 g, 97% yield). ^1^H NMR (500 MHz, DMSO-*d*_6_) δ 11.06 (s, 1H), 8.23 (d, *J* = 8.2 Hz, 1H), 7.95 (dd, *J* = 7.9, 1.3 Hz, 1H), 7.92-7.79 (series of m, 4H), 7.69 (ddd, *J* = 8.4, 8.4, 1.2 Hz, 1H), 7.30 (ddd, *J* = 7.9, 7.9, 0.8 Hz, 1H), 3.82 (s, 3H); ^13^C NMR (125 MHz, DMSO-*d*_6_) ppm 167.5, 165.4, 138.8, 135.7 (q, ^3^*J*_C-F_ = 2.1 Hz), 134.0, 133.0, 130.7, 130.5, 128.2, 126.7 (q, ^3^*J*_C-F_ = 4.9 Hz), 126.0 (q, ^2^*J*_C-F_ = 31 Hz), 124.3, 123.7 (q, ^1^*J*_C-F_ = 272 Hz), 121.8, 119.2, 52.5; ^19^F NMR (470 MHz, DMSO-*d*_6_) δ -57.8; HRMS (ESI) m/z: [M+Na]^+^ calcd for C_16_H_12_F_3_NNaO_3_ 346.0661; found 346.0665.

3-Amino-2-(2-(trifluoromethyl)phenyl)quinazolin-4(3*H*)-one (S4)**.** Following General Procedure B the ester (1.00 g, 3.09 mmol) in H_2_NNH_2_•H_2_O (6.2 mL) and 1-butanol (6.2 mL), after filtration, afforded the product as a free-flowing, colorless solid (688 mg).^[14]^ HRMS (ESI) m/z: [M+H]+ calcd for C_15_H_11_F_3_N_3_O 306.0849; found 306.0850.

*N*-(4-Oxo-2-(2-(trifluoromethyl)phenyl)quinazolin-3(4*H*)-yl)benzamide (25)**.** HRMS (ESI) m/z: [M+H]^+^ calcd for C_22_H_15_F_3_N_3_O_2_ 410.1111; found 410.1111.

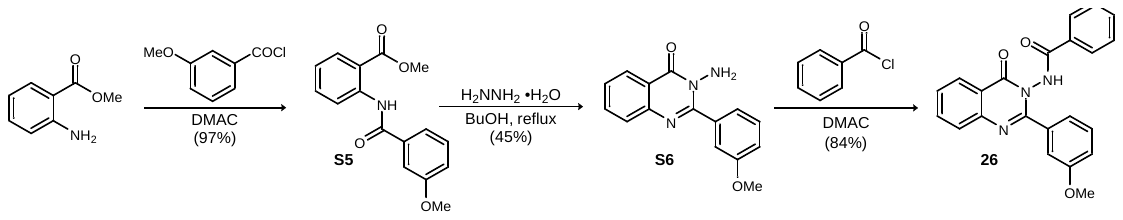

Methyl 2-(3-methoxybenzamido)benzoate (S5)**.** Following General Procedure A, methyl anthranilate (1.00 mL, 7.73 mmol) and the acid chloride (1.19 mL, 8.50 mmol) in *N,N*-dimethylacetamide (77 mL) after filtration afforded the product as a colorless solid (2.14 g, 97% yield). LRMS (APCI) m/z: [M+H]^+^ calcd for C_16_H_16_NO_4_ 286.1; found 286.1.

3-Amino-2-(3-methoxyphenyl)quinazolin-4(3*H*)-one (S6)**.** Following General Procedure B, the ester (1.05 g, 3.68 mmol) in H_2_NNH_2_•H_2_O (13 mL) and 1-butanol (13 mL), after filtration, afforded the product as a free-flowing, colorless solid (441 mg, 45% yield). LRMS (APCI) m/z: [M+H]^+^ calcd for C_15_H_14_N_3_O_2_ 268.1; found 268.2.

*N*-(2-(3-Methoxyphenyl)-4-oxoquinazolin-3(4*H*)-yl)benzamide (26)**.** Following General Procedure C, the amine (100.0 mg, 374 µmol) and acid chloride (48.0 µL, 445 µmol) in dimethylacetamide (3.3 mL), after filtration, afforded the amide as a colorless solid (104 mg, 84% yield). ^1^H NMR (500 MHz, DMSO-*d_6_*) δ 11.72 (s, 1H), 8.23 (br dd, *J* = 7.9, 0.85 Hz, 1H), 7.95 (ddd, *J* = 8.2, 1.3, 1.3 Hz, 1H), 7.81 (d, *J* = 8.0 Hz, 1H), 7.74 (d, *J* = 7.3 Hz, 2H), 7.65 (dd, *J* = 7.7, 7.7 Hz, 1H), 7.61 (dd, *J* = 7.4, 7.4 Hz, 1H), 7.51 (dd, *J* = 7.7, 7.7 Hz, 2H), 7.38 (dd, *J* = 7.9, 7.9 Hz, 1H), 7.32-7.24 (series of br m, 2H), 7.05 (dd, *J* = 8.2, 2.2 Hz, 1H), 3.77 (s, 3H); ^13^C NMR (125 MHz, DMSO-*d_6_*) ppm 165.4, 159.6, 158.5, 156.0, 146.6, 135.4, 134.6, 132.7, 131.2, 129.2, 128.7, 127.8, 127.6, 127.4, 126.6, 120.78, 120.76, 116.1, 113.8, 55.3; HRMS (ESI) m/z: [M+H]^+^ calcd for C_22_H_18_N_3_O_3_ 372.1343; found 372.1344.

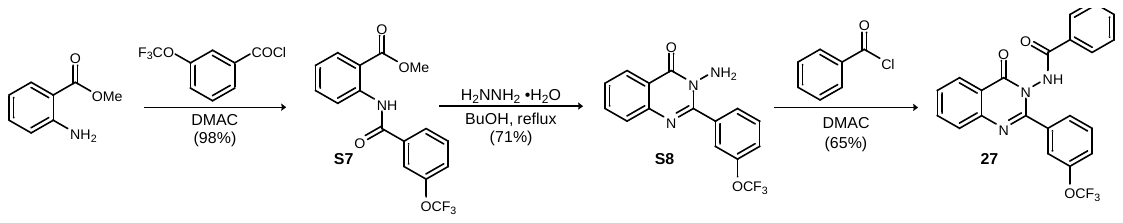

Methyl 2-(3-(trifluoromethoxy)benzamido)benzoate (S7)**.** Following General Procedure A, methyl anthranilate (1.28 mL, 9.92 mmol) and the acid chloride (1.72 mL, 10.9 mmol) in *N,N*-dimethylacetamide (100 mL) after filtration afforded the product as a colorless solid (3.30 g, 98% yield). ^1^H NMR (500 MHz, DMSO-*d*_6_) δ 11.56 (s, 1H), 8.44 (d, *J* = 8.5 Hz, 1H), 7.97 (ddd, *J* = 8.1, 8.1, 1.4 Hz, 1H), 7.86 (br s, 1H), 7.74 (dd, *J* = 8.1 Hz, 1H), 7.70-7.61 (series of m, 2H), 7.25 (dd, *J* = 7.3, 7.3 Hz, 1H), 3.87 (s, 3H); ^13^C NMR (125 MHz, DMSO-*d*_6_) ppm 167.9, 163.2, 148.6 (q, ^3^*J*_C-F_ = 1.8 Hz), 139.6, 136.6, 134.2, 131.2, 130.6, 125.9, 124.6, 123.8, 121.2, 120.0 (q, ^1^*J*_C-F_ = 256 Hz), 119.7, 117.9, 52.6; ^19^F NMR (470 MHz, DMSO-*d*_6_) δ -56.9; HRMS (ESI) m/z: [M+Na]^+^ calcd for C_16_H_12_F_3_NNaO_4_ 362.0611; found 362.0613.

3-Amino-2-(3-(trifluoromethoxy)phenyl)quinazolin-4(3*H*)-one (S8)**.** Following General Procedure B, the ester (1.50 g, 4.42 mmol) in H_2_NNH_2_•H_2_O (8.0 mL) and 1-butanol (8.0 mL) after filtration afforded the product as a free-flowing, colorless solid (1.01 g, 71% yield). ^1^H NMR (500 MHz, DMSO-*d*_6_) δ 8.20 (d, *J* = 7.8 Hz, 1H), 7.92-7.79 (series of br m, 3H), 7.73 (d, *J* = 8.0 Hz, 1H), 7.63 (dd, *J* = 7.9, 7.9 Hz, 1H), 7.59 (dd, *J* = 7.6, 7.6 Hz, 1H), 7.52 (br d, *J* = 7.6 Hz, 1H), 5.66 (s, 2H); ^13^C NMR (125 MHz, DMSO-*d*_6_) ppm 161.2, 154.4, 147.4, 146.5, 136.9, 134.4, 129.6, 128.8, 127.5, 127.0, 126.1, 122.5, 122.1, 120.3, 120.1 (q, ^1^*J*_C-F_ = 255 Hz); ^19^F NMR (470 MHz, DMSO-*d*_6_) δ -56.7; HRMS (ESI) m/z: [M+H]^+^ calcd for C_15_H_11_F_3_N_3_O_2_ 322.0798; found 322.0800.

*N*-(4-Oxo-2-(3-(trifluoromethoxy)phenyl)quinazolin-3(4*H*)-yl)benzamide (27)**.** Following General Procedure C, the amine (100 mg, 311 µmol) and acid chloride (45 µL, 389 µmol) in dimethylacetamide (1.0 mL), after filtration, afforded the amide as a colorless solid (85.6 mg, 65% yield). ^1^H NMR (500 MHz, DMSO-*d_6_*) δ 11.78 (s, 1H), 8.24 (br dd, *J* = 7.9, 0.9 Hz, 1H), 7.97 (br dd, *J* = 7.7, 7.7 Hz, 1H), 7.84 (d, *J* = 8.1 Hz, 1H), 7.80-7.57 (series of m, 7H), 7.55-7.47 (series of m, 3H); ^13^C NMR (125 MHz, DMSO-*d_6_*) ppm 165.6, 159.4, 154.6, 147.5 (q, ^3^*J*_C-F_ = 1.8 Hz), 146.4, 135.5, 135.3, 132.8, 130.9, 130.4, 128.7, 128.0, 127.9, 127.8, 127.4, 126.6, 123.0, 121.0, 120.9, 120.0 (q, ^1^*J*_C-F_ = 255 Hz); ^19^F NMR (470 MHz, CDCl_3_) δ -56.9; HRMS (ESI) m/z: [M+H]^+^ calcd for C_22_H_15_F_3_N_3_O_3_ 426.1060; found 426.1061.

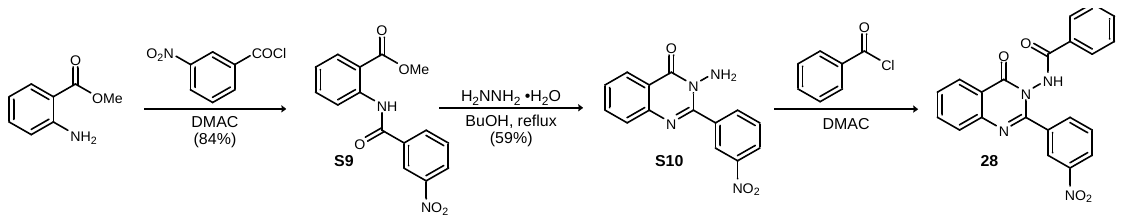

Methyl 2-(3-nitrobenzamido)benzoate (S9)**.** Following General Procedure A, methyl anthranilate (1.71 mL, 13.2 mmol) and the acid chloride (2.93 g, 15.8 mmol) in *N,N*-dimethylacetamide (135 mL) after filtration afforded the product as a colorless solid (3.32 g, 84% yield). ^1^H NMR (500 MHz, DMSO-*d*_6_) δ 11.55 (s, 1H), 8.73 (dd, *J* = 1.8, 1.8 Hz, 1H), 8.48 (dd, *J* = 8.2, 1.5 Hz, 1H), 8.40-8.33 (br m, 2H), 7.99 (dd, *J* = 7.9, 1.4 Hz, 1H), 7.91 (dd, *J* = 7.9, 7.9 Hz, 1H), 7.70 (ddd, *J* = 8.5, 8.5, 1.4 Hz, 1H), 7.30 (ddd, *J* = 8.1, 8.1, 0.8 Hz, 1H), 3.88 (s, 3H); ^13^C NMR (125 MHz, DMSO-*d*_6_) ppm 167.7, 162.8, 148.0, 139.1, 135.8, 134.0, 133.3, 130.8, 130.7, 126.6, 124.2, 122.1, 122.0, 119.1, 52.6; HRMS (ESI) m/z: [M+H]^+^ calcd for C_15_H_13_N_2_O_5_ 301.0819; found 301.0819.

3-Amino-2-(3-nitrophenyl)quinazolin-4(3*H*)-one (S10)**.** Following General Procedure B, the ester (1.50 g, 5.00 mmol) in H_2_NNH_2_•H_2_O (8.0 mL) and 1-butanol (8.0 mL) after filtration afforded the product as a free-flowing, colorless solid (825 mg, 59% yield). ^1^H NMR (500 MHz, DMSO-*d*_6_) δ 8.68 (br s, 1H), 8.36 (dd, *J* = 8.3, 1.5 Hz, 1H), 8.27 (d, *J* = 7.8 Hz, 1H), 8.20 (d, *J* = 7.1 Hz, 1H), 7.87 (ddd, *J* = 8.3, 8.3, 1.3 Hz, 1H), 7.79 (dd, *J* = 8.0, 8.0 Hz, 1H), 7.76 (d, *J* = 8.0 Hz, 1H), 7.60 (dd, *J* = 7.3 Hz, 1H), 5.69 (s, 2H); ^13^C NMR (125 MHz, DMSO-*d*_6_) ppm 161.2, 154.0, 147.0, 146.5, 136.3, 136.2, 134.5, 129.1, 127.5, 127.2, 126.1, 124.6, 124.2, 120.4; HRMS (ESI) m/z: [M+H]^+^ calcd for C_14_H_11_O_3_N_4_ 283.0826; found 283.0826.

*N*-(2-(3-nitrophenyl)-4-oxoquinazolin-3(4*H*)-yl)benzamide (28)**.** Following General Procedure C, the amine (100 mg, 354 µmol) and acid chloride (52 µL, 443 µmol) in dimethylacetamide (1.0 mL), after filtration, afforded the amide as a tan solid (137 mg, 99% yield).^[15]^ ^1^H NMR (500 MHz, DMSO-*d_6_*) δ 11.89 (s, 1H), 8.59 (br s, 1H), 8.35 (br dd, *J* = 8.2, 1.4 Hz, 1H), 8.25 (d, *J* = 7.5 Hz, 1H), 8.19 (d, *J* = 7.8 Hz, 1H), 8.01-7.93 (br m, 1H), 7.87 (d, *J* = 8.1 Hz, 1H), 7.80 (t, *J* = 8.1 Hz, 1H), 7.74 (d, *J* = 7.6 Hz, 2H), 7.69 (t, *J* = 7.7 Hz, 1H), 7.61 (t, *J* = 7.4 Hz, 1H), 7.50 (t, *J* = 7.8 Hz, 2H); ^13^C NMR (125 MHz, DMSO-*d_6_*) ppm 165.8, 159.6, 154.1, 147.2, 146.4, 135.6, 135.0, 134.6, 132.8, 130.9, 129.9, 128.8, 128.1, 128.0, 127.4, 126.7, 125.1, 123.3, 121.0; HRMS (ESI) m/z: [M+Na]^+^ calcd for C_21_H_15_N_4_NaO_4_ 409.0907; found 409.0908.

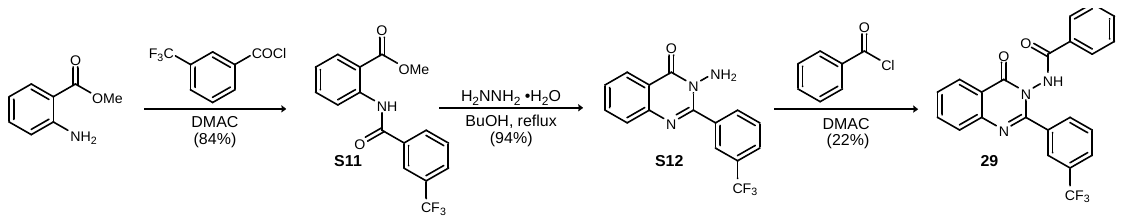

Methyl 2-(3-(trifluoromethyl)benzamido)benzoate (S11)**.** Following General Procedure A, methyl anthranilate (1.71 mL, 13.2 mmol) and the acid chloride (2.19 mL, 14.5 mmol) in *N,N*-dimethylacetamide (132 mL) after filtration afforded the product as a colorless solid (3.94 g, 92% yield). ^1^H NMR (500 MHz, DMSO-*d*_6_) δ 11.51 (s, 1H), 8.38 (d, *J* = 8.1 Hz, 1H), 8.28-8.22 (series of br s, 2H), 8.03 (d, *J* = 7.8 Hz, 1H), 8.01 (dd, *J* = 7.9, 1.4 Hz, 1H), 7.86 (dd, *J* = 7.7, 7.7 Hz, 1H), 7.69 (ddd, *J* = 8.6, 8.6, 1.5 Hz, 1H), 7.29 (ddd, *J* = 8.1, 8.1, 0.8 Hz, 1H), 3.87 (s, 3H); ^13^C NMR (125 MHz, DMSO-*d*_6_) ppm 167.8, 163.5, 139.3, 135.4, 134.1, 131.0, 130.7, 130.3, 129.6 (q, ^2^*J*_C-F_ = 32 Hz), 128.7 (q, ^3^*J*_C-F_ = 3.6 Hz), 124.1, 123.9 (q, ^3^*J*_C-F_ = 3.9 Hz), 123.8 (q, ^1^*J*_C-F_ = 271 Hz), 121.8, 118.8, 52.8; ^19^F NMR (470 MHz, DMSO-*d*_6_) δ -61.3; HRMS (ESI) m/z: [M+Na]^+^ calcd for C_16_H_12_F_3_NNaO_3_ 346.0661; found 346.0662.

**3**-Amino-2-(3-(trifluoromethyl)phenyl)quinazolin-4(3*H*)-one (S12)**.** Following General Procedure B, ester (1.25 g, 3.87 mmol) in H_2_NNH_2_•H_2_O (7.5 mL) and 1-butanol (7.5 mL) after filtration afforded the product as a free-flowing, colorless solid (1.11 g, 94% yield). ^1^H NMR (500 MHz, DMSO-*d*_6_) δ 8.21 (d, *J* = 7.2 Hz, 1H), 8.17 (br s, 1H), 8.11 (d, *J* = 7.8 Hz, 1H), 7.91-7.82 (series of br m, 2H), 7.78-7.69 (series of br m, 2H), 7.59 (dd, *J* = 7.4, 7.4 Hz, 1H), 5.69 (s, 2H); ^13^C NMR (125 MHz, DMSO-*d*_6_) ppm 161.2, 154.6, 146.6, 135.9, 134.4, 133.7, 128.7, 128.3 (q, ^2^*J*_C-F_ = 32 Hz), 127.5, 127.1, 126.4 (q, ^3^*J*_C-F_ = 3.9 Hz), 126.1, 124.1 (q, ^1^*J*_C-F_ = 271 Hz), 120.3;^[16]^ ^19^F NMR (470 MHz, DMSO-*d*_6_) δ -61.0; HRMS (ESI) m/z: [M+H]^+^ calcd for C_15_H_11_F_3_N_3_O 306.0849; found 306.0849.

*N*-(4-Oxo-2-(3-(trifluoromethyl)phenyl)quinazolin-3(4*H*)-yl)benzamide (29)**.** Following General Procedure C, the amine (100 mg, 328 µmol) and acid chloride (48 µL, 410 µmol) in dimethylacetamide (1.0 mL), after filtration, afforded the amide as a pale yellow solid (29.1 mg, 22% yield). ^1^H NMR (500 MHz, DMSO-*d_6_*) δ 10.17 (s, 1H), 8.36 (d, *J* = 7.9 Hz, 1H), 8.16 (br s, 1H), 8.02 (d, *J* = 7.8 Hz, 1H), 7.92-7.77 (series of m, 3H), 73.72 (d, *J* = 7.8 Hz, 1H), 7.64-7.55 (m, 3H), 7.42 (t, *J* = 7.5 Hz, 1H), 7.31-7.21 (series of m, 2H); ^13^C NMR (125 MHz, DMSO-*d_6_*) ppm 166.0, 161.3, 153.9, 146.6, 135.1, 134.3, 133.6, 132.3, 131.7, 129.9, 128.5, 128.2, 127.6 (q, ^2^*J_C-F_* = 45 Hz), 126.9, 126.7, 126.6 (q, ^3^*J_C-F_* = 3.6 Hz), 126.2, 125.3 (q, ^3^*J_C-F_* = 3.8 Hz), 123.3 (q, ^1^*J_C-F_* = 271 Hz), 120.4; ^19^F NMR (470 MHz, DMSO-*d_6_*) δ -61.0; HRMS (ESI) m/z: [M+H]^+^ calcd for C_22_H_15_F_3_N_3_O_2_ 410.1111; found 410.1113.

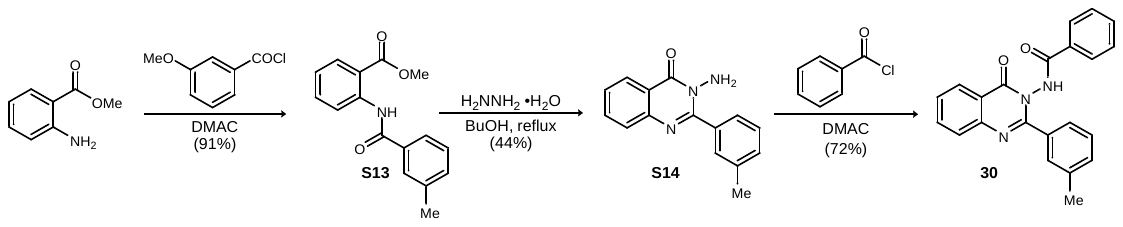

Methyl 2-(3-methylbenzamido)benzoate (S13)**.** Following General Procedure A, methyl anthranilate (1.00 mL, 7.73 mmol) and acid chloride (1.12 mL, 8.50 mmol) in *N,N*-dimethylacetamide (77 mL) after filtration afforded the product as a colorless solid (1.89 g, 91% yield). LRMS (APCI) m/z: [M+H]^+^ calcd for C_16_H_16_NO_3_ 270.1; found 270.1.

3-Amino-2-(*m*-tolyl)quinazolin-4(3*H*)-one (S14). Following General Procedure B, the ester (1.03 g, 3.82 mmol) in H_2_NNH_2_•H_2_O (12 mL) and 1-butanol (5 mL), after filtration, afforded the product as a free-flowing, colorless solid (441 mg, 45% yield). LRMS (APCI) m/z: [M+H]^+^ calcd for C_15_H_14_N_3_O 252.1; found 252.2.

*N*-(4-Oxo-2-(m-tolyl)quinazolin-3(4*H*)-yl)benzamide (30)**.** Following General Procedure C, the amine (100.0 mg, 398 µmol) and acid chloride (56.0 µL, 478 µmol) in dimethylacetamide (4.0 mL), after filtration, afforded the amide as a colorless solid (102 mg, 72% yield). ^1^H NMR (500 MHz, DMSO-*d_6_*) δ 11.72 (s, 1H), 8.22 (dd, *J* = 8.0, 1.0 Hz, 1H), 7.95 (ddd, *J* = 8.3, 8.3, 1.4 Hz, 1H), 7.81 (d, *J* = 8.2 Hz, 1H), 7.72 (d, *J* = 7.3 Hz, 2H), 7.69-7.47 (series of br m, 6H), 7.35 (dd, *J* = 7.6, 7.6 Hz, 1H), 7.30 (br d, *J* = 7.6 Hz, 1H), 2.34 (s, 3H); ^13^C NMR (125 MHz, DMSO-*d_6_*) ppm 165.4, 159.6, 156.4, 146.7, 137.2, 135.4, 133.3, 132.6, 131.3, 130.8, 129.1, 128.7, 127.8, 127.7, 127.5, 127.4, 126.6, 125.6, 120.7, 20.9; HRMS (ESI) m/z: [M+H]^+^ calcd for C_22_H_18_N_3_O_2_ 356.1394 found 356.1395.

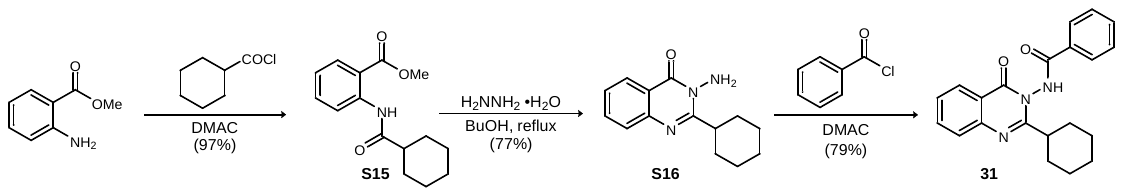

Methyl 2-(cyclohexanecarboxamido)benzoate (S15)**.** Following General Procedure A, methyl anthranilate (1.00 mL, 7.73 mmol) and acid chloride (1.24 mL, 9.28 mmol) in *N,N*-dimethylacetamide (77 mL) after filtration afforded the product as a colorless solid (1.95 g, 97% yield). ^1^H NMR (500 MHz, CDCl_3_) δ11.08 (s, 1H), 8.75 (d, *J* = 8.5 Hz, 1H), 8.02 (d, *J* = 7.0 Hz, 1H), 7.53 (dd, *J* = 7.5, 7.5 Hz, 1H), 7.05 (dd, *J* = 7.5, 7.5 Hz, 1H), 3.93 (s, 3H), 2.33 (tt, *J* = 11.7, 3.4 Hz, 1H), 2.02 (br d, *J* = 12.0 Hz, 2H), 1.89-1.64 (series of br m, 4H), 1.55 (dddd, *J* = 12.1, 12.1, 12.1, 2.6 Hz, 2H), 1.41-1.19 (series of br m, 3H); ^13^C NMR (125 MHz, CDCl_3_) ppm 175.4, 168.8, 141.8, 134.6, 130.8, 122.2, 120.4, 114.8, 52.3, 47.2, 29.6, 25.7;^[17]^ LRMS (APCI) m/z: [M+H]^+^ calcd for C_15_H_20_NO_3_ 262.1; found 262.2.

3-Amino-2-cyclohexylquinazolin-4(3*H*)-one (S16)**.** Following General Procedure B, the ester (1.00 g, 3.83 mmol) in H_2_NNH_2_•H_2_O (12.8 mL) and 1-butanol (26 mL), after filtration, afforded the product as a free-flowing, colorless solid (716 mg, 77% yield). ^1^H NMR (500 MHz, DMSO-*d*_6_) δ 8.10 (d, *J* = 7.6 Hz, 1H), 7.77 (ddd. *J* = 7.6, 7.6, 1.1 Hz, 1H), 7.61 (d, *J* = 8.1 Hz, 1H), 7.47 (dd, *J* = 7.5, 7.5 Hz, 1H), 5.67 (s, 2H), 3.44 (tt, *J* = 11.6, 3.0 Hz, 1H), 1.98 (br d, *J* = 12.1 Hz, 2H), 1.80 (br d, *J* = 12.8 Hz, 2H), 1.71 (br d, *J* = 12.4 Hz, 1H), 1.61-1.44 (br m, 2H), 1.43-1.30 (br m, 2H), 1.30-1.17 (br m, 1H); ^13^C NMR (125 MHz, DMSO-*d*_6_) ppm 162.0, 161.3, 147.0, 134.4, 127.4, 126.4, 126.3, 120.2, 40.4, 30.6, 26.13, 26.08; LRMS (APCI) m/z: [M+H]^+^ calcd for C_14_H_18_N_3_O 244.1; found 244.0.

*N*-(2-Cyclohexyl-4-oxoquinazolin-3(4*H*)-yl)benzamide (31)**.** Following General Procedure C, the amine (100 mg, 411 µmol) and acid chloride (57.3 µL, 493 µmol) in dimethylacetamide (2 mL), after filtration, afforded the amide as a colorless solid (112 mg, 79% yield). ^1^H NMR (500 MHz, DMSO-*d_6_*) δ 11.57 (s, 1H), 8.14 (d, *J* = 7.9 Hz, 1H), 7.99 (d, *J* = 7.3 Hz, 2H), 7.88 (dt, *J* = 7.7, 1.2 Hz, 1H), 7.71 (d, *J* = 7.7 Hz, 1H), 7.70 (br d, *J* = 7.1 Hz, 1H), 7.62 (dd, *J* = 7.7, 7.7 Hz, 2H), 7.56 (t, *J* = 7.5 Hz, 1H), 2.87 (tt, *J* = 11.5, 3.1 Hz, 1H), 2.09 (br d, *J* = 12.7 Hz, 1H), 1.87-1.73 (series of m, 3H), 1.73-1.58 (series of m, 2H), 1.50 (dddd, *J* = 12.7, 12.7, 12.7, 3.3 Hz, 1H), 1.39-1.15 (series of m, 3H); ^13^C NMR (125 MHz, DMSO-*d_6_*) 166.5, 161.6, 159.1, 146.6, 135.1, 132.8, 131.4, 128.9, 127.7, 127.3, 126.9, 126.4, 120.5, 40.5, 31.0, 29.9, 25.7, 25.42, 25.35; HRMS (ESI) m/z: [M+H]^+^ calcd for C_21_H_22_N_3_O_2_ 348.1707 found 348.1708.

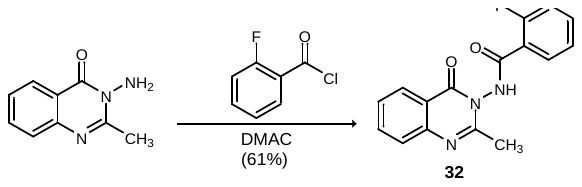

2-Fluoro-N-(2-methyl-4-oxoquinazolin-3(4*H*)-yl)benzamide (32)**.** Following General Procedure A, the commercially available amine (40 mg, 228 µmol) and acid chloride (33 µL, 274 µmol) in dimethylacetamide (2.5 mL), after filtration, afforded the amide as a colorless solid (41.4 mg, 61% yield). ^1^H NMR (500 MHz, DMSO-*d_6_*) δ 11.50 (s, 1H), 8.15 (br d, *J* = 7.2 Hz, 1H), 7.89 (dt, *J* = 7.6, 1.1 Hz, 1H), 7.83 (dt, *J* = 7.2, 1.4 Hz, 1H), 7.74-7.65 (series of m, 2H), 7.57 (t, *J* = 7.6 Hz, 1H), 7.48-7.38 (series of m, 2H);^[18]^ ^13^C NMR (125 MHz, DMSO-*d_6_*) ppm 163.2, 159.5 (d, *J* = 250 Hz), 158.7, 155.9, 146.6, 135.1, 134.2 (d, *J* = 8.6 Hz), 127.0, 126.9, 126.5, 124.9 (d, *J* = 3.4 Hz), 120.63 (d, *J* = 14 Hz), 120.60, 116.6 (d, *J* = 22 Hz), 21.0;^[19]^ ^19^F NMR (470 MHz, DMSO-*d_6_*) δ -112.6^4^; HRMS (ESI) m/z: [M+H]^+^ calcd for C_16_H_13_FN_3_O_2_ 298.0986; found 298.0986.

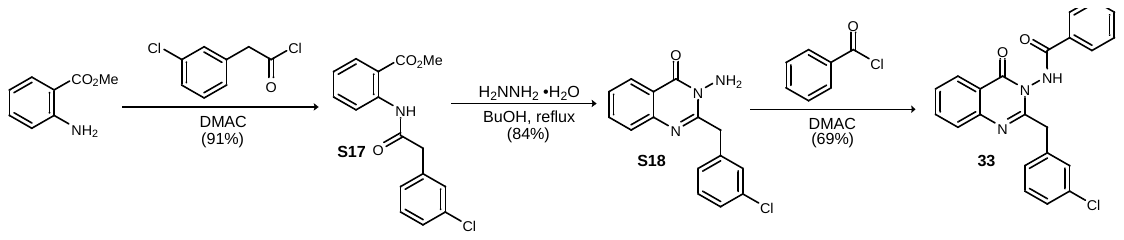

Methyl 2-(2-(3-chlorophenyl)acetamido)benzoate (S17)**.** Following General Procedure A, methyl anthranilate (856 µL, 6.62 mmol) and the acid chloride (1.15 mL, 7.94 mmol) in *N,N*-dimethylacetamide (66 mL), after filtration, afforded the amide as a pale yellow solid (1.83 g, 91% yield). ^1^H NMR (500 MHz, DMSO-*d*_6_) δ 10.62 (s, 1H), 8.17 (d, *J* = 8.2 Hz, 1H), 7.87 (dd, *J* = 7.9, 1.4 Hz, 1H), 7.59 (ddd, *J* = 8.6, 1.5, 1.5 Hz, 1H), 7.45 (s, 1H), 7.41-7.29 (series of m, 3H), 7.19 (ddd, *J* = 7.9, 7.9, 0.8 Hz, 1H), 3.78 (s, 2H), 3.77 (s, 3H); ^13^C NMR (125 MHz, DMSO-*d*_6_) ppm 168.8, 167.4, 139.1, 137.6, 133.8, 133.0, 130.5, 130.2, 129.3, 128.2, 126.8, 123.5, 121.4, 118.5, 25.3, 43.2; HRMS (ESI) m/z: [M+Na]^+^ calcd for C_16_H_14_ClNNaO_3_ 326.0554; found 326.0557.

3-Amino-2-(3-chlorobenzyl)quinazolin-4(3*H*)-one (S18)**.** Following General Procedure B, the ester (1.00 g, 3.29 mmol) in H_2_NNH_2_•H_2_O (6.5 mL) and 1-butanol (6.5 mL), after filtration, afforded the product as a free-flowing, colorless solid (793 mg, 84% yield). ^1^H NMR (500 MHz, DMSO-*d*_6_) δ 8.12 (dd, *J* = 8.0, 0.9 Hz, 1H), 7.78 (ddd, *J* = 8.3, 8.3, 1.3 Hz, 1H), 7.61 (d, *J* = 8.1 Hz, 1H), 7.50 (dd, *J* = 7.3, 7.3 Hz, 1H), 7.39 (br s, 1H), 7.36-7.26 (series of br m, 3H), 5.73 (s, 2H), 4.35 (s, 2H); ^13^C NMR (125 MHz, DMSO-*d*_6_) ppm 160.7, 156.7, 146.5, 139.2, 134.1, 132.8, 130.1, 129.0, 127.9, 127.0, 126.5, 126.4, 125.9, 120.0, 39.3; HRMS (ESI) m/z: [M+H]^+^ calcd for C_15_H_13_ClN_3_O 286.0742; found 286.0744.

*N*-(2-(3-Chlorobenzyl)-4-oxoquinazolin-3(4*H*)-yl)benzamide (33)**.** Following General Procedure C, the amine (100 mg, 350 µmol) and acid chloride (51 µL, 438 µmol) in dimethylacetamide (1 mL), after filtration, afforded the amide as a colorless solid (93.3 mg, 69% yield). ^1^H NMR (500 MHz, DMSO-*d_6_*) δ 11.54 (s, 1H), 8.15 (d, *J* = 7.2 Hz, 1H), 7.97-7.85 (series of m, 3H), 7.74 (d, *J* = 8.1 Hz, 1H), 7.67 (t, *J* = 7.4 Hz, 1H), 7.60 (d, *J* = 6.6 Hz, 1H), 7.57 (d, *J* = 7.4 Hz, 2H), 7.34-7.26 (series of m, 3H), 7.26-7.17 (m, 1H), 4.15 (s, 2H); ^13^C NMR (125 MHz, DMSO-*d_6_*) ppm 166.1, 159.1, 156.8, 146.4, 137.9, 135.2, 133.0, 132.7, 131.2, 130.2, 128.8, 128.7, 127.8, 127.6, 127.4, 127.3, 126.8, 126.5, 120.7;^[20]^ HRMS (ESI) m/z: [M+H]^+^ calcd for C_22_H_17_ClN_3_O_2_ 390.1004; found 390.1005.

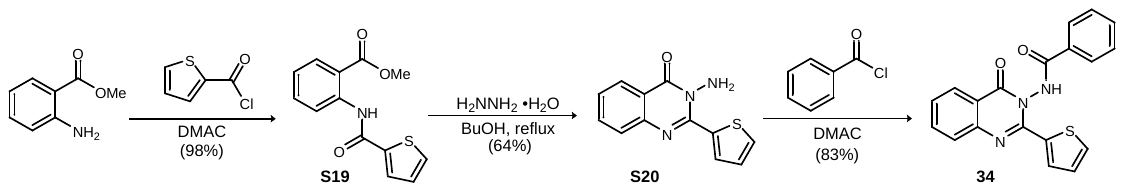

Methyl 2-(thiophene-2-carboxamido)benzoate (S19)**.** Following General Procedure A, methyl anthraniliate (856 µL, 6.62 mmol) and the acid chloride (849 µL, 7.94 mmol) in *N,N*-dimethylacetamide (15 mL) after filtration afforded the product as a colorless solid (1.69 g, 98% yield). ^1^H NMR (500 MHz, DMSO-*d*_6_) δ 11.53 (s, 1H), 8.43 (d, *J* = 8.2 Hz, 1H), 8.00 (dd, *J* = 8.0, 1.4 Hz, 1H), 7.93 (dd, *J* = 5.0, 0.70 Hz, 1H), 7.81 (dd, *J* = 3.7, 0.7 Hz, 1H), 7.67 (ddd, *J* = 8.6, 8.6, 1.4 Hz, 1H), 7.28 (dd, *J* = 4.8, 3.9 Hz, 1H), 7.23 (ddd, *J* = 7.9, 7.9, 0.7 Hz, 1H), 3.89 (s, 3H); ^13^C NMR (125 MHz, DMSO-*d*_6_) ppm 168.0, 159.5, 139.8, 139.4, 134.3, 132.6, 130.7, 129.0, 128.5, 123.4, 120.9, 117.1, 52.6; HRMS (ESI) m/z: [M+Na]^+^ calcd for C_13_H_11_NNaO_3_S 284.0352; found 284.0354.

3-Amino-2-(thiophen-2-yl)quinazolin-4(3*H*)-one (S20)**.** Following General Procedure B, the ester (1.00 g, 3.83 mmol) in H_2_NNH_2_•H_2_O (7.6 mL) and 1-butanol (7.6 mL), after filtration, afforded the product as a free-flowing, colorless solid (593 mg, 64% yield). ^1^H NMR (500 MHz, DMSO-*d*_6_) δ 8.42 (dd, *J* = 3.8, 1.0 Hz, 1H), 8.13 (dd, *J* = 8.0, 1.0 Hz, 1H), 7.87 (dd, *J* = 5.1, 1.0 Hz, 1H), 7.82 (ddd, *J* = 8.3, 8.3, 1.4 Hz, 1H), 7.68 (d, *J* = 8.2 Hz, 1H), 7.49 (br dd, *J* = 7.3, 7.3 Hz, 1H), 7.22 (dd, *J* = 4.9, 4.0 Hz, 1H), 5.97 (s, 2H); ^13^C NMR (125 MHz, DMSO-*d*_6_) ppm 161.3, 149.6, 146.9, 135.2, 134.4, 133.9, 133.7, 127.2, 127.1, 126.2, 126.0, 19.3; HRMS (ESI) m/z: [M+H]^+^ calcd for C_12_H_10_N_3_OS 244.0539; found 244.0539.

*N*-(4-Oxo-2-(thiophen-2-yl)quinazolin-3(4*H*)-yl)benzamide (34)**.** Following General Procedure C, the amine (100.0 mg, 411 µmol) and acid chloride (60.0 µL, 514 µmol) in dimethylacetamide (1.0 mL), after filtration, afforded the amide as a colorless solid (119 mg, 83% yield). ^1^H NMR (500 MHz, DMSO-*d_6_*) δ 11.94 (s, 1H), 8.17 (br dd, *J* = 7.9, 0.9 Hz, 1H), 8.09 (dd, *J* = 3.8 Hz, 0.9 Hz, 1H), 8.05 (br d, *J* = 7.3 Hz, 2H), 7.97-7.89 (m, 1H), 7.87 (dd, *J* = 5.0, 0.9 Hz, 1H), 7.78 (d, *J* = 8.1 Hz, 1H), 7.72 (br dd, *J* = 7.4, 7.4 Hz, 1H), 7.63 (t, *J* = 7.8 Hz, 2H), 7.59 (t, *J* = 7.8 Hz, 1H), 7.21 (dd, *J* = 4.9, 4.1 Hz, 1H); ^13^C NMR (125 MHz, DMSO-*d_6_*) ppm 166.3, 159.4, 149.6, 146.7, 135.5, 134.1, 133.01, 132.95, 132.7, 131.0, 128.9, 127.89, 127.88, 127.5, 127.2, 126.6, 120.0; HRMS (ESI) m/z: [M+H]^+^ calcd for C_19_H_14_N_3_O_2_S 348.0801; found 348.0803.

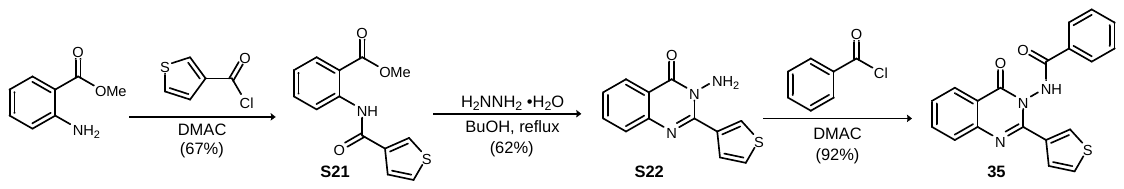

Methyl 2-(thiophene-3-carboxamido)benzoate (S21)**.** Following General Procedure A, methyl anthranilate (856 µL, 6.62 mmol) and the acid chloride (1.16 g, 7.94 mmol) in *N,N*-dimethylacetamide (66 mL) after filtration afforded the product as a colorless solid (1.16 g, 67% yield). ^1^H NMR (500 MHz, DMSO-*d*_6_) δ 11.41 (s, 1H), 8.50 (d, *J* = 8.2 Hz, 1H), 8.29 (dd, *J* = 2.8, 1.2 Hz, 1H), 8.00 (dd, *J* = 8.0, 1.4 Hz, 1H), 7.74 (dd, *J* = 5.0, 3.0 Hz, 1H), 7.67 (ddd, *J* = 8.6, 8.6, 1.4 Hz, 1H), 7.56 (dd, *J* = 5.0, 1.1 Hz, 1H), 7.23 (ddd, *J* = 8.0, 8.0, 0.7 Hz, 1H), 3.83 (s, 3H); ^13^C NMR (125 MHz, DMSO-*d*_6_) ppm 168.0, 160.4, 140.2, 137.6, 134.3, 130.7, 130.2, 128.1, 126.1, 123.3, 120.8, 117.0, 52.6; HRMS (ESI) m/z: [M+Na]^+^ calcd for C_13_H_11_NNaO_3_S 284.0352; found 284.0354.

3-Amino-2-(thiophen-3-yl)quinazolin-4(3*H*)-one (S22)**.** Following General Procedure B, the ester (1.00 g, 3.83 mmol) in H_2_NNH_2_•H_2_O (7.6 mL) and 1-butanol (7.6 mL), after filtration, afforded the product as a free-flowing, colorless solid (575 mg, 62% yield). ^1^H NMR (500 MHz, DMSO-*d*_6_) δ 8.49 (dd, *J* = 2.9, 0.9 Hz, 1H), 8.16 (dd, *J* = 7.9, 0.9 Hz, 1H), 7.86-7.79 (series of m, 2H), 7.70 (d, *J* = 8.2 Hz, 1H), 7.61 (dd, *J* = 5.1, 3.1 Hz, 1H), 7.53 (dd, *J* = 7.3, 7.3 Hz, 1H), 5.86 (s, 2H); ^13^C NMR (125 MHz, DMSO-*d*_6_) ppm 161.2, 150.6, 146.8, 135.0, 134.3, 131.0, 130.0, 127.3, 126.5, 126.0, 124.8, 119.6; HRMS (ESI) m/z: [M+H]^+^ calcd for C_12_H_10_N_3_OS 244.0539; found 244.0540.

*N*-(4-Oxo-2-(thiophen-3-yl)quinazolin-3(4*H*)-yl)benzamide (35)**.** Following General Procedure C, the amine (100.0 mg, 411 µmol) and acid chloride (60.0 µL, 514 µmol) in dimethylacetamide (1.0 mL), after filtration, afforded the amide as a colorless solid (132 mg, 92% yield). ^1^H NMR (500 MHz, DMSO-*d_6_*) δ 11.79 (s, 1H), 8.22-8.16 (br m, 2H), 7.93 (br td, *J* = 7.7, 1.3 Hz, 1H), 7.88 (br d, *J* = 7.3 Hz, 2H), 7.79 (d, *J* = 8.1 Hz, 1H), 7.68-7.58 (series of br m, 4H), 7.56 (br t, *J* = 7.6 Hz, 2H); ^13^C NMR (125 MHz, DMSO-*d_6_*) ppm 165.7, 159.6, 151.7, 146.7, 135.4, 133.8, 132.7, 131.2, 129.8, 128.8, 128.2, 127.7, 127.6, 127.4, 126.6, 126.3, 120.5; HRMS (ESI) m/z: [M+H]^+^ calcd for C_19_H_14_N_3_O_2_S 348.0801; found 348.0804.

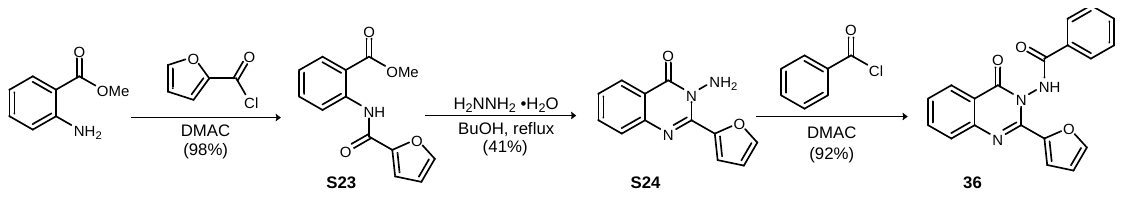

Methyl 2-(furan-2-carboxamido)benzoate (S23)**.** Following General Procedure A, methyl anthranilate (856 µL, 6.62 mmol) and the acid chloride (783 µL, 7.94 mmol) in *N,N*-dimethylacetamide (15 mL) after filtration afforded the product as a colorless solid (1.59 g, 98% yield). ^1^H NMR (500 MHz, DMSO-*d*_6_) δ 11.79 (s, 1H), 8.63 (d, *J* = 8.3 Hz, 1H), 8.05-8.00 (series of m, 2H), 7.68 (ddd, *J* = 8.6, 8.6, 1.4 Hz, 1H), 7.31 (d, *J* = 3.4 Hz, 1H), 7.23 (ddd, *J* = 8.0, 8.0, 0.8 Hz, 1H), 6.76 (dd, *J* = 3.4, 1.7 Hz, 1H), 3.92 (s, 3H); ^13^C NMR (125 MHz, DMSO-*d*_6_) ppm 167.9, 155.8, 147.3, 146.3, 140.1, 134.6, 130.8, 123.2, 120.2, 115.9, 115.7, 112.8, 52.7; HRMS (ESI) m/z: [M+Na]^+^ calcd for C_13_H_11_NNaO_4_ 268.0580; found 268.0583.

3-Amino-2-(furan-2-yl)quinazolin-4(3H)-one (S24)**.** Following General Procedure B, the ester (1.00 g, 4.08 mmol) in H_2_NNH_2_•H_2_O (8.0 mL) and 1-butanol (8.0 mL), after filtration, afforded the product as a free-flowing, colorless solid (377 mg, 41% yield). ^1^H NMR (500 MHz, DMSO-*d*_6_) δ 8.15 (br d, *J* = 7.8 Hz, 1H), 8.01 (br s, 1H), 7.83 (ddd, *J* = 8.3, 1.3 Hz, 1H), 7.78 (d, *J* = 3.4 Hz, 1H), 7.73 (d, *J* = 8.1 Hz, 1H), 7.53 (dd, *J* = 7.4 Hz, 1H), 6.75 (dd, *J* = 3.4, 1.7 Hz, 1H), 5.92 (s, 2H); ^13^C NMR (125 MHz, DMSO-*d*_6_) ppm 160.8, 146.8, 146.0, 145.4, 145.3, 134.4, 127.3, 126.6, 126.0, 119.7, 119.4, 112.2; HRMS (ESI) m/z: [M+H]^+^ calcd for C_12_H_10_N_3_O_2_ 228.0768; found 228.0768.

*N*-(2-(Furan-2-yl)-4-oxoquinazolin-3(4*H*)-yl)benzamide (36)**.** Following General Procedure C, the amine (100.0 mg, 440 µmol) and acid chloride (64 µL, 550 µmol) in dimethylacetamide (1.0 mL), after filtration, afforded the amide as a pale yellow solid (134 mg, 92% yield). ^1^H NMR (500 MHz, DMSO-*d_6_*) δ 11.87 (s, 1H), 8.19 (d, *J* = 7.1 Hz, 1H), 8.09-7.97 (series of m, 3H), 7.94 (br dt, *J* = 7.7, 1.3 Hz, 1H), 7.82 (d, *J* = 8.1 Hz, 1H), 7.70 (br dd, *J* = 7.4 Hz, 1H), 7.66-7.56 (m, 3H)^[21]^, 7.29 (d, *J* = 3.5 Hz, 1H), 6.70 (dd, *J* = 3.5, 1.3 Hz, 1H); ^13^C NMR (125 MHz, DMSO-*d_6_*) ppm 165.8, 159.3, 146.7, 146.6, 145.9, 144.7, 135.5, 132.9, 131.1, 128.9, 127.73, 127.71, 127.5, 126.6, 120.4, 117.0, 112.4; HRMS (ESI) m/z: [M+H]^+^ calcd for C_19_H_14_N_3_O_3_ 332.1030; found 332.1031.

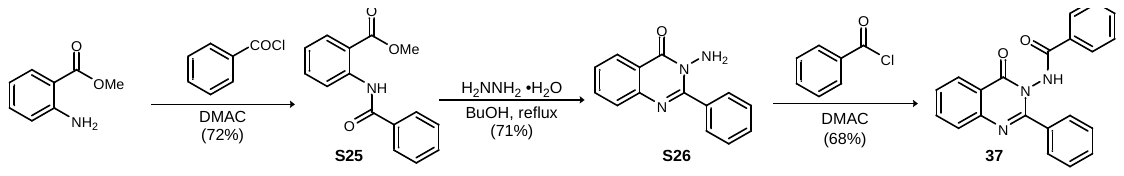

Methyl 2-benzamidobenzoate (25)**.** Following General Procedure A, methyl anthranilate (1.00 mL, 7.73 mmol) and acid the chloride (1.08 mL, 9.28 mmol) in *N,N*-dimethylacetamide (77 mL) after filtration afforded the product as a colorless solid (1.42 g, 84% yield). ^1^H NMR (500 MHz, CDCl_3_) δ12.04 (s, 1H), 8.94 (d, *J* = 8.4 Hz, 1H), 8.12-7.99 (series of m, 3H), 7.61 (ddd, *J* = 7.9, 7.9, 1.5 Hz, 1H), 7.59-7.49 (series of m, 3H), 7.13 (dt, *J* = 7.7, 0.8 Hz, 1H), 3.97 (s, 3H); ^13^C NMR (125 MHz, DMSO-*d*_6_) ppm 169.0, 165.7, 141.8, 134.85, 134.81, 131.9, 130.9, 128.8, 127.3, 122.6, 120.4, 115.1, 52.5; LRMS (APCI) m/z: [M+H]^+^ calcd for C_15_H_14_NO_3_ 256.1; found 256.0.

3-Amino-2-phenylquinazolin-4(3*H*)-one (26)**.** Following General Procedure B, the ester (850 mg, 3.33 mmol) in H_2_NNH_2_•H_2_O (11 mL) and 1-butanol (22 mL), after filtration, afforded the product as a free-flowing, colorless solid (558 mg, 71% yield). ^1^H NMR (500 MHz, DMSO-*d*_6_) δ8.19 (d, *J* = 8.0 Hz, 1H), 7.88-7.77 (series of m, 3H), 7.71 (d, *J* = 8.1 Hz, 1H), 7.57 (t, *J* = 7.4 Hz, 1H), 7.53-7.44 (series of m, 3H), 5.67 (s, 2H); ^13^C NMR (125 MHz, DMSO-*d*_6_) ppm 161.6, 156.2, 147.1, 135.3, 134.7, 130.0, 129.9, 127.84, 127.80, 127.2, 126.4, 120.5; LRMS (APCI) m/z: [M+H]^+^ calcd for C_14_H_12_N_3_O 238.1; found 238.0.

N-(4-Oxo-2-phenylquinazolin-3(4*H*)-yl)benzamide (37)**.** Following General Procedure C, the amine (100.0 mg, 422 µmol) and acid chloride (58.8 µL, 506 µmol) in dimethylacetamide (2 mL), after filtration, afforded the amide as a colorless solid (97.9 mg, 68% yield). ^1^H NMR (500 MHz, DMSO-*d_6_*) δ 11.74 (s, 1H), 8.22 (d, *J* – 7.9 Hz, 1H), 7.95 (dt, *J* = 7.1, 1.1 Hz, 1H), 7.81 (d, *J* = 8.1 Hz, 1H), 7.75-7.68 (series of m, 4H), 7.65 (t, *J* = 7.4 Hz, 1H), 7.60 (t, *J* = 7.5 Hz, 1H), 7.53-7.42 (series of m, 5H); ^13^C NMR (125 MHz, DMSO-*d_6_*) ppm 165.45, 159.6, 156.3, 146.6, 135.4, 133.4, 132.6, 131.2, 130.2, 128.7, 128.5, 127.9, 127.8, 127.6, 127.4, 126.6, 120.8; HRMS (ESI) m/z: [M+H]^+^ calcd for C_21_H_16_N_3_O_2_ 342.1237; found 342.1238.

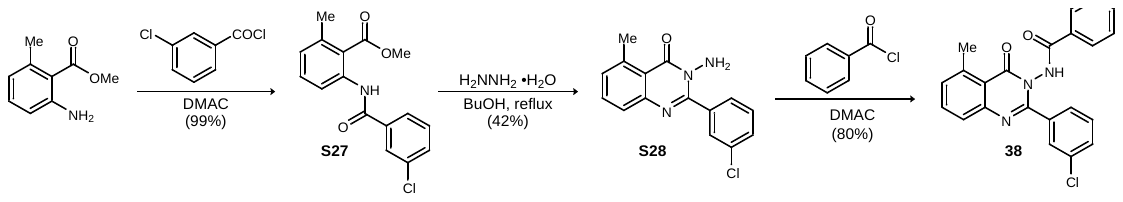

Methyl 2-(3-chlorobenzamido)-6-methylbenzoate (S27)**.** Following General Procedure A, the amine (500 mg, 3.03 mmol) and acid chloride (466 µL, 3.64 mmol) in *N,N*-dimethylacetamide (30 mL) after filtration afforded the product as a colorless solid (910 mg, 99% yield). LRMS (APCI) m/z: [M+H]^+^ calcd for C_16_H_15_ClNO_3_ 304.1; found 304.0.

3-Amino-2-(3-chlorophenyl)-5-methylquinazolin-4(3*H*)-one (S28)**.** Following General Procedure B, the ester (710 mg, 2.34 mmol) in H_2_NNH_2_•H_2_O (7.8 mL) and 1-butanol (11.7 mL), after filtration, afforded the product as a free-flowing, colorless solid (284 mg, 42% yield). LRMS (APCI) m/z: [M+H]^+^ calcd for C_15_H_13_ClN_3_O 286.1; found 286.1.

*N*-(2-(3-Chlorophenyl)-5-methyl-4-oxoquinazolin-3(4*H*)-yl)benzamide (38)**.** Following General Procedure C, the amine (100.0 mg, 350 µmol) and acid chloride (49.0 µL, 420 µmol) in dimethylacetamide (3.5 mL), after filtration, afforded the amide as a colorless solid (109 mg, 80% yield). ^1^H NMR (500 MHz, DMSO-*d_6_*) δ 11.64 (s, 1H), 7.83-7.70 (series of m, 4H), 7.68 (br d, *J* = 7.7 Hz, 1H), 7.65-7.58 (series of m, 2H), 7.58-7.47 (series of m, 4H), 7.42 (d, *J* = 7.4 Hz, 1H), 2.80 (s, 3H); ^13^C NMR (125 MHz, DMSO-*d_6_*) ppm 165.7, 159.7, 154.5, 147.9, 140.7, 135.2, 134.5, 132.7, 132.5, 131.1, 130.13, 130.09, 129.9, 128.8, 128.2, 127.4, 127.1, 126.1, 119.2, 22.4; HRMS (ESI) m/z: [M+H]^+^ calcd for C_22_H_17_ClN_3_O_2_ 390.1004; found 390.1007.

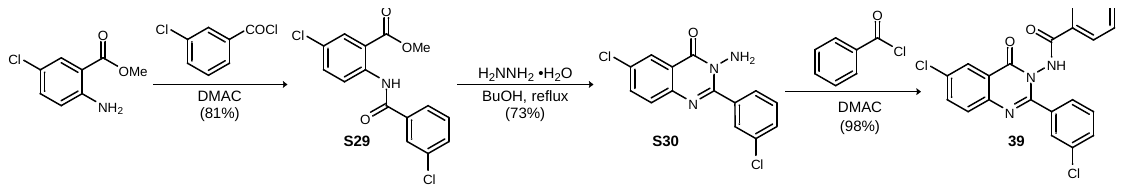

Methyl 5-chloro-2-(3-chlorobenzamido)benzoate (S29)**.** Following General Procedure A, the amine (1.00 g, 5.39 mmol) and acid chloride (823 µL, 6.47 mmol) in *N,N*-dimethylacetamide (54 mL) after filtration afforded the product as a colorless solid (1.42 g, 81% yield). ^1^H NMR (500 MHz, CDCl_3_) δ 11.96 (s, 1H), 8.87 (d, *J* = 9.1 Hz, 1H), 8.05 (d, *J* = 2.6 Hz, 1H), 8.02 (app br s, 1H), 7.87 (d, *J* = 7.8 Hz, 1H), 7.57-7.51 (series of br m, 2H), 7.46 (dd, *J* = 7.8, 7.8 Hz, 1H), 3.98 (s, 3H); ^13^C NMR (125 MHz, CDCl_3_) ppm 168.0, 164.2, 140.1, 136.3, 135.1, 134.7, 132.2, 130.6, 130.1, 128.0, 125.1, 121.9, 116.4, 52.9;^17^ LRMS (APCI) m/z: [M+H]^+^ calcd for C_15_H_12_Cl_2_N_3_O 324.0; found 324.0.

3-Amino-6-chloro-2-(3-chlorophenyl)quinazolin-4(3*H*)-one (S30)**.** Following General Procedure B, the ester (600 mg, 1.85 mmol) in H_2_NNH_2_•H_2_O (6.2 mL) and 1-butanol (12 mL), after filtration, afforded the product as a free-flowing, colorless solid (277 mg, 49% yield). ^1^H NMR (500 MHz, DMSO-*d*_6_) δ 8.13 (d, *J* = 2.5 Hz, 1H), 7.91-7.85 (m, 2H), 7.78-7.72 (m, 2H), 7.62-7.55 (br m, 1H), 7.52 (dd, *J* = 7.9, 7.9 Hz, 1H), 5.69 (s, 2H); ^13^C NMR (125 MHz, DMSO-*d*_6_) ppm 160.7, 155.4, 145.7, 136.9, 135.0, 132.5, 131.7, 130.2, 129.94, 129.86, 128.7, 125.4, 121.9; LRMS (APCI) m/z: [M+H]^+^ calcd for C_14_H_10_Cl_2_N_3_O 306.0; found 306.0.

*N*-(6-Chloro-2-(3-chlorophenyl)-4-oxoquinazolin-3(4*H*)-yl)benzamide (39)**.** Following General Procedure C, the amine (60 mg, 197 µmol) and acid chloride (27.4 µL, 236 µmol) in dimethylacetamide (1.0 mL), after filtration, afforded the amide as a colorless solid (79.2 mg, 98% yield). ^1^H NMR (500 MHz, DMSO-*d_6_*) δ 11.87 (s, 1H), 8.19 (d, *J* = 2.4 Hz, 1H), 8.01 (dd, *J* = 8.7, 2.4 Hz, 1H), 7.95 (br d, *J* = 7.3 Hz, 1H), 7.87 (d, *J* = 8.7 Hz, 1H), 7.79 (br s, 1H), 7.74 (d, *J* = 7.3 Hz, 1H), 7.69 (br d, *J* = 7.7 Hz, 1H), 7.63 (t, *J* = 7.4 Hz, 1H), 7.58 (br d, *J* = 8.8 Hz, 1H), 7.55-7.47 (series of m, 3H); ^13^C NMR (125 MHz, DMSO-*d_6_*) 165.5, 158.6, 155.2, 145.2, 135.6, 134.8, 132.8, 132.6, 132.2, 130.9, 130.4, 130.2, 130.0, 129.2, 128.8, 128.6, 128.3, 127.4, 127.2, 125.6, 122.2; HRMS (ESI) m/z: [M+H]^+^ calcd for C_21_H_14_Cl_2_N_3_O_2_ 410.0458; found 410.0458.

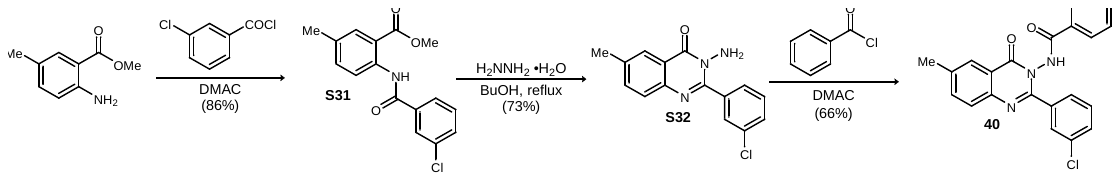

Methyl 2-(3-chlorobenzamido)-5-methylbenzoate (S31)**.** Following General Procedure A, the amine (1.00 g, 6.05 mmol) and acid chloride (929 µL, 7.26 mmol) in *N,N*-dimethylacetamide (61 mL) after filtration afforded the product as a colorless solid (1.58 g, 86% yield). ^1^H NMR (500 MHz, CDCl_3_) δ 11.94 (s, 1H), 8.77 (d, *J* = 8.6 Hz, 1H), 8.04 (br s, 1H), 7.92-7.86 (br m, 2H), 7.52 (br d, *J* = 8.8 Hz, 1H), 7.48-7.39 (series of br m, 2H), 3.96 (s, 3H), 2.36 (s, 3H); ^13^C NMR (125 MHz, CDCl_3_) ppm 169.1, 164.1, 139.1, 136.8, 135.6, 135.0, 132.5, 131.8, 131.1, 130.0, 128.0, 125.0, 120.4, 115.1, 52.5, 20.7; LRMS (APCI) m/z: [M+H]^+^ calcd for C_16_H_15_ClNO_3_ 304.1; found 304.0.

3-Amino-2-(3-chlorophenyl)-6-methylquinazolin-4(3*H*)-one (S32)**.** Following General Procedure B, the ester (600 mg, 1.98 mmol) in H_2_NNH_2_•H_2_O (6.6 mL) and 1-butanol (13 mL), after filtration, afforded the product as a free-flowing, colorless solid (411 mg, 73% yield). ^1^H NMR (500 MHz, DMSO-*d*_6_) δ 7.99 (s, 1H), 7.86 (s, 1H), 7.75 (br d, *J* = 7.6 Hz, 1H), 7.68 (br d, *J* = 7.6 Hz, 1H), 7.63 (d, *J* = 8.3 Hz, 1H), 7.56 (br d, *J* = 8.1 Hz, 1H), 7.51 (dd, *J* = 7.9 Hz, 1H), 5.65 (s, 2H), 2.48 (s, 3H); ^13^C NMR (125 MHz, DMSO-*d*_6_) ppm 161.5, 154.1, 145.1, 137.3, 137.2, 136.2, 132.5, 129.84, 129.79, 129.67, 128.7, 127.8, 125.7, 120.4, 21.3; LRMS (APCI) m/z: [M+H]^+^ calcd for C_15_H_13_ClN_3_O 286.1; found 286.1.

*N*-(2-(3-Chlorophenyl)-6-methyl-4-oxoquinazolin-3(4*H*)-yl)benzamide (40)**.** Following General Procedure C, the amine (100 mg, 350 µmol) and acid chloride (49.0 µL, 420 µmol) in dimethylacetamide (3.5 mL), after filtration, afforded the amide as a colorless solid (109 mg, 80% yield). ^1^H NMR (500 MHz, DMSO-*d_6_*) δ 11.75 (s, 1H), 8.02 (br s, 1H), 7.82-7.43 (series of br m, 11H), 2.49 (s, 3H); ^13^C NMR (125 MHz, DMSO-*d_6_*) 165.9, 159.8, 154.4, 144.9, 138.3, 137.1, 135.7, 133.1, 133.0, 131.5, 130.5, 130.3, 129.7, 129.2, 129.0, 128.7, 128.2, 127.8, 127.6, 126.3, 121.1, 21.3; HRMS (ESI) m/z: [M+H]^+^ calcd for C_22_H_17_ClN_3_O_2_ 390.1004; found 390.1005.

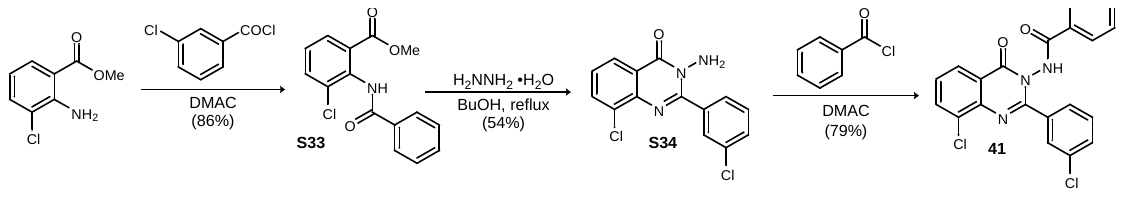

Methyl 2-benzamido-3-chlorobenzoate (S33)**.** Following General Procedure A, the amine (1.00 g, 5.39 mmol) and acid chloride (828 µL, 6.47 mmol) in *N,N*-dimethylacetamide (54 mL) after filtration afforded the product as a colorless solid (1.51 g, 86% yield). LRMS (APCI) m/z: [M+H]^+^ calcd for C_15_H_12_Cl_2_N_3_O 324.0; found 324.1.

3-Amino-8-chloro-2-(3-chlorophenyl)quinazolin-4(3*H*)-one (S34)**.** Following General Procedure B, the ester (1.00 g, 3.08 mmol) in H_2_NNH_2_•H_2_O (10 mL) and 1-butanol (10 mL), after filtration, afforded the product as a free-flowing, colorless solid (479 mg, 54% yield). LRMS (APCI) m/z: [M+H]^+^ calcd for C_14_H_10_Cl_2_N_3_O 306.0; found 306.0.

*N*-(8-Chloro-2-(3-chlorophenyl)-4-oxoquinazolin-3(4*H*)-yl)benzamide (41)**.** Following General Procedure C, the amine (100.0 mg, 327 µmol) and acid chloride (46.0 µL, 392 µmol) in dimethylacetamide (3.3 mL), after filtration, afforded the amide as a colorless solid (106 mg, 79% yield).^[22]^ ^1^H NMR (500 MHz, DMSO-*d_6_*) δ 11.91 (s, 1H), 8.20 (dd, *J* = 7.9, 1.0 Hz, 1H), 8.13 (dd, *J* = 7.9, 1.0 Hz, 1H), 7.95 (br d, *J* = 7.2 Hz, 1H), 7.81 (br s, 1H), 7.78-7.46 (series of m, 8H); ^13^C NMR (125 MHz, DMSO-*d_6_*) 165.5, 159.0, 155.6, 142.9, 135.6, 134.9, 132.8, 132.6, 131.5, 130.9, 130.5, 130.1, 128.8, 128.6, 128.4, 127.5, 127.3, 125.9, 122.7; HRMS (ESI) m/z: [M+H]^+^ calcd for C_21_H_14_Cl_2_N_3_O_2_ 410.0458 found 410.0459.

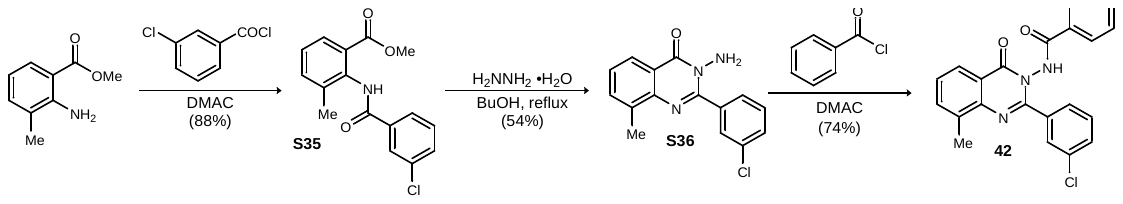

Methyl 2-(3-chlorobenzamido)-3-methylbenzoate (S35)**.** Following General Procedure A, the amine (1.00 g, 6.05 mmol) and acid chloride (929 µL, 7.29 mmol) in *N,N*-dimethylacetamide (61 mL) after filtration afforded the amide as a colorless solid (1.62 g, 88% yield).^[23]^

3-Amino-2-(3-chlorophenyl)-8-methylquinazolin-4(3*H*)-one (S36)**.** Following General Procedure B, the ester (1.18 g, 3.88 mmol) in H_2_NNH_2_•H_2_O (13 mL) and 1-butanol (13 mL), after filtration, afforded the product as a free-flowing, colorless solid (597 mg, 54% yield). ^1^H NMR (500 MHz, DMSO-*d*_6_) δ 8.03 (d, *J* = 7.9 Hz, 1H), 7.91 (br s, 1H), 7.80 (d, *J* = 7.7 Hz, 1H), 7.71 (d, *J* = 7.2 Hz, 1H), 7.58 (d, *J* = 8.3 Hz, 1H), 7.53 (dd, *J* = 7.8, 7.8 Hz, 1H), 7.46 (dd, *J* = 7.6 Hz, 1H), 5.67 (s, 2H), 2.55 (s, 3H); ^13^C NMR (125 MHz, DMSO-*d*_6_) ppm 161.3, 153.3, 145.0, 137.1, 135.7, 134.6, 132.1, 129.5, 129.4, 129.3, 128.5, 126.7, 123.7, 120.2, 17.1; LRMS (APCI) m/z: [M+H]^+^ calcd for C_15_H_13_ClN_3_O 286.1; found 286.1.

*N*-(2-(3-Chlorophenyl)-8-methyl-4-oxoquinazolin-3(4*H*)-yl)benzamide (42)**.** Following General Procedure C, the amine (100.0 mg, 350 µmol) and acid chloride (49.0 µL, 420 µmol) in dimethylacetamide (3.5 mL), after filtration, afforded the amide as a colorless solid (101 mg, 74% yield). ^1^H NMR (500 MHz, DMSO-*d_6_*) δ 11.79 (s, 1H), 8.06 (d, *J* = 7.8 Hz, 1H), 7.82 (br d, *J* = 6.7 Hz, 2H), 7.78-7.67 (series of br m, 3H), 7.62 (br dd, *J* = 7.4, 7.4 Hz, 1H), 7.60-7.45 (series of br m, 5H), 2.60 (s, 3H); ^13^C NMR (125 MHz, DMSO-*d_6_*) 165.5, 159.7, 153.6, 144.9, 136.3, 135.8, 135.5, 132.7, 132.6, 131.1, 130.2, 130.0, 128.8, 128.4, 127.5, 127.4, 127.3, 124.3, 120.9, 17.1; HRMS (ESI) m/z: [M+H]^+^ calcd for C_22_H_17_ClN_3_O_2_ 390.1004 found 390.1006.

### Supplementary Figure 6. ^1^H NMR (500 MHz, DMSO-*d*_6_) of S1

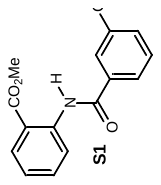

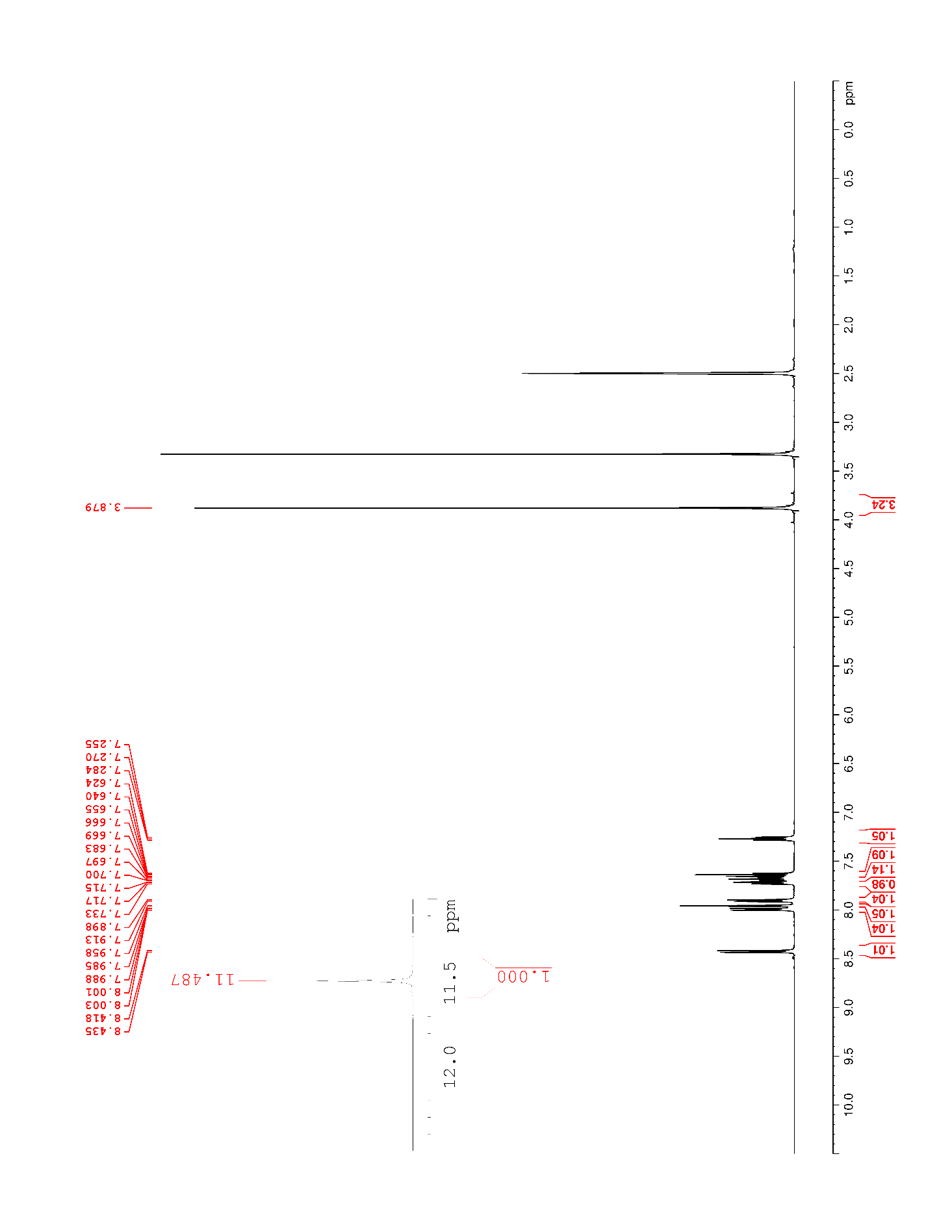

### Supplementary Figure 7. ^13^C NMR (500 MHz, DMSO-*d*_6_) of S1

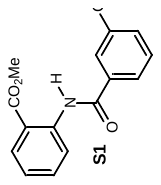

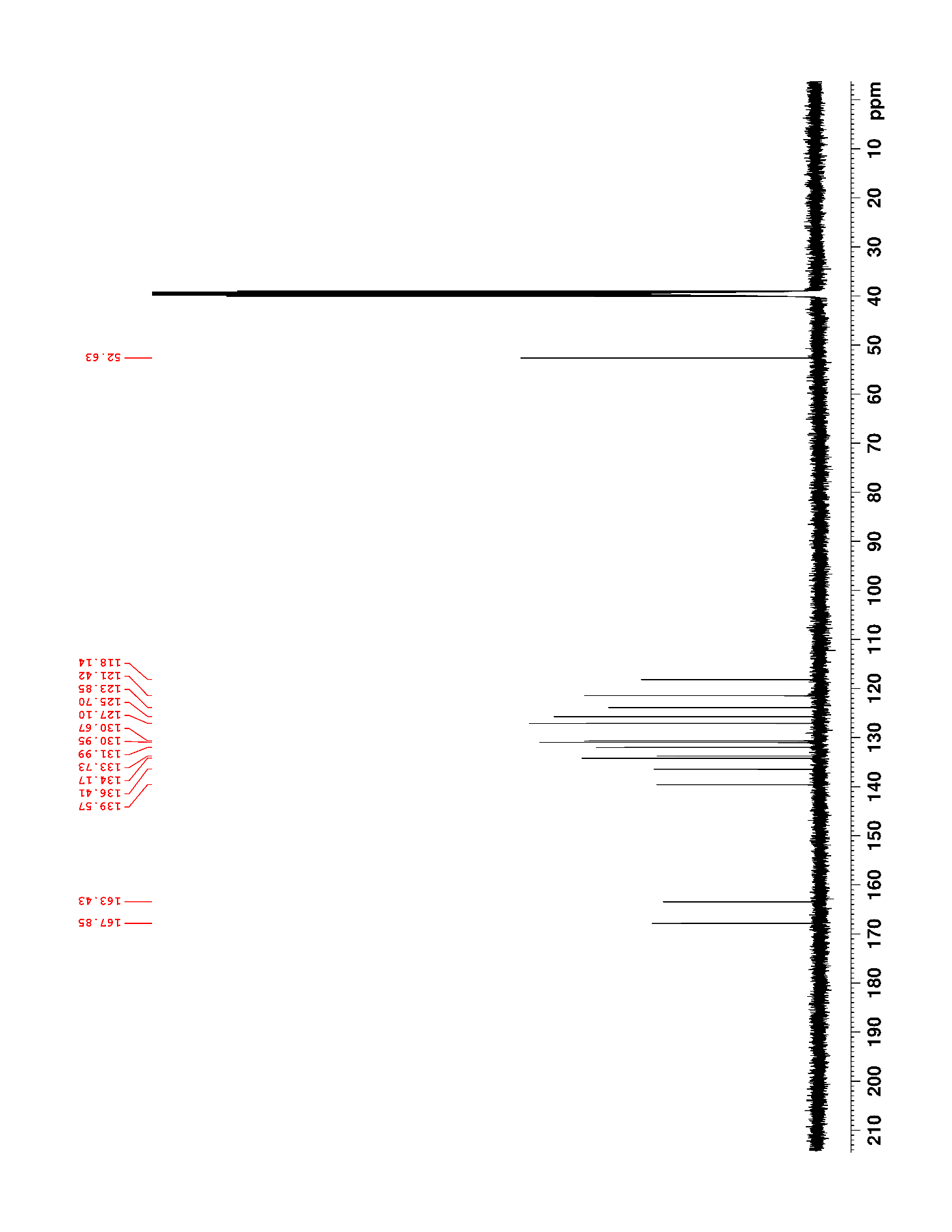

### Supplementary Figure 8. ^1^H NMR (500 MHz, DMSO-*d*_6_) of S2

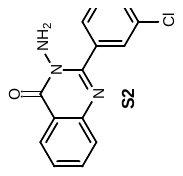

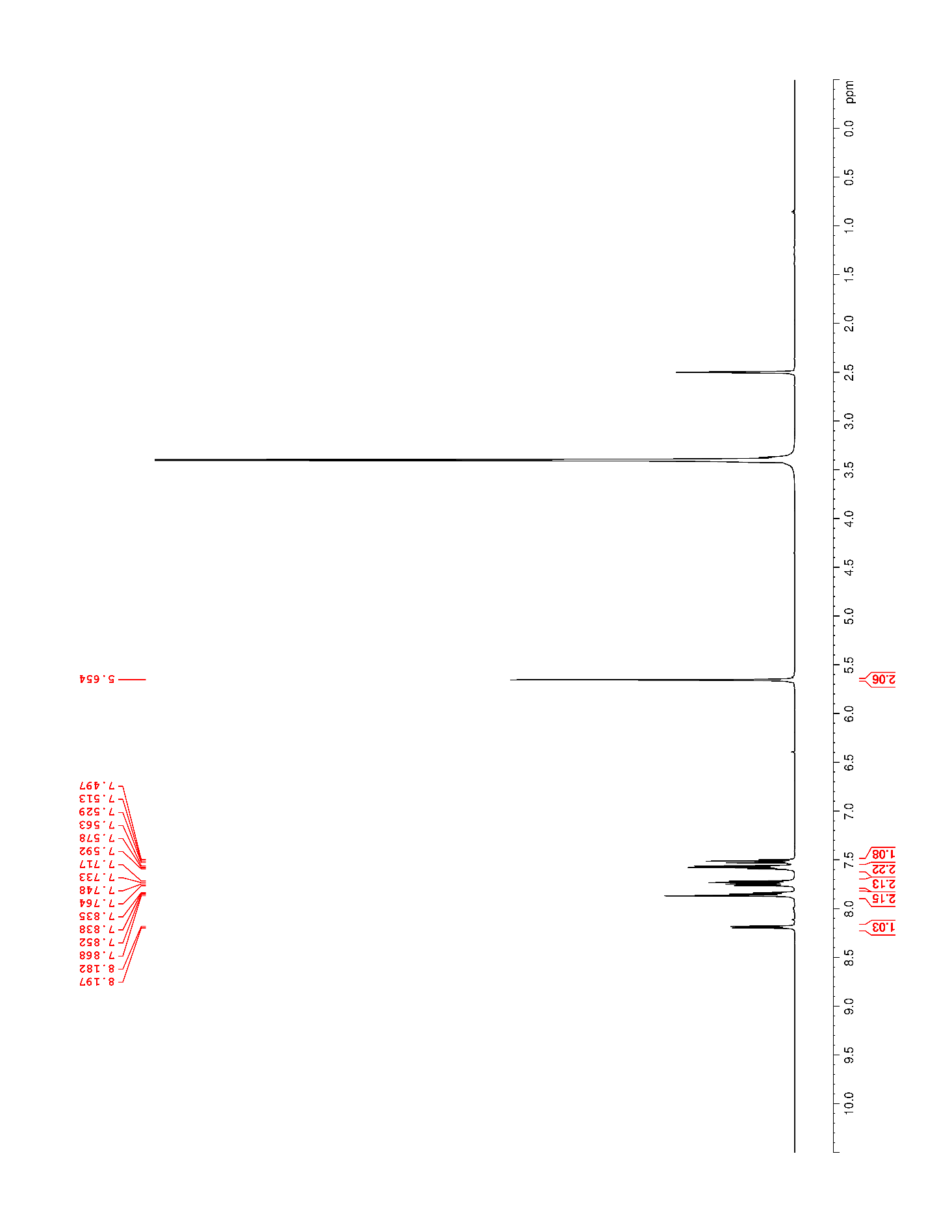

### Supplementary Figure 9. ^13^C NMR (500 MHz, DMSO-*d*_6_) of S2

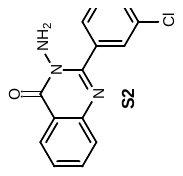

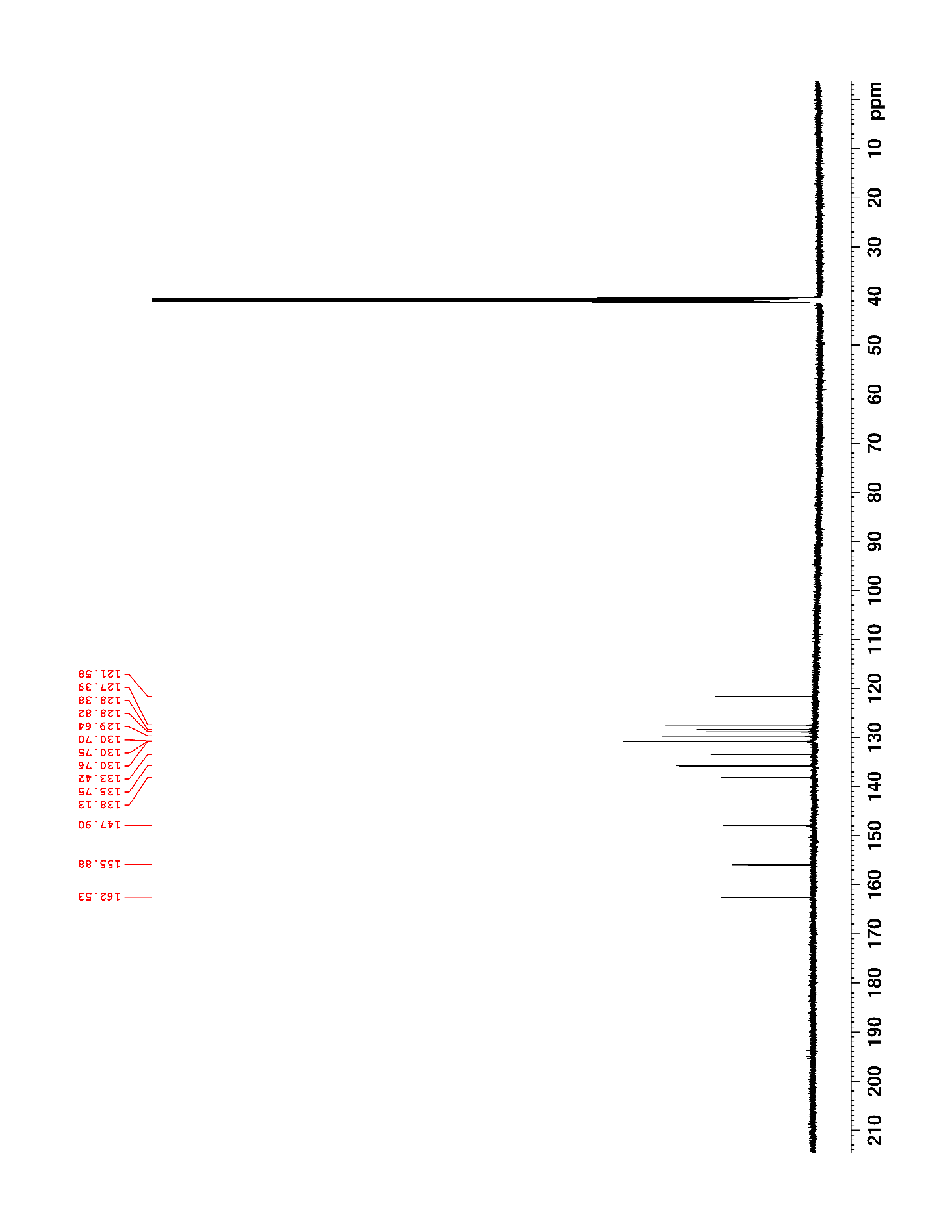

### Supplementary Figure 10. ^1^H NMR (500 MHz, CDCl_3_) of 1

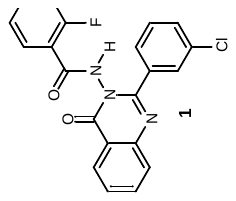

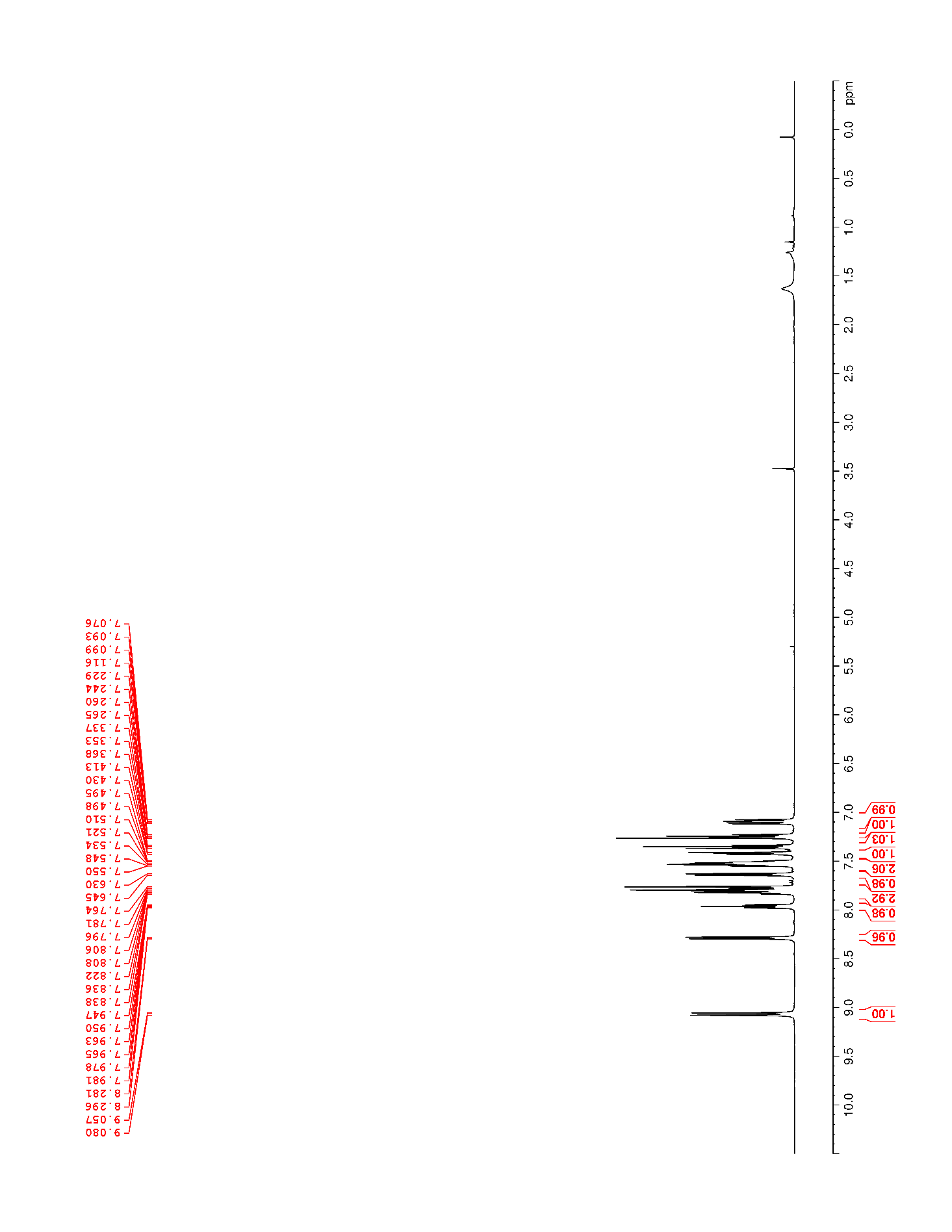

### Supplementary Figure 11. ^13^C NMR (500 MHz, CDCl_3_) of 1

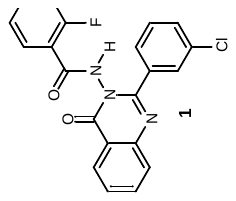

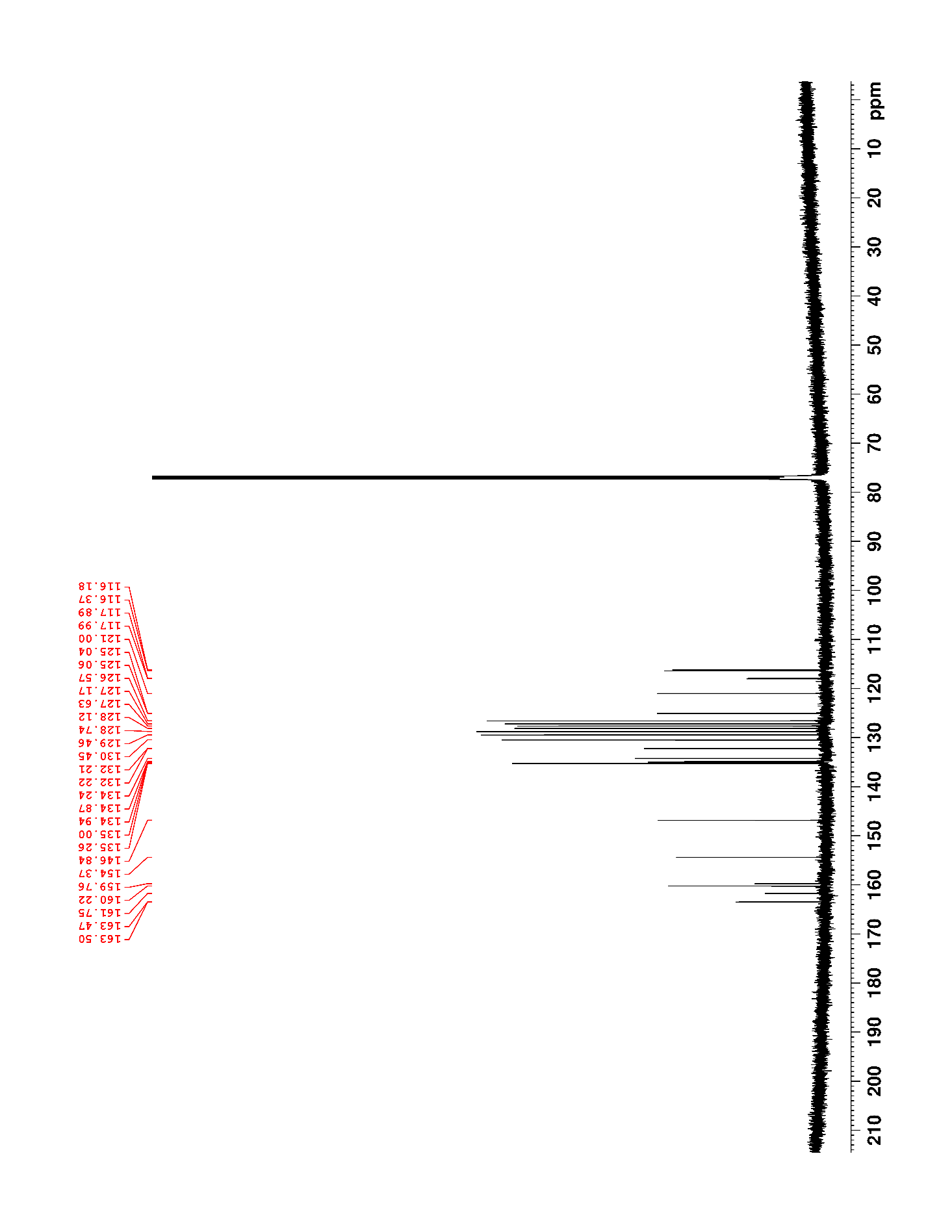

### Supplementary Figure 12. ^19^F NMR (470 MHz, CDCl_3_) of 1

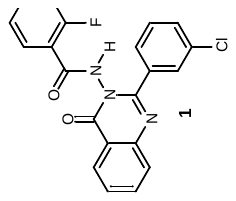

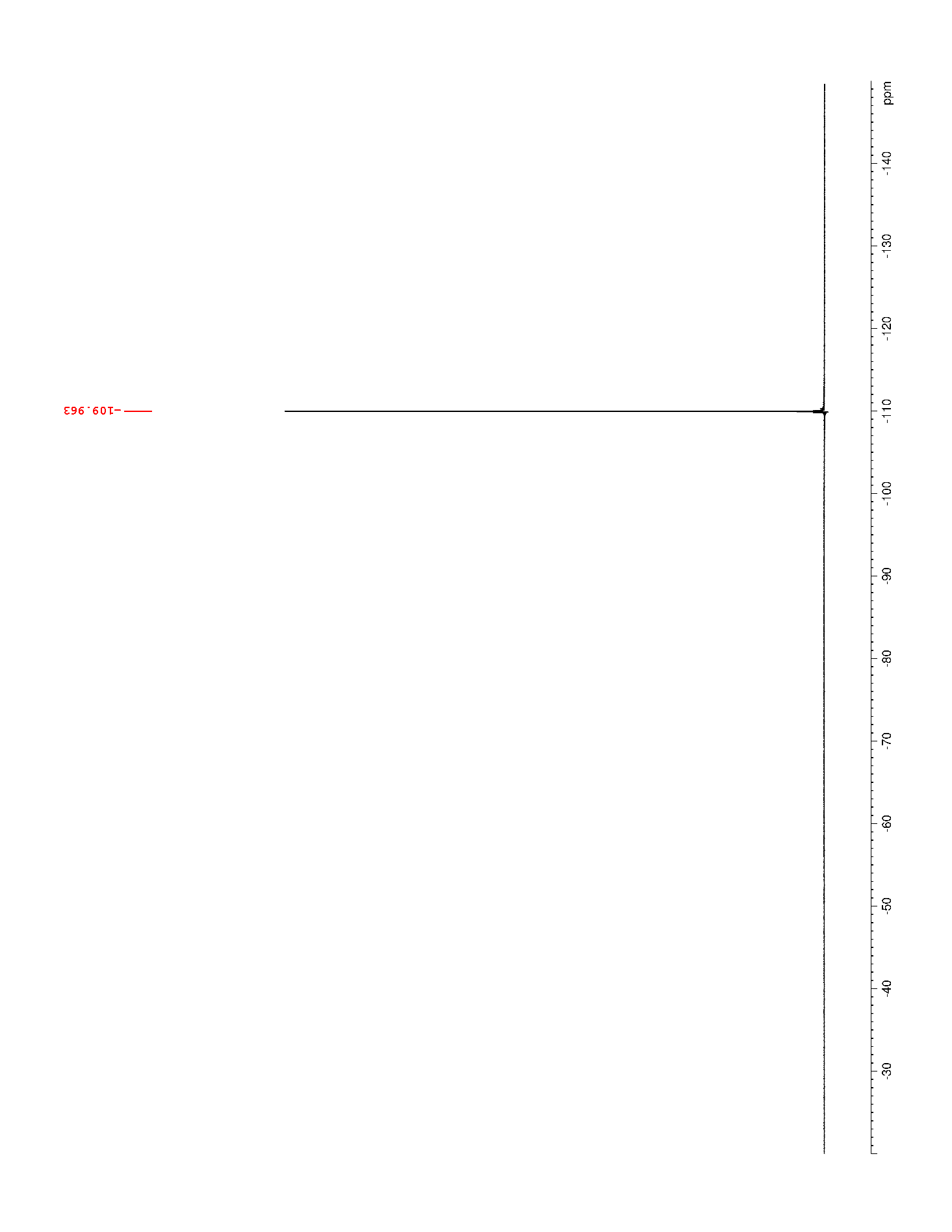

### Supplementary Figure 13. ^1^H NMR (500 MHz, DMSO-*d*_6_) of 2

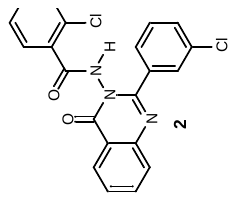

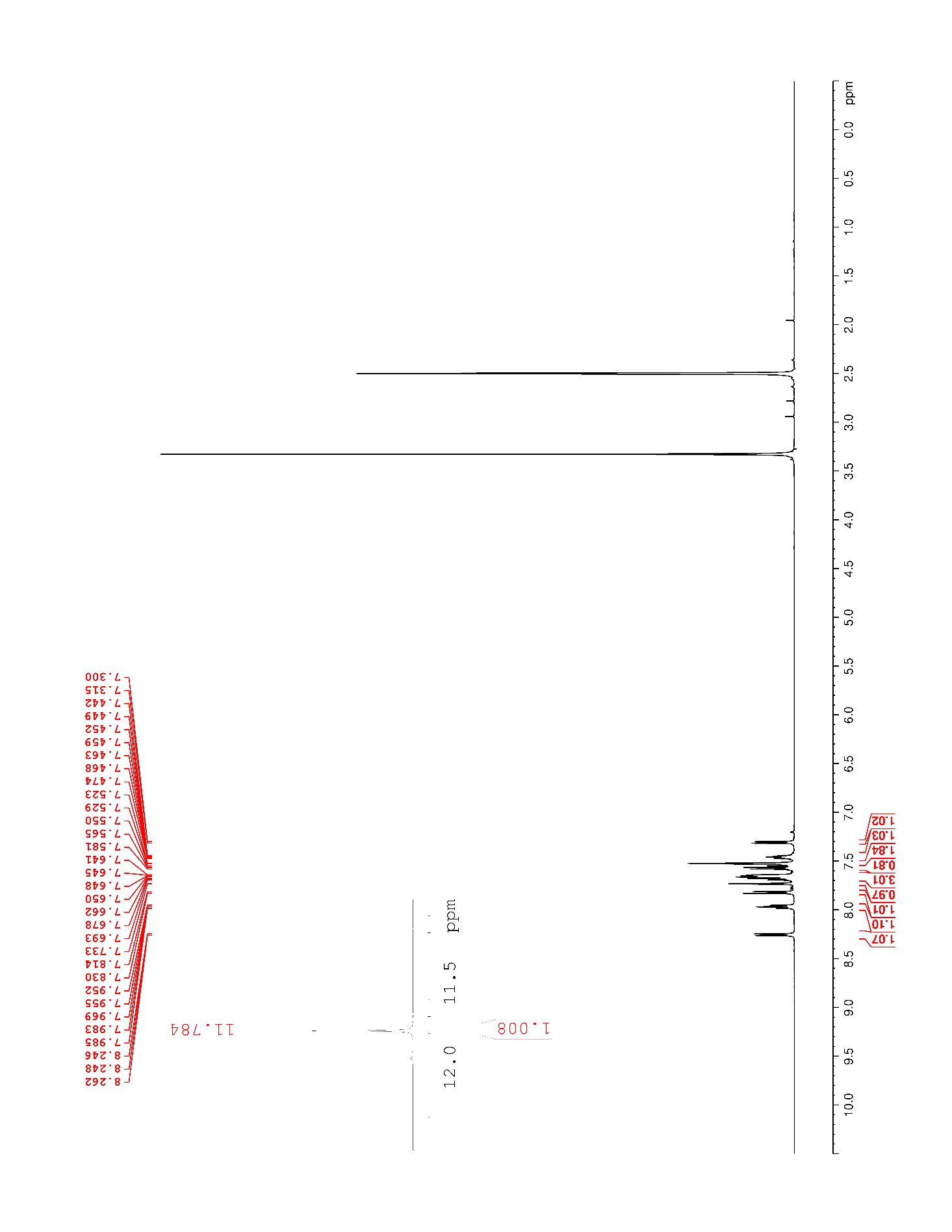

### Supplementary Figure 14. ^13^C NMR (125 MHz, DMSO-*d*_6_) of 2

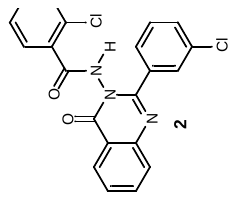

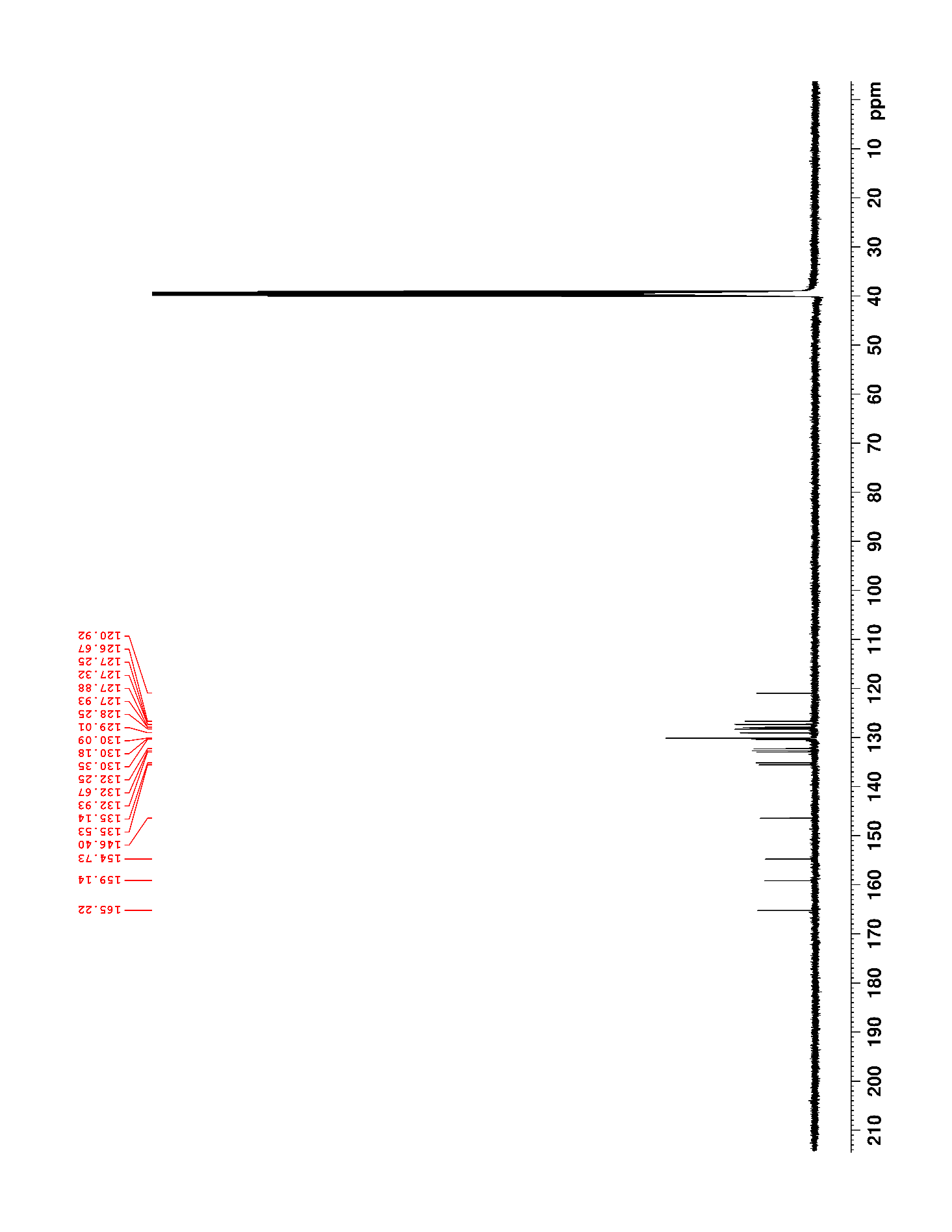

### Supplementary Figure 15. ^1^H NMR (500 MHz, DMSO-*d*_6_) of 3

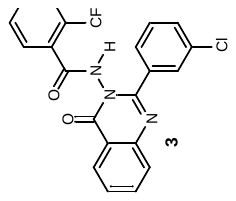

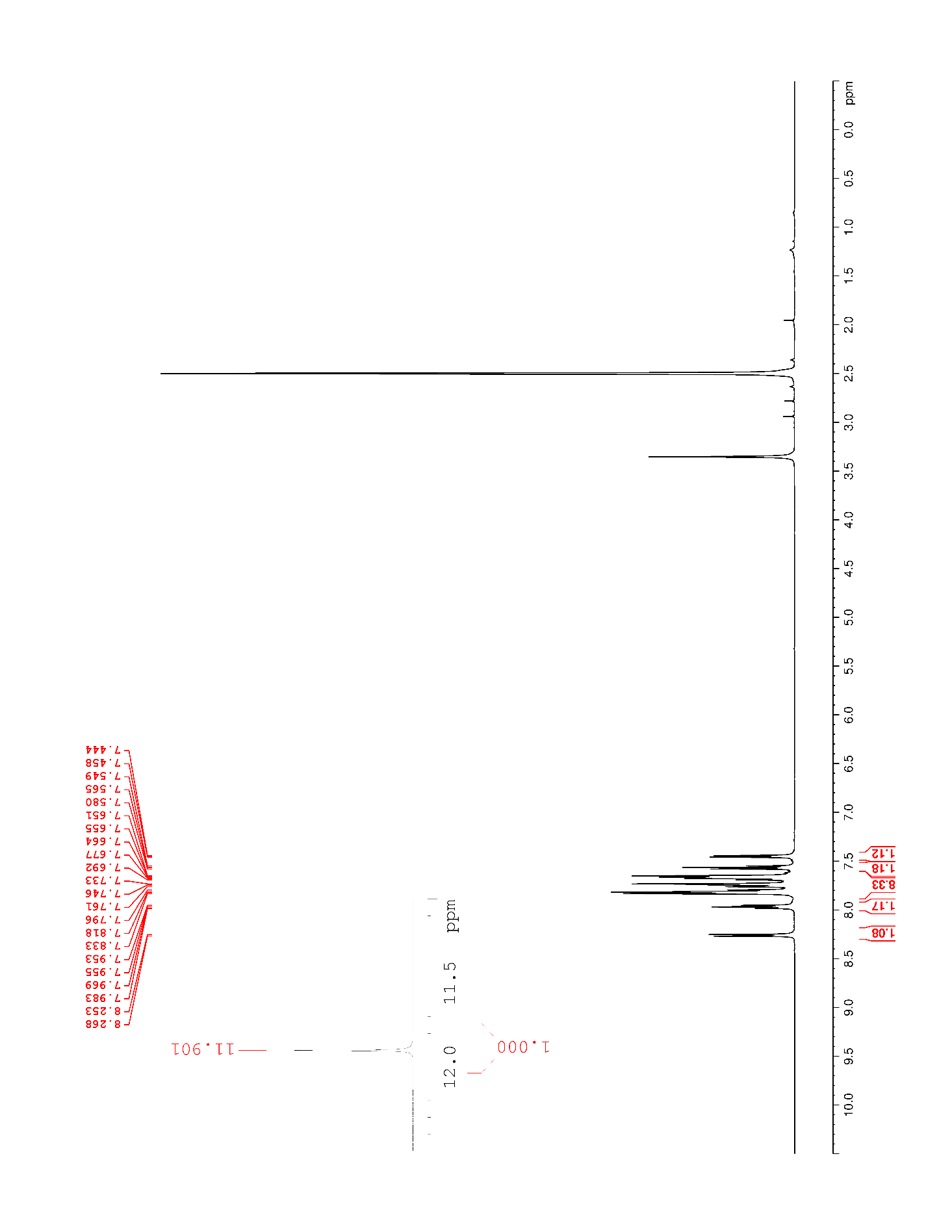

### Supplementary Figure 16. ^13^C NMR (125 MHz, DMSO-*d*_6_) of 3

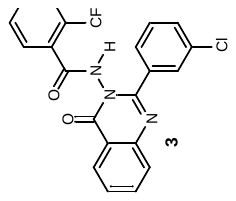

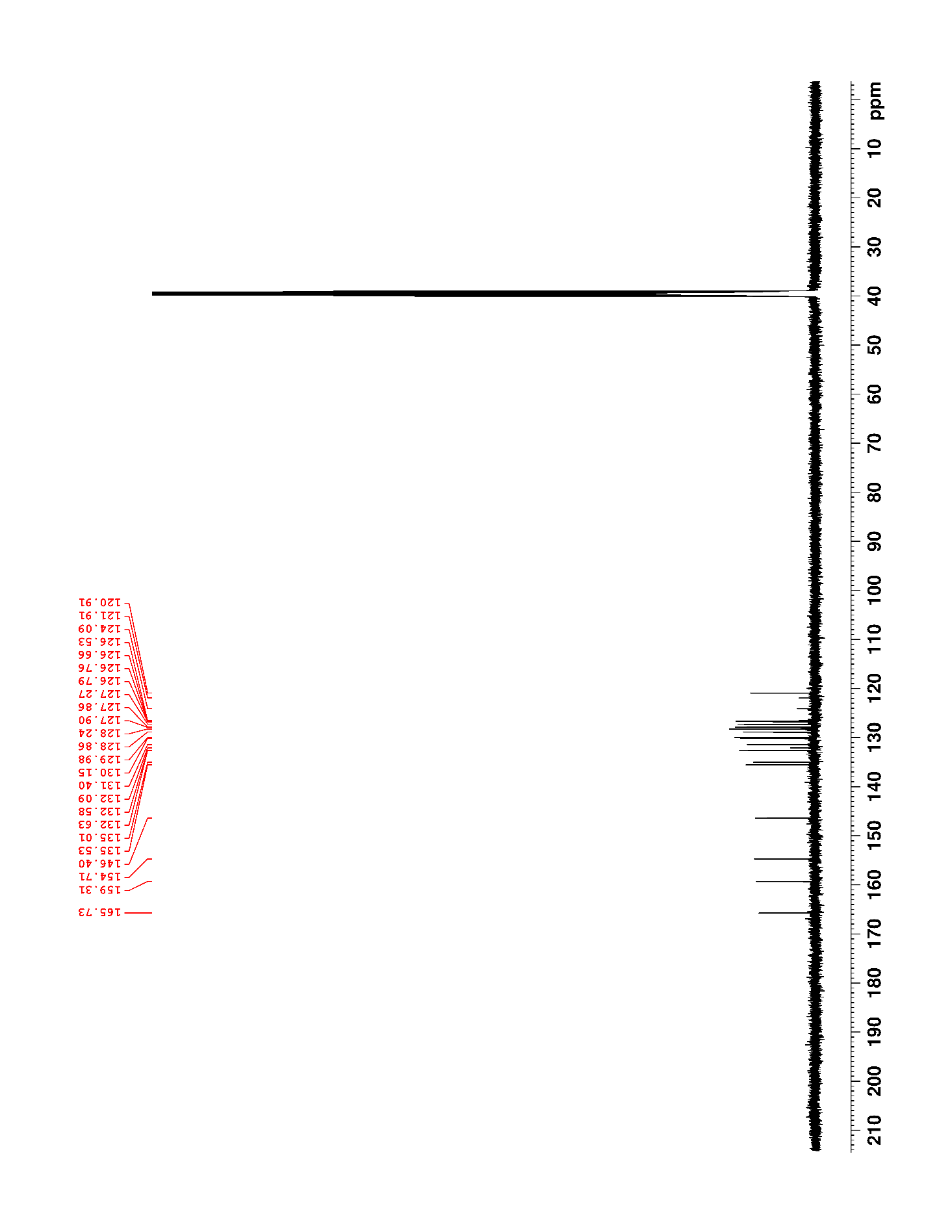

### Supplementary Figure 17. ^19^F NMR (470 MHz, DMSO-*d*_6_) of 3

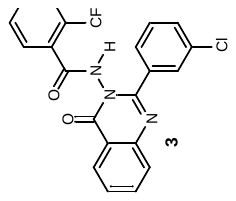

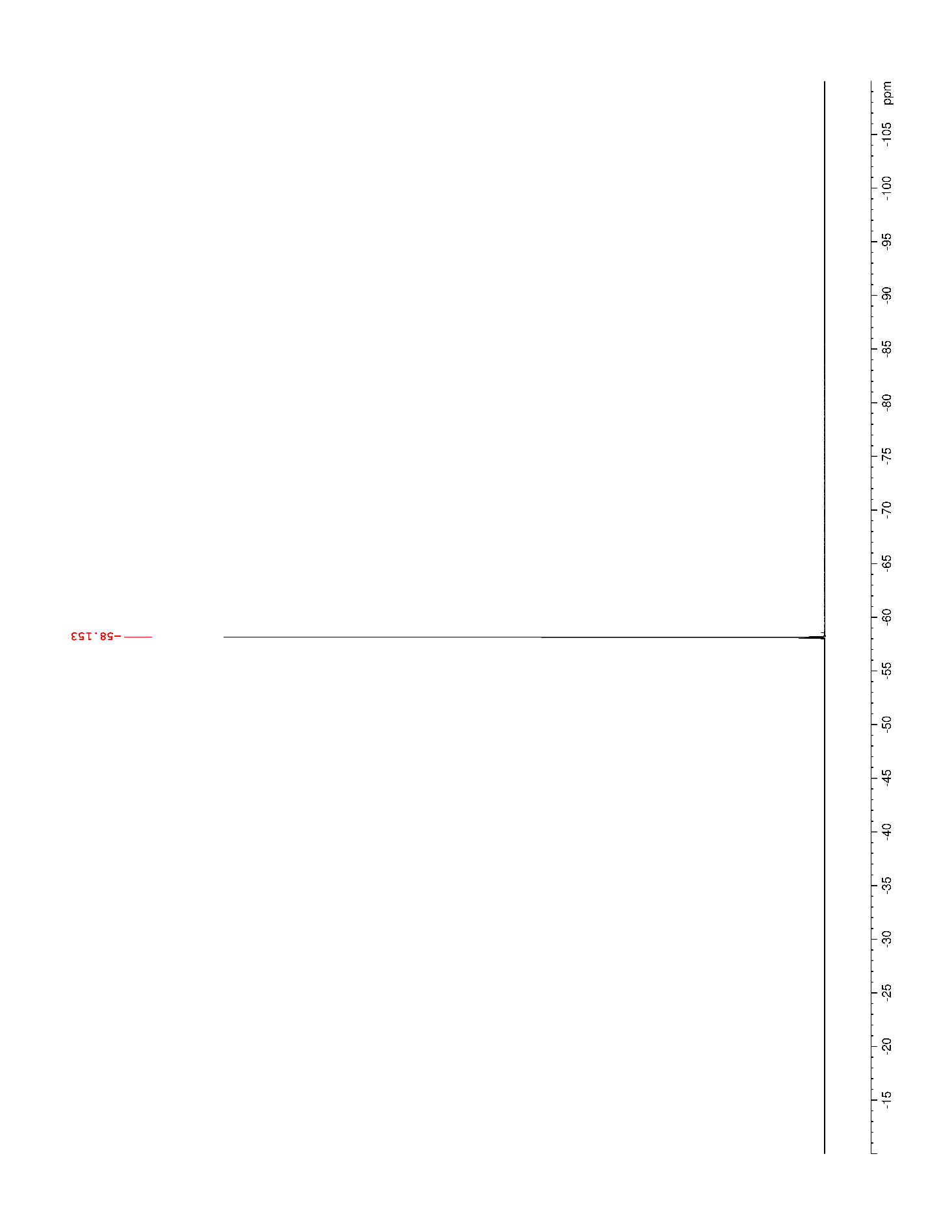

### Supplementary Figure 18. ^1^H NMR (500 MHz, DMSO-*d*_6_) of 4

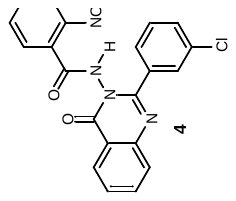

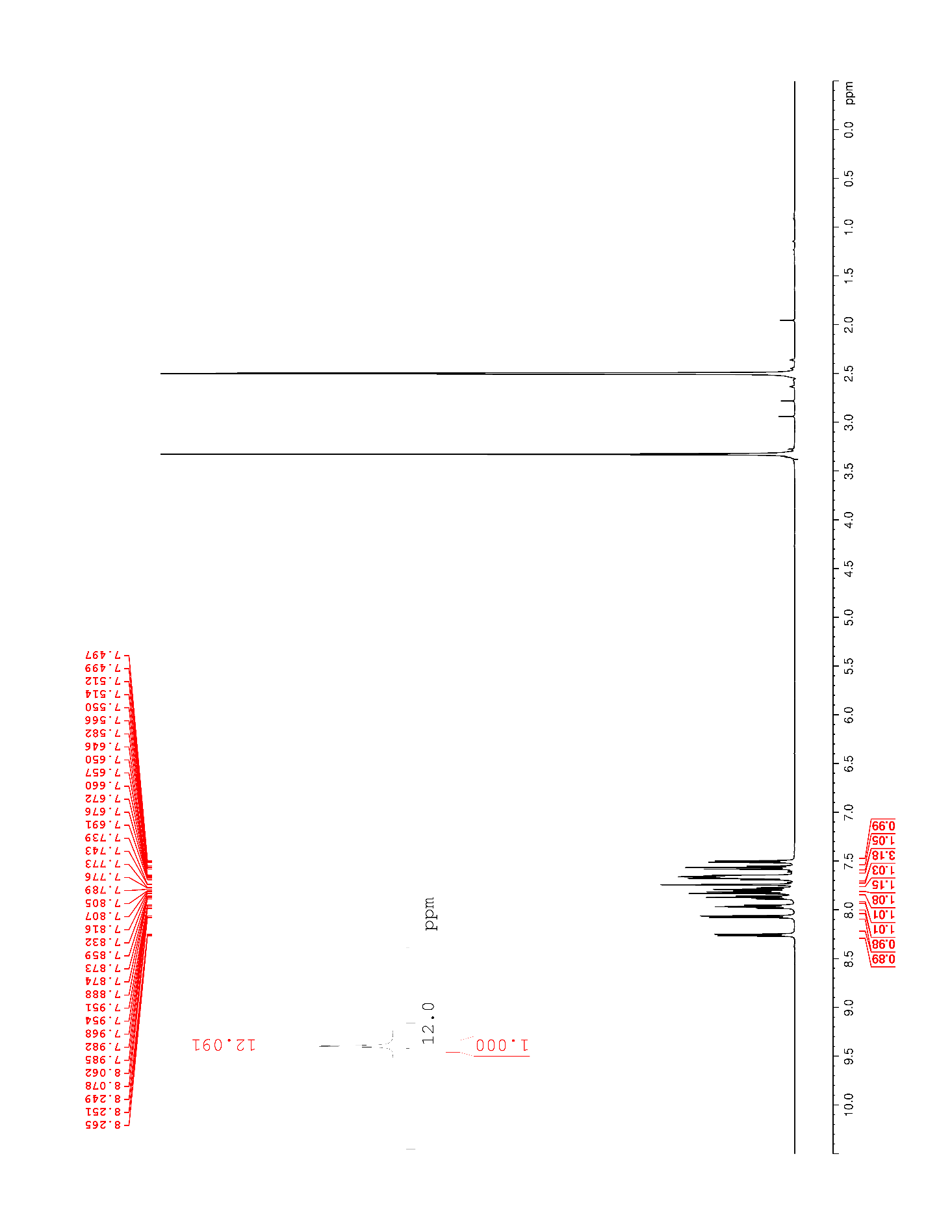

### Supplementary Figure 19. ^13^C NMR (125 MHz, DMSO-*d*_6_) of 4

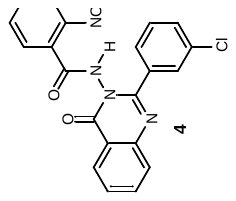

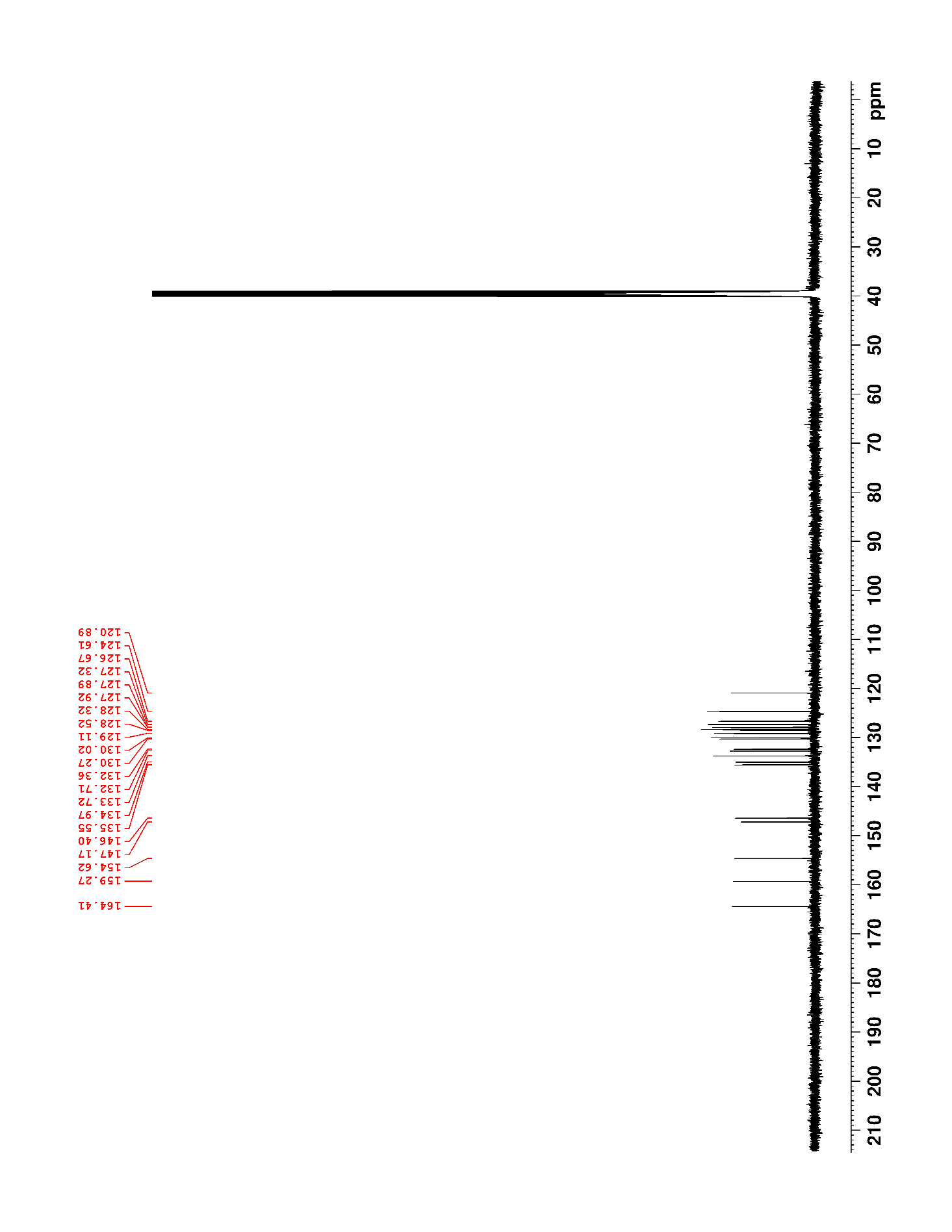

### Supplementary Figure 20. ^1^H NMR (500 MHz, DMSO-*d*_6_) of 5

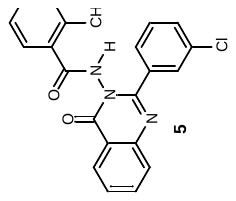

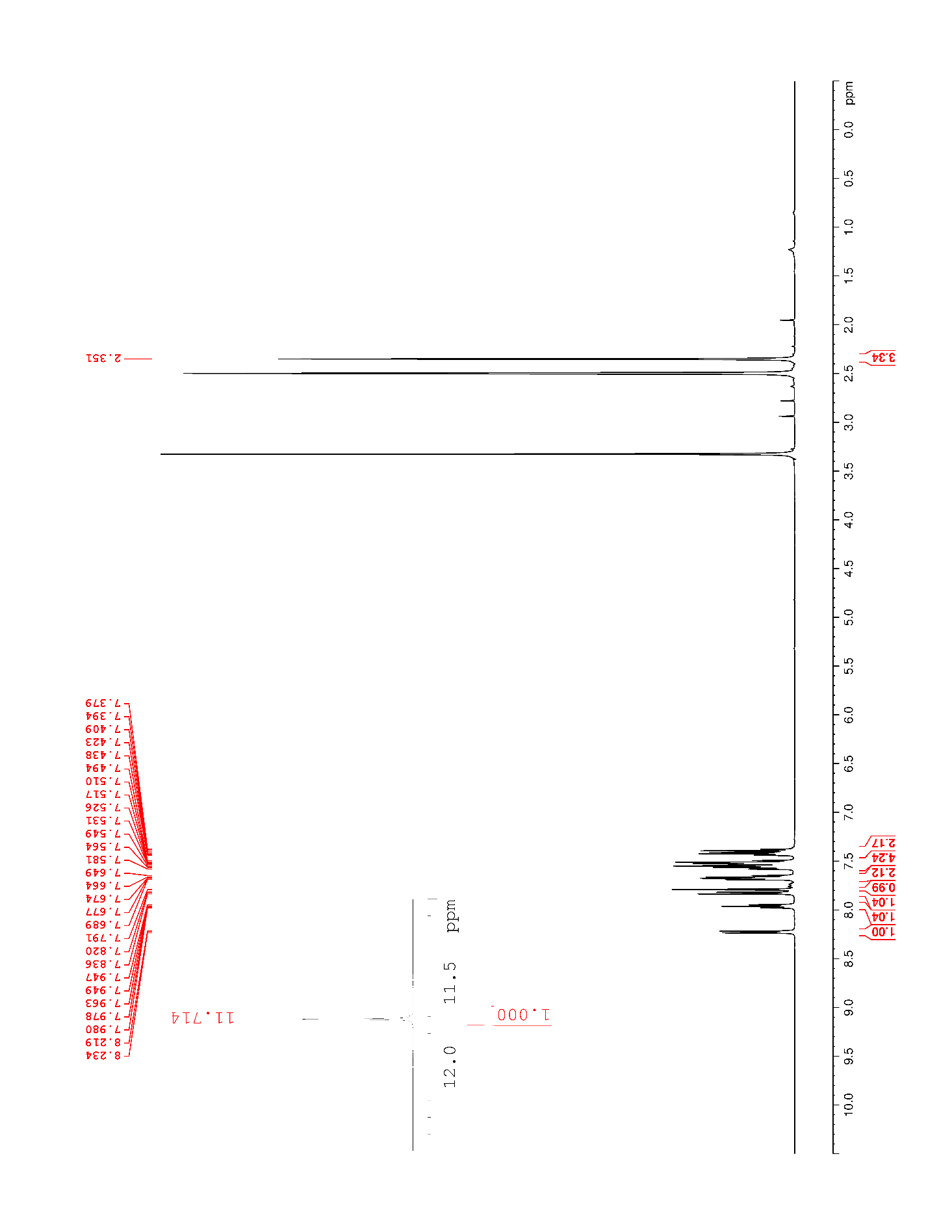

### Supplementary Figure 21. ^13^C NMR (125 MHz, DMSO-*d*_6_) of 5

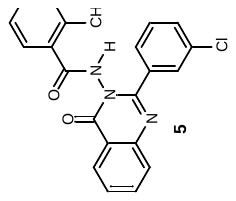

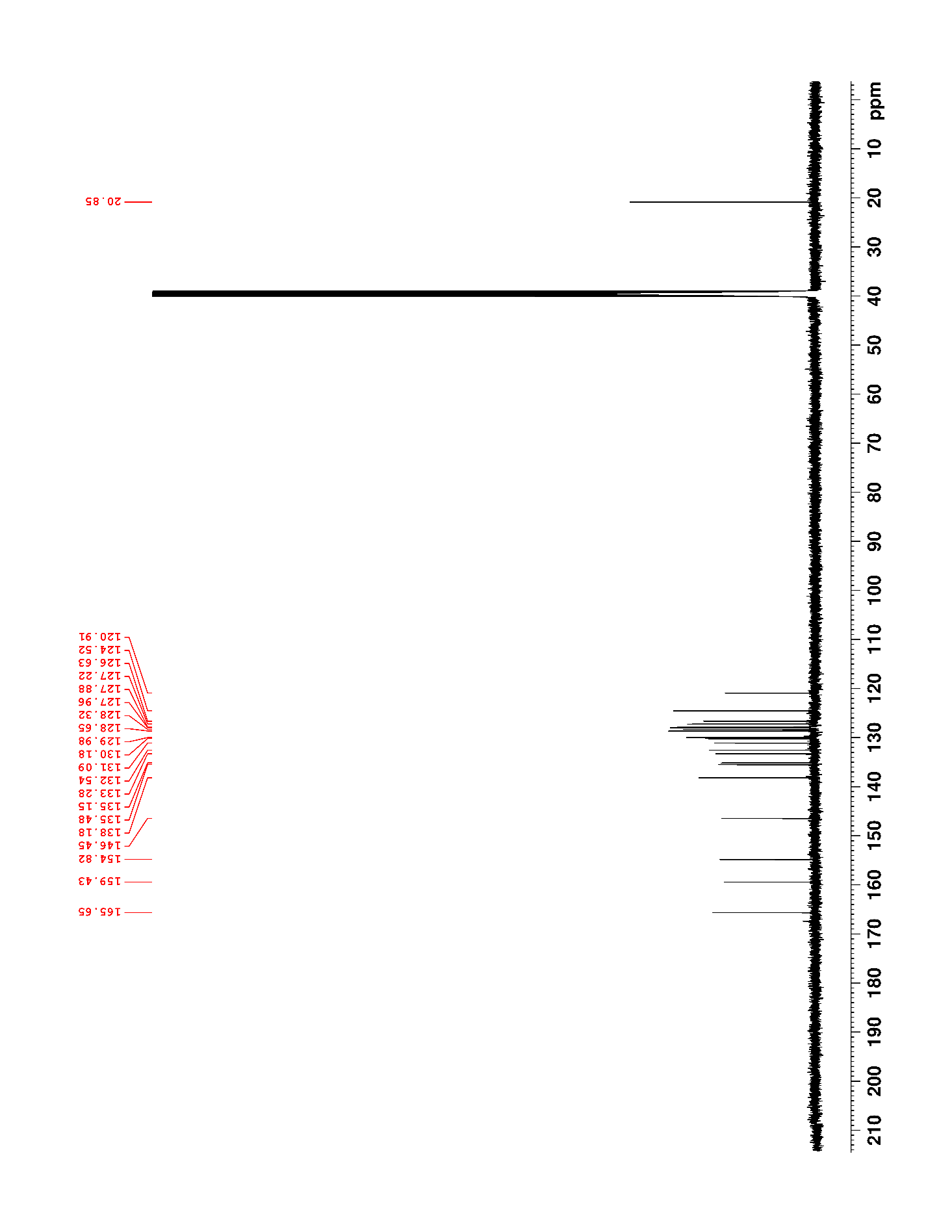

### Supplementary Figure 22. ^1^H NMR (500 MHz, CDCl_3_) of 6

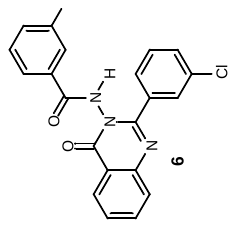

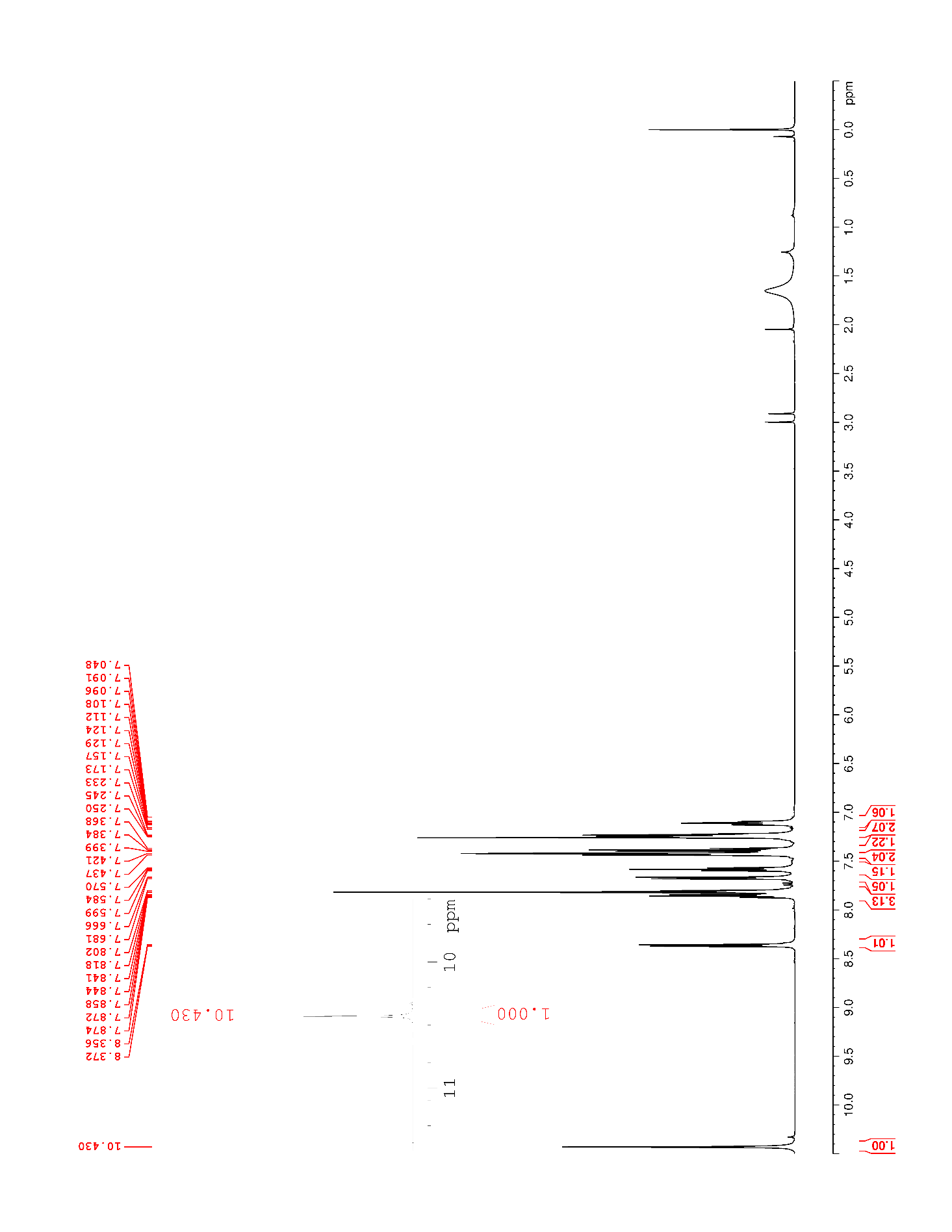

### Supplementary Figure 23. ^13^C NMR (125 MHz, CDCl_3_) of 6

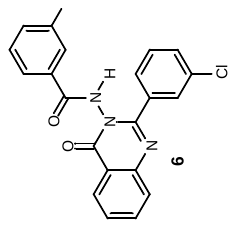

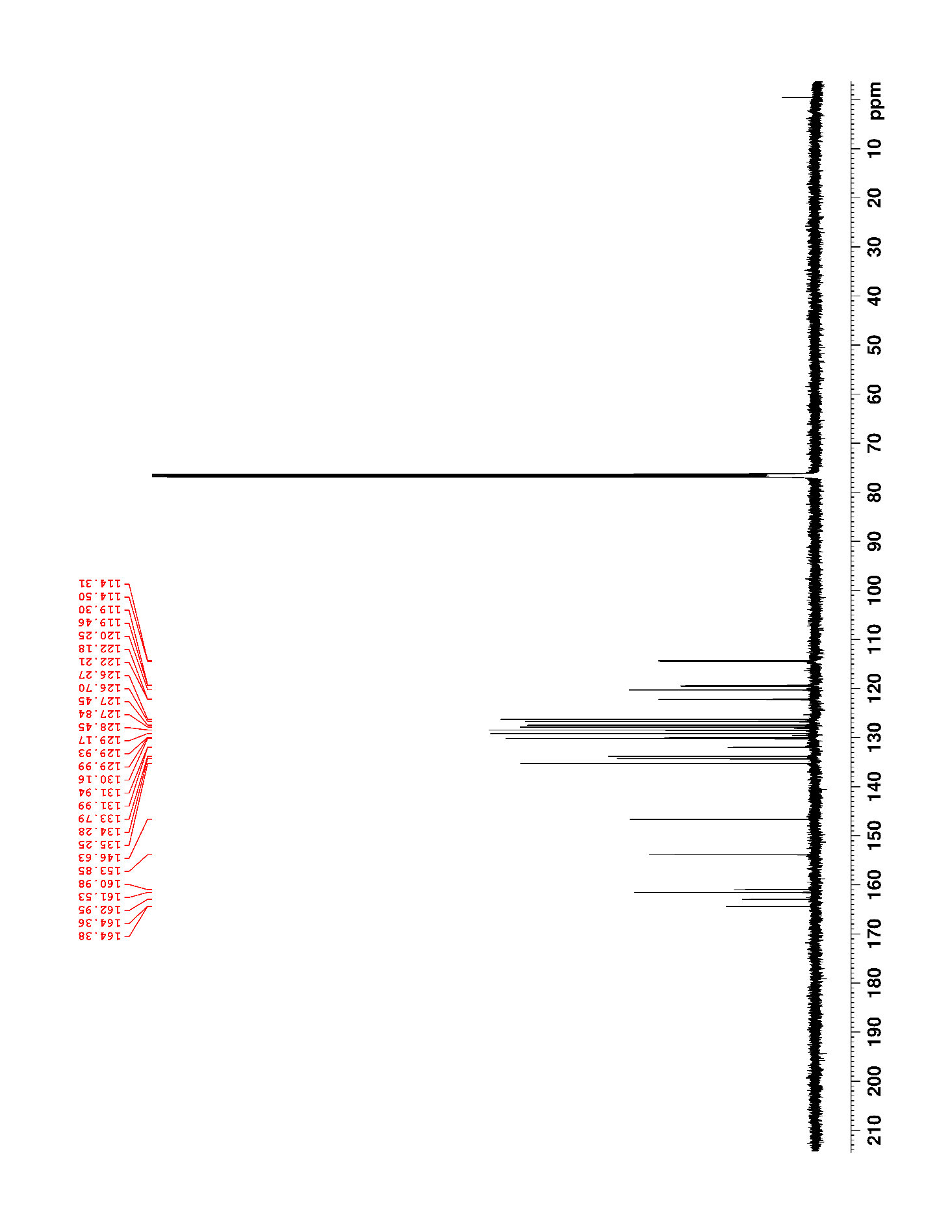

### Supplementary Figure 24. ^19^F NMR (470 MHz, CDCl_3_) of 6

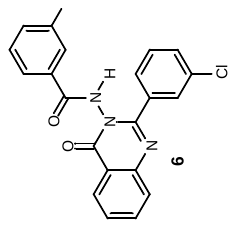

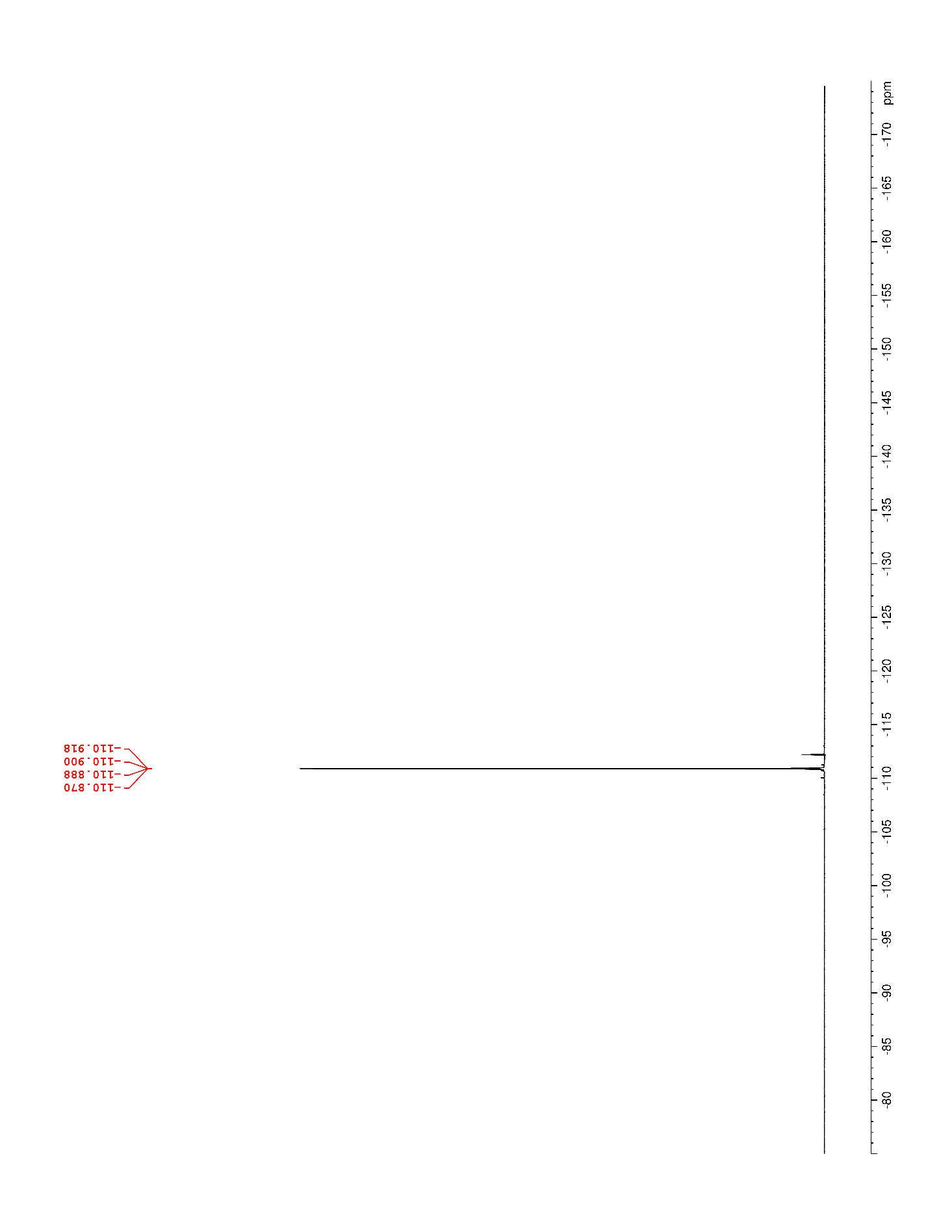

### Supplementary Figure 25. ^1^H NMR (500 MHz, DMSO-*d*_6_) of 7

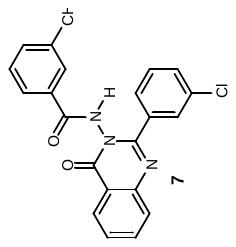

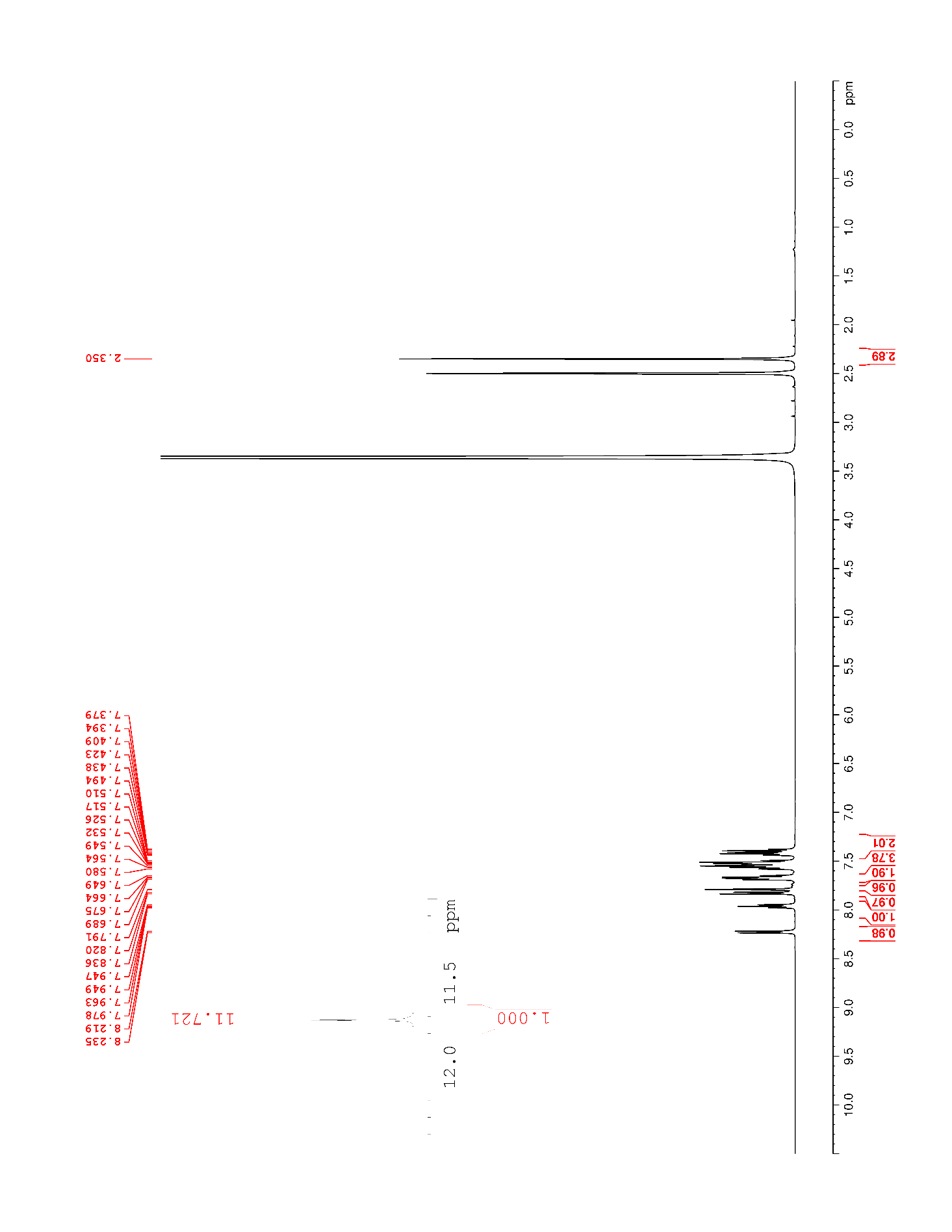

### Supplementary Figure 26. ^13^C NMR (125 MHz, DMSO-*d*_6_) of 7

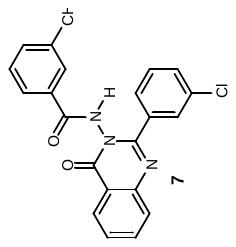

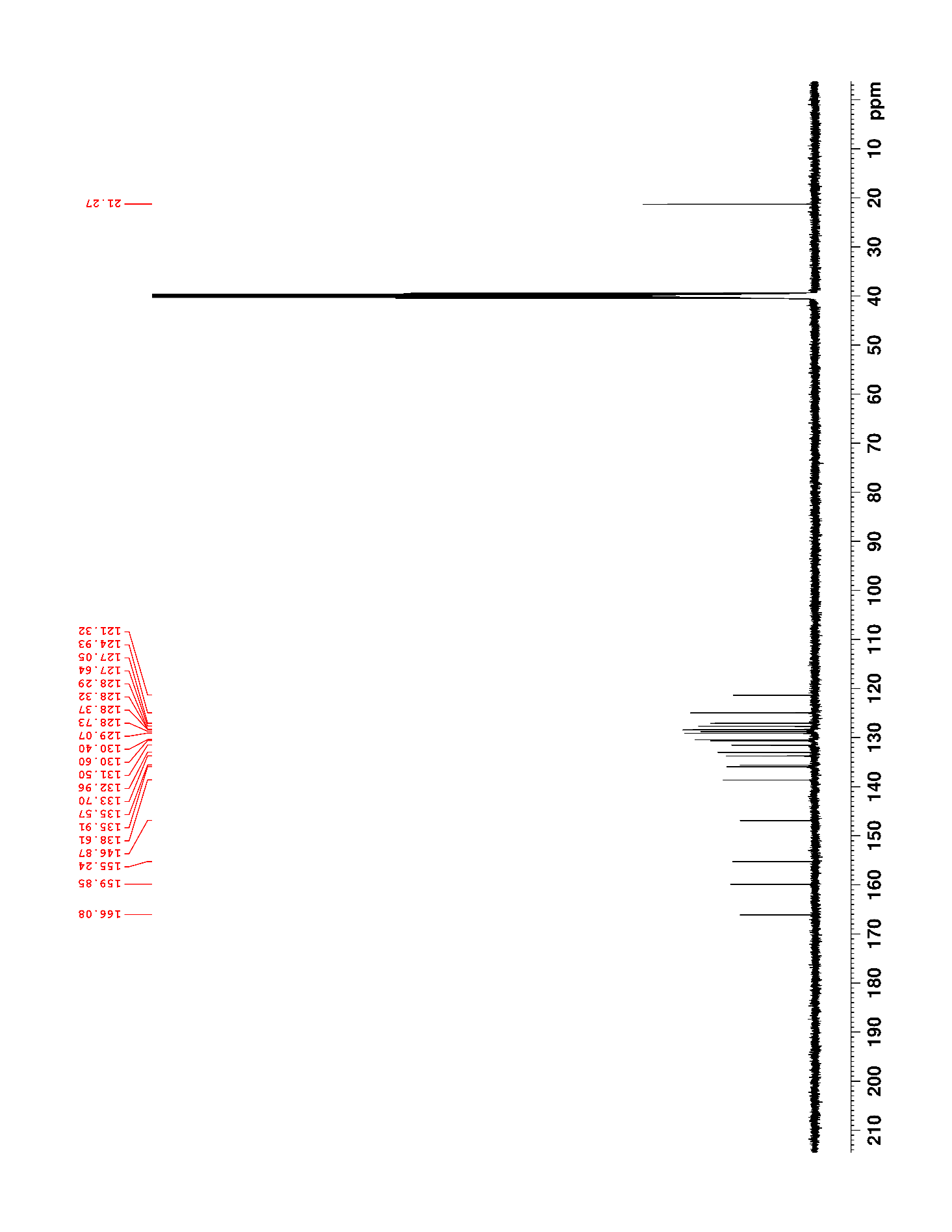

### Supplementary Figure 27. ^1^H NMR (500 MHz, DMSO-*d*_6_) of 8

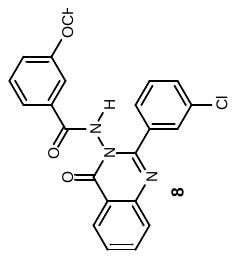

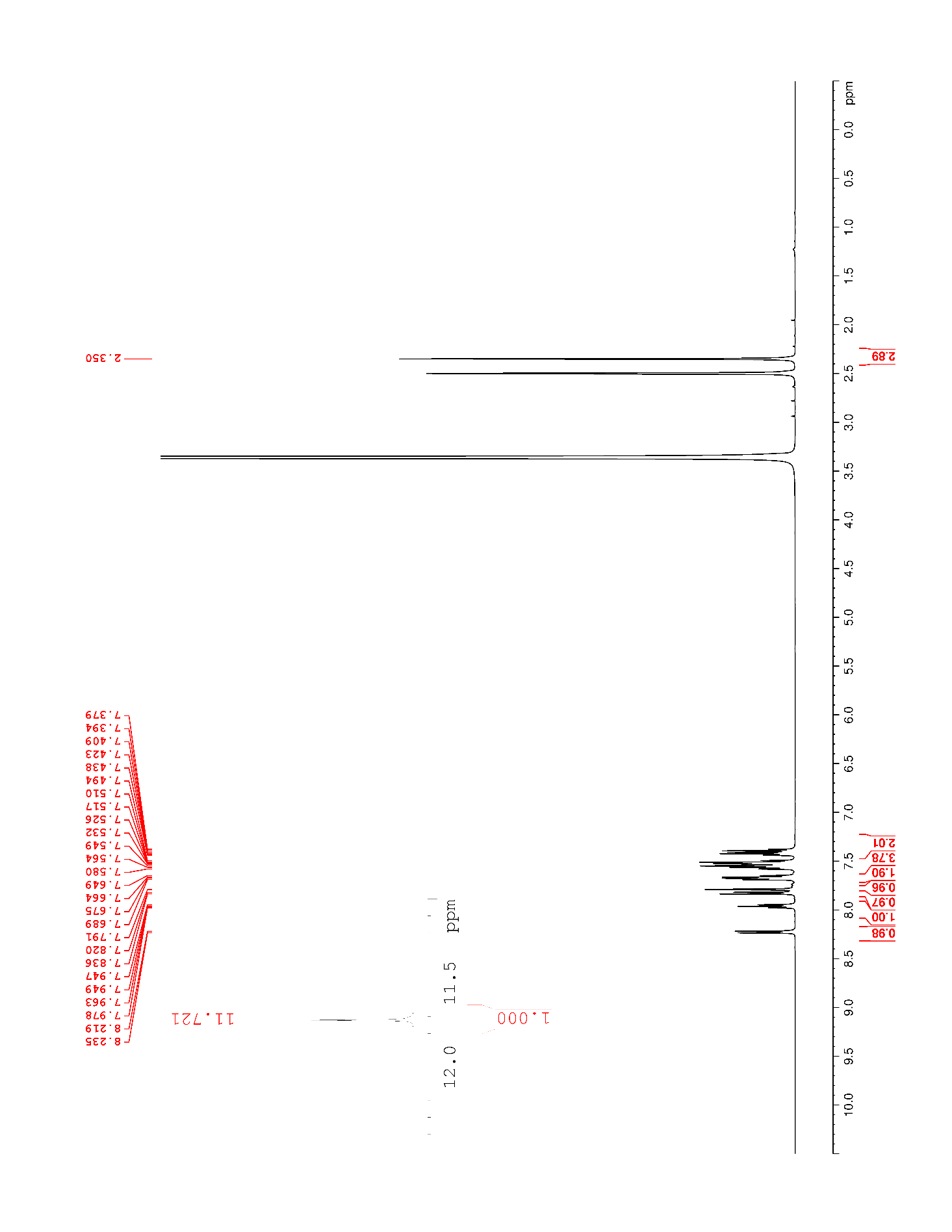

### Supplementary Figure 28. ^13^C NMR (125 MHz, DMSO-*d*_6_) of 8

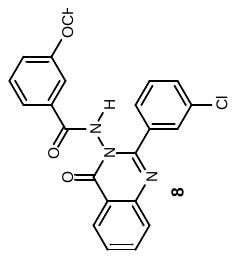

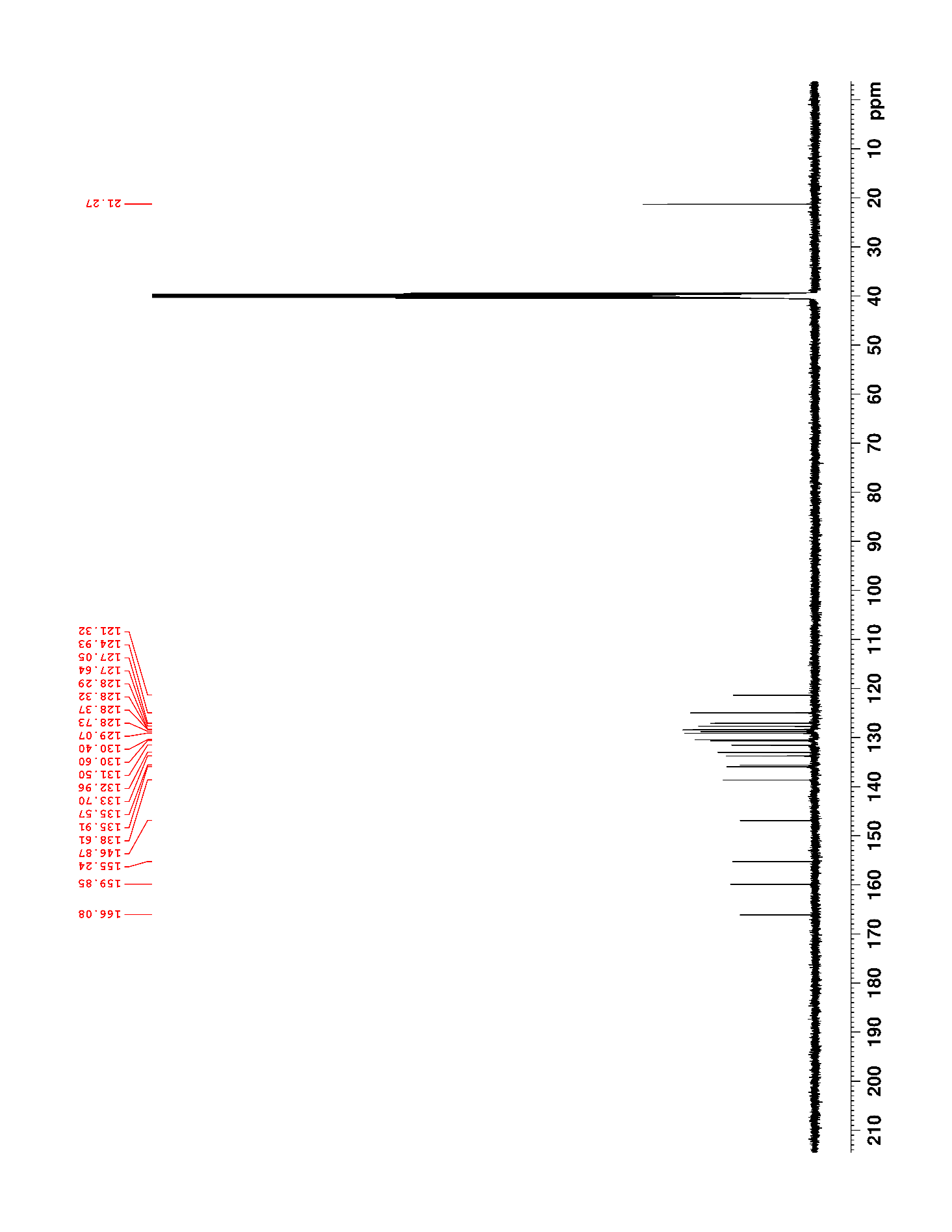

### Supplementary Figure 29. ^1^H NMR (500 MHz, DMSO-*d*_6_) of 9

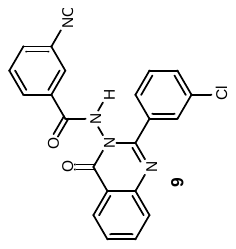

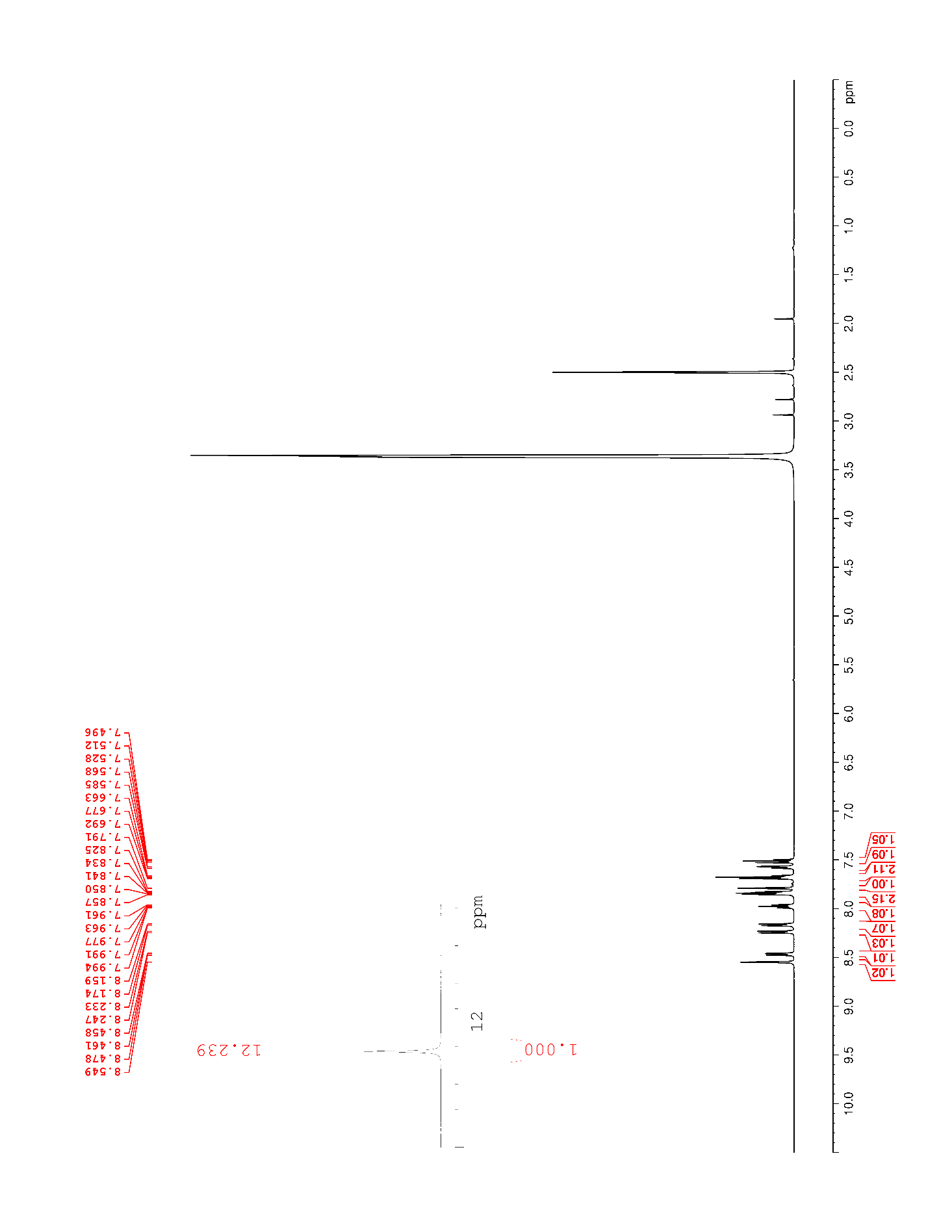

### Supplementary Figure 30. ^13^C NMR (125 MHz, DMSO-*d*_6_) of 9

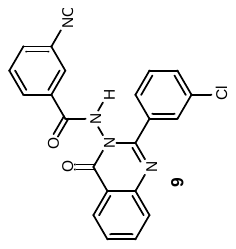

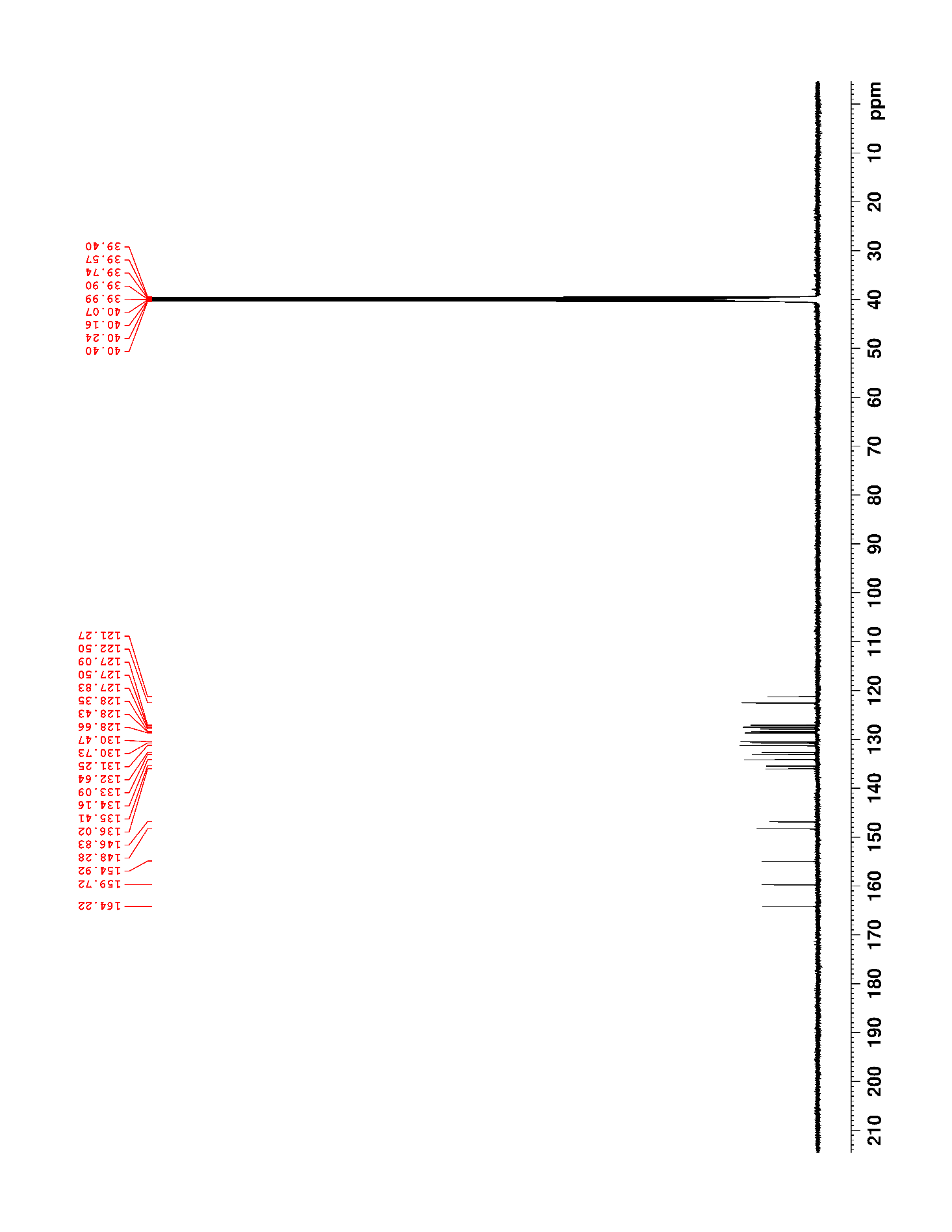

### Supplementary Figure 31. ^1^H NMR (500 MHz, DMSO-*d*_6_) of 10

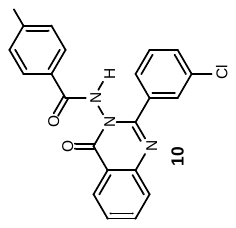

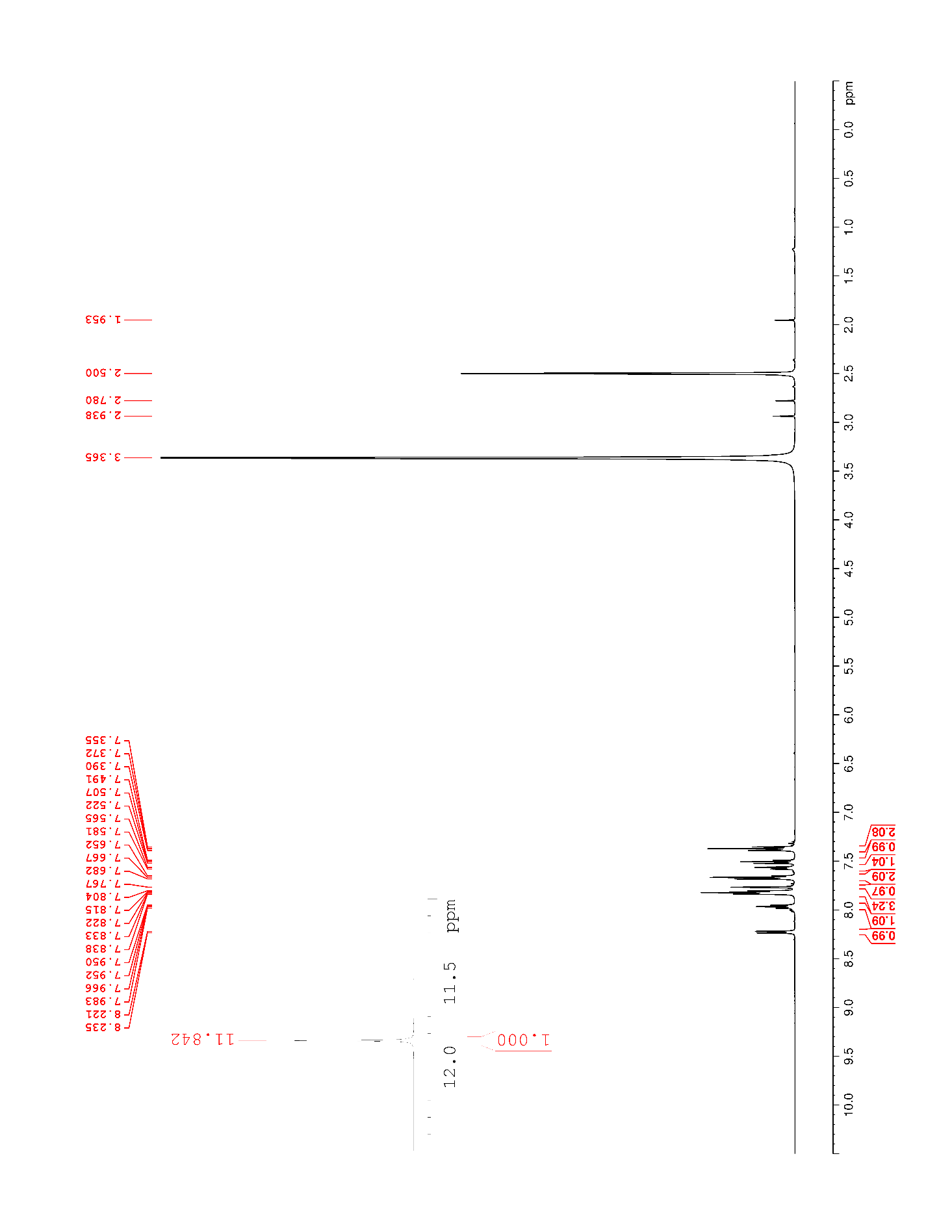

### Supplementary Figure 32. ^13^C NMR (125 MHz, DMSO-*d*_6_) of 10

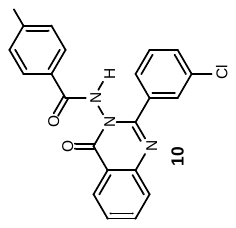

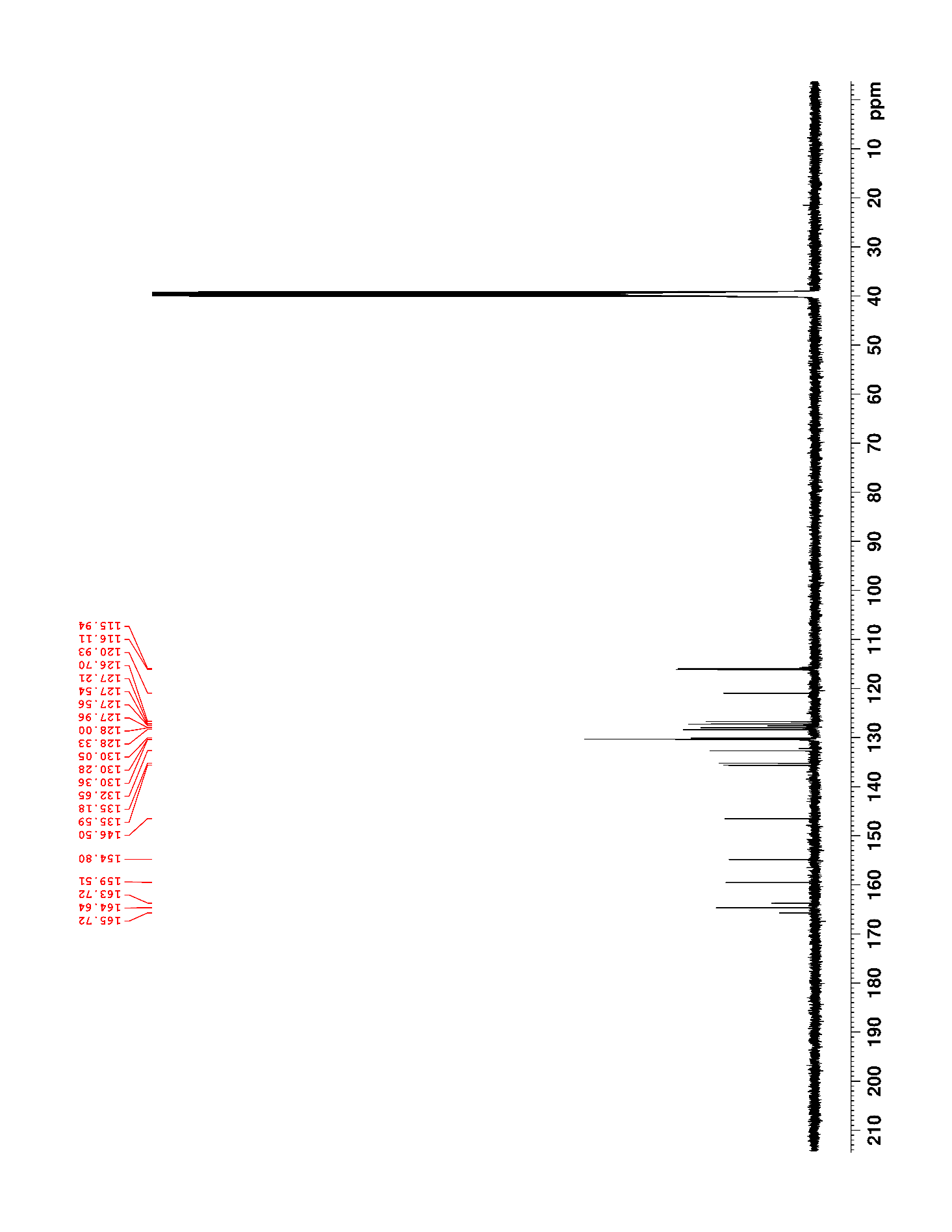

### Supplementary Figure 33. ^19^F NMR (470 MHz, DMSO-*d*_6_) of 10

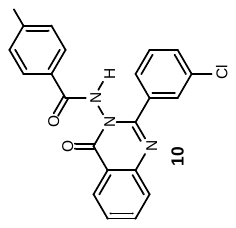

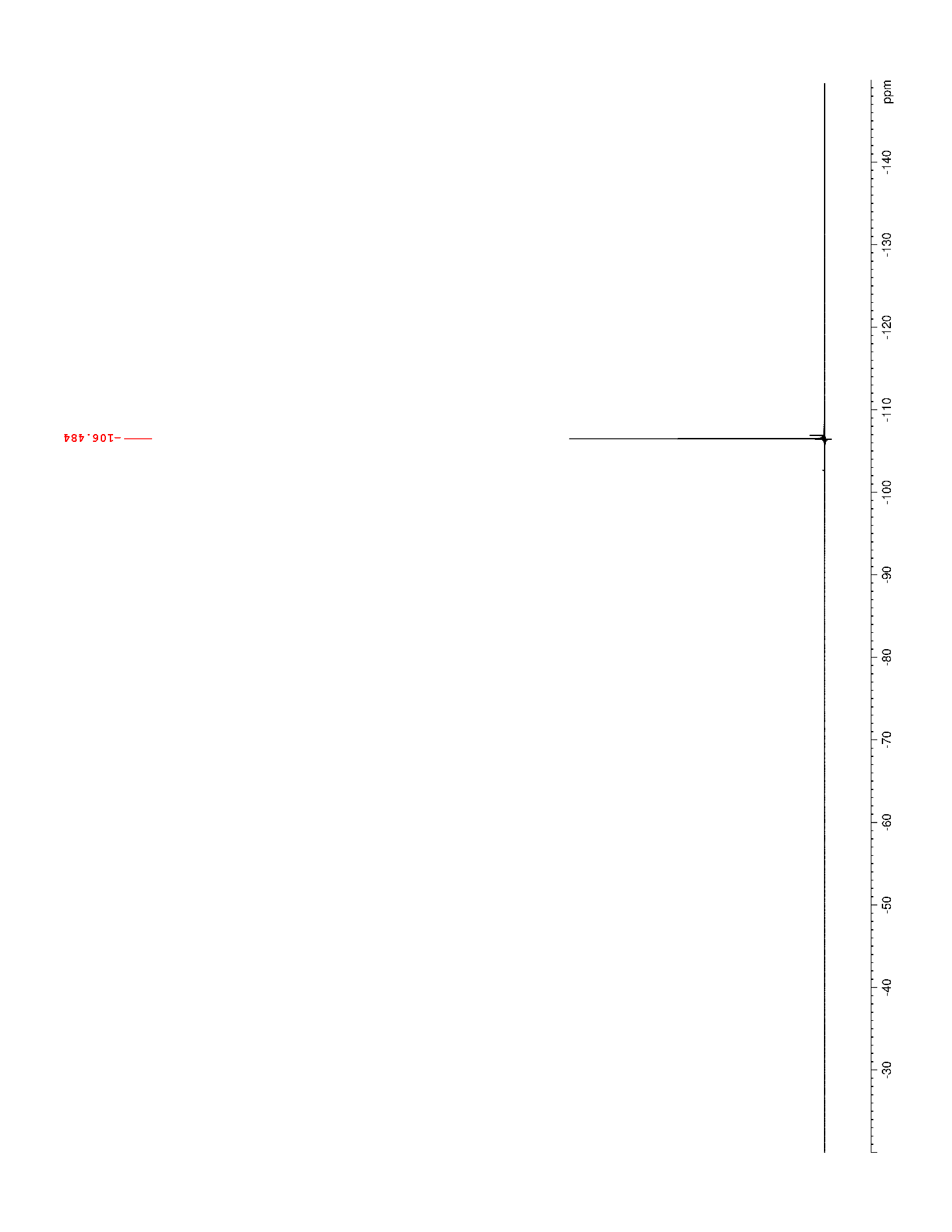

### Supplementary Figure 34. ^1^H NMR (500 MHz, DMSO-*d*_6_) of 11

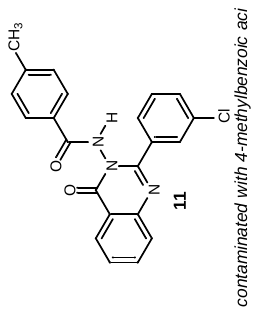

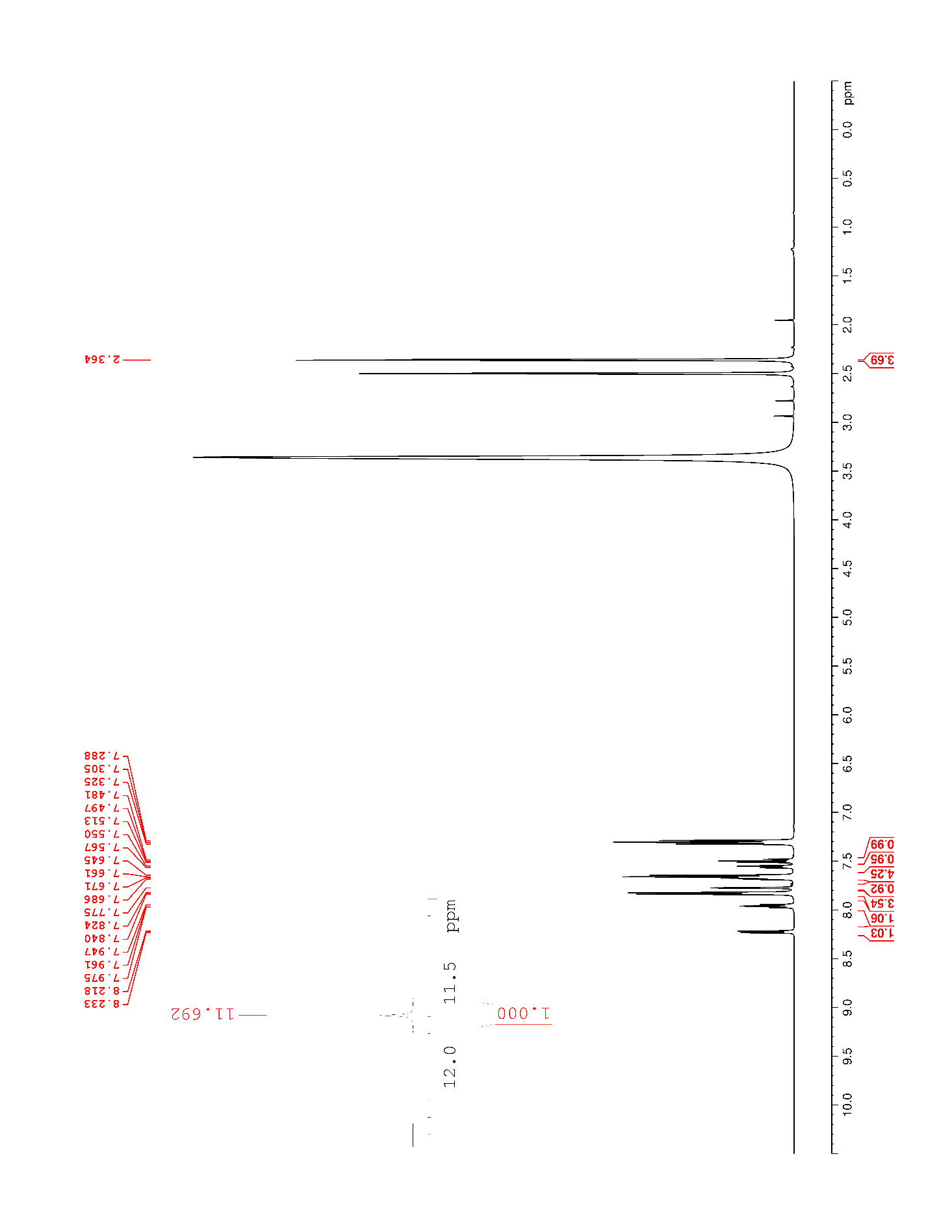

### Supplementary Figure 35. ^1^H NMR (500 MHz, DMSO-*d*_6_) of 12

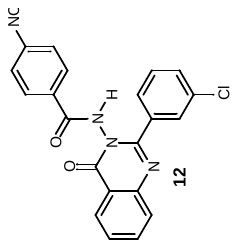

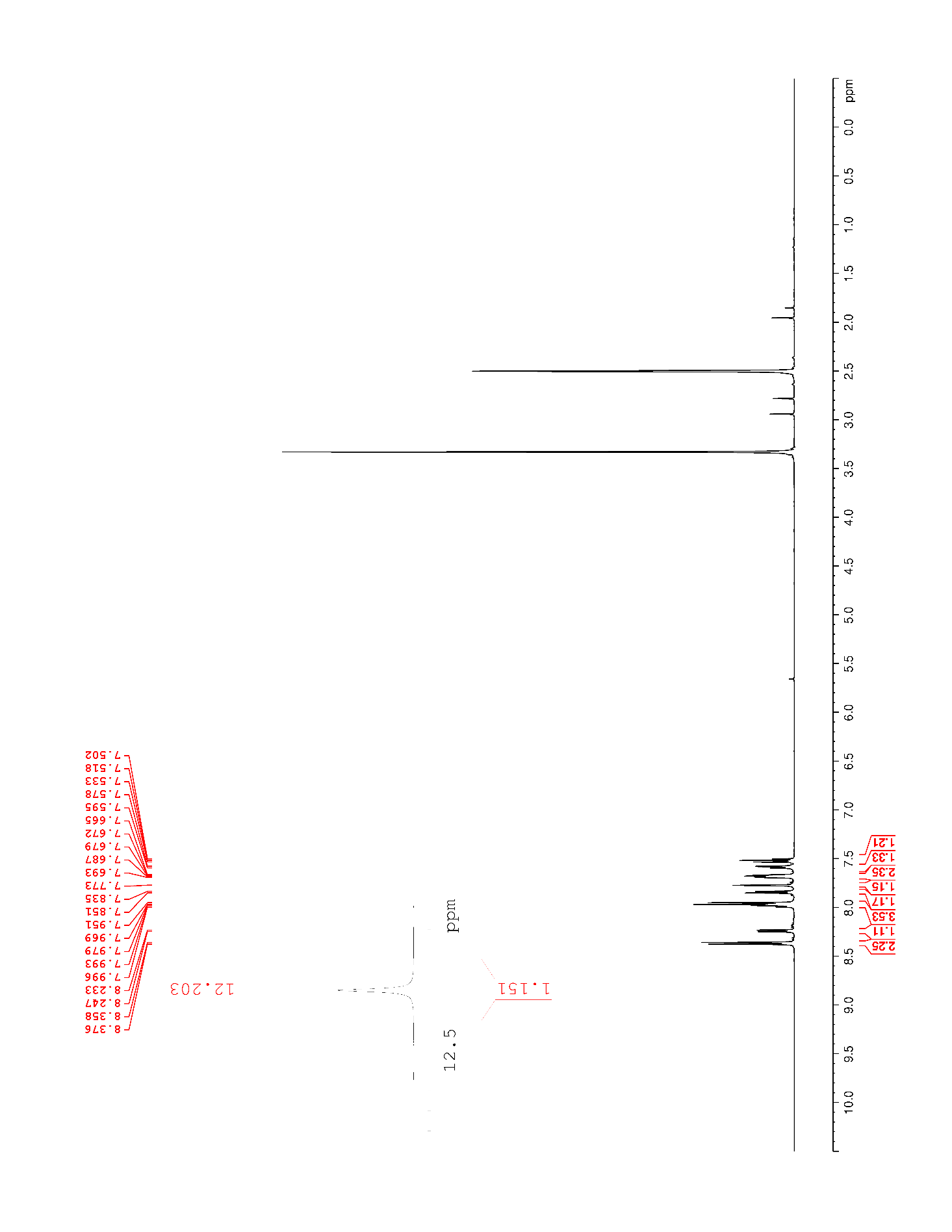

### Supplementary Figure 36. ^13^C NMR (125 MHz, DMSO-*d*_6_) of 12

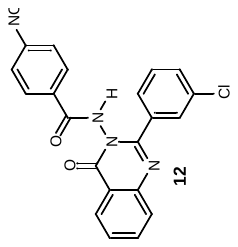

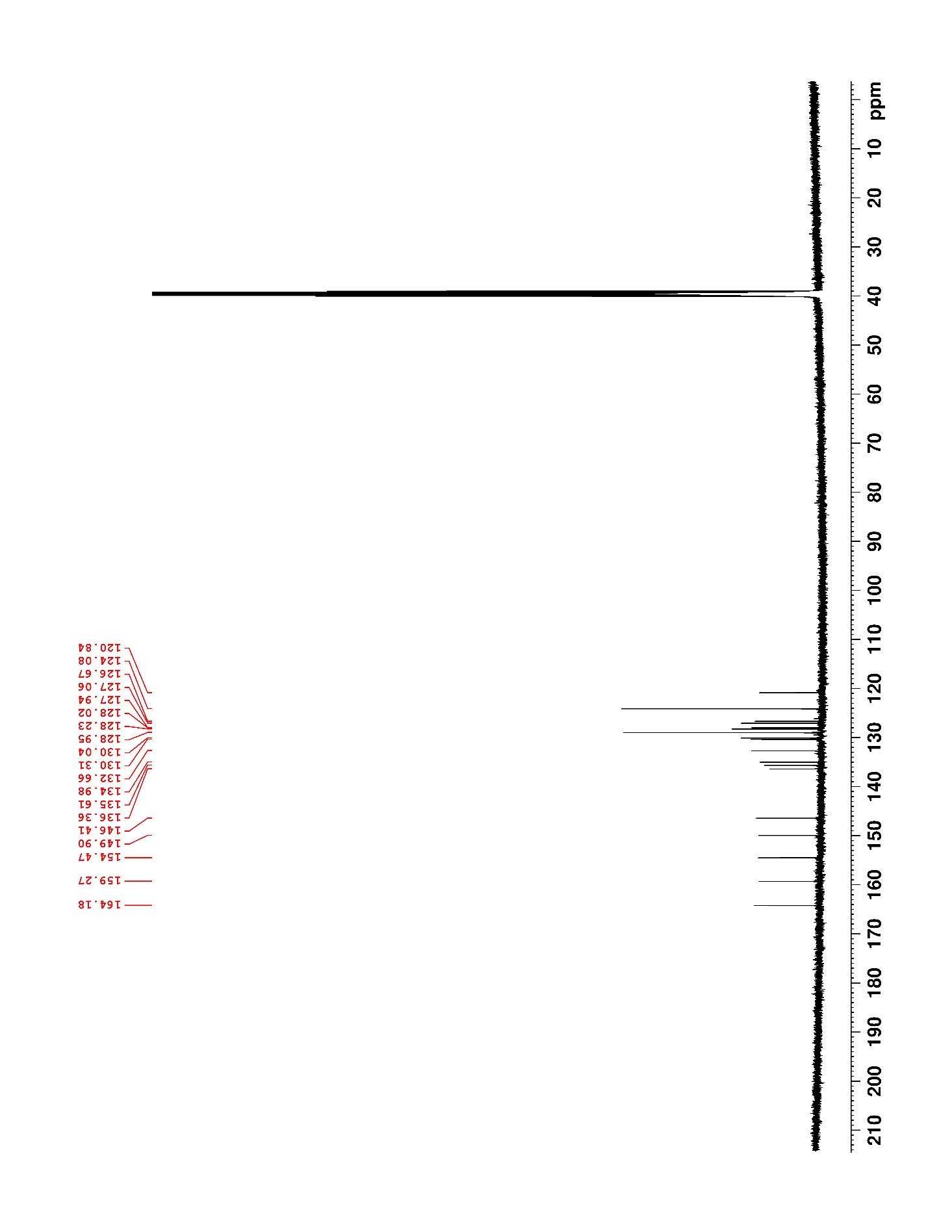

### Supplementary Figure 37. ^1^H NMR (500 MHz, DMSO-*d*_6_) of 13

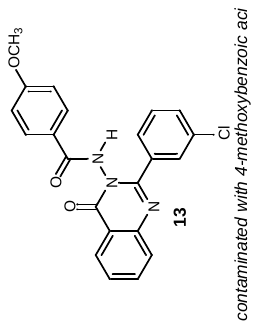

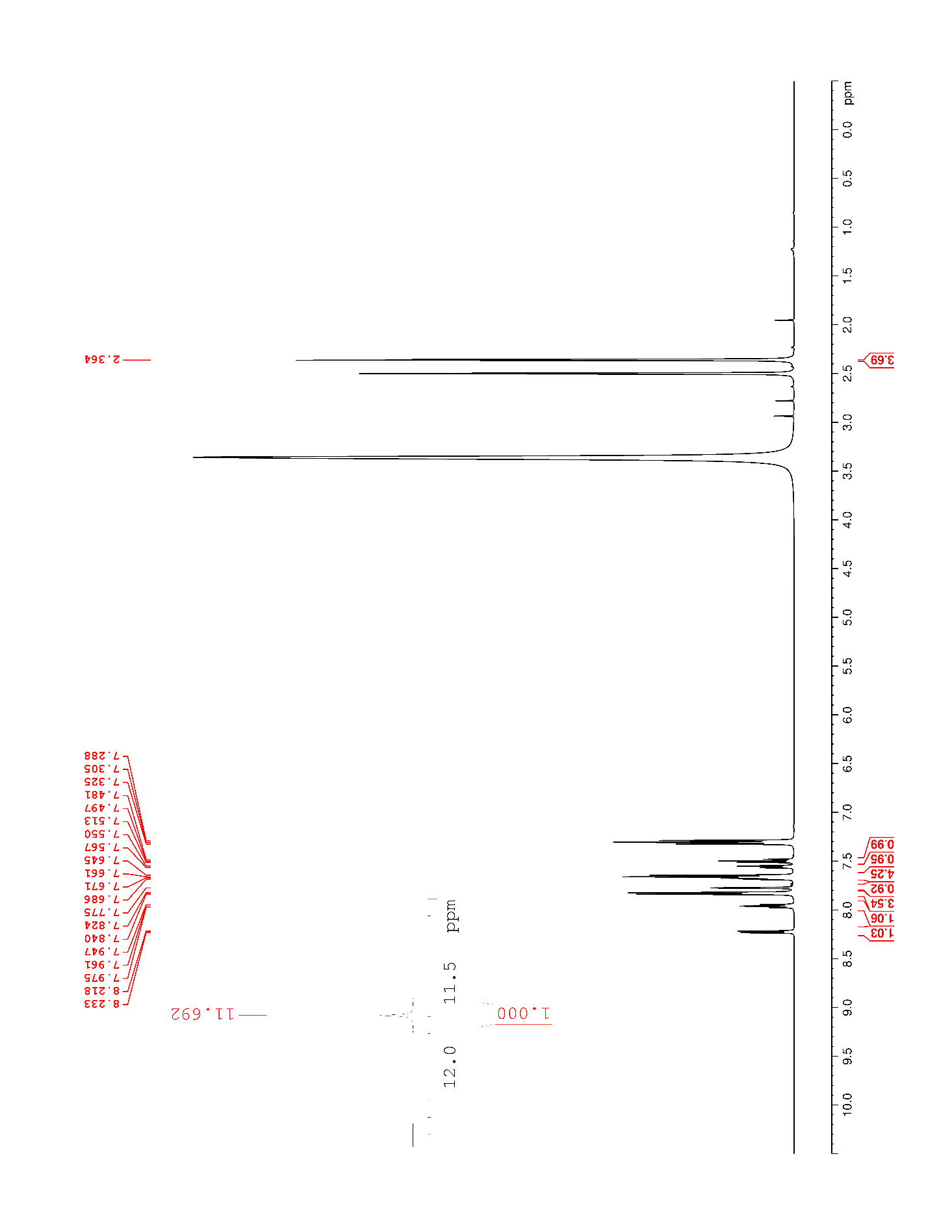

### Supplementary Figure 38. ^1^H NMR (500 MHz, DMSO-*d*_6_) of 14

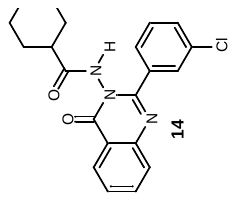

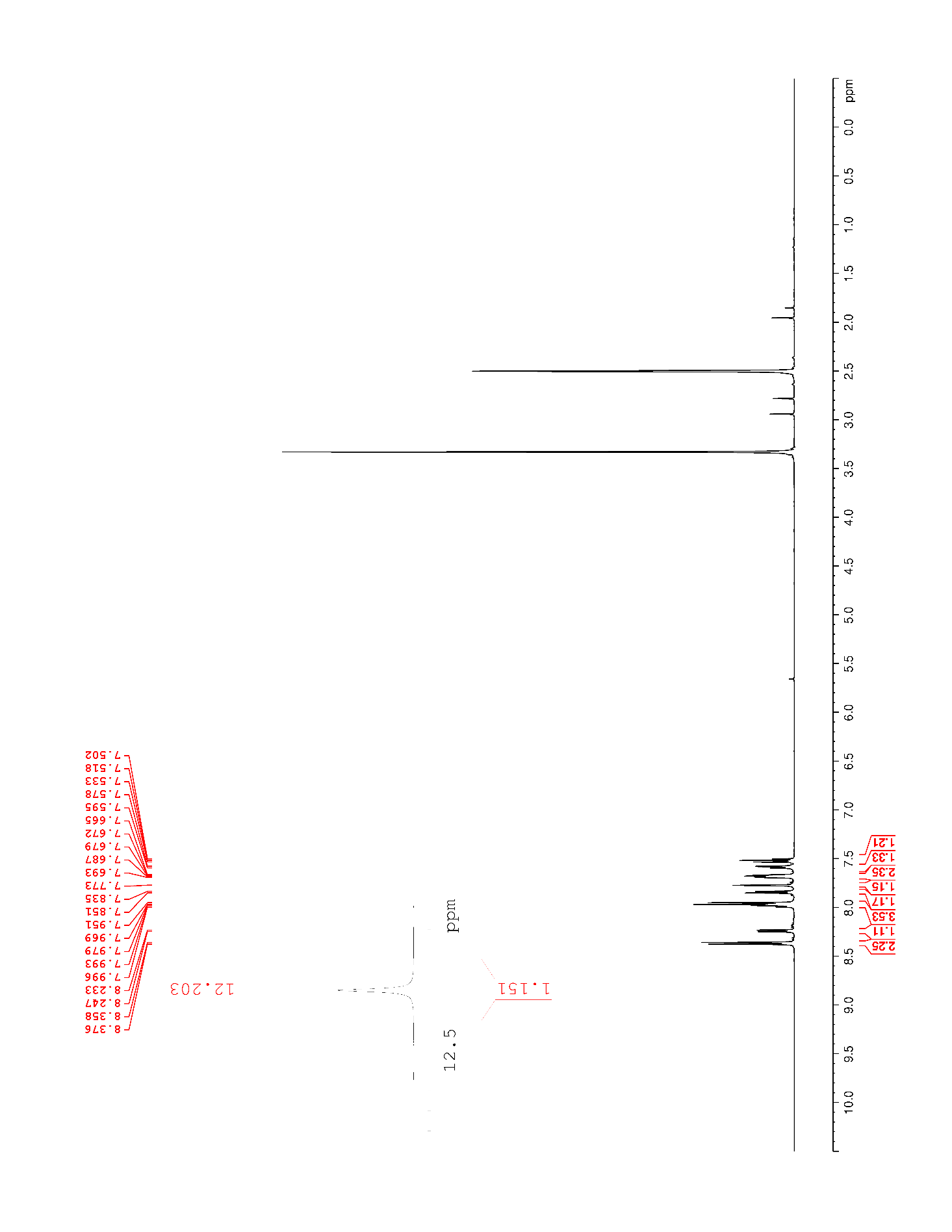

### Supplementary Figure 39. ^13^C NMR (125 MHz, DMSO-*d*_6_) of 14

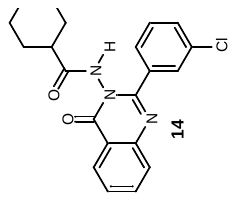

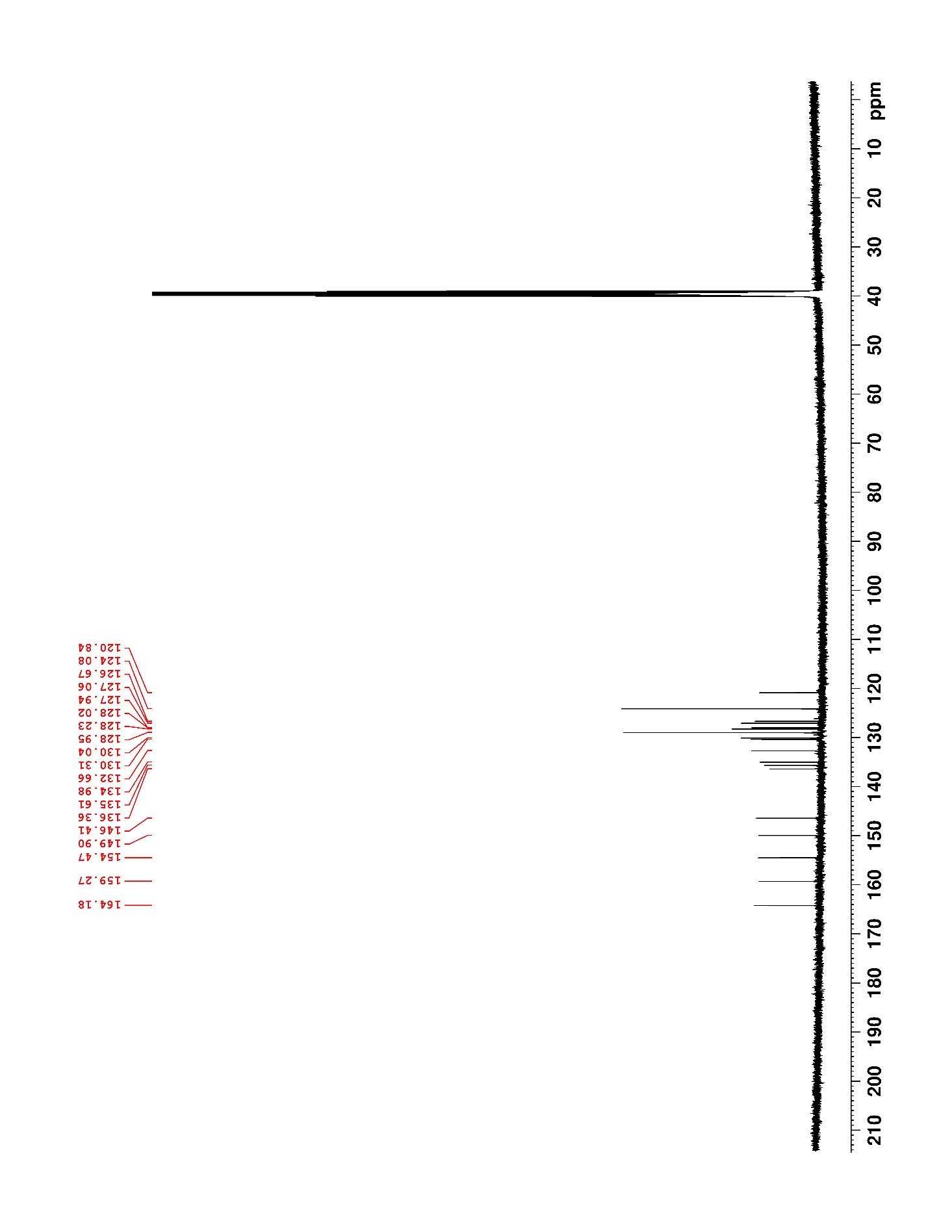

### Supplementary Figure 40. ^1^H NMR (500 MHz, DMSO-*d*_6_) of 15

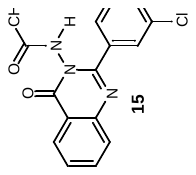

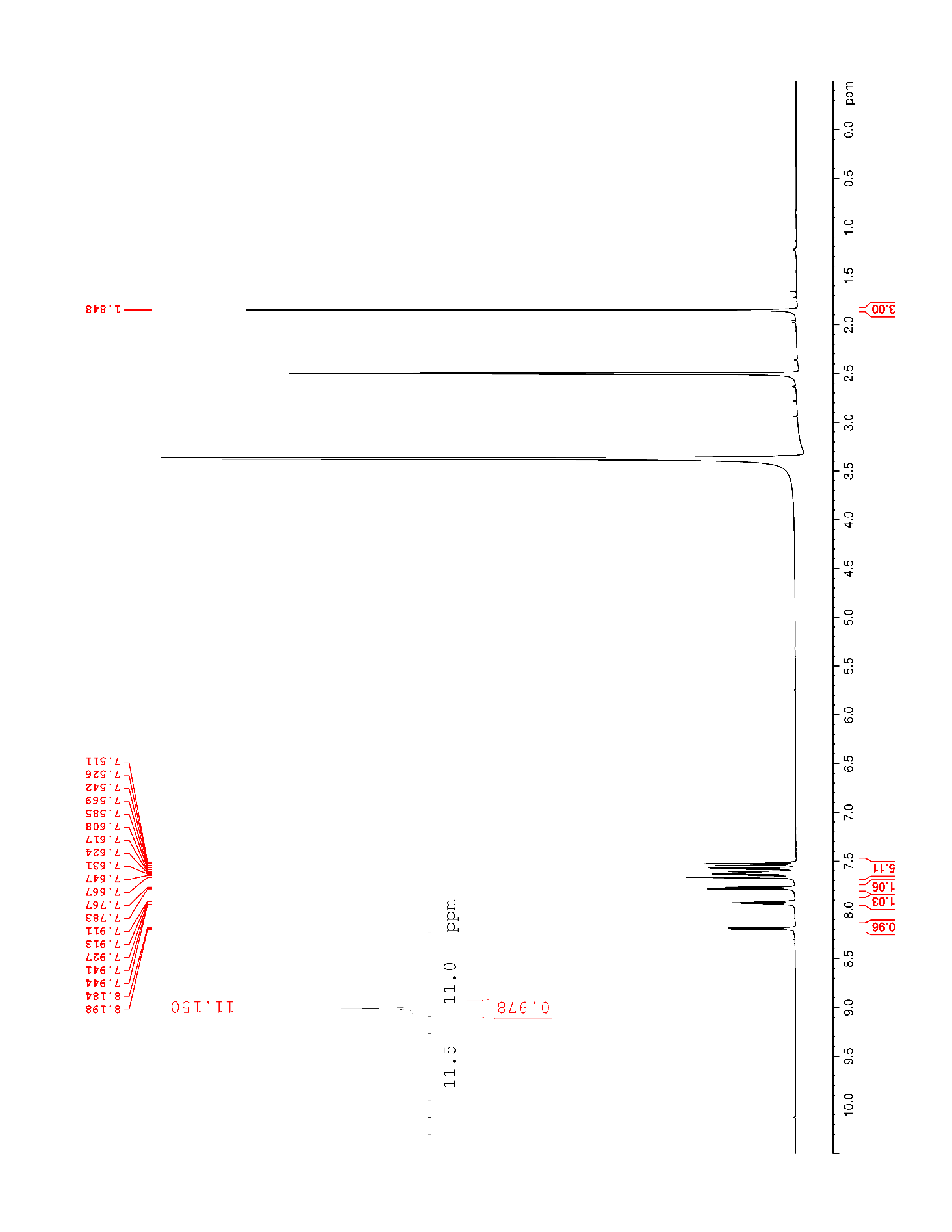

### Supplementary Figure 41. ^13^C NMR (125 MHz, DMSO-*d*_6_) of 15

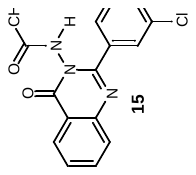

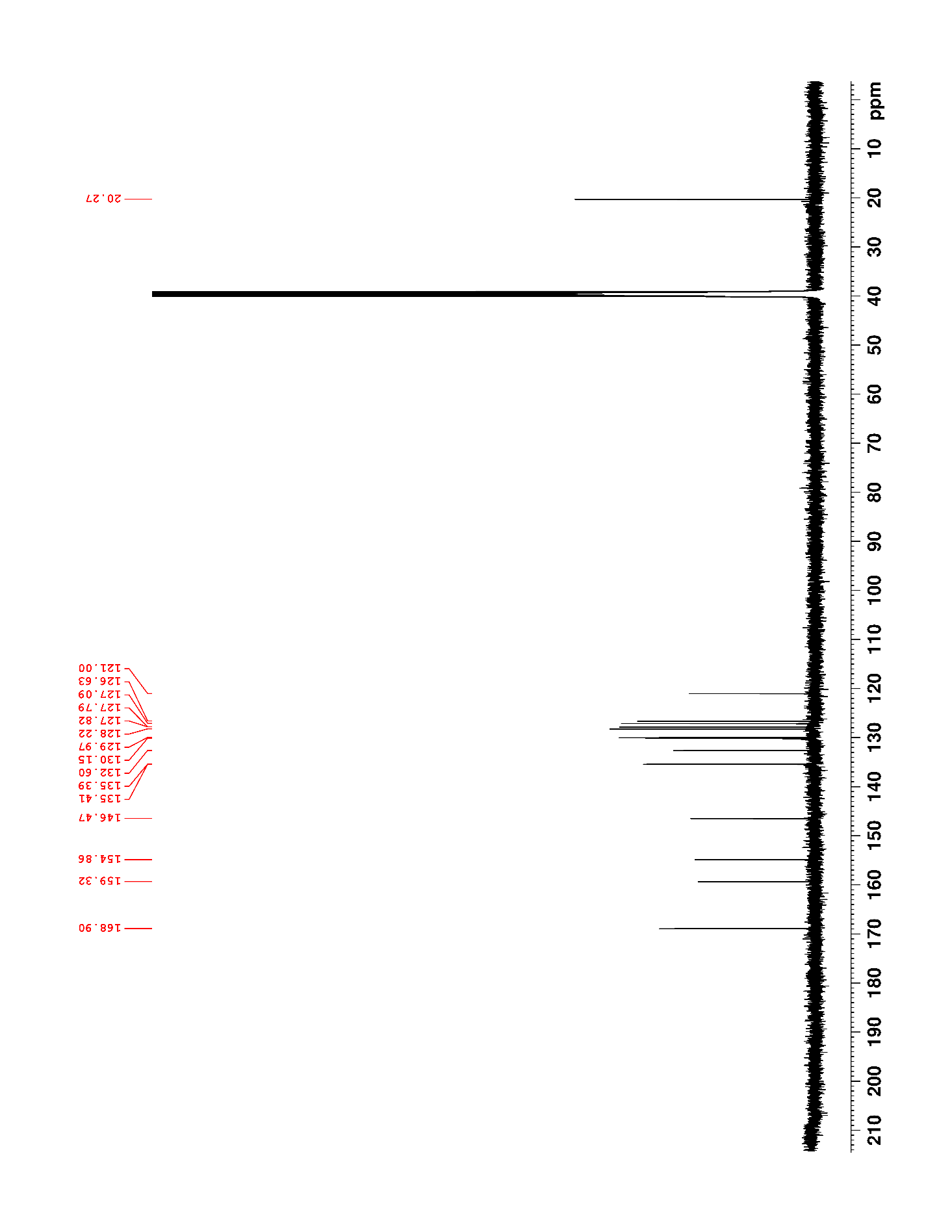

### Supplementary Figure 42. ^1^H NMR (500 MHz, DMSO-*d*_6_) of 16

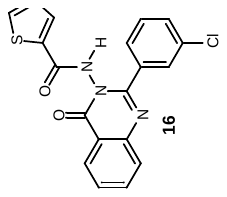

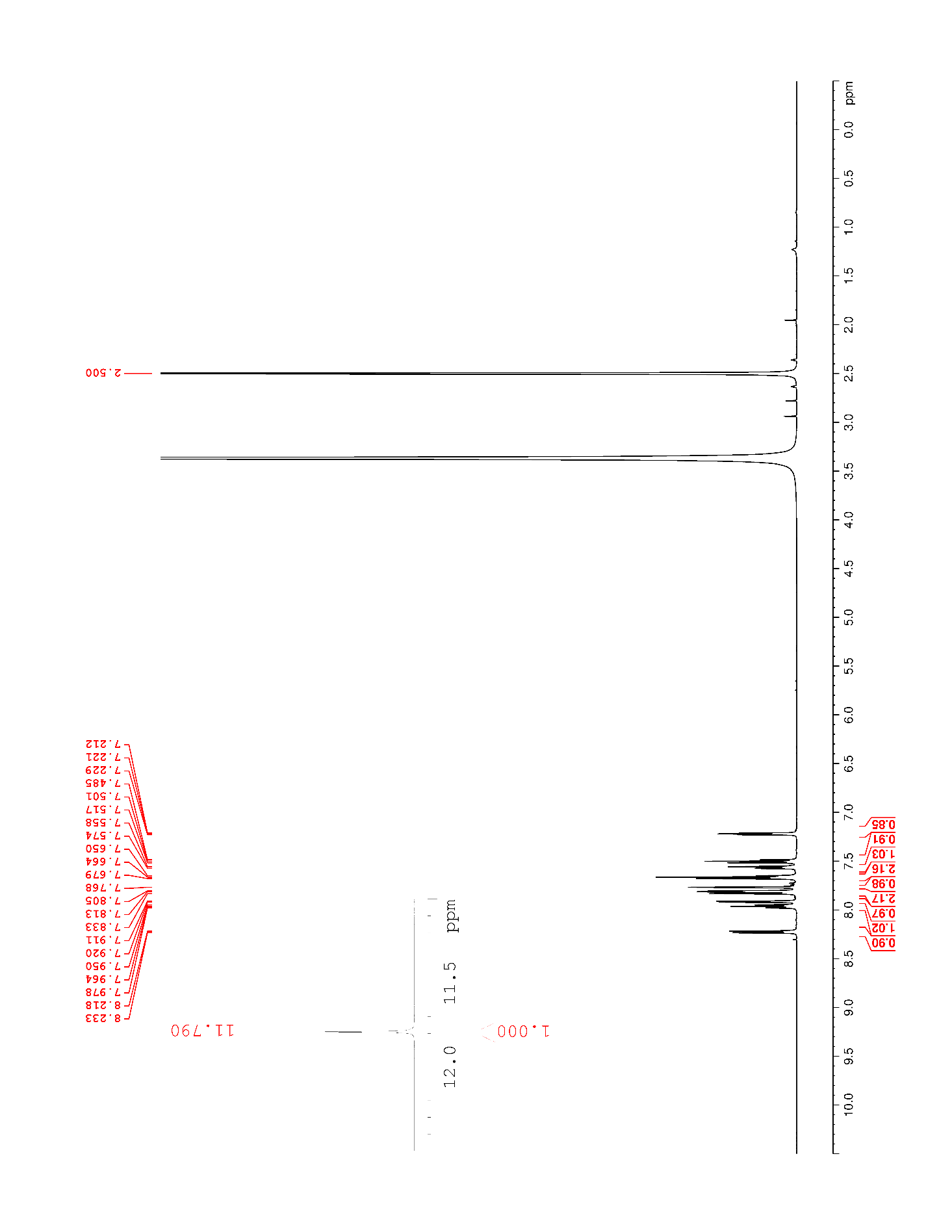

### Supplementary Figure 43. ^13^C NMR (125 MHz, DMSO-*d*_6_) of 16

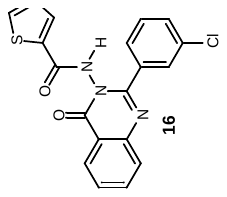

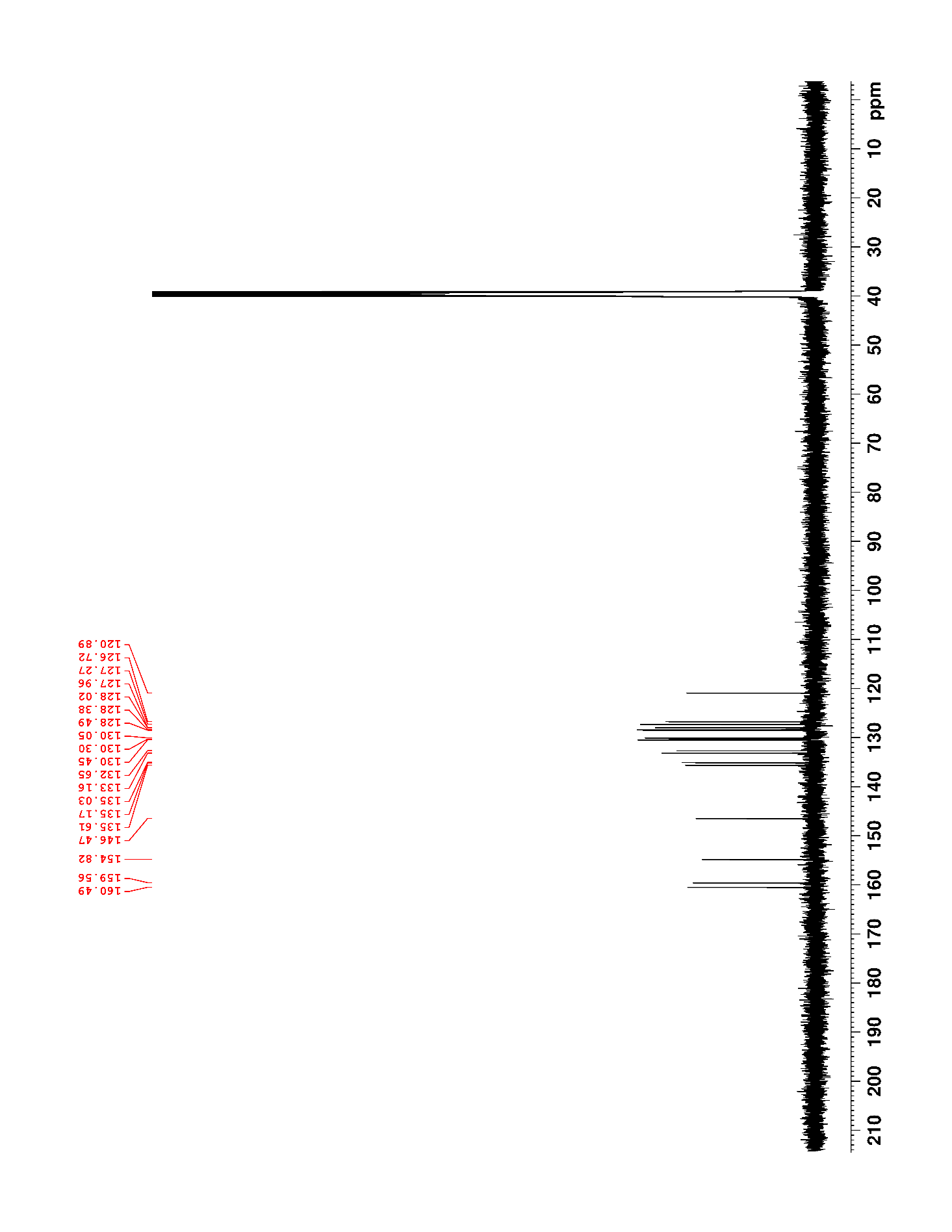

### Supplementary Figure 44. ^1^H NMR (500 MHz, DMSO-*d*_6_) of 17

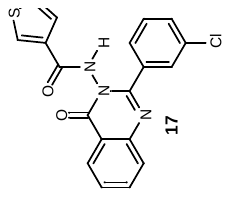

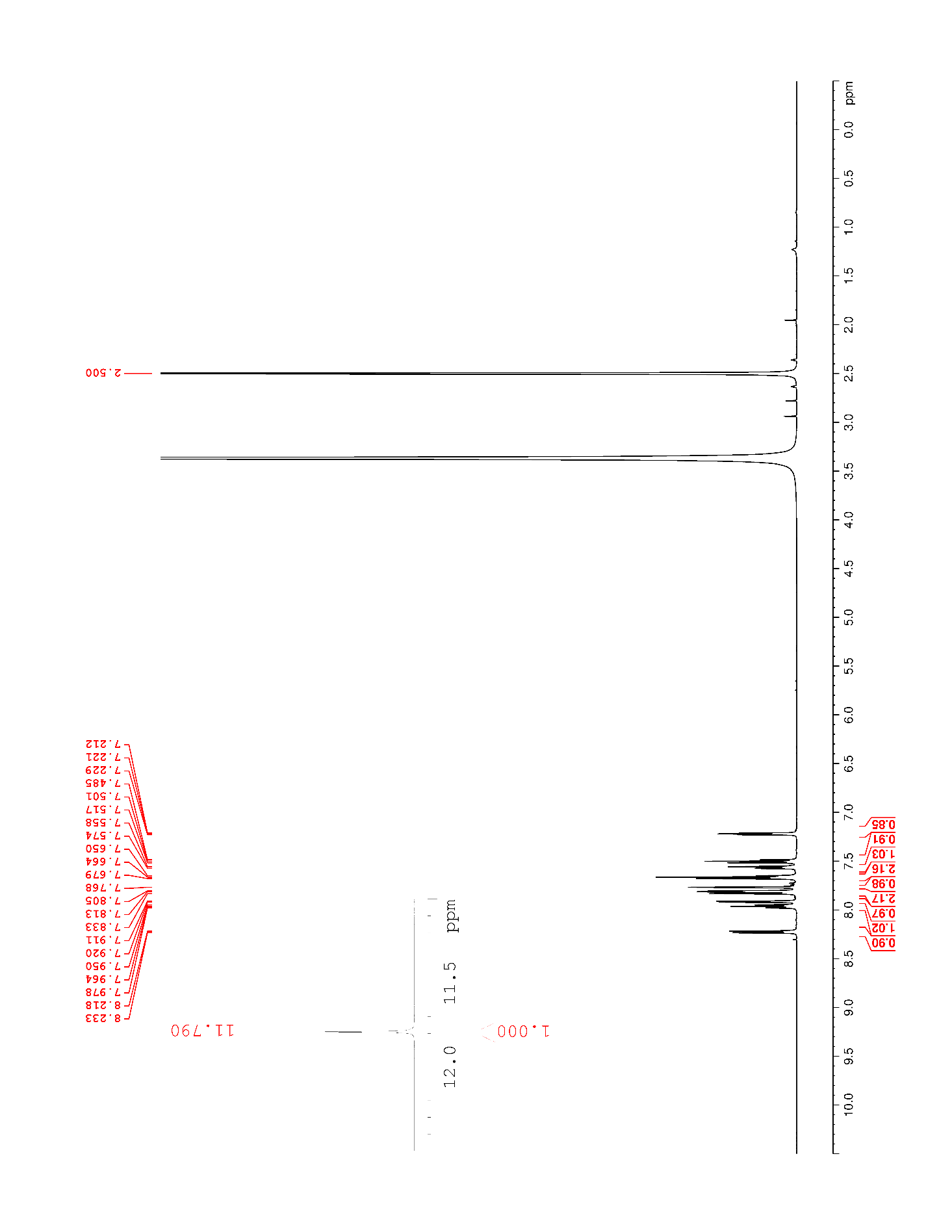

### Supplementary Figure 45. ^13^C NMR (125 MHz, DMSO-*d*_6_) of 17

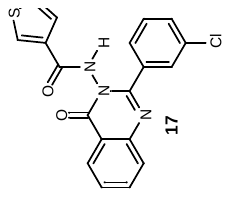

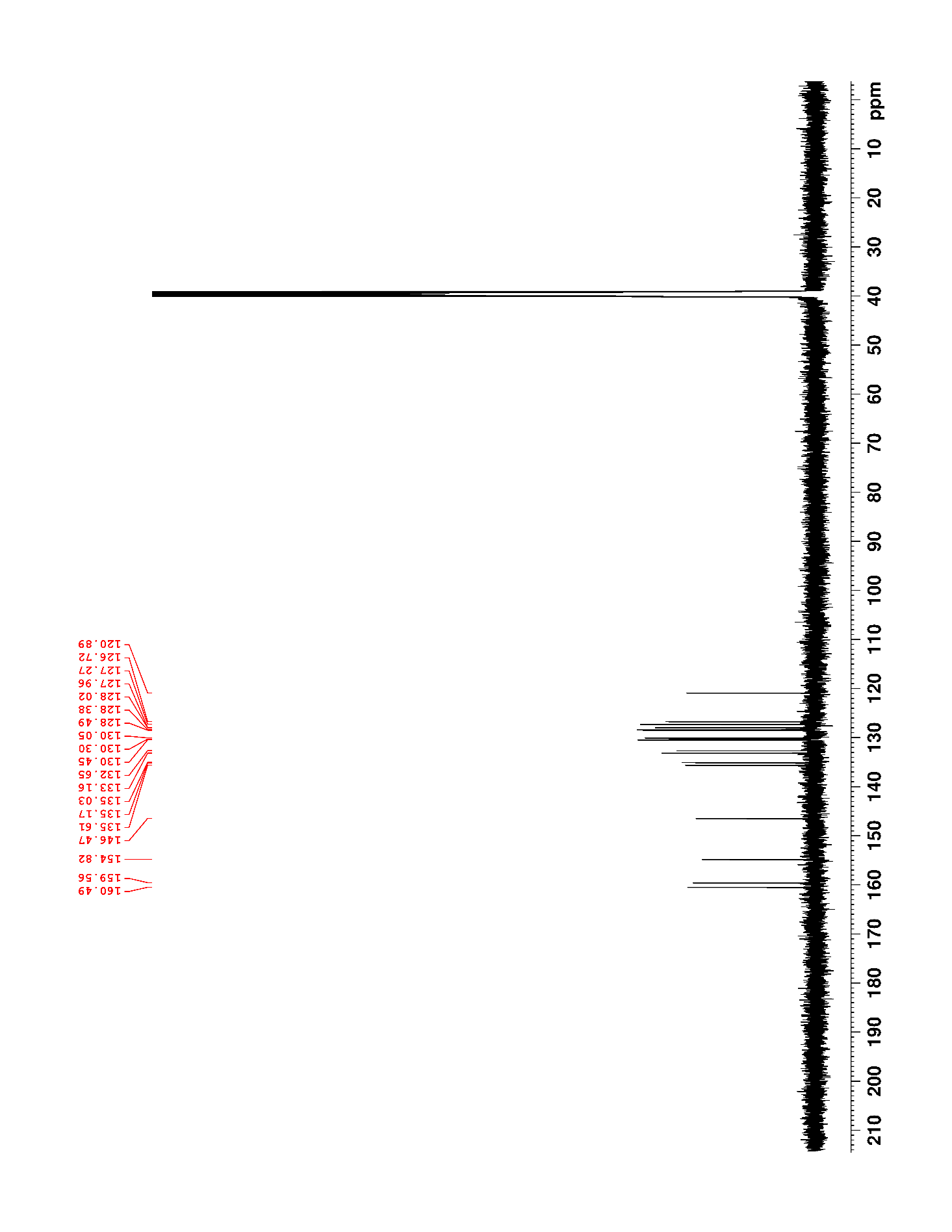

### Supplementary Figure 46. ^1^H NMR (500 MHz, DMSO-*d*_6_) of 18

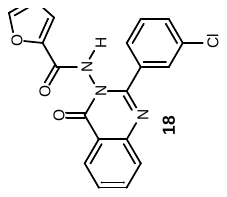

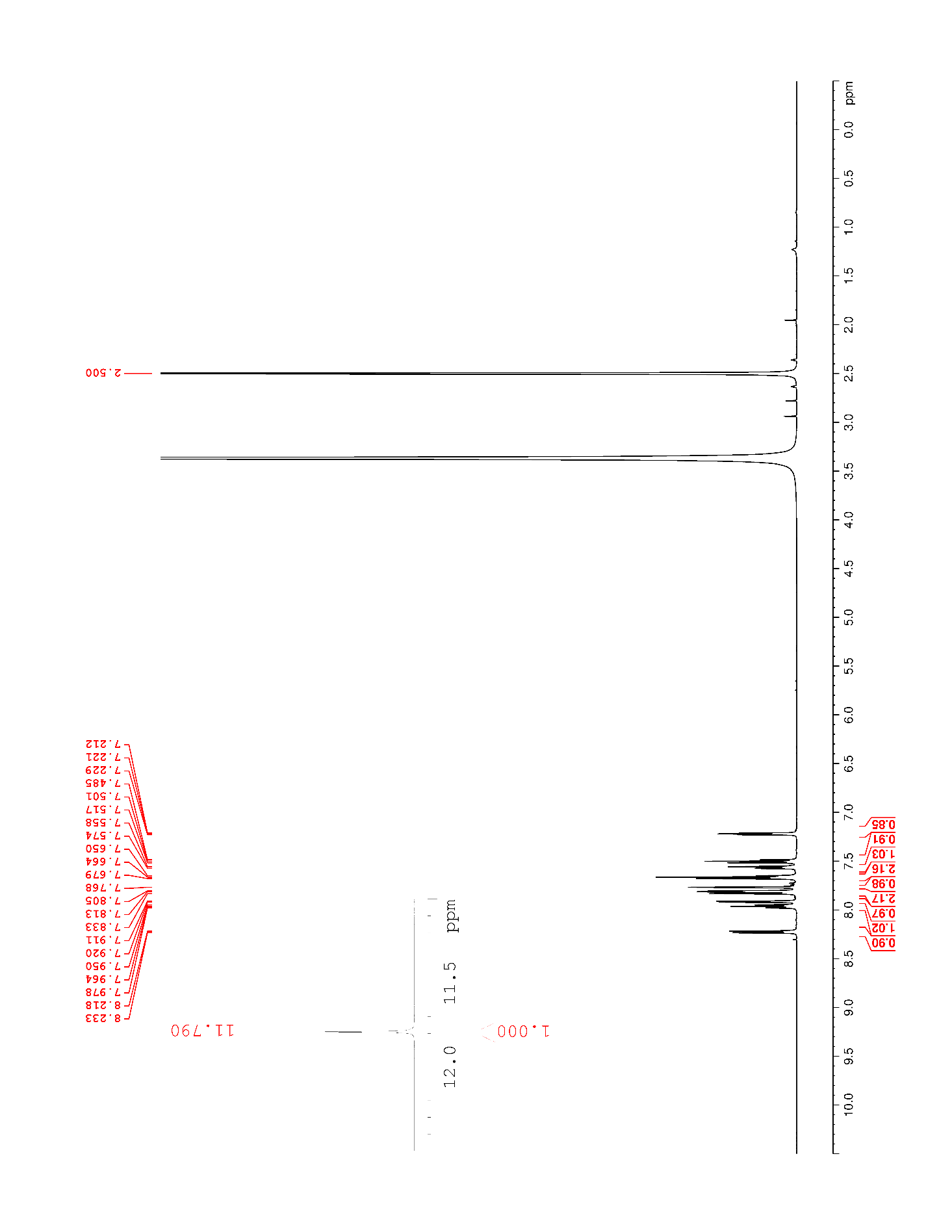

### Supplementary Figure 47. ^13^C NMR (125 MHz, DMSO-*d*_6_) of 18

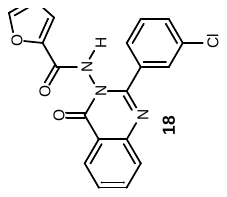

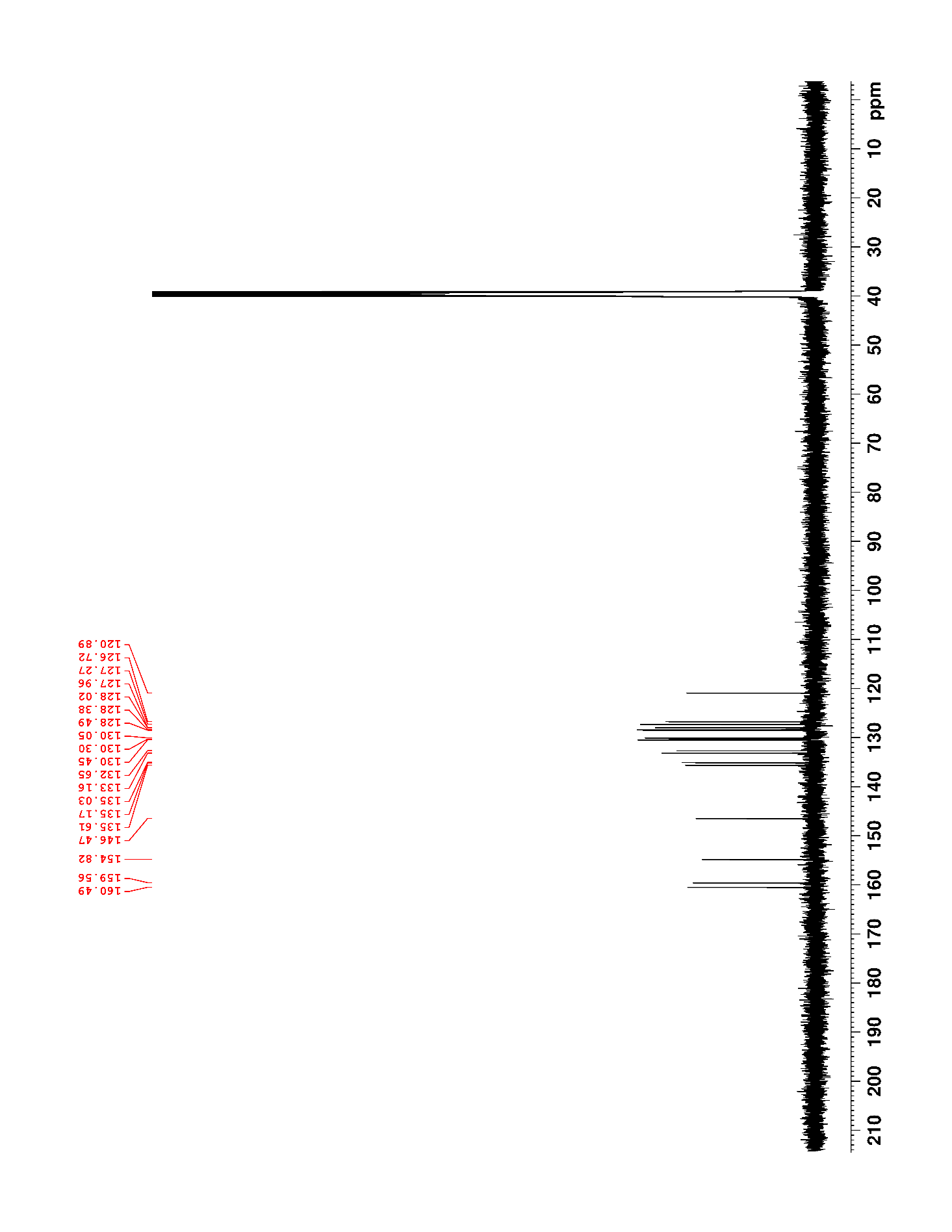

### Supplementary Figure 48. ^1^H NMR (500 MHz, DMSO-*d*_6_) of 19

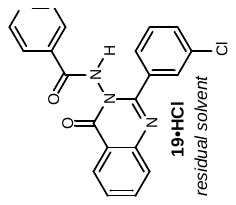

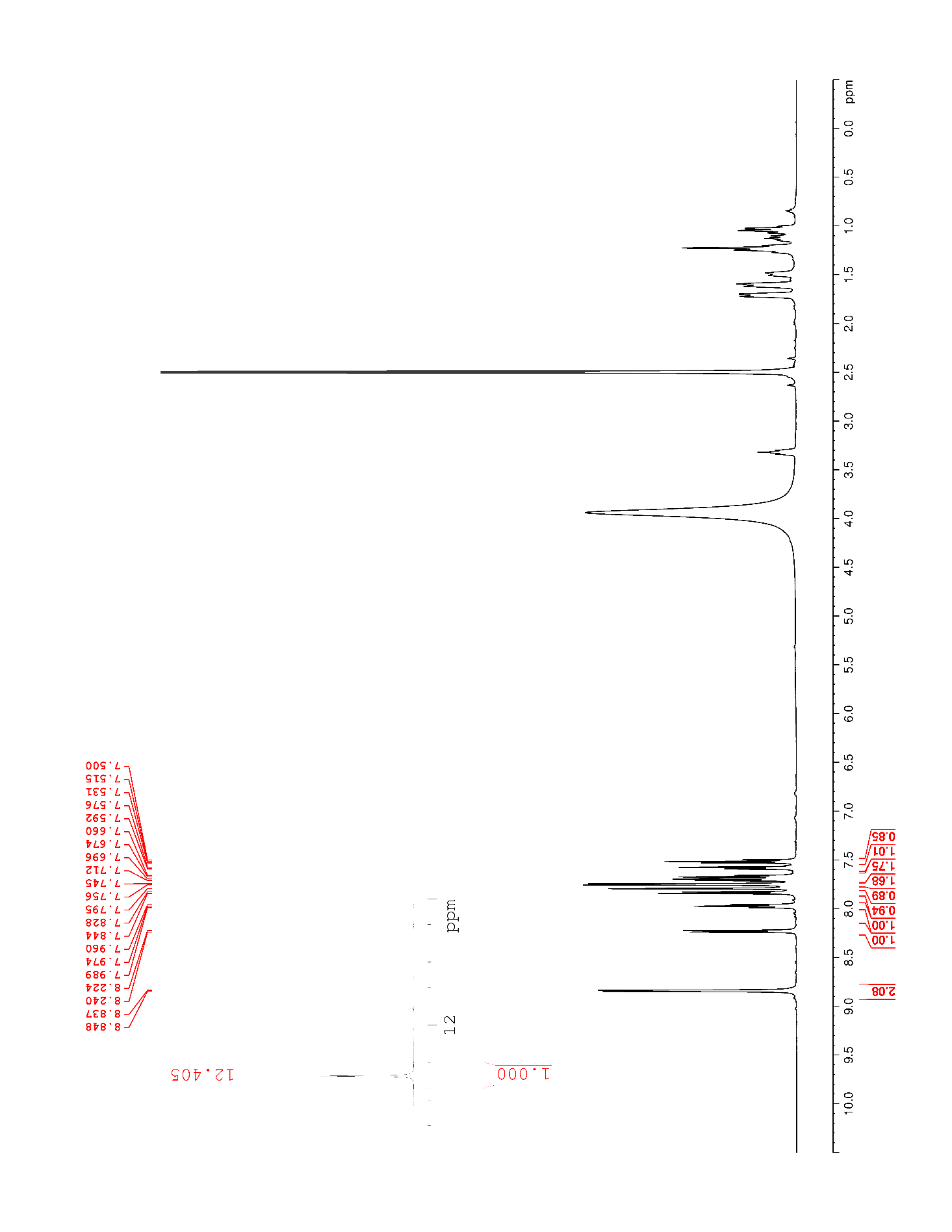

### Supplementary Figure 49. ^13^C NMR (125 MHz, DMSO-*d*_6_) of 19

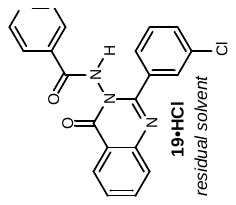

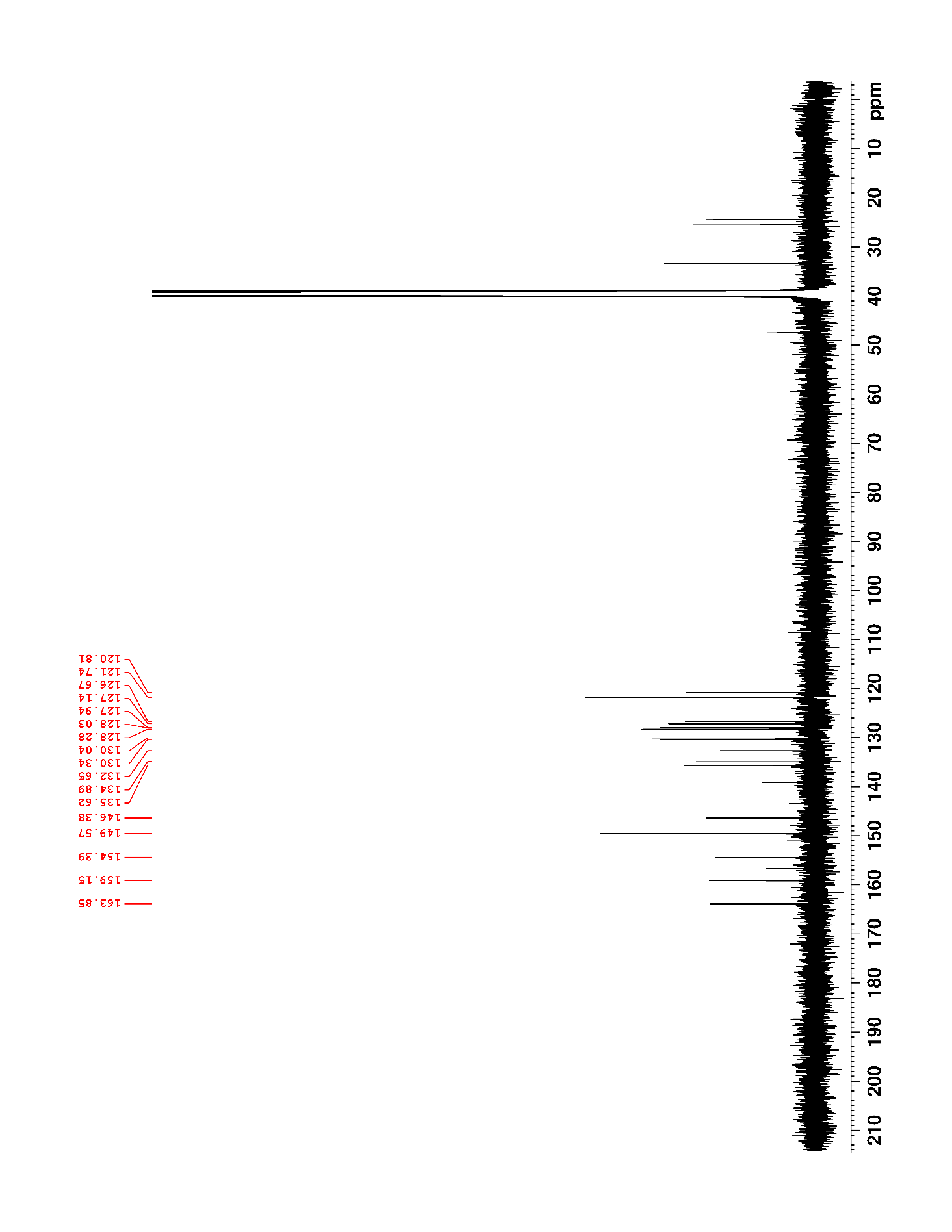

### Supplementary Figure 50. ^1^H NMR (500 MHz, DMSO-*d*_6_) of 20 (BZQ)

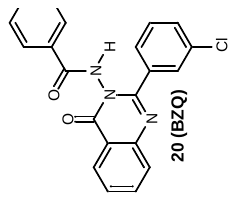

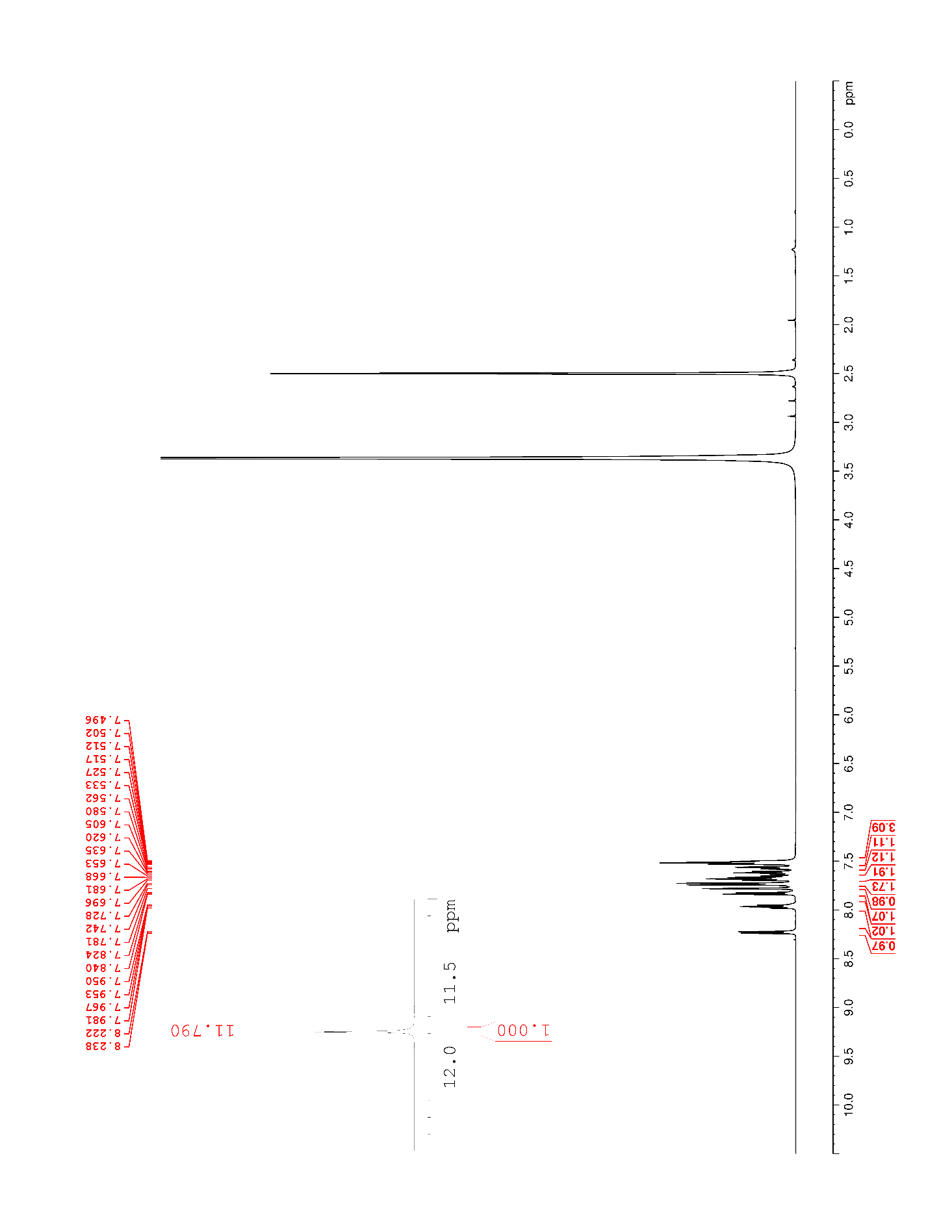

### Supplementary Figure 51. ^13^C NMR (125 MHz, DMSO-*d*_6_) of 20 (BZQ)

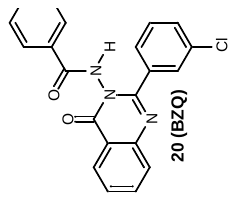

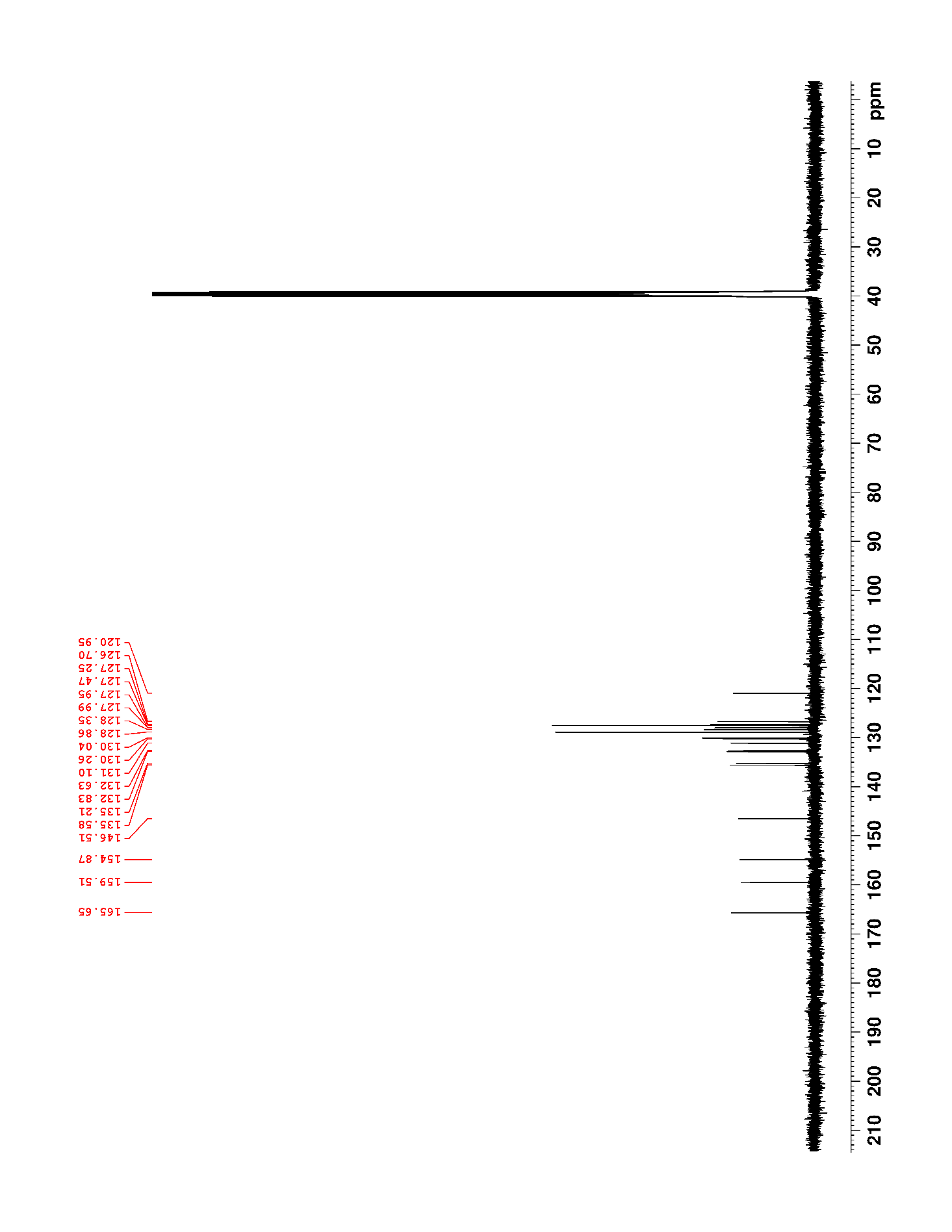

### Supplementary Figure 52. ^1^H NMR (500 MHz, DMSO-*d*_6_) of 21

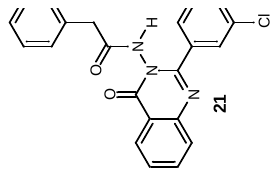

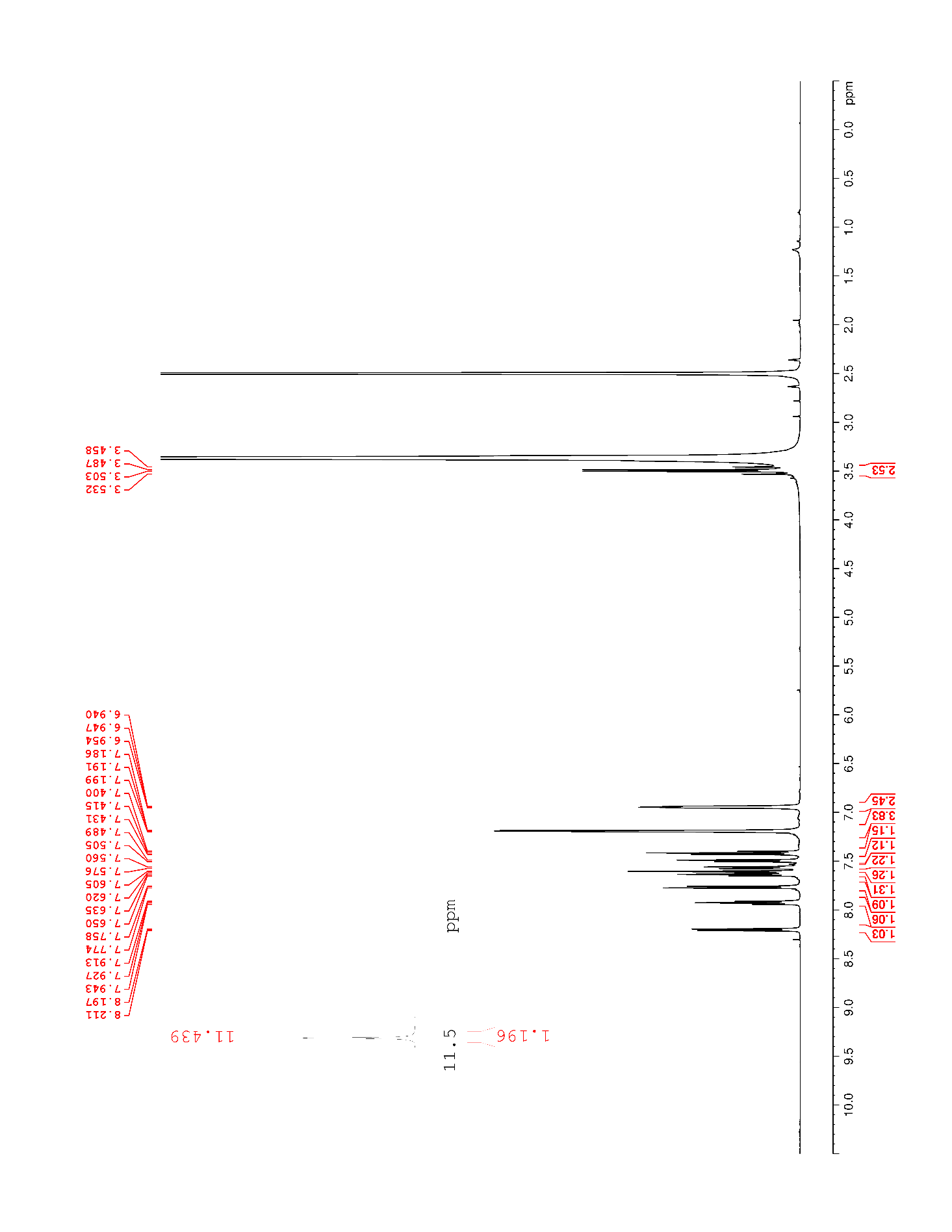

### Supplementary Figure 53. ^13^C NMR (125 MHz, DMSO-*d*_6_) of 21

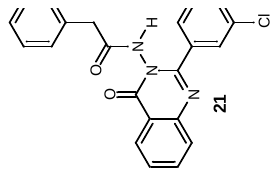

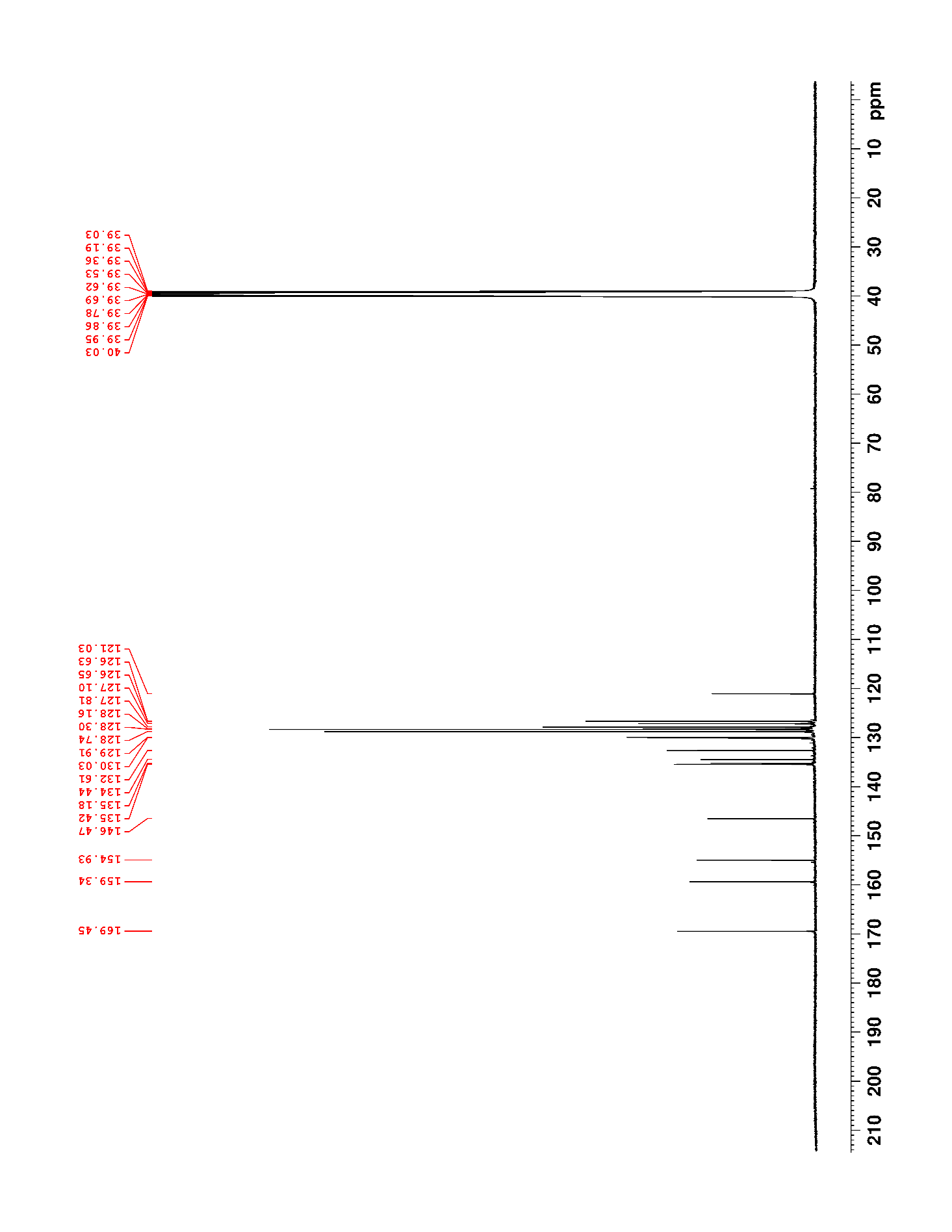

### Supplementary Figure 54. ^1^H NMR (500 MHz, DMSO-*d*_6_) of 22

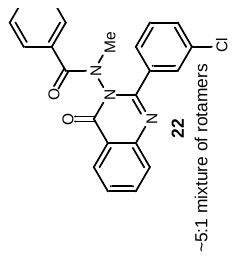

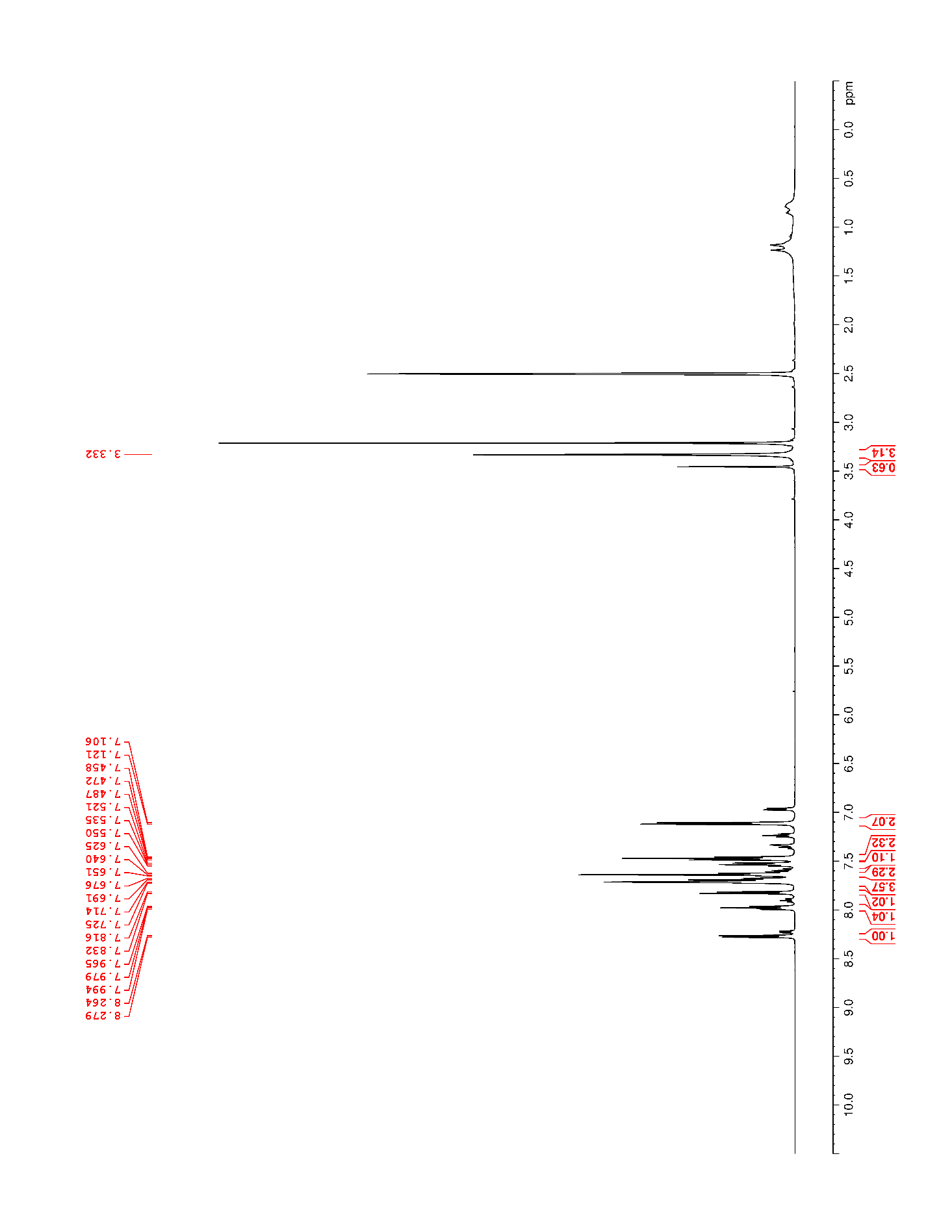

### Supplementary Figure 55. ^13^C NMR (125 MHz, DMSO-*d*_6_) of 22

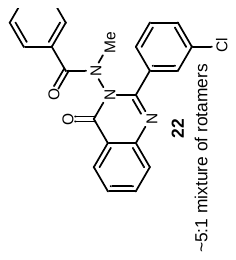

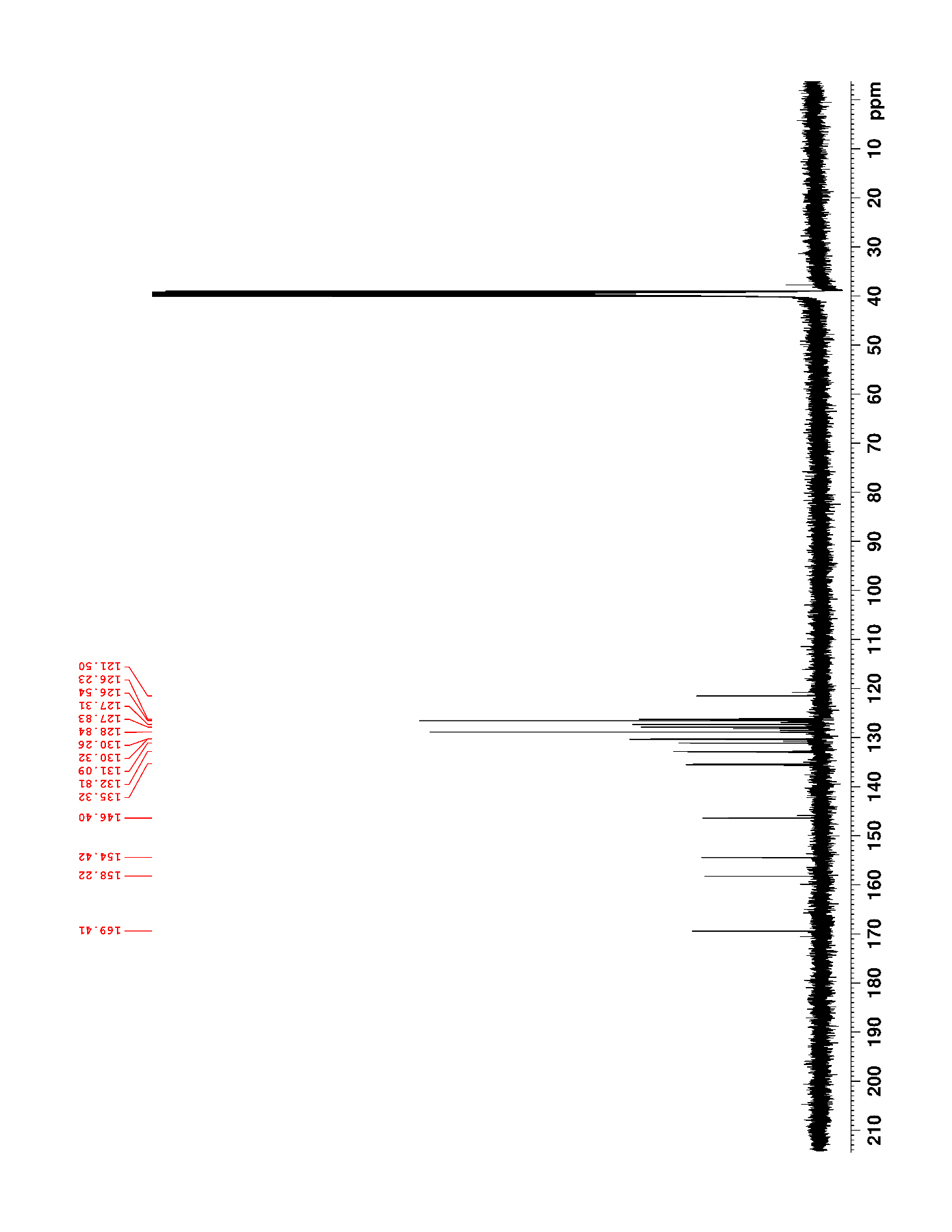

### Supplementary Figure 56. ^1^H NMR (500 MHz, DMSO-*d*_6_) of S3

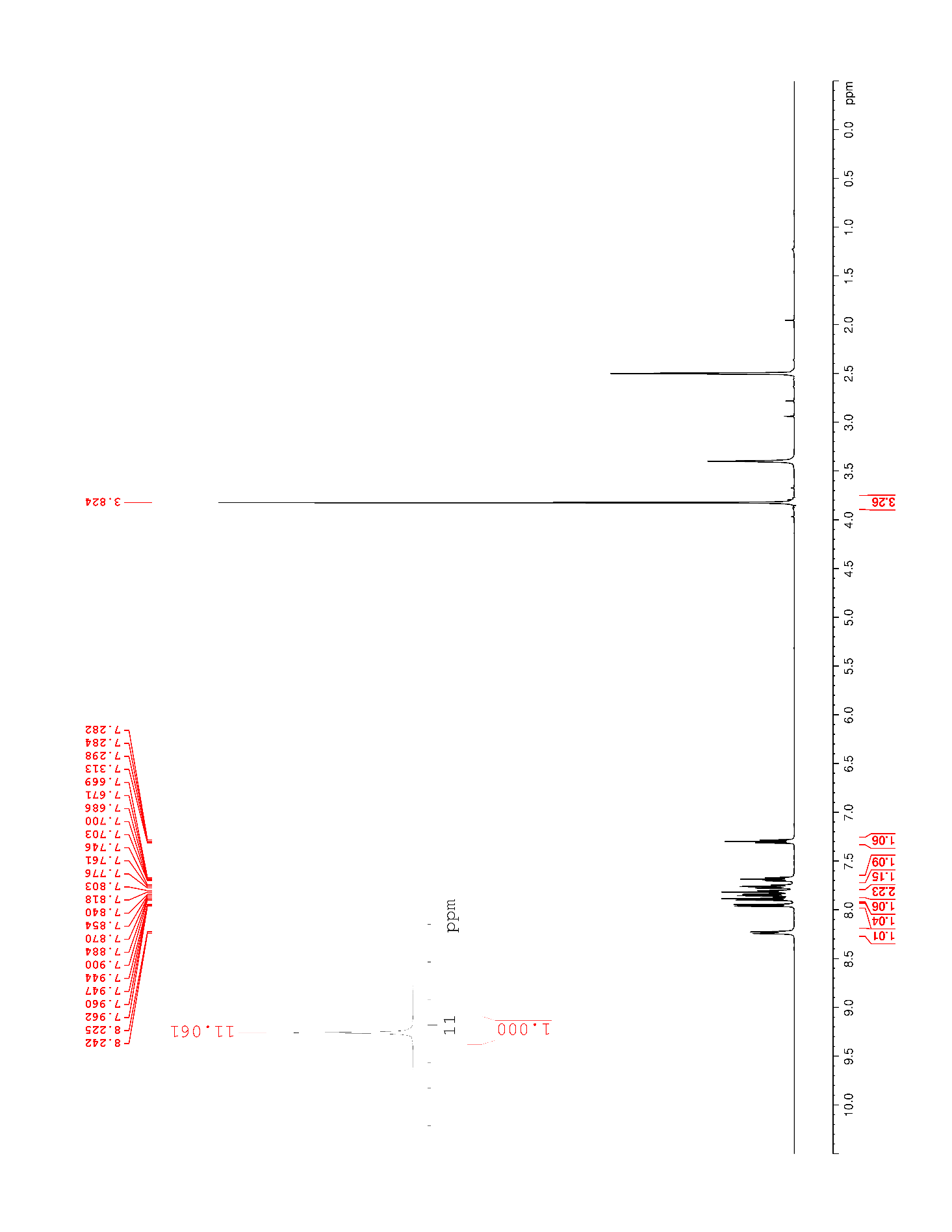

### Supplementary Figure 57. ^13^C NMR (125 MHz, DMSO-*d*_6_) of S3

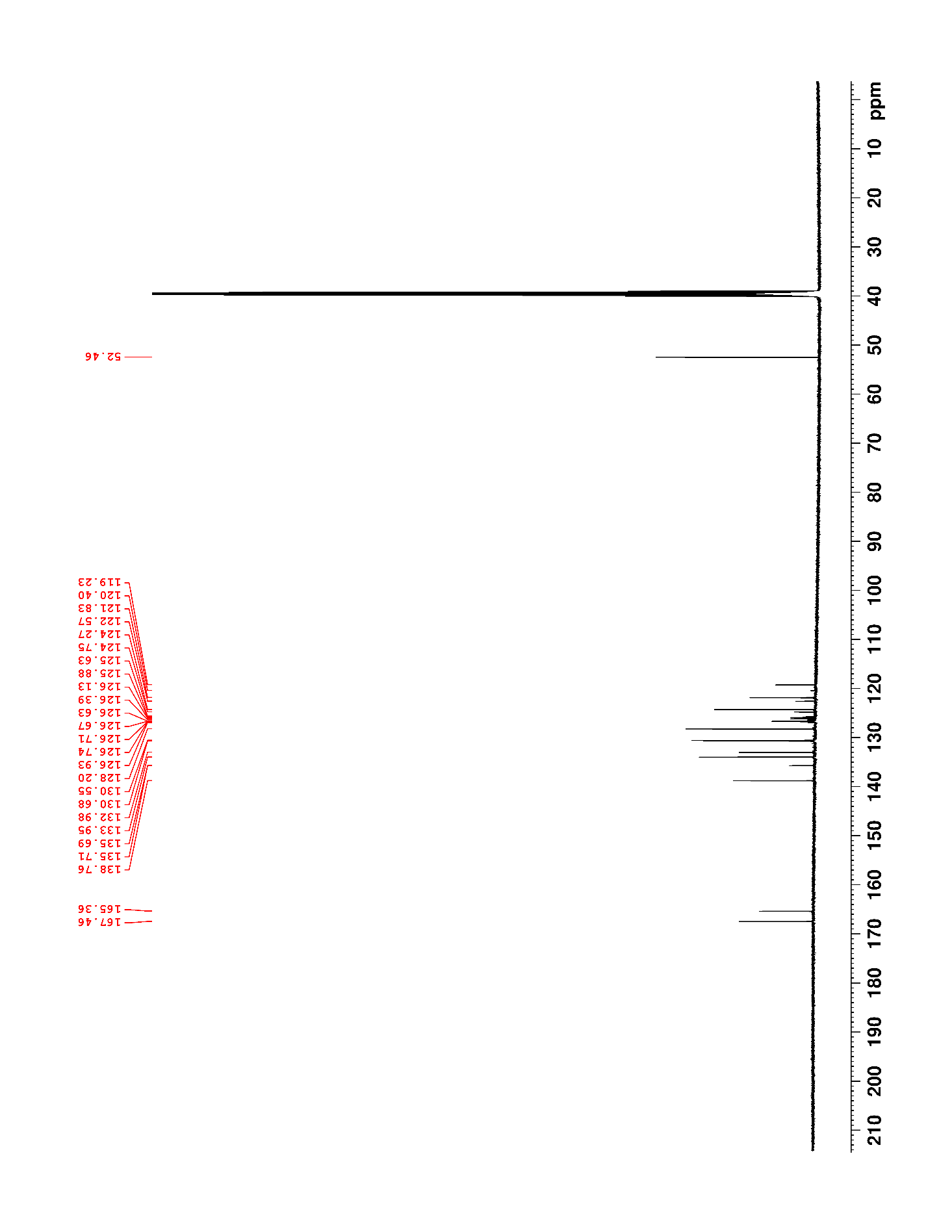

### Supplementary Figure 58. ^19^F NMR (470 MHz, DMSO-*d*_6_) of S3

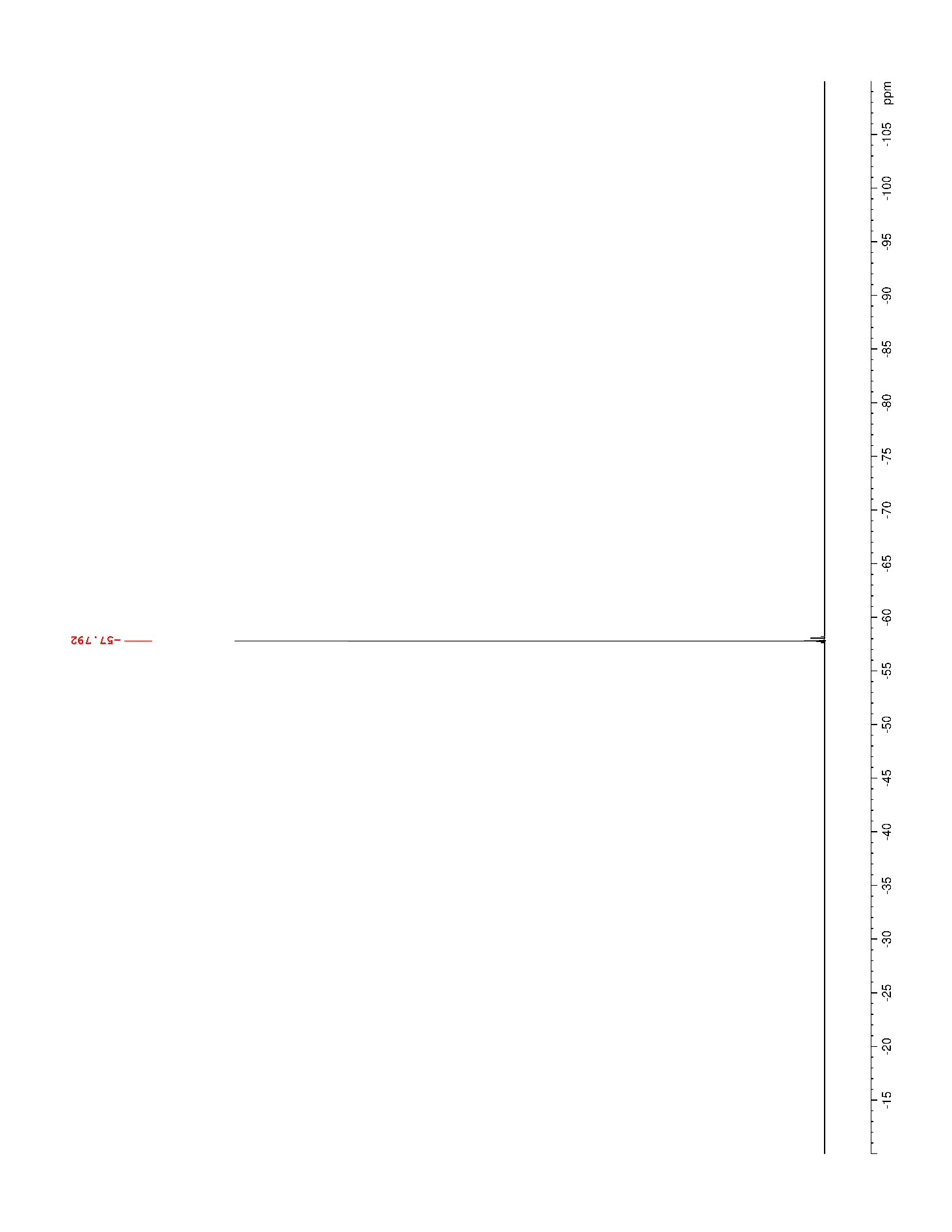

### Supplementary Figure 59. ^1^H NMR (500 MHz, DMSO-*d*_6_) of 25

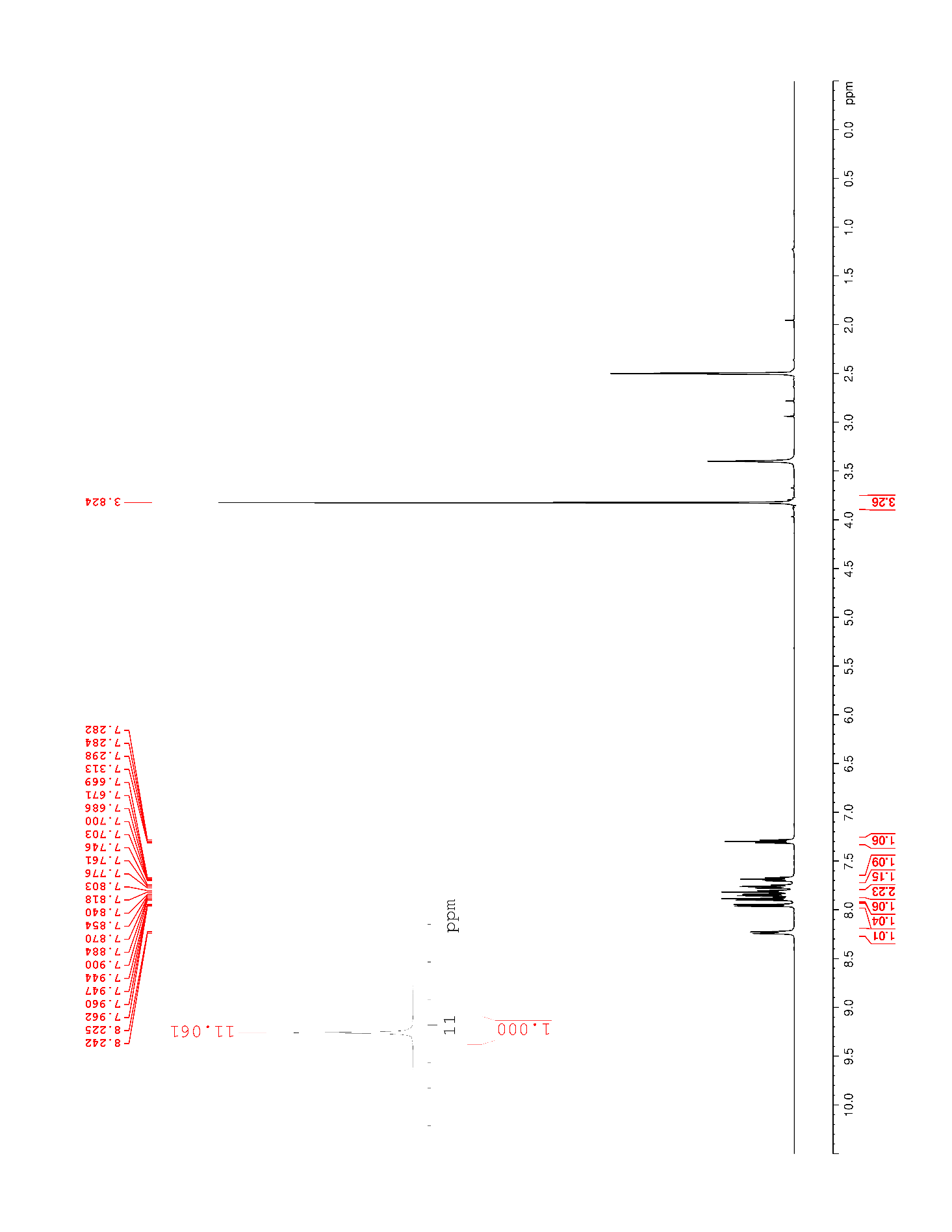

### Supplementary Figure 60. ^13^C NMR (125 MHz, DMSO-*d*_6_) of 25

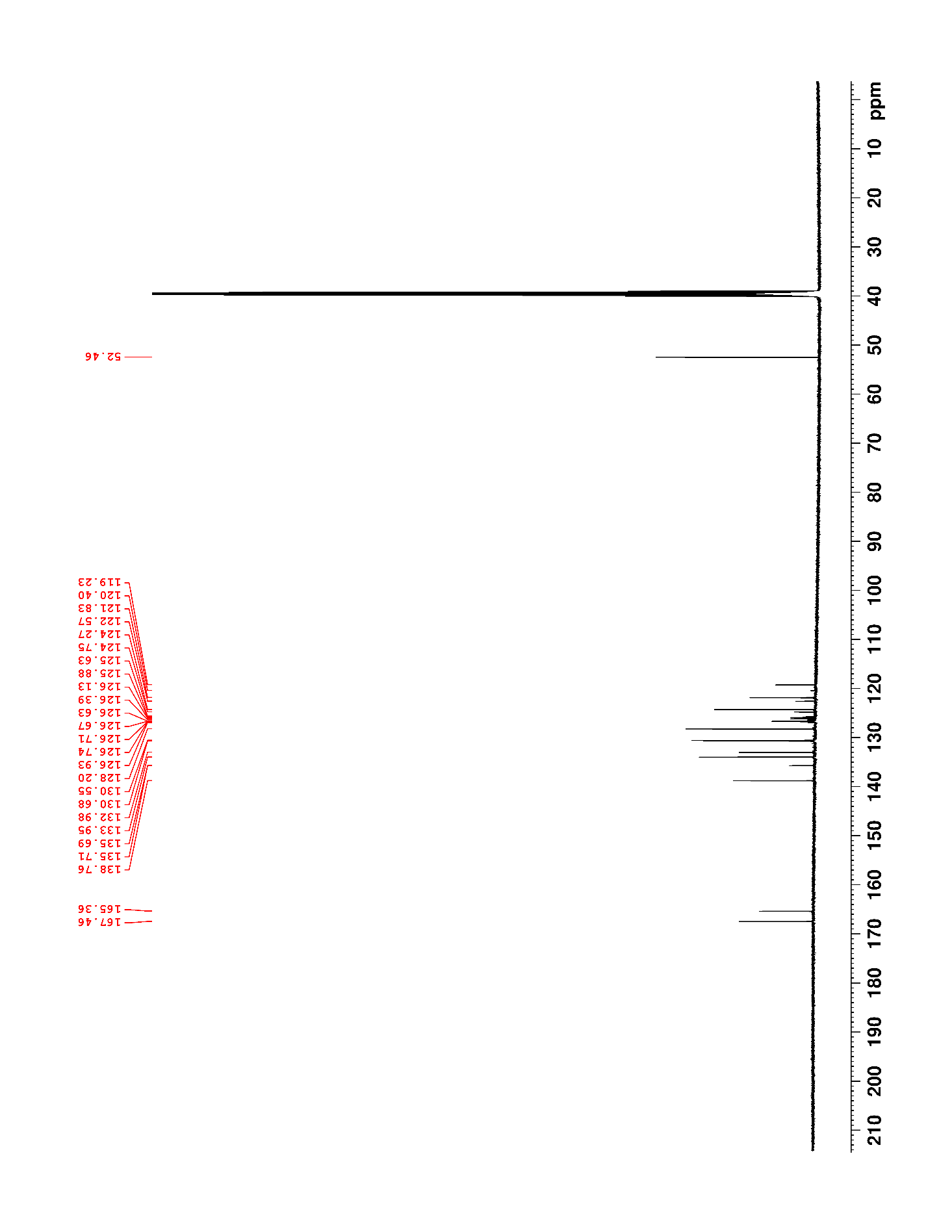

### Supplementary Figure 61. ^19^F NMR (470 MHz, DMSO-*d*_6_) of 25

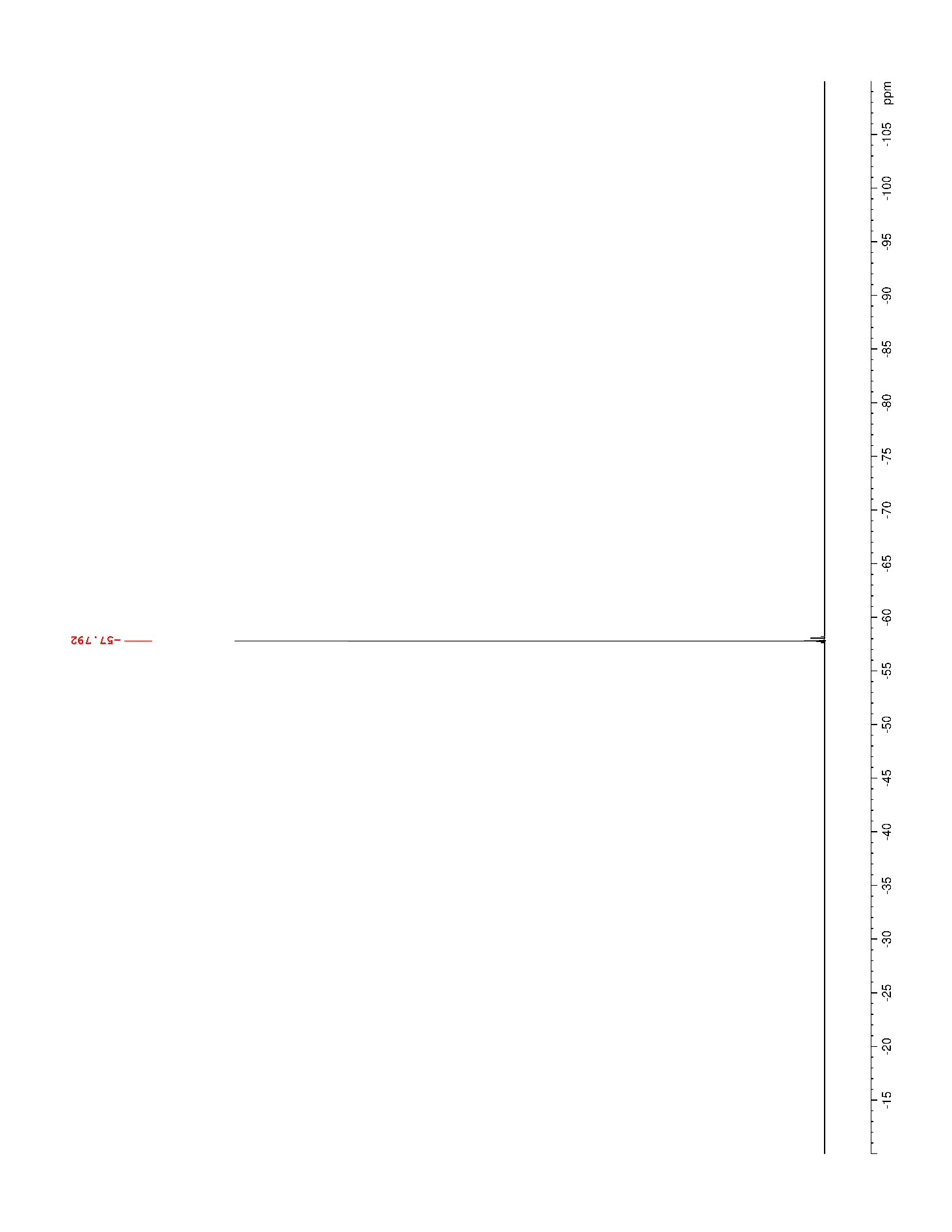

### Supplementary Figure 62. ^1^H NMR (500 MHz, DMSO-*d*_6_) of 26

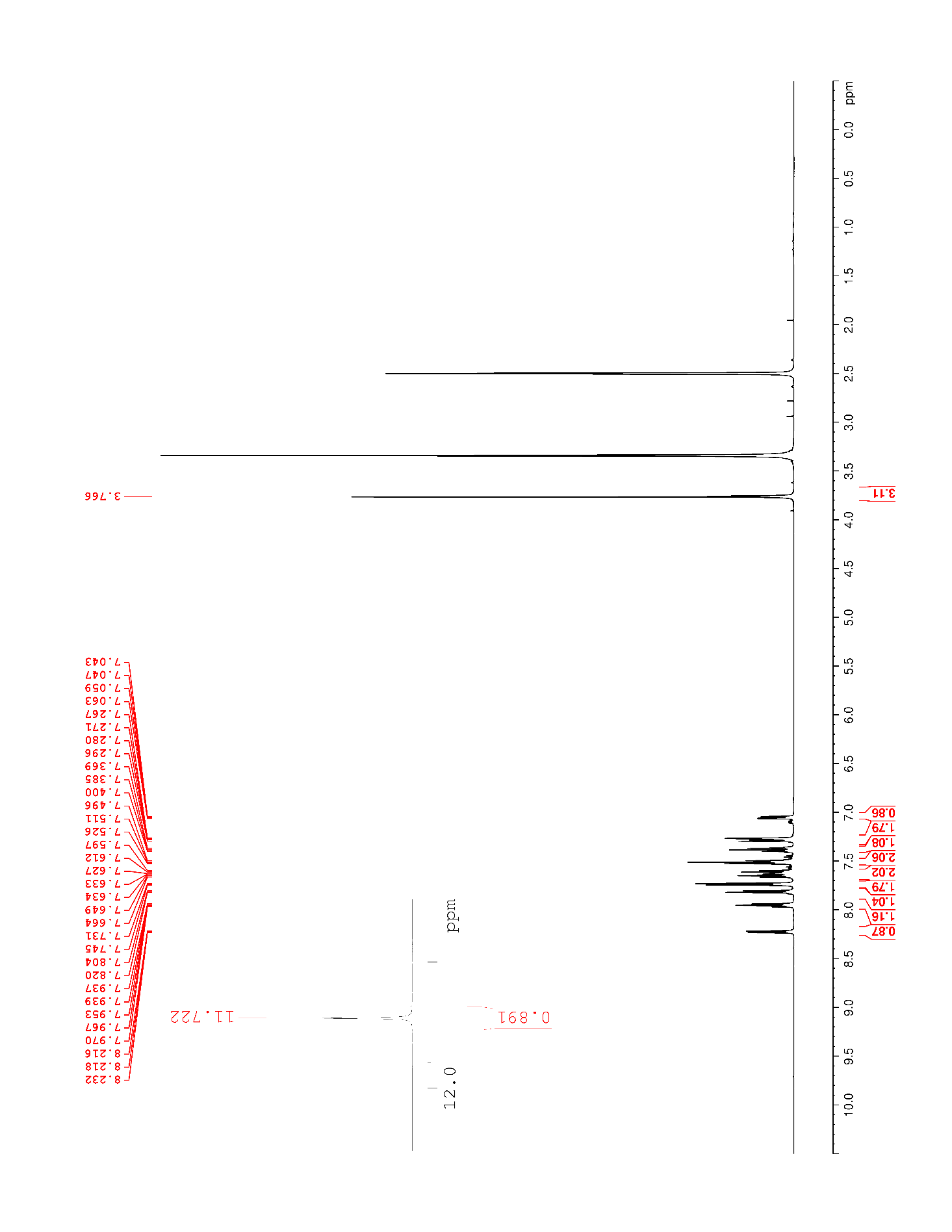

### Supplementary Figure 63. ^13^C NMR (125 MHz, DMSO-*d*_6_) of 26

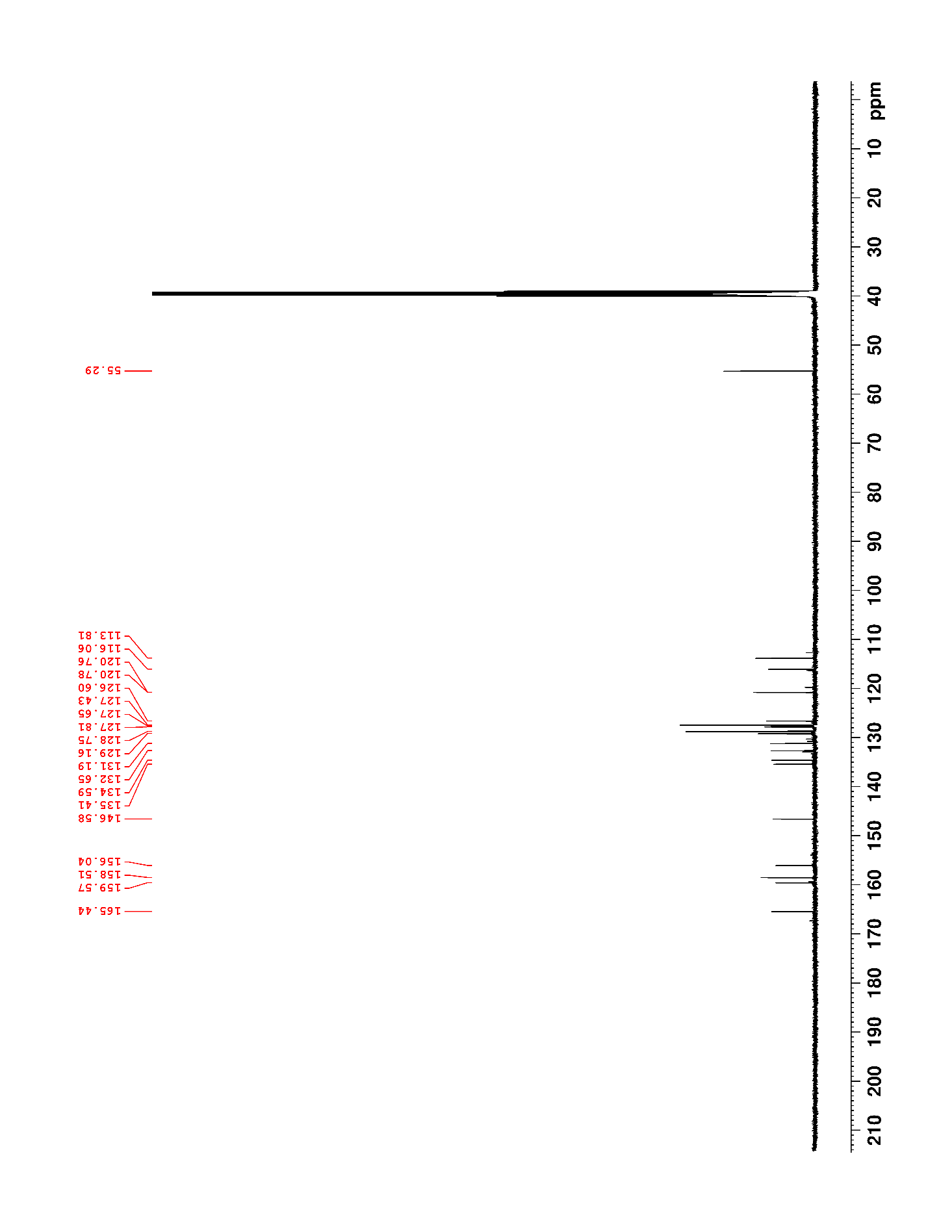

### Supplementary Figure 64. ^1^H NMR (500 MHz, DMSO-*d*_6_) of S7

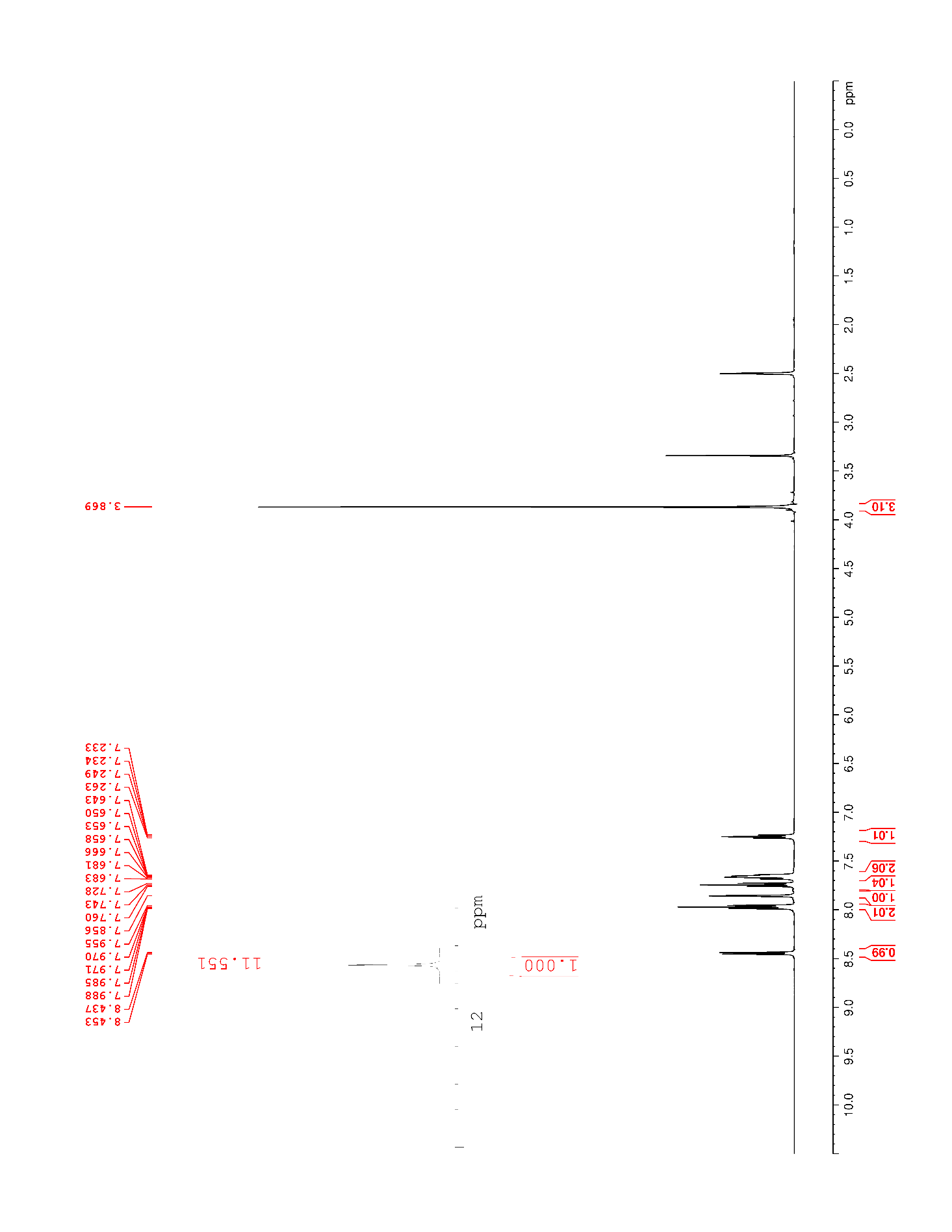

### Supplementary Figure 65. ^13^C NMR (125 MHz, DMSO-*d*_6_) of S7

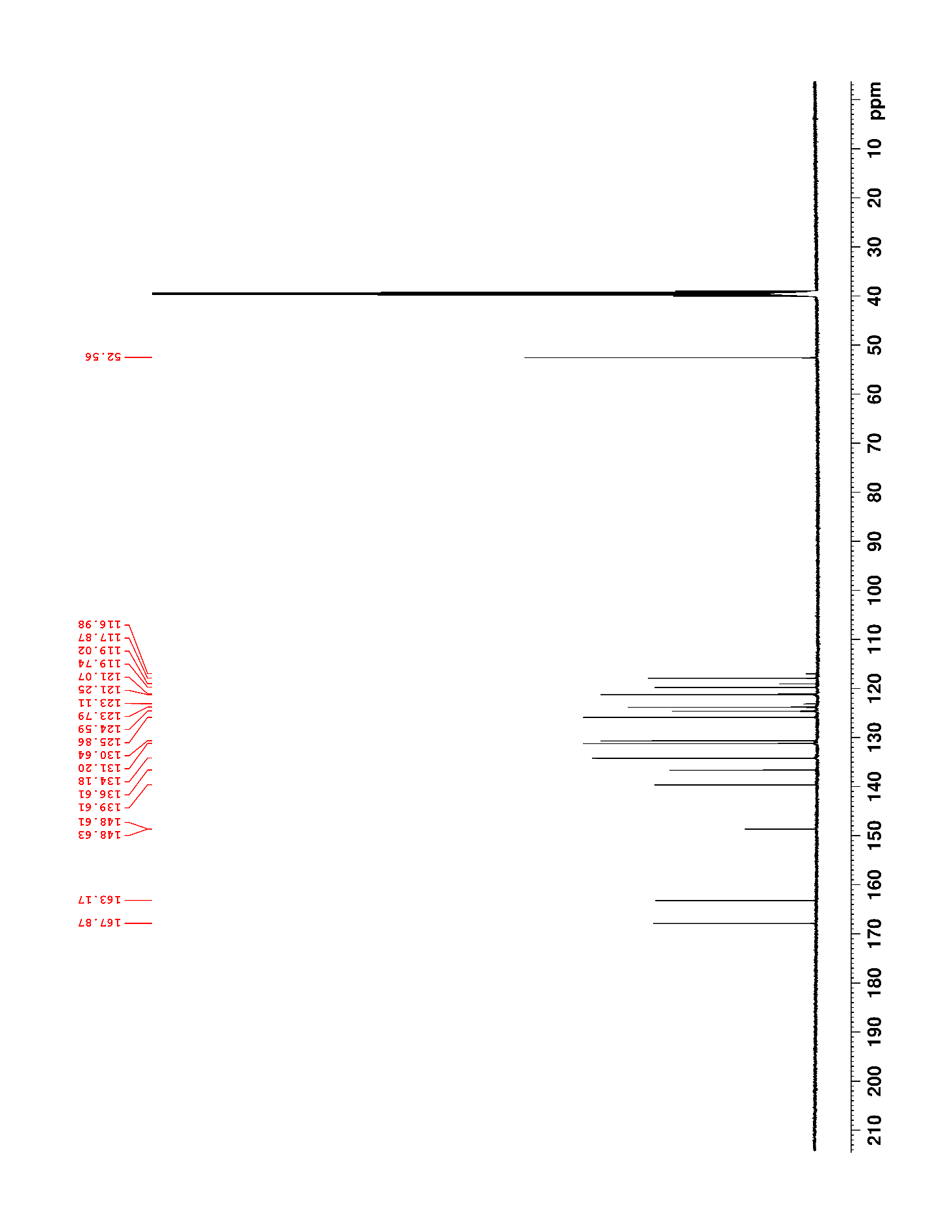

### Supplementary Figure 66. ^19^F NMR (470 MHz, DMSO-*d*_6_) of S7

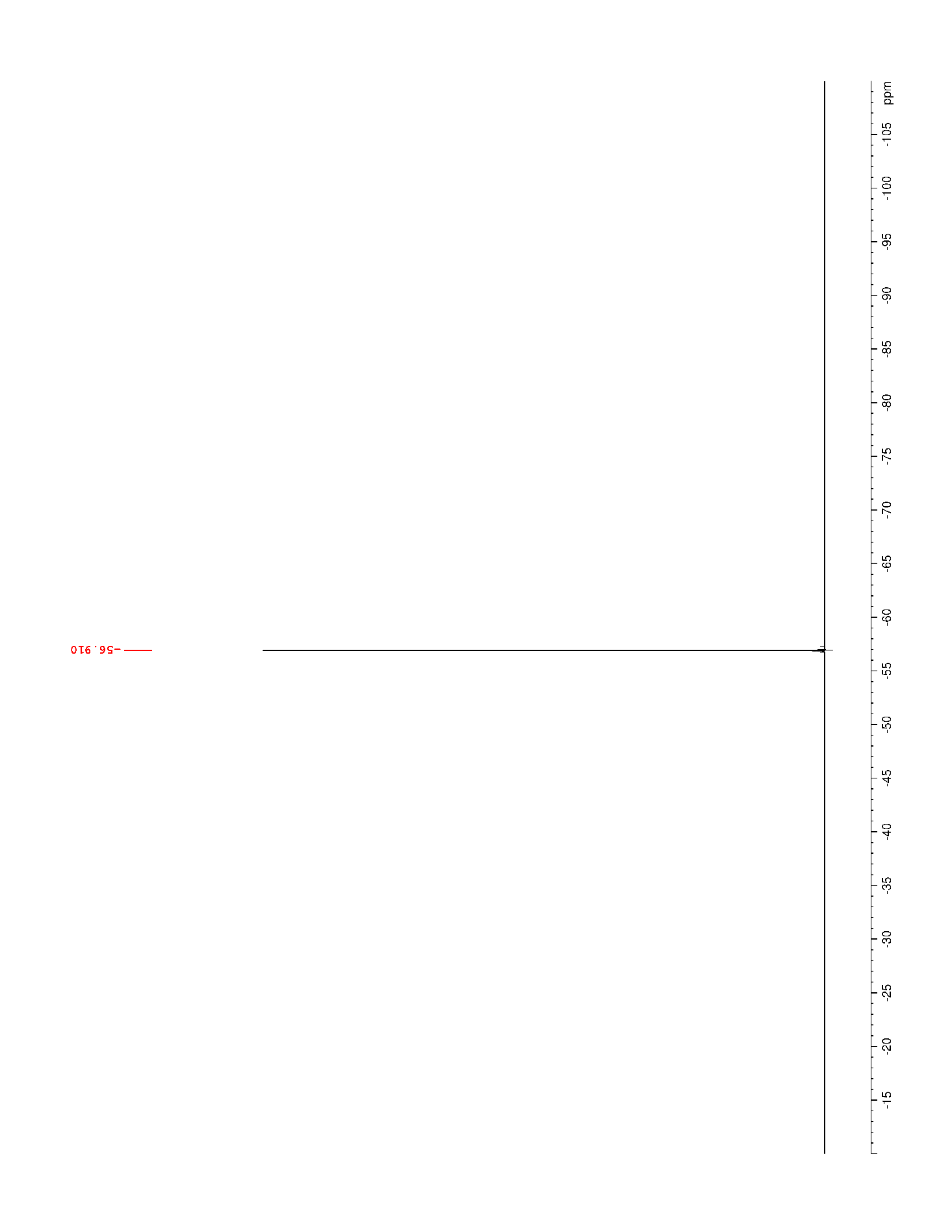

### Supplementary Figure 67. ^1^H NMR (500 MHz, DMSO-*d*_6_) of S8

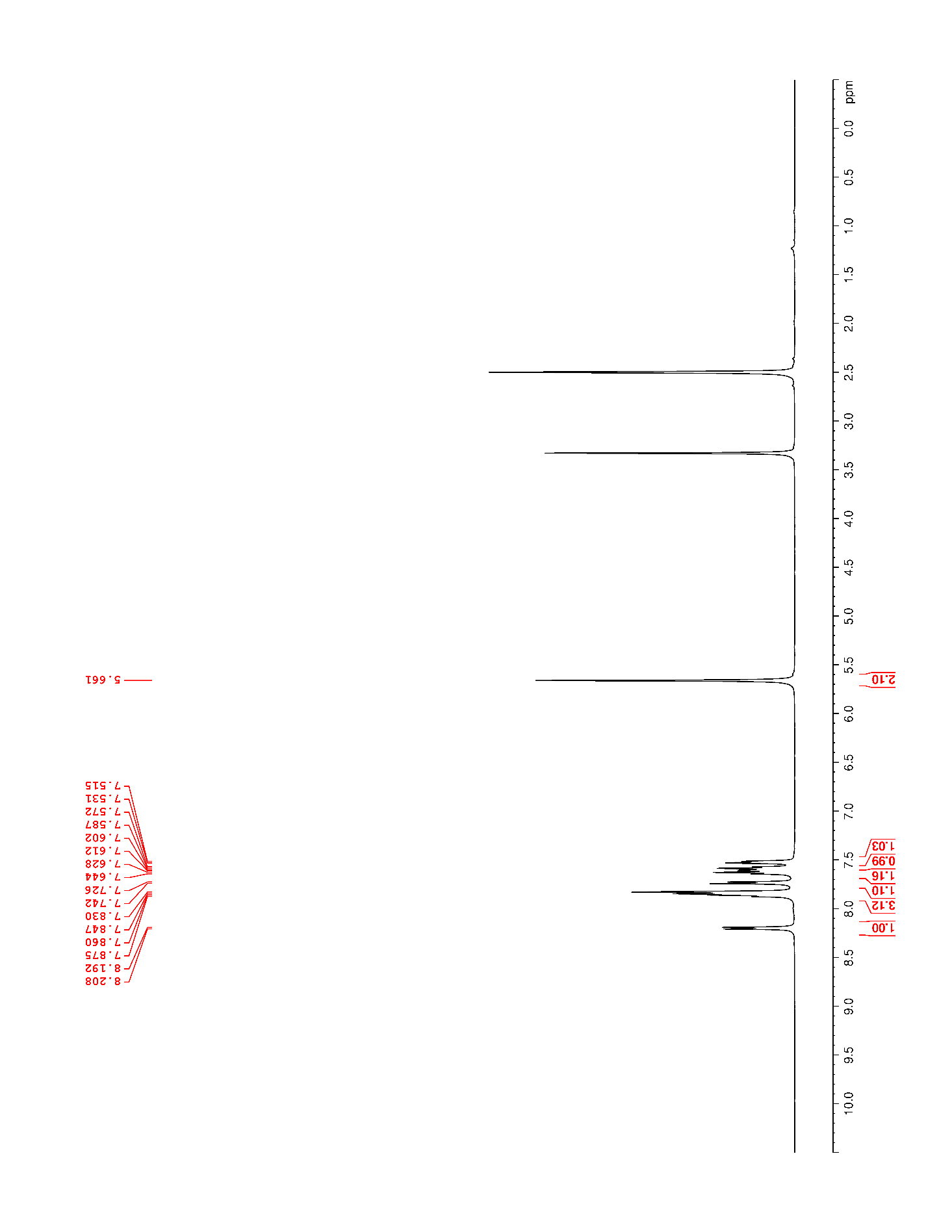

### Supplementary Figure 68. ^13^C NMR (125 MHz, DMSO-*d*_6_) of S8

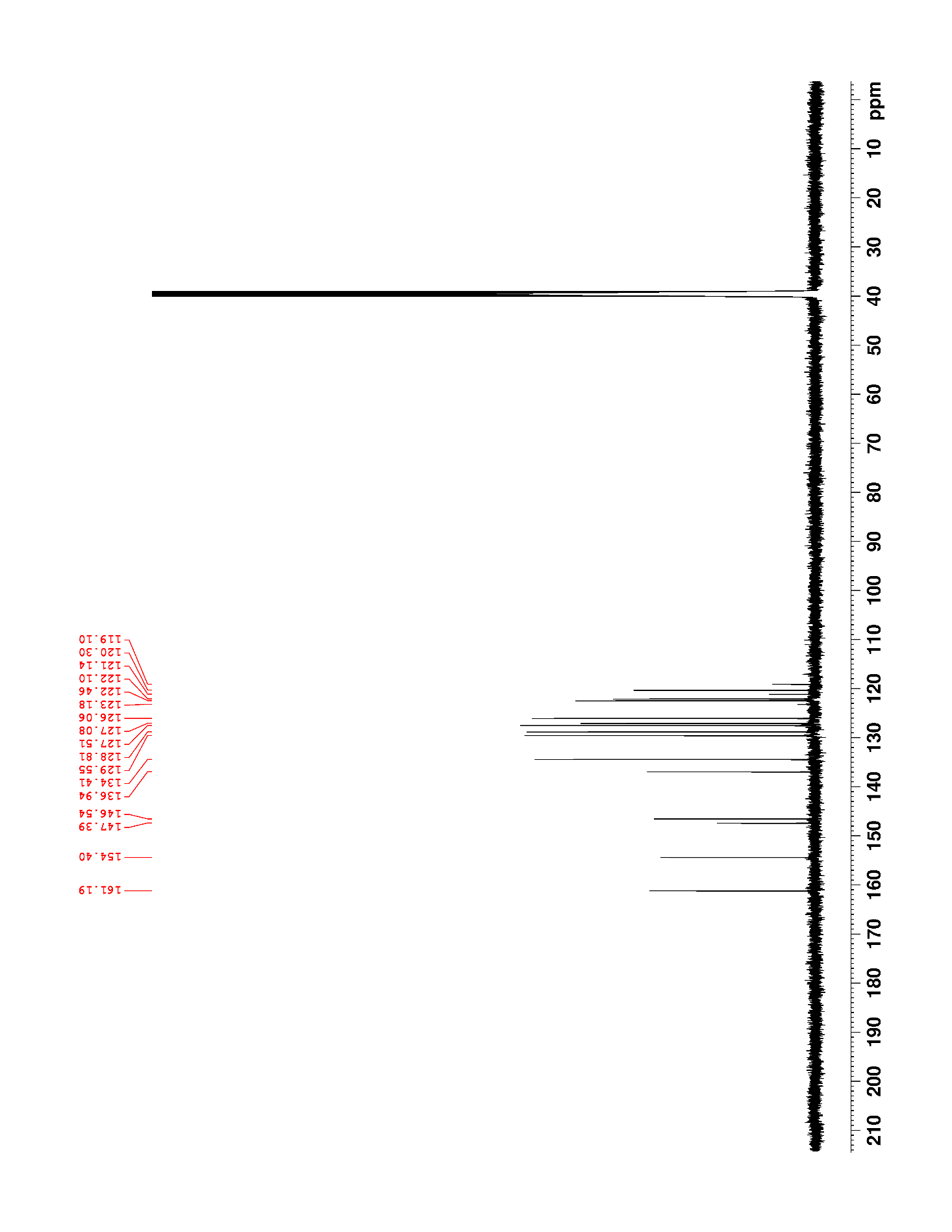

### Supplementary Figure 69. ^19^F NMR (470 MHz, DMSO-*d*_6_) of S8

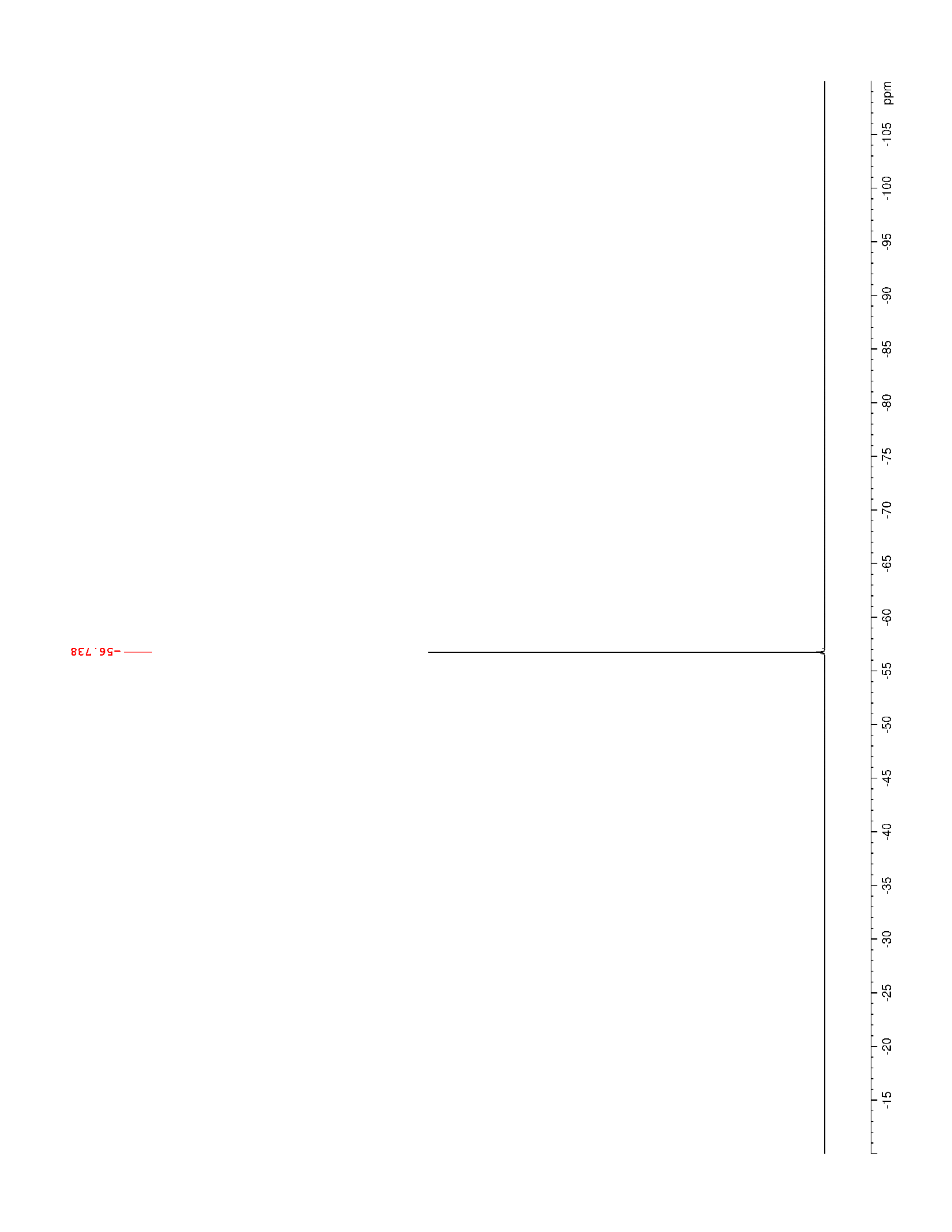

### Supplementary Figure 70. ^1^H NMR (500 MHz, DMSO-*d*_6_) of 27

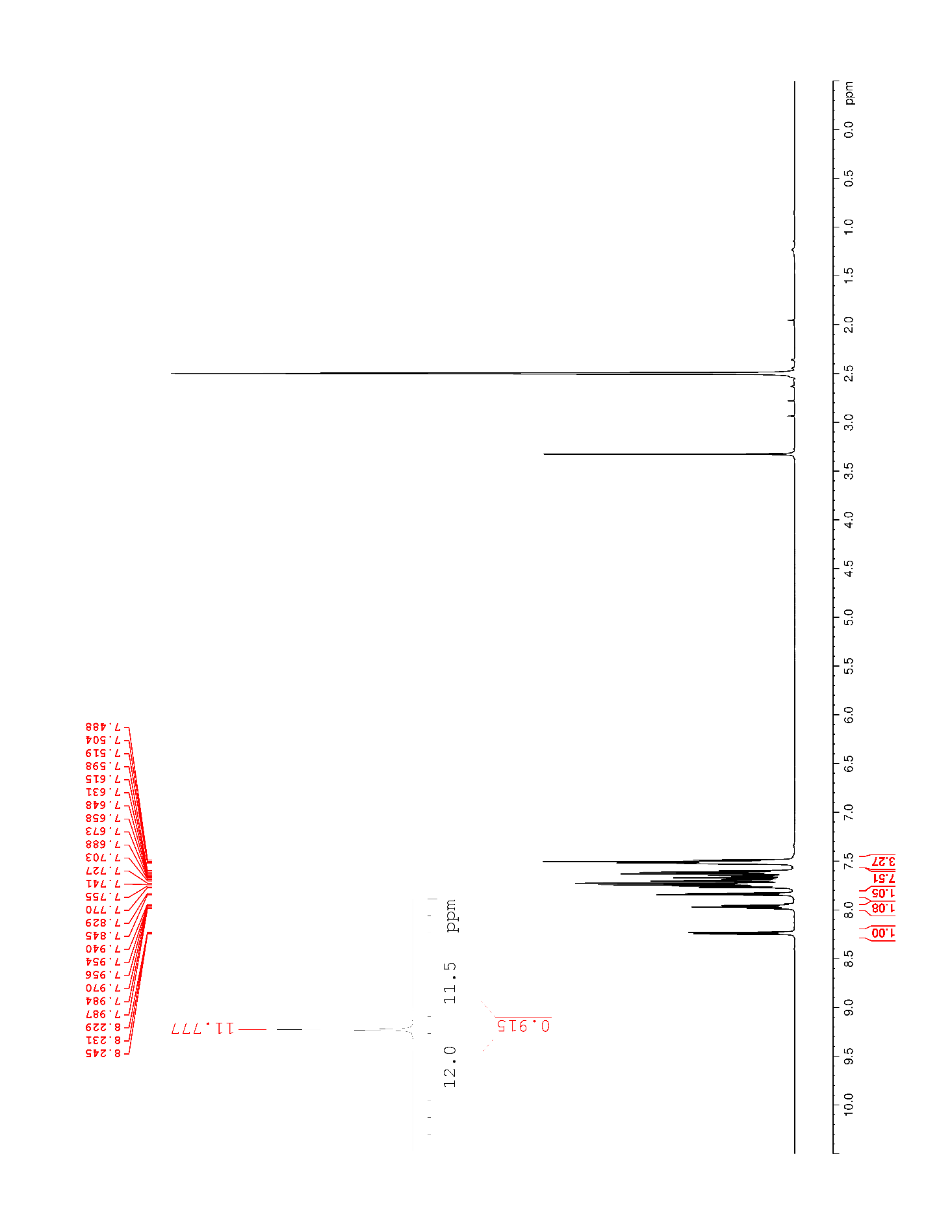

### Supplementary Figure 71. ^13^C NMR (125 MHz, DMSO-*d*_6_) of 27

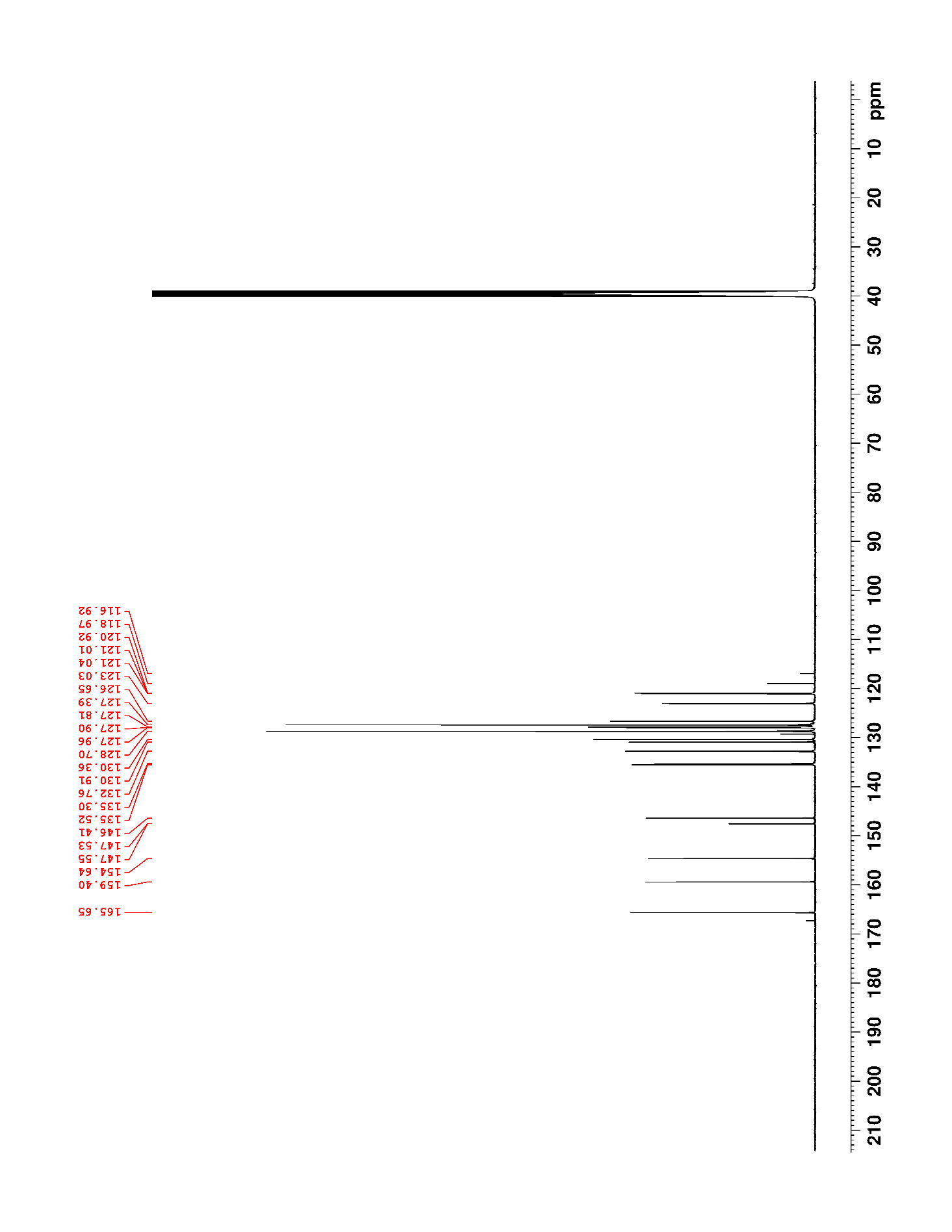

### Supplementary Figure 72. ^19^F NMR (470 MHz, DMSO-*d*_6_) of 27

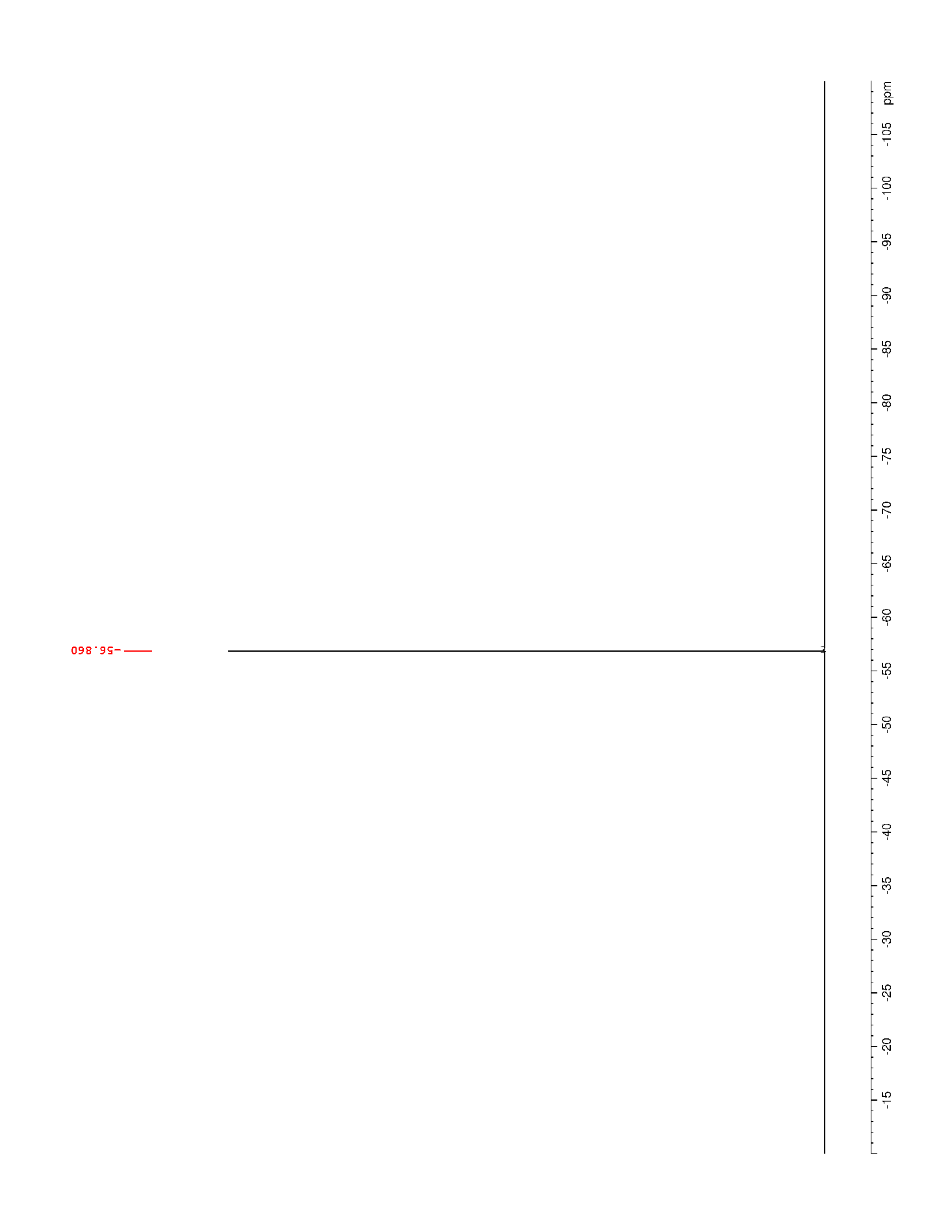

### Supplementary Figure 73. ^1^H NMR (500 MHz, DMSO-*d*_6_) of S9

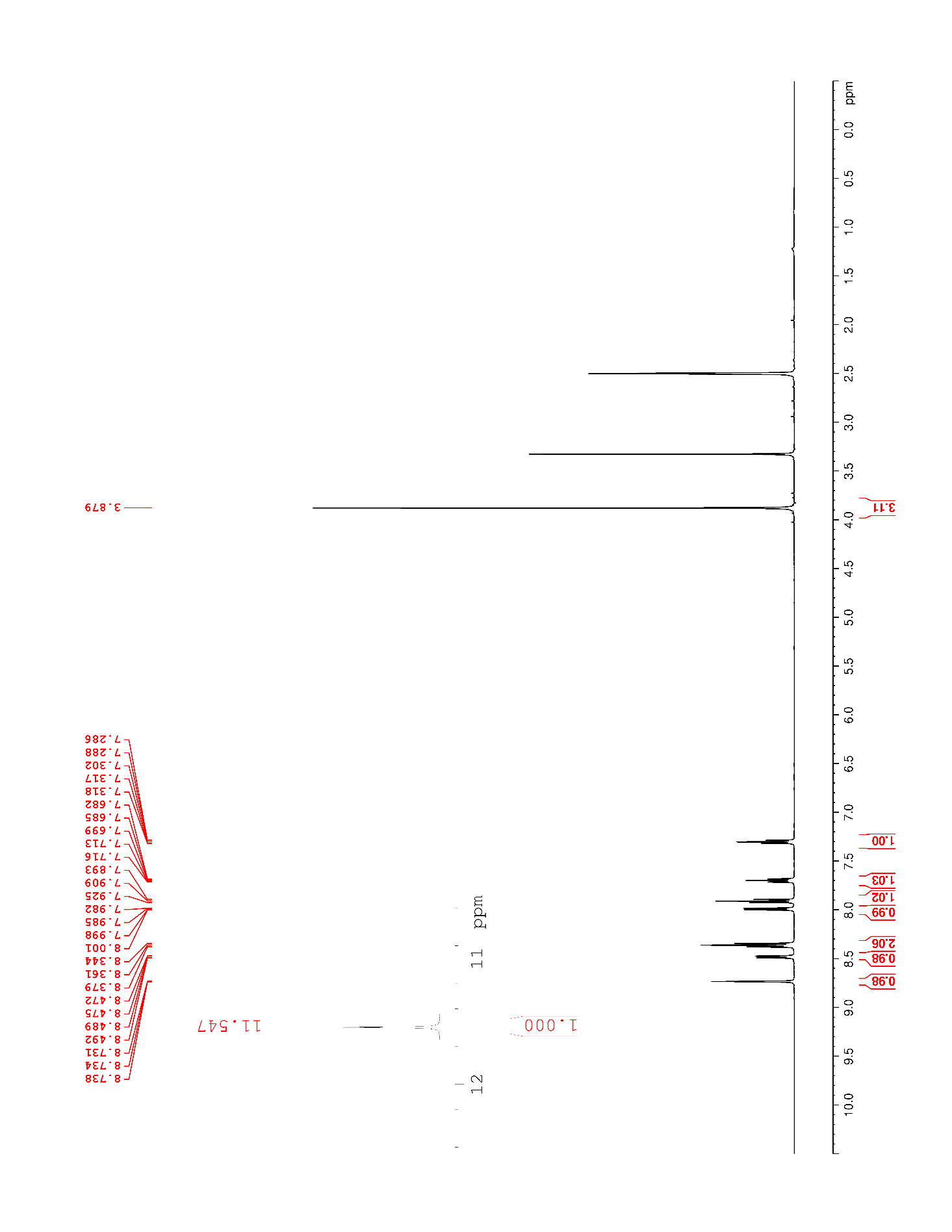

### Supplementary Figure 74. ^13^C NMR (125 MHz, DMSO-*d*_6_) of S9

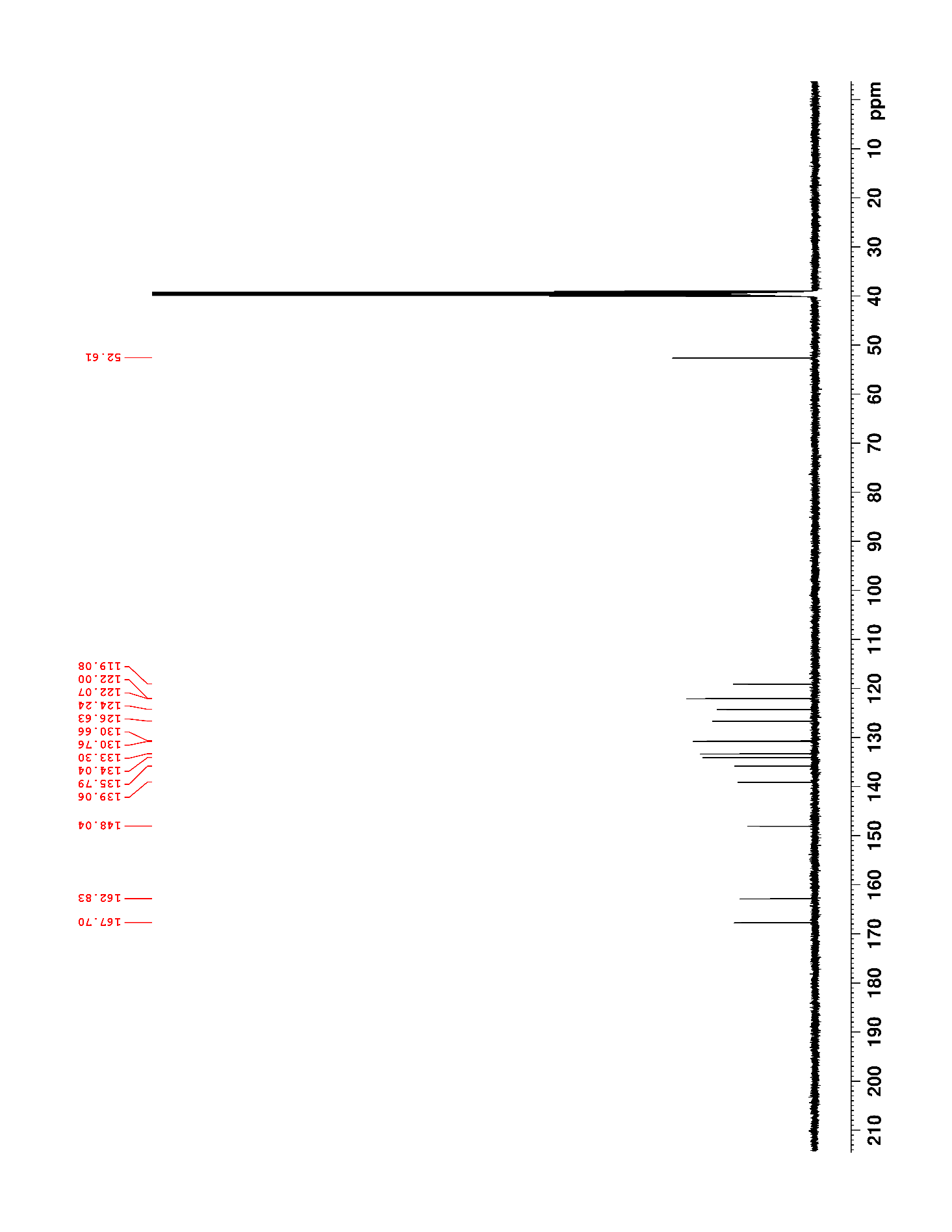

### Supplementary Figure 75. ^1^H NMR (500 MHz, DMSO-*d*_6_) of S10

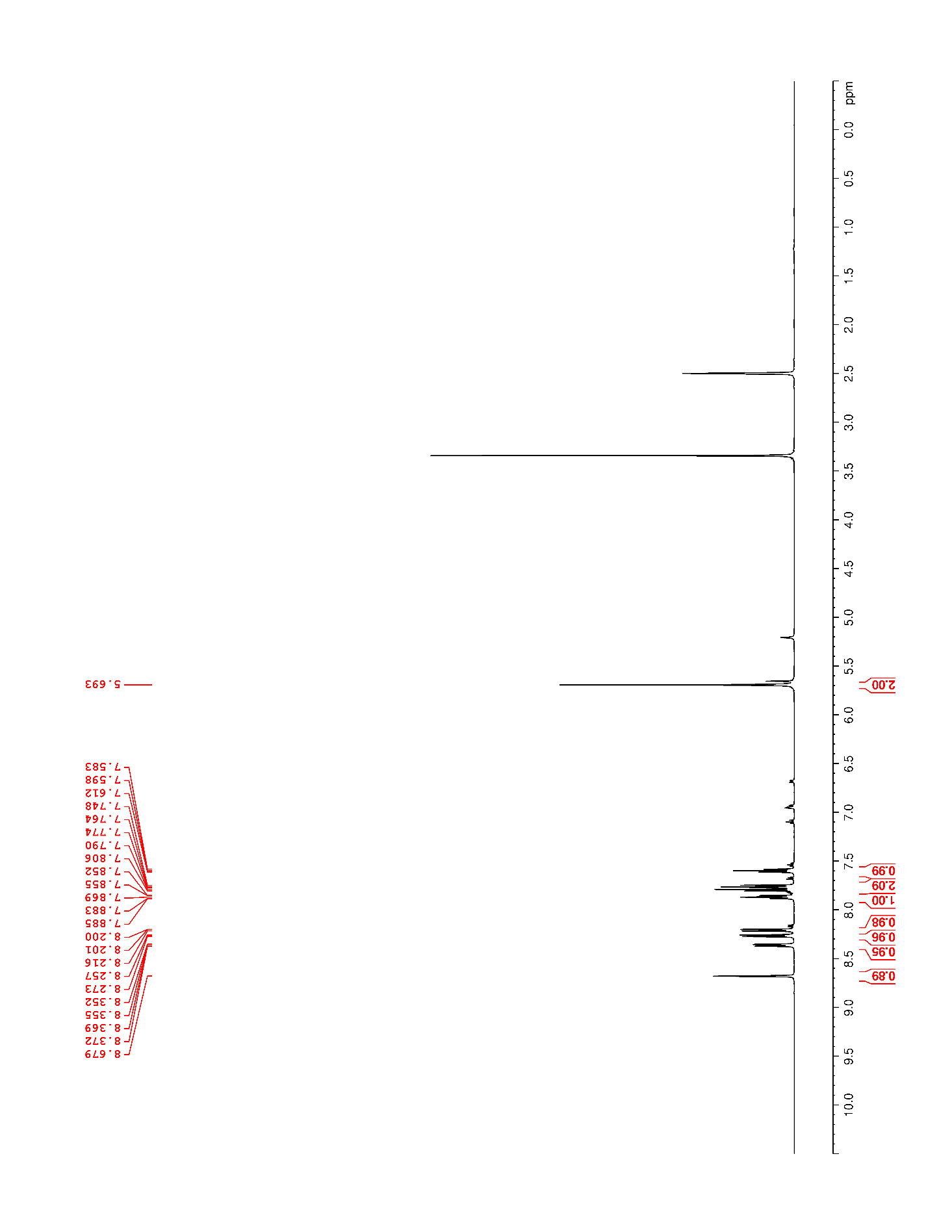

### Supplementary Figure 76. ^13^C NMR (125 MHz, DMSO-*d*_6_) of 28

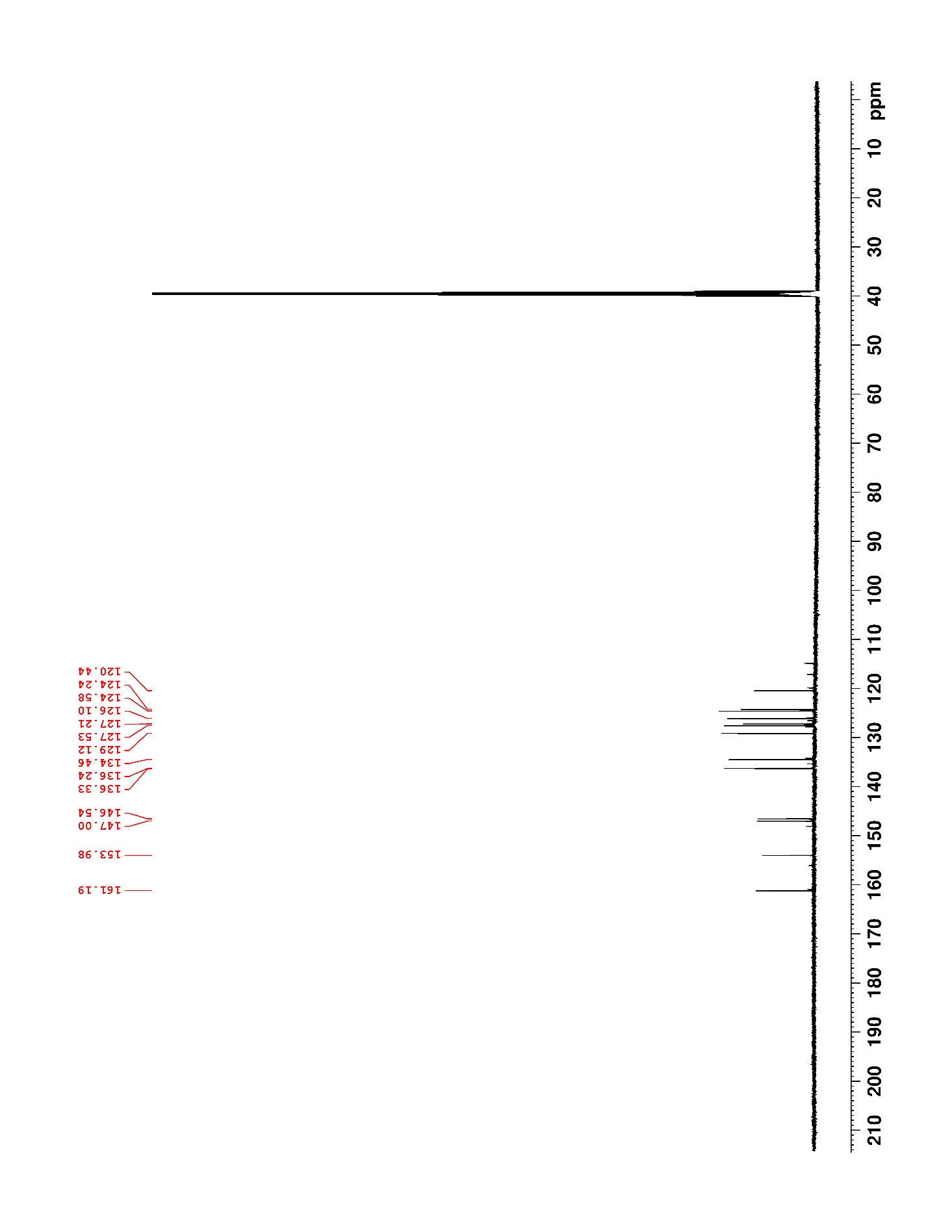

### Supplementary Figure 77. ^1^H NMR (500 MHz, DMSO-*d*_6_) of 28

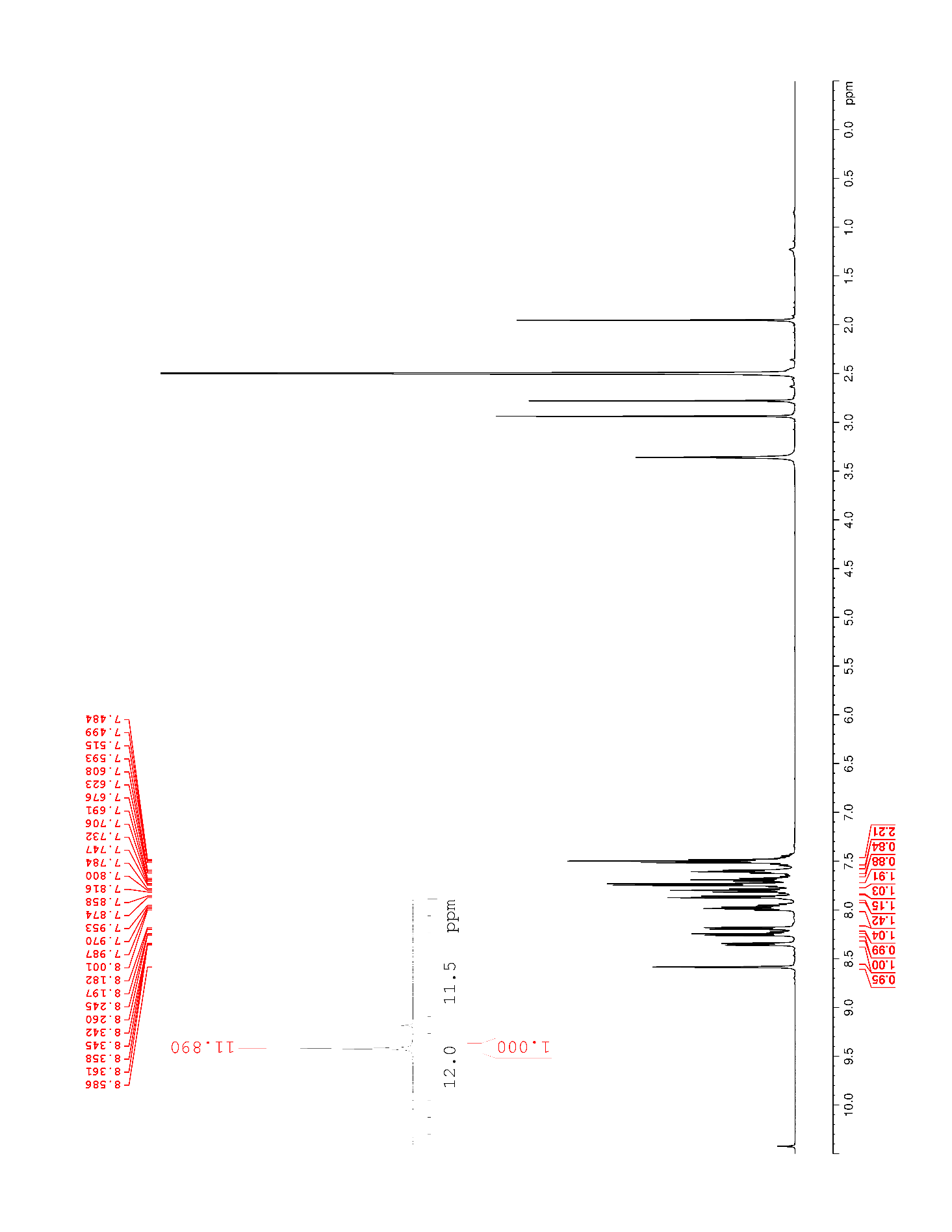

### Supplementary Figure 78. ^13^C NMR (125 MHz, DMSO-*d*_6_) of 28

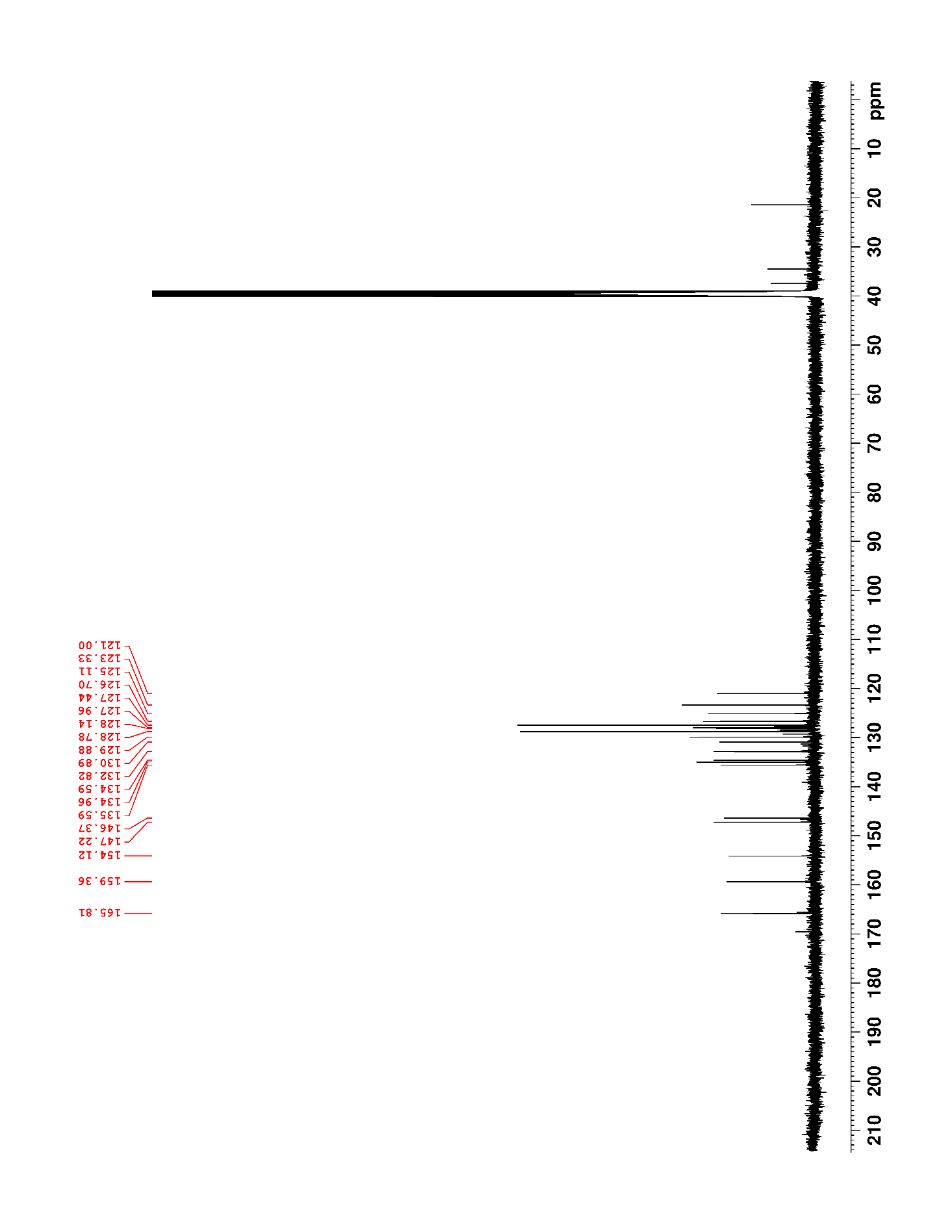

### Supplementary Figure 79. ^1^H NMR (500 MHz, DMSO-*d*_6_) of S11

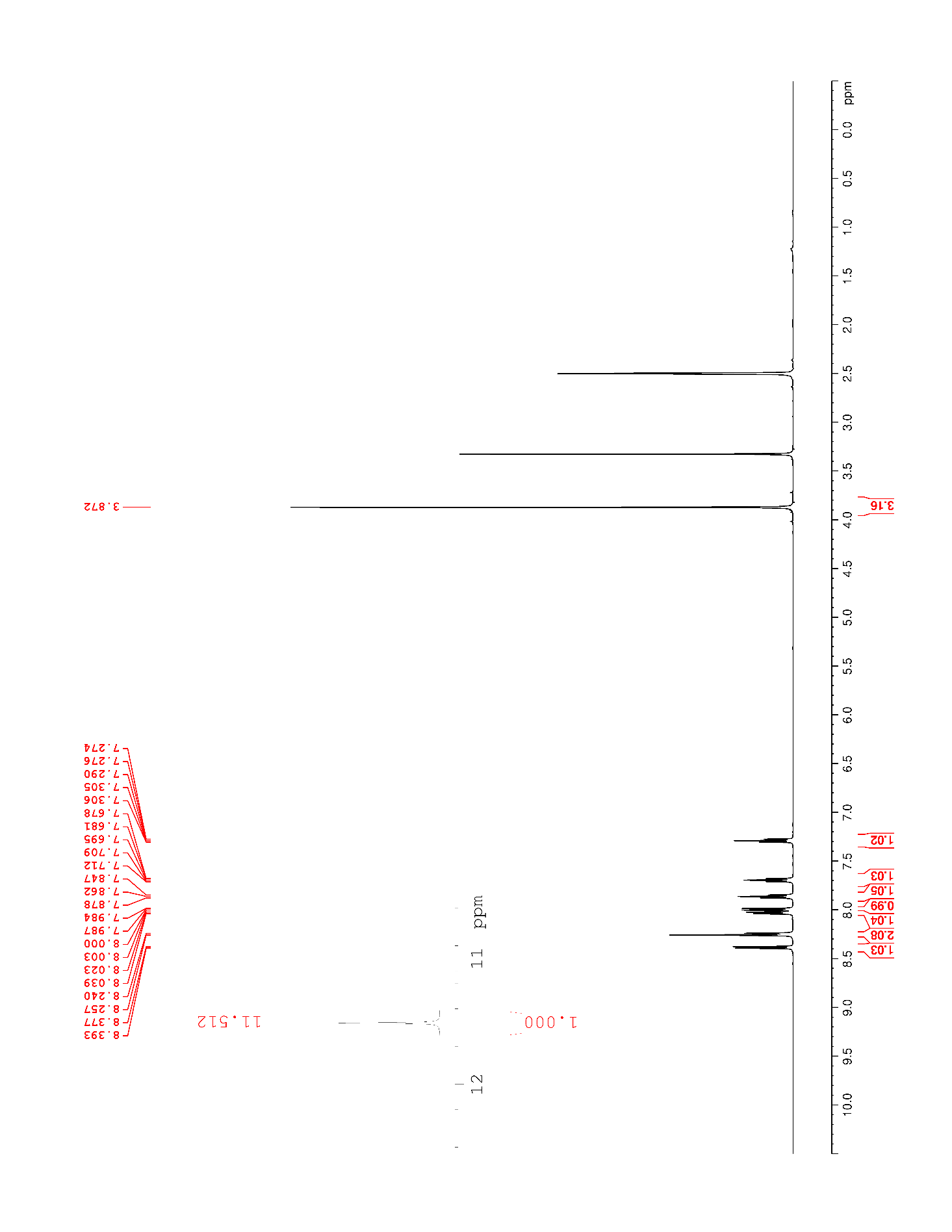

### Supplementary Figure 80. ^13^C NMR (125 MHz, DMSO-*d*_6_) of S11

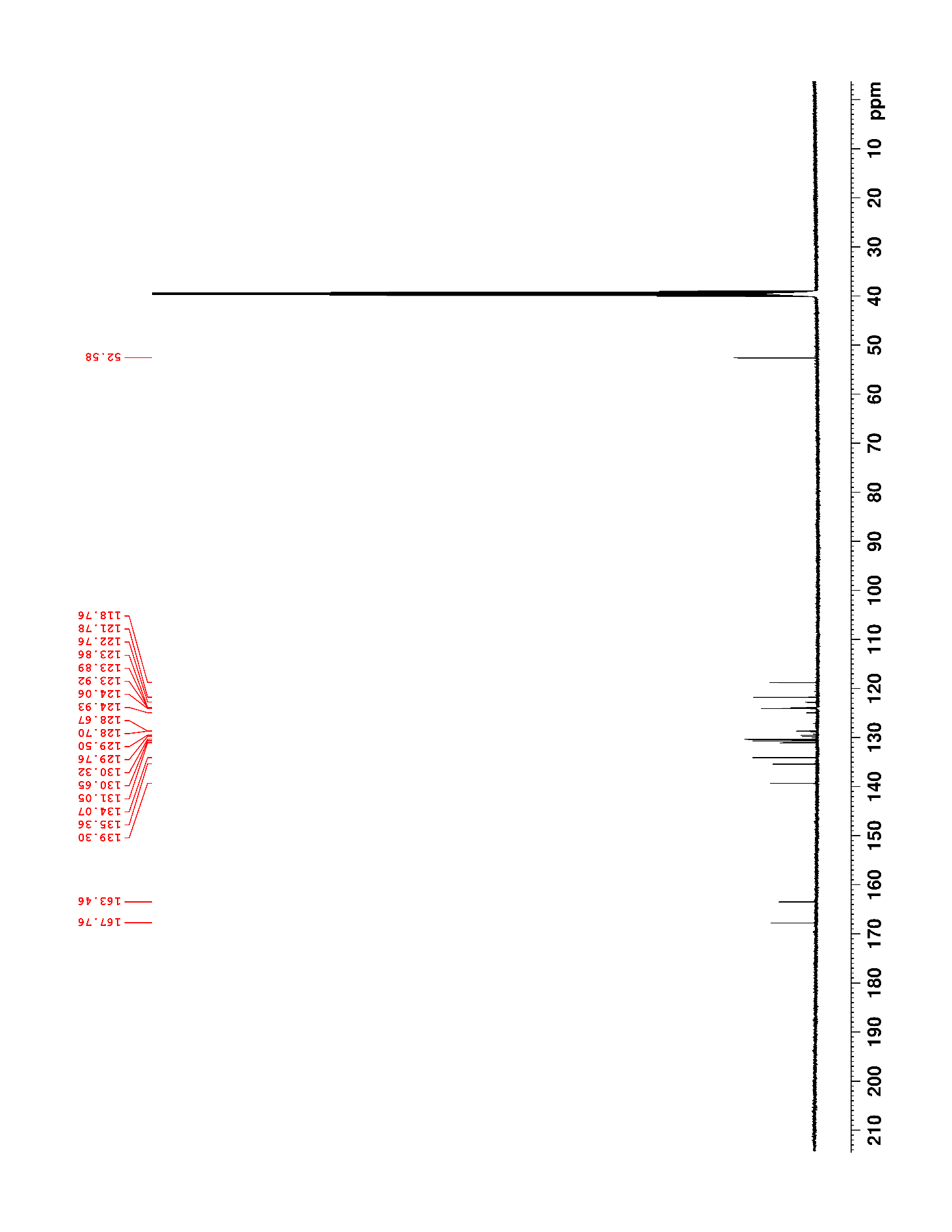

### Supplementary Figure 81. ^19^F NMR (470 MHz, DMSO-*d*_6_) of S11

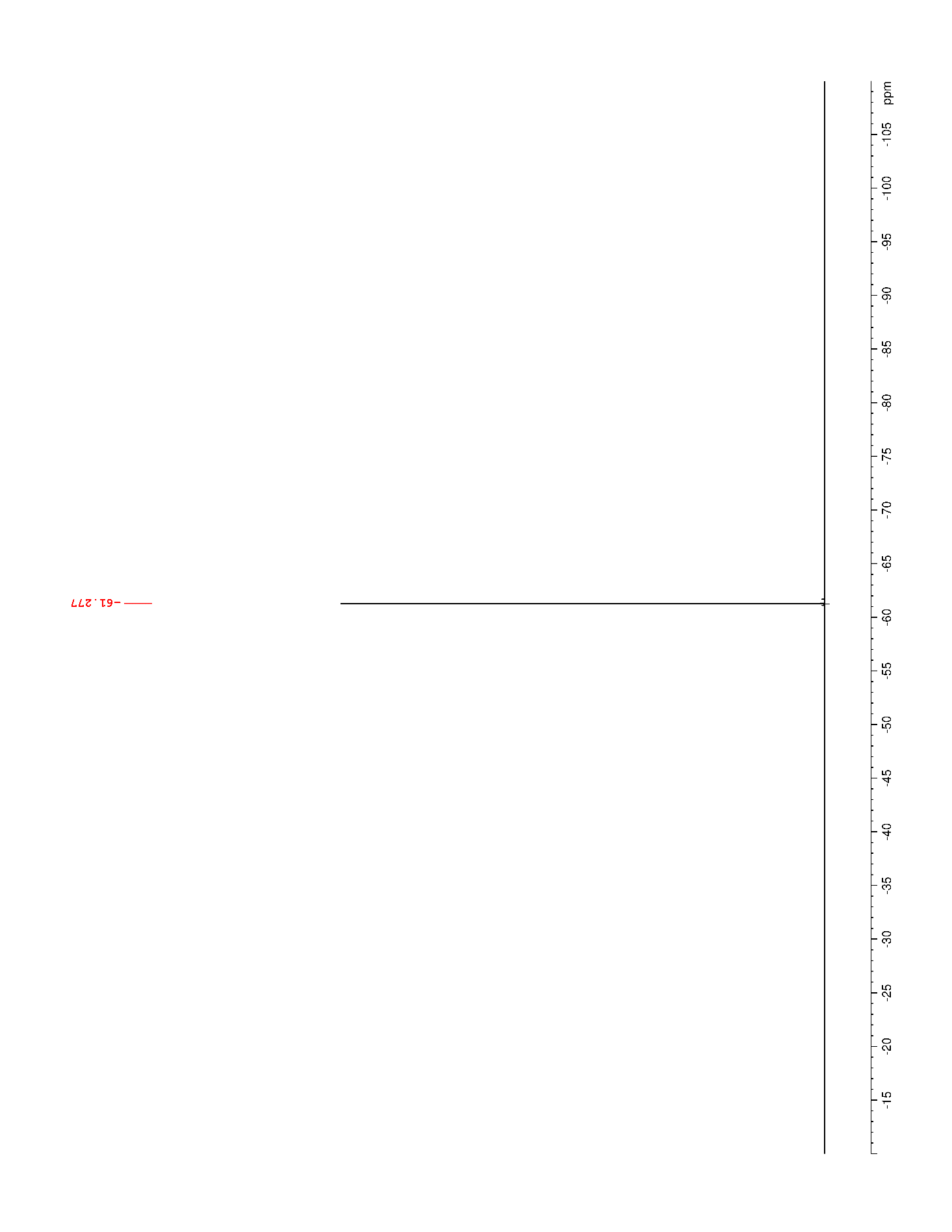

### Supplementary Figure 82. ^1^H NMR (500 MHz, DMSO-*d*_6_) of S12

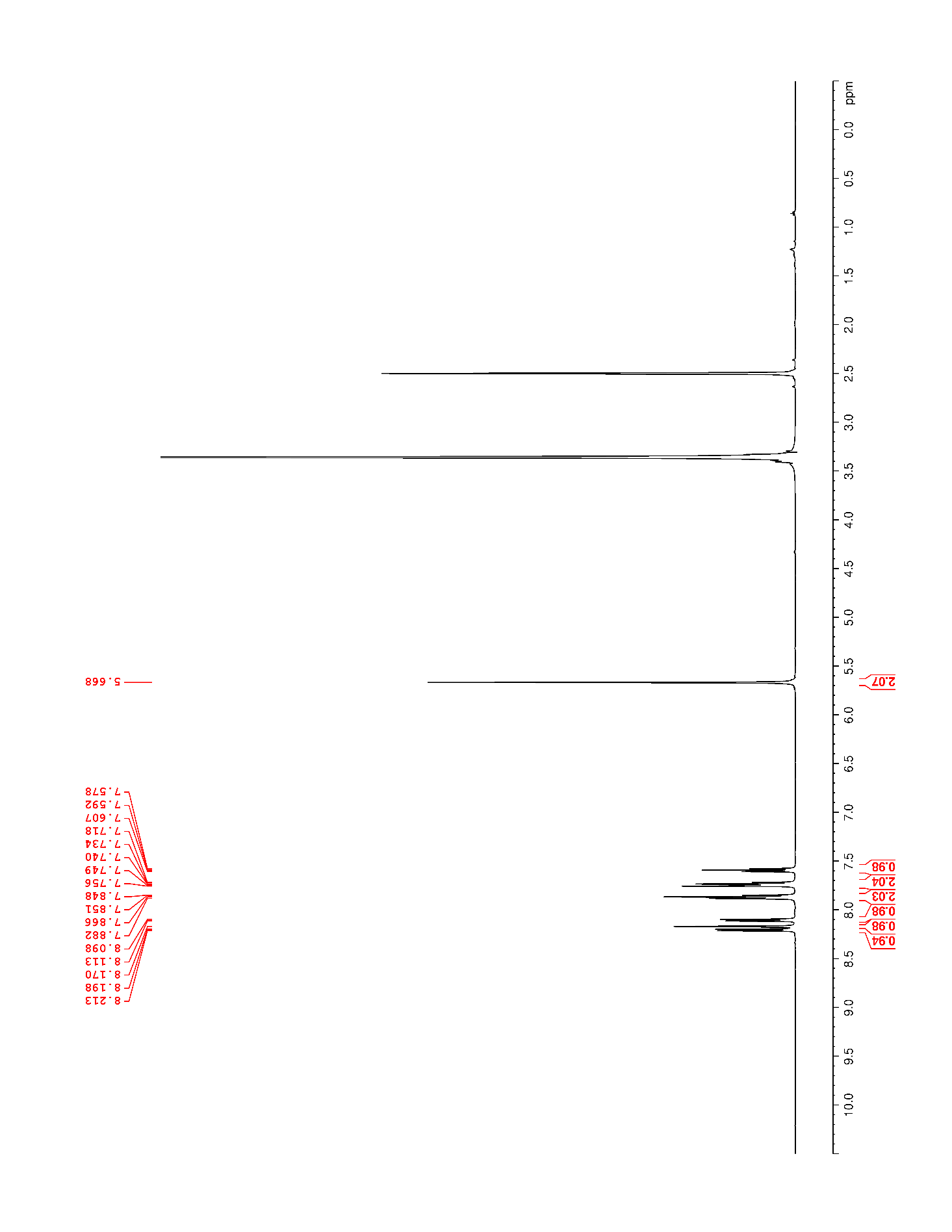

### Supplementary Figure 83. ^13^C NMR (125 MHz, DMSO-*d*_6_) of S12

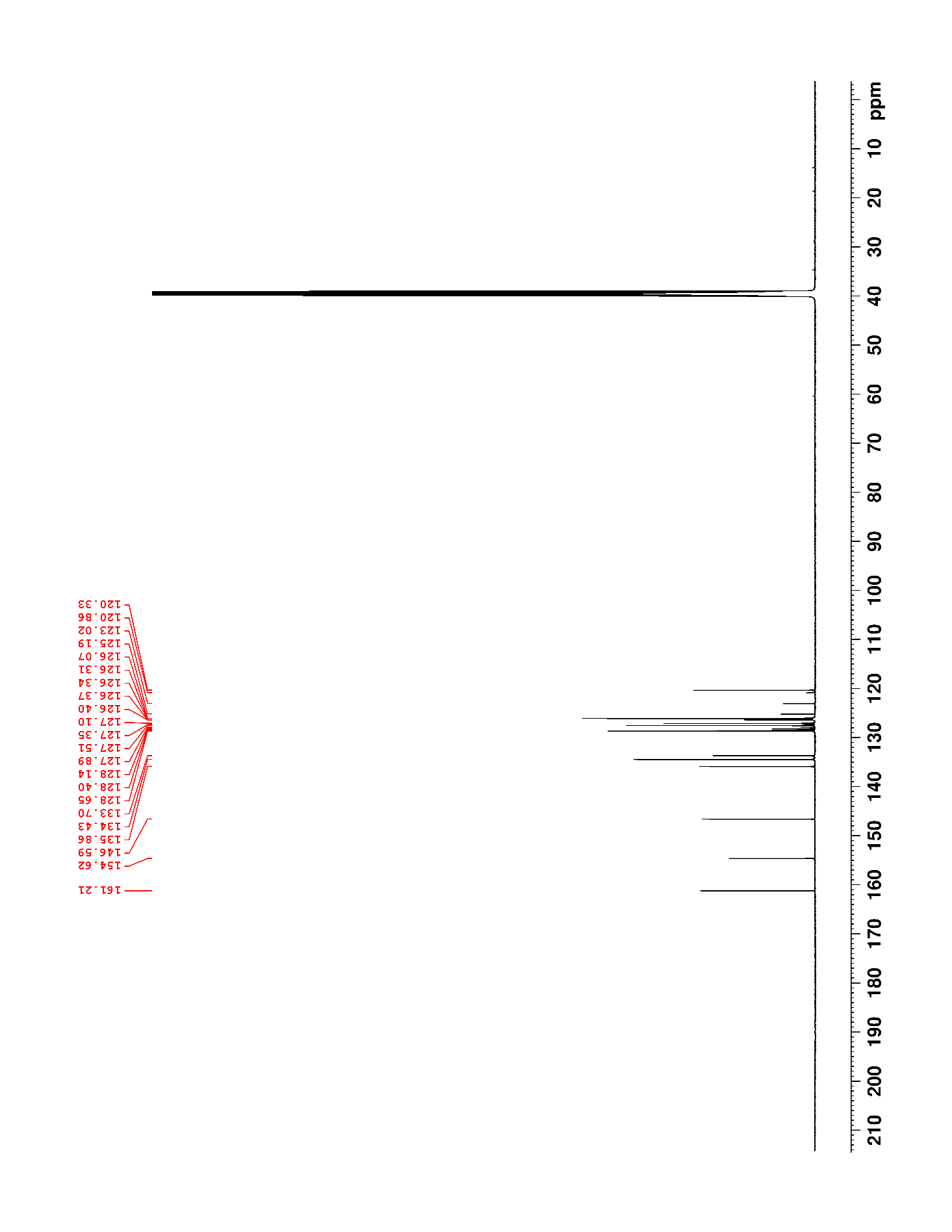

### Supplementary Figure 84. ^19^F NMR (470 MHz, DMSO-*d*_6_) of S12

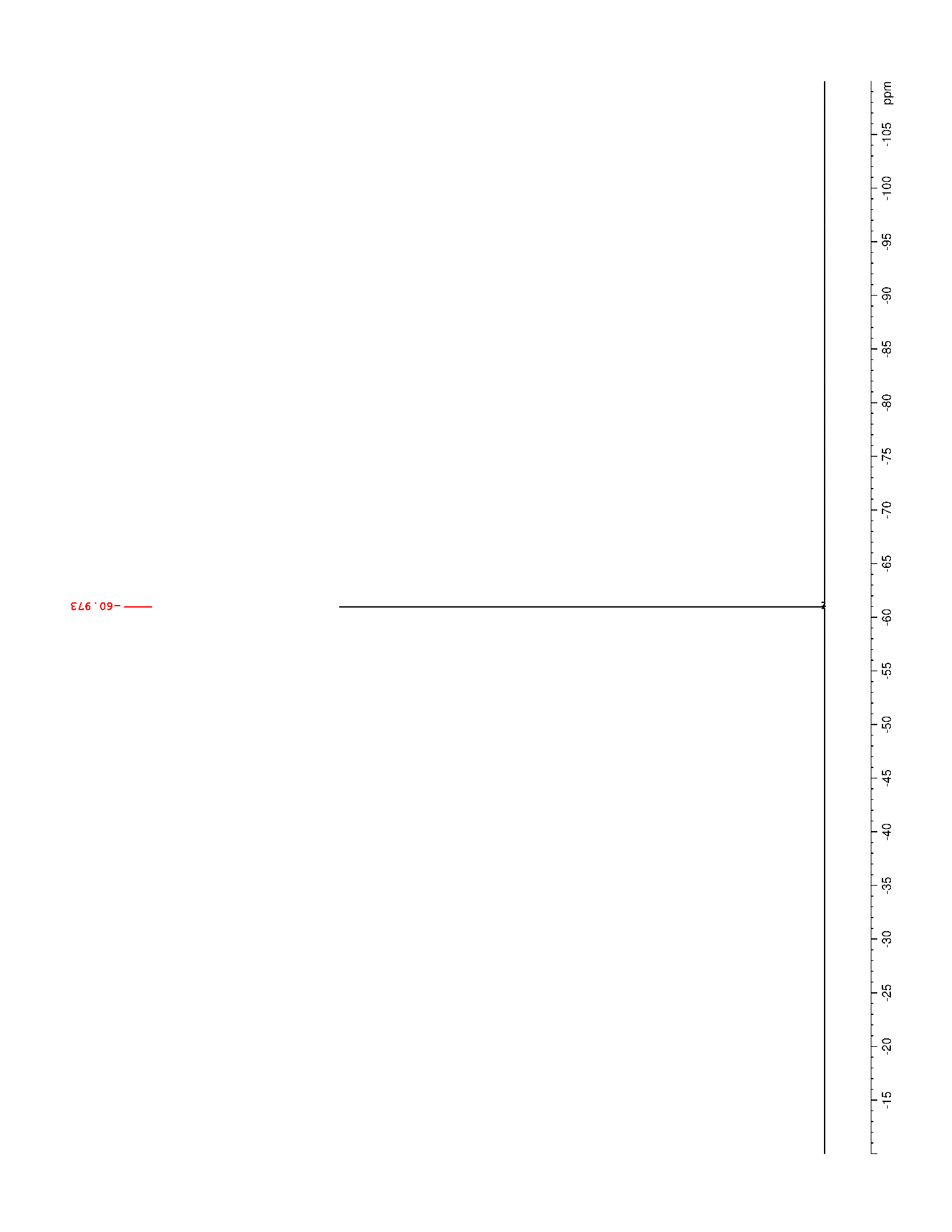

### Supplementary Figure 85. ^1^H NMR (500 MHz, CDCl_3_) of 29

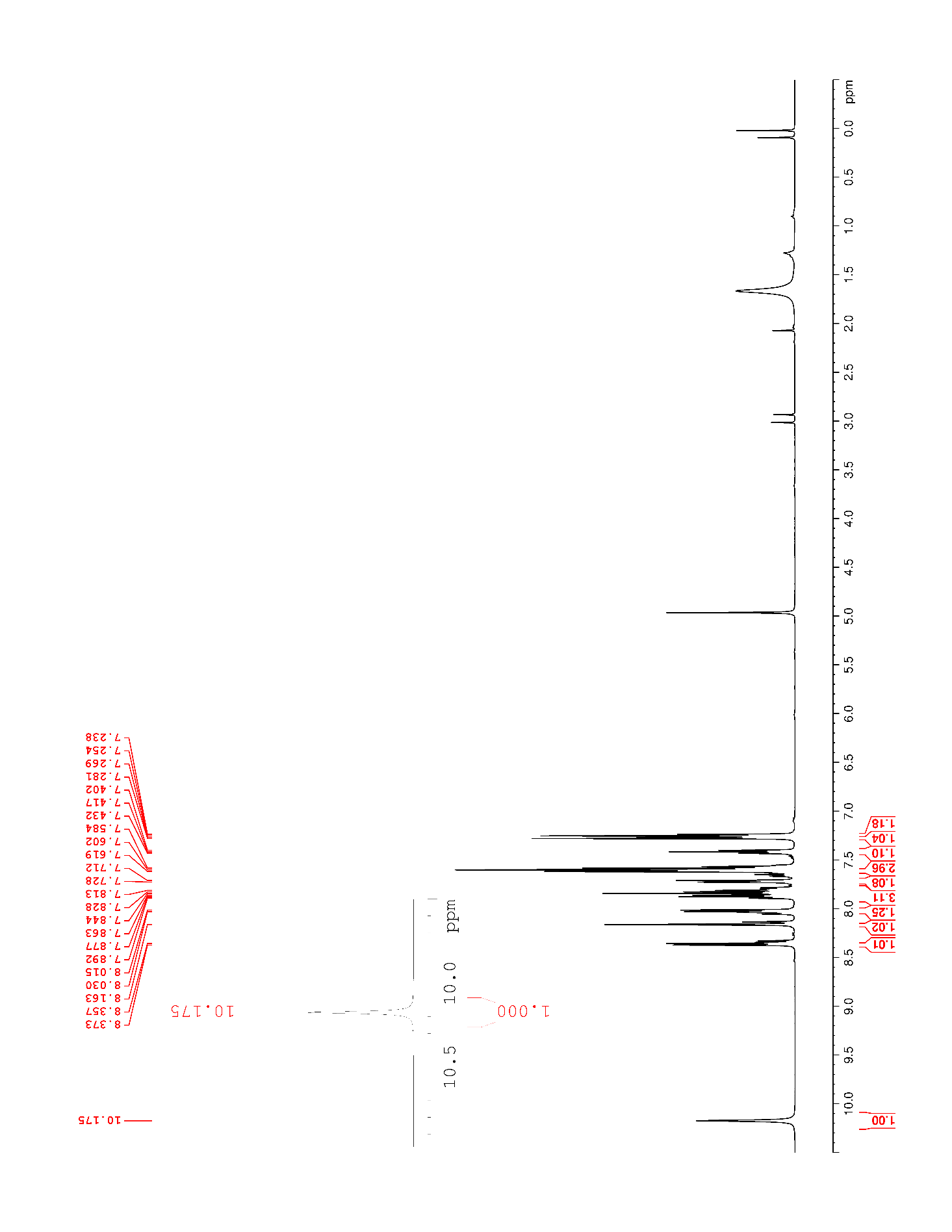

### Supplementary Figure 86. ^13^C NMR (125 MHz, CDCl_3_) of 29

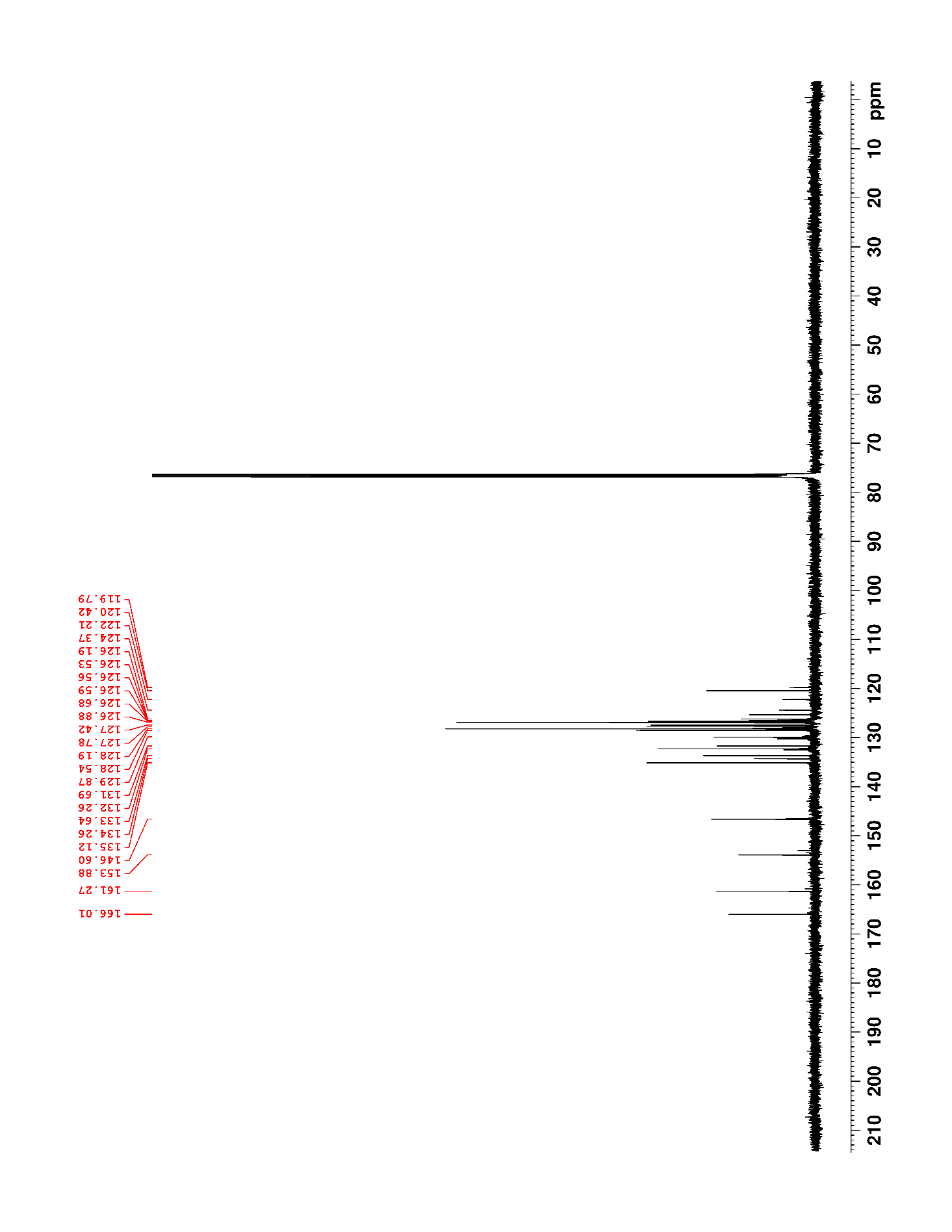

### Supplementary Figure 87. ^19^F NMR (470 MHz, CDCl_3_) of 29

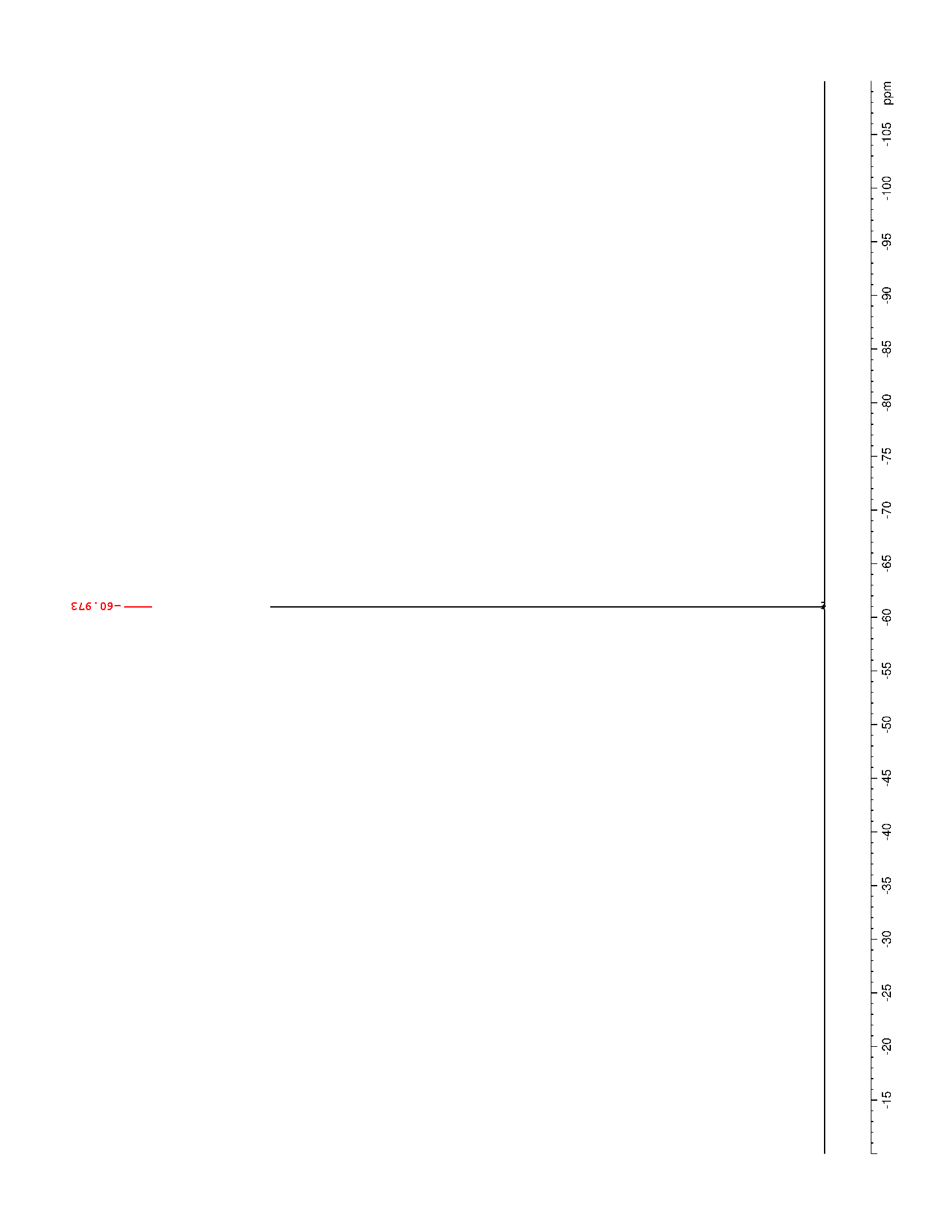

### Supplementary Figure 88. ^1^H NMR (500 MHz, DMSO-*d*_6_) of 30

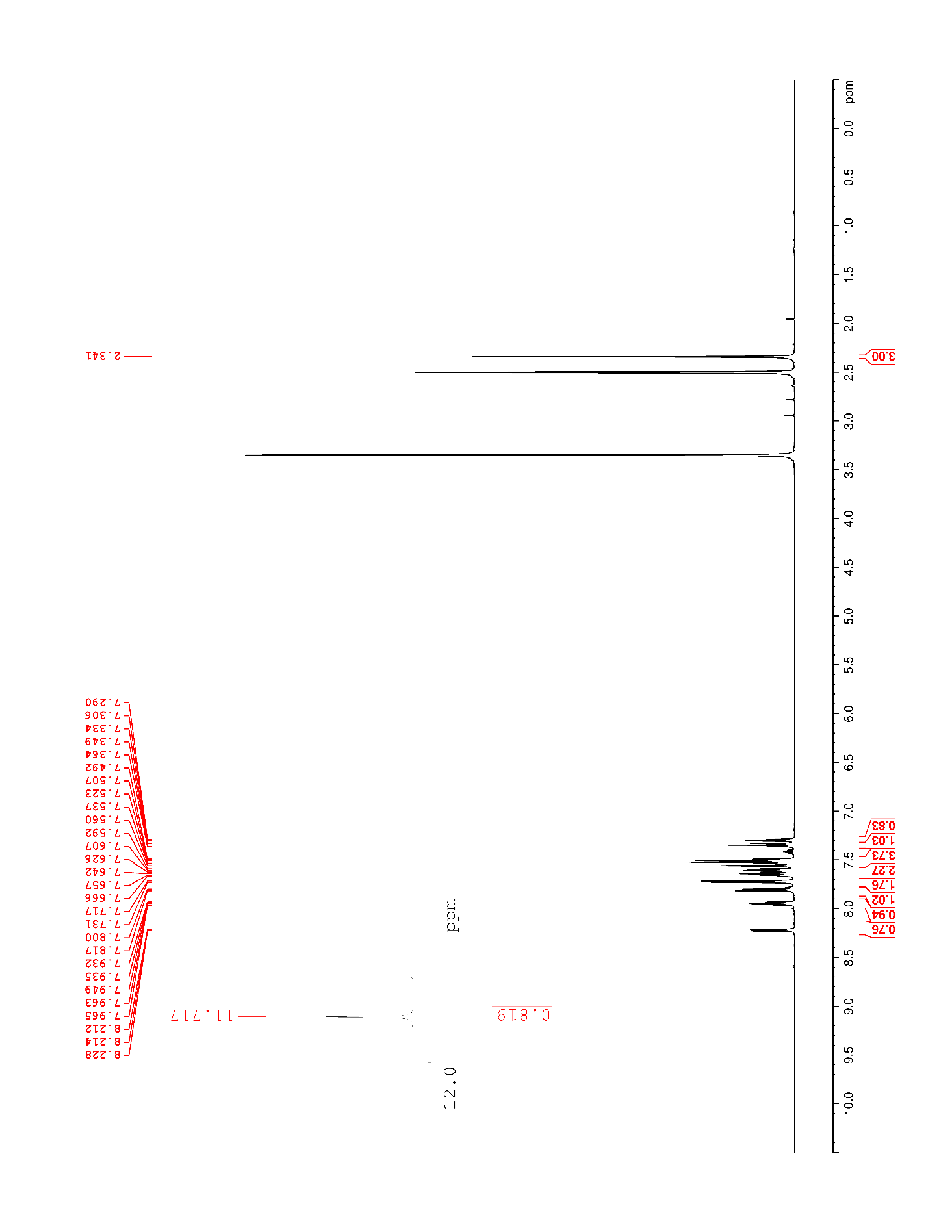

### Supplementary Figure 89. ^13^C NMR (125 MHz, DMSO-*d*_6_) of 30

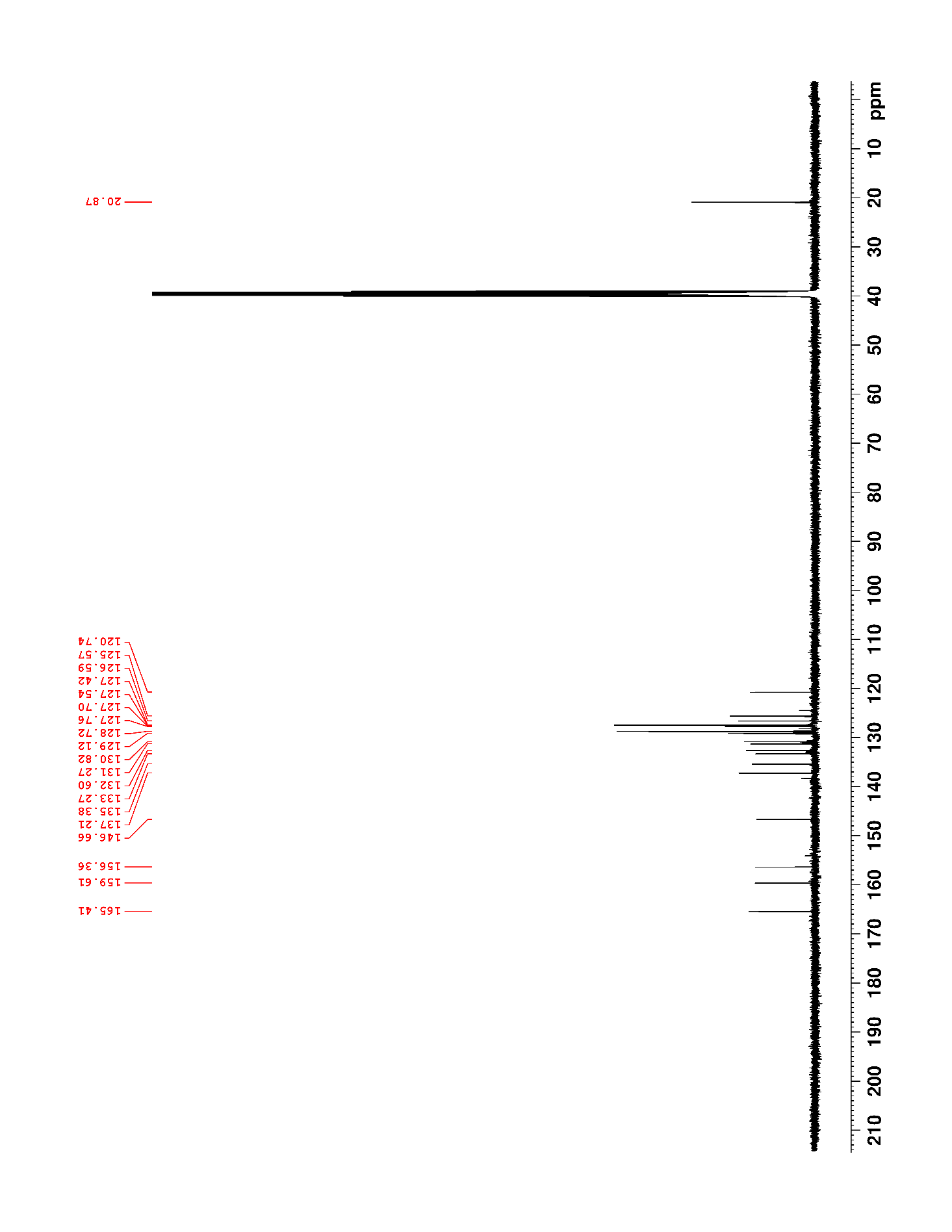

### Supplementary Figure 90. ^1^H NMR (500 MHz, CDCl_3_) of S15

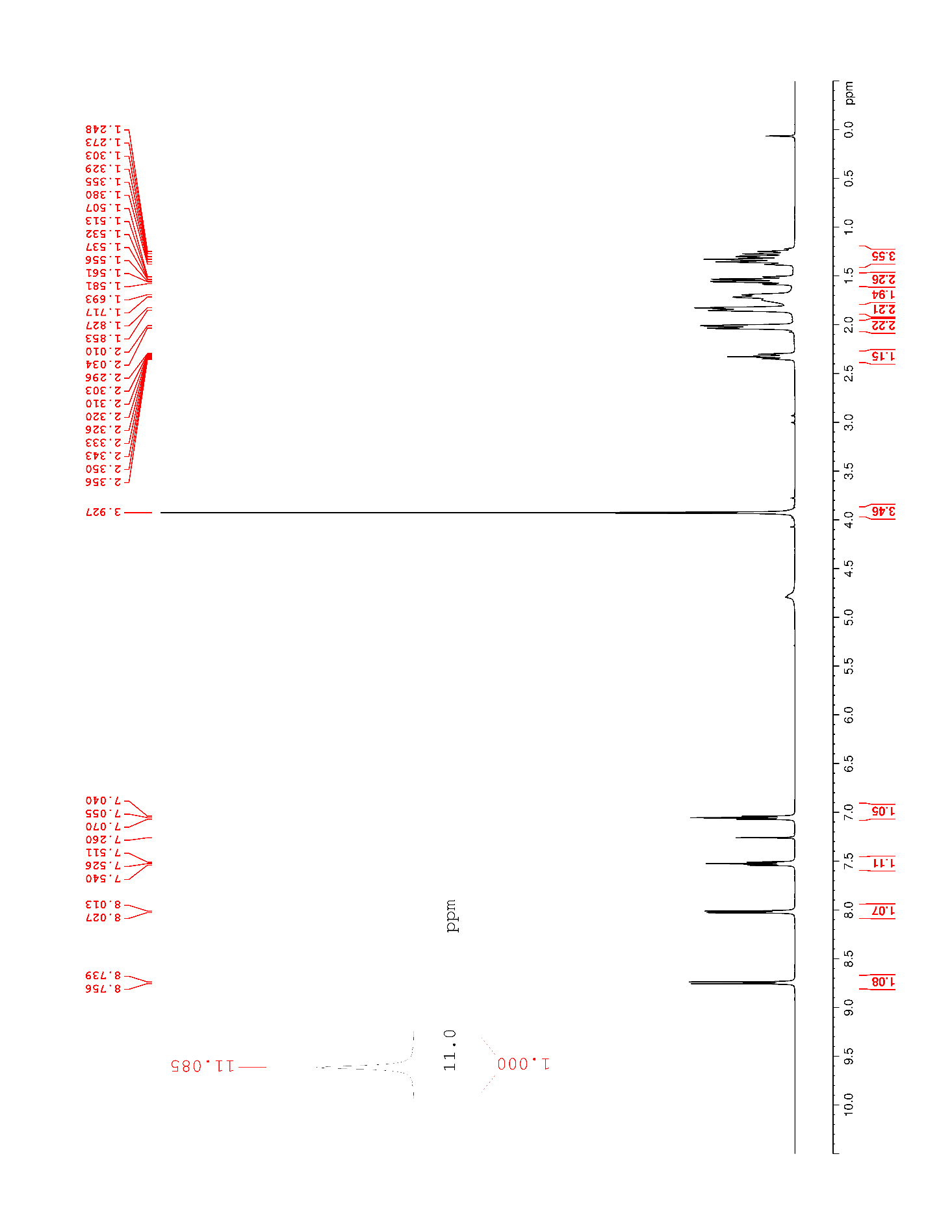

### Supplementary Figure 91. ^13^C NMR (125 MHz, CDCl_3_) of S15

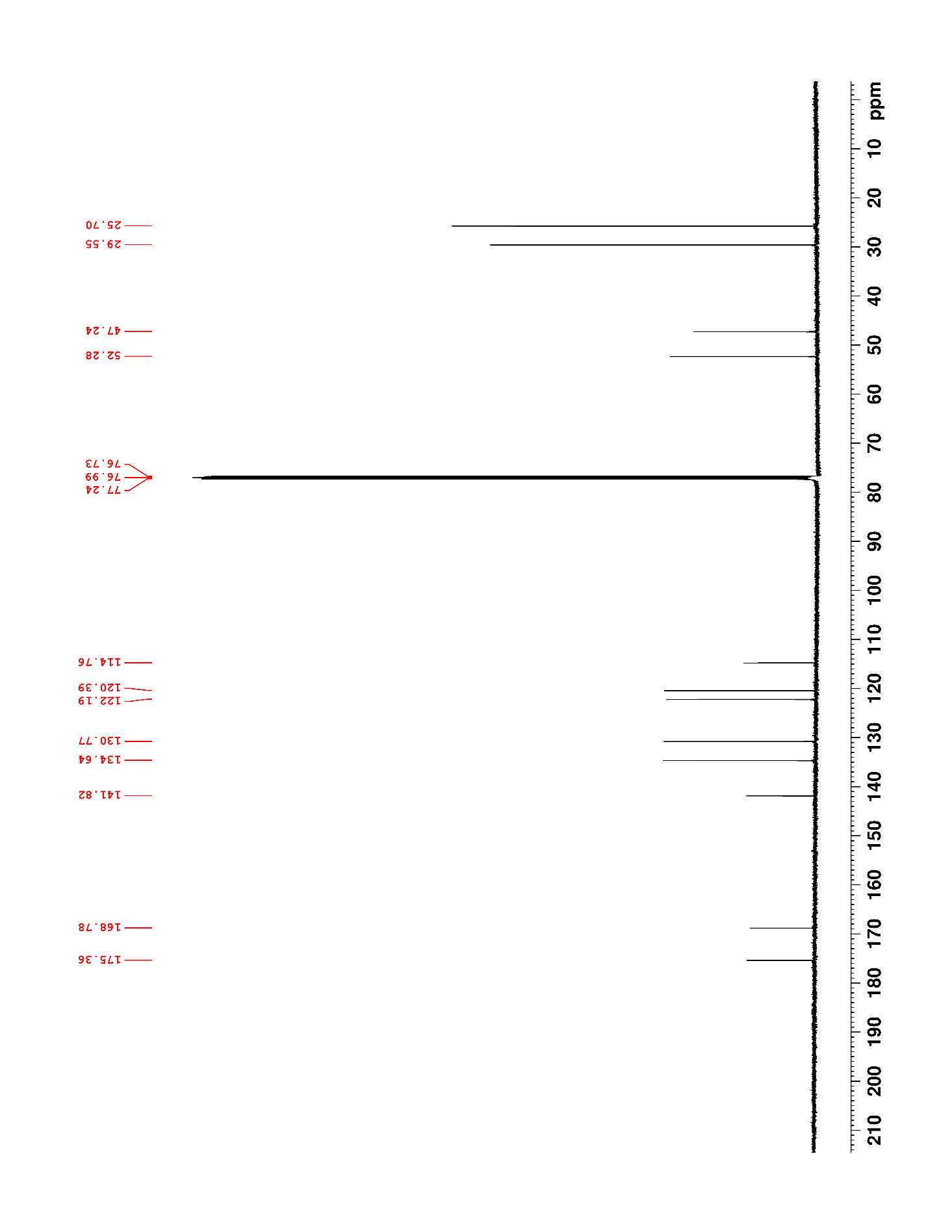

### Supplementary Figure 92. ^1^H NMR (500 MHz, DMSO-*d*_6_) of S16

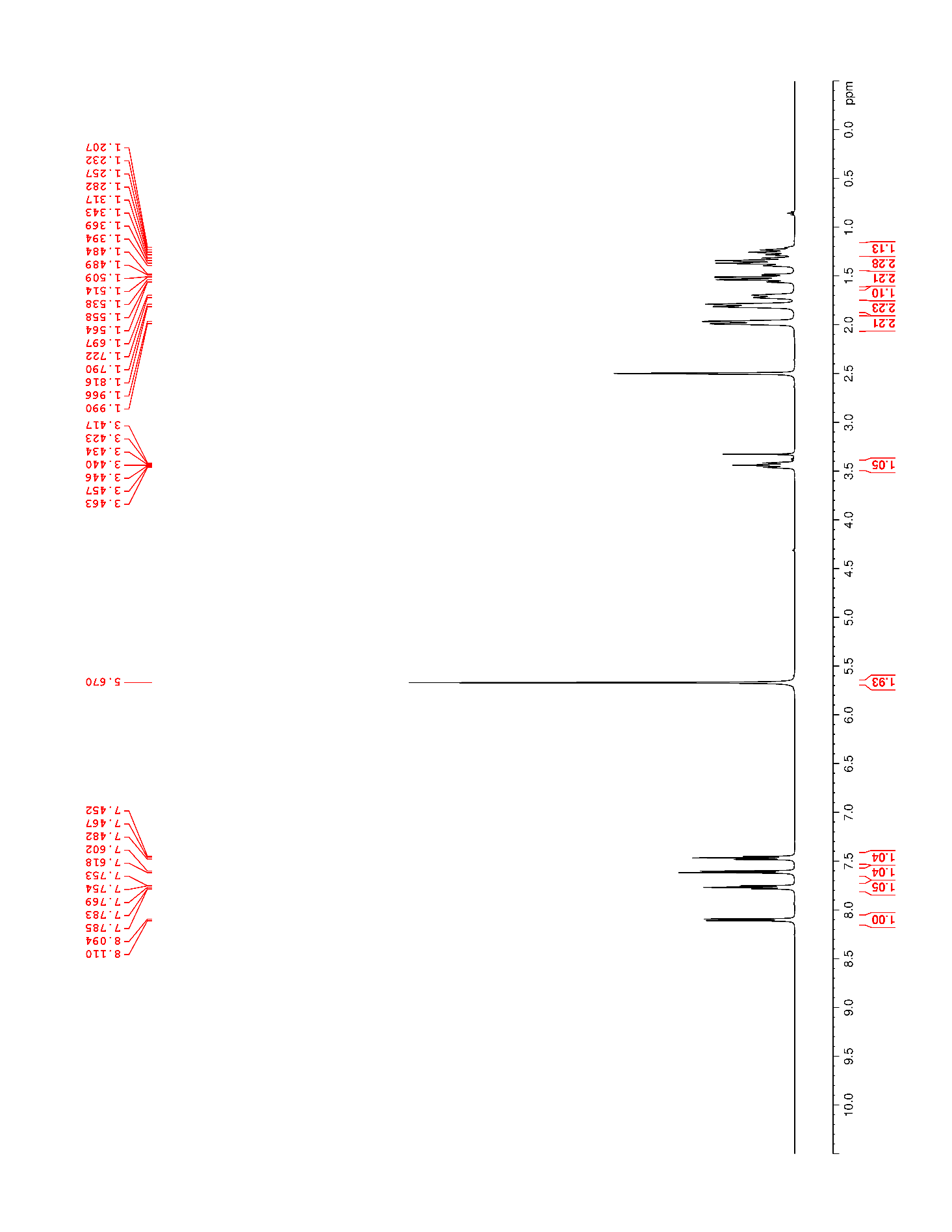

### Supplementary Figure 93. ^13^C NMR (125 MHz, DMSO-*d*_6_) of S16

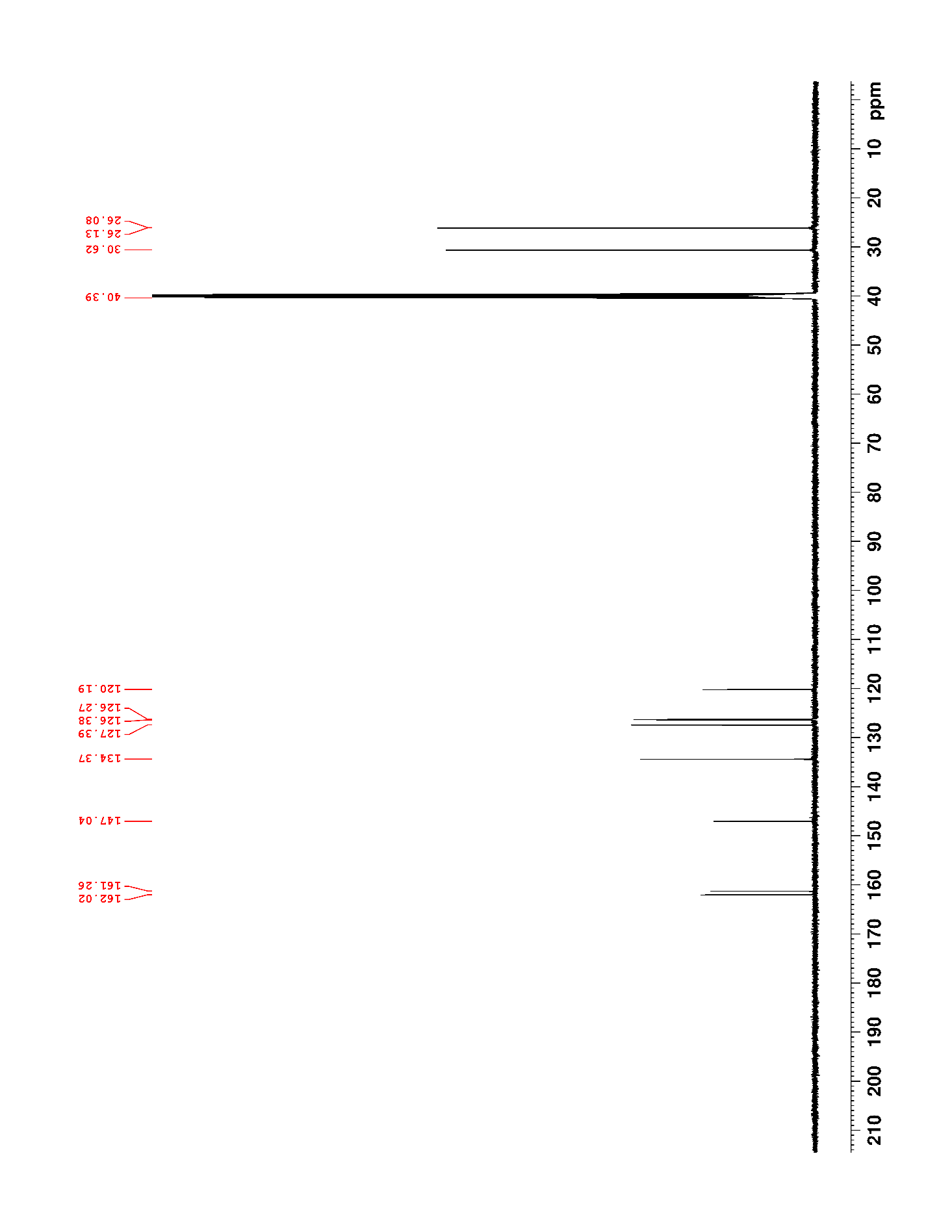

### Supplementary Figure 94. ^1^H NMR (500 MHz, DMSO-*d*_6_) of 31

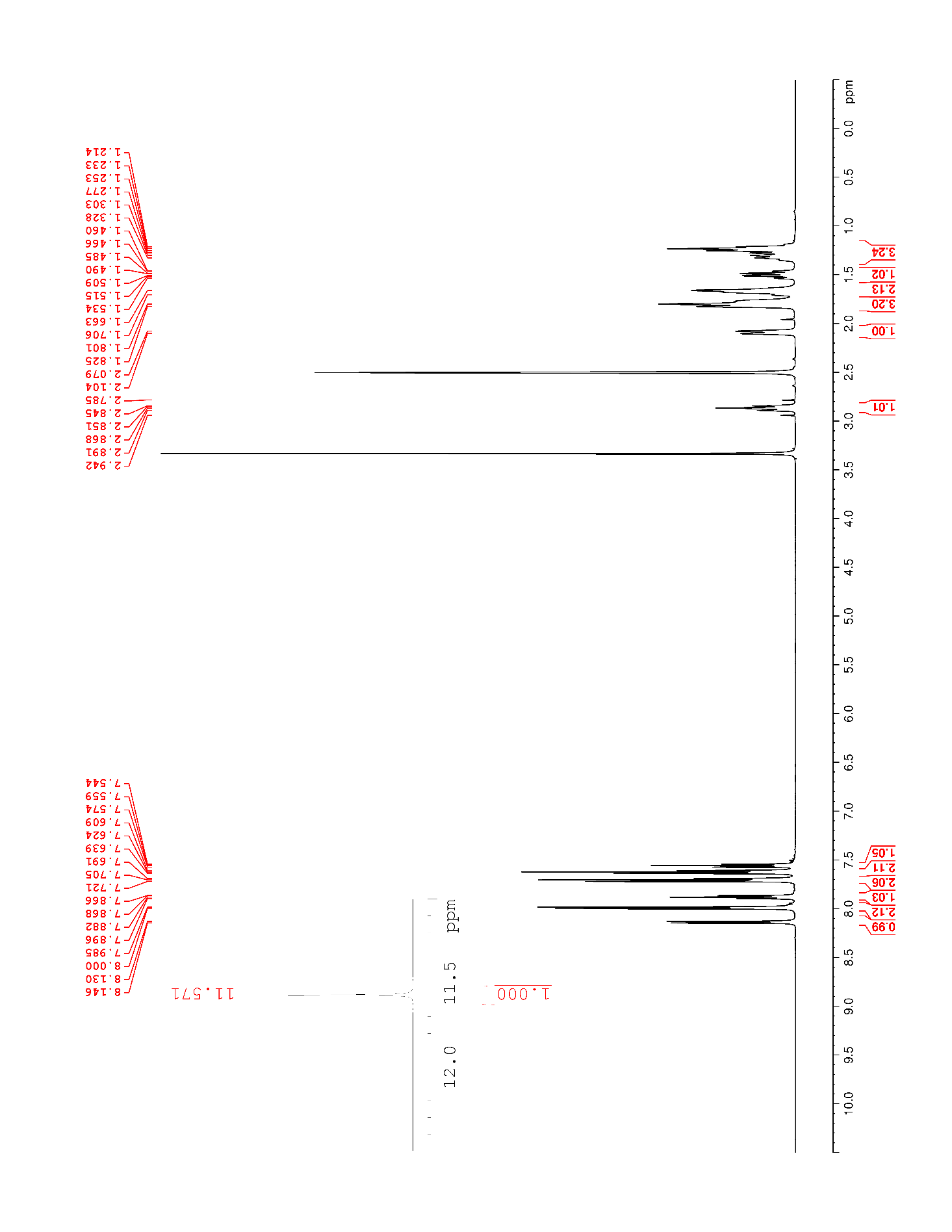

### Supplementary Figure 95. ^13^C NMR (125 MHz, DMSO-*d*_6_) of 31

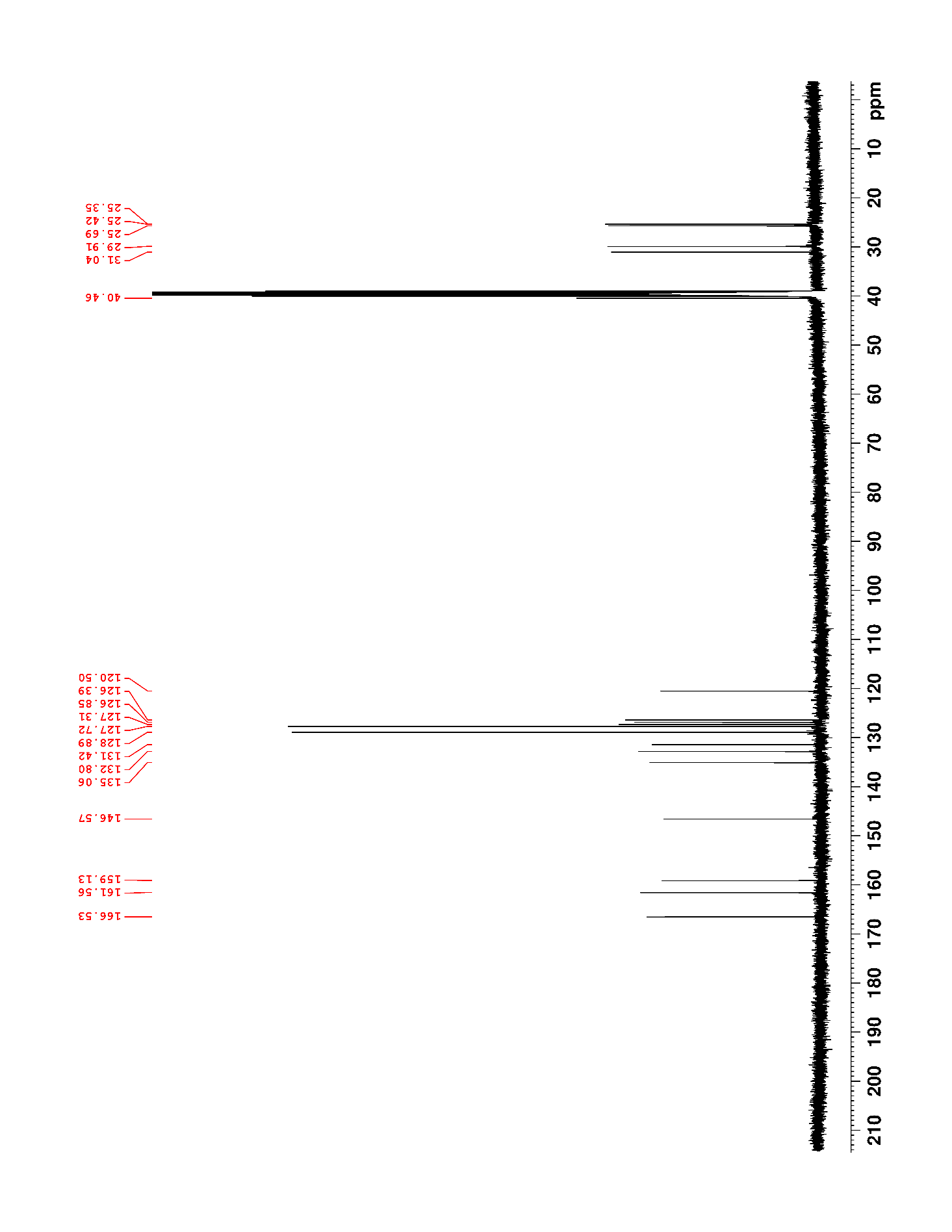

### Supplementary Figure 96. ^1^H NMR (500 MHz, DMSO-*d*_6_) of 32

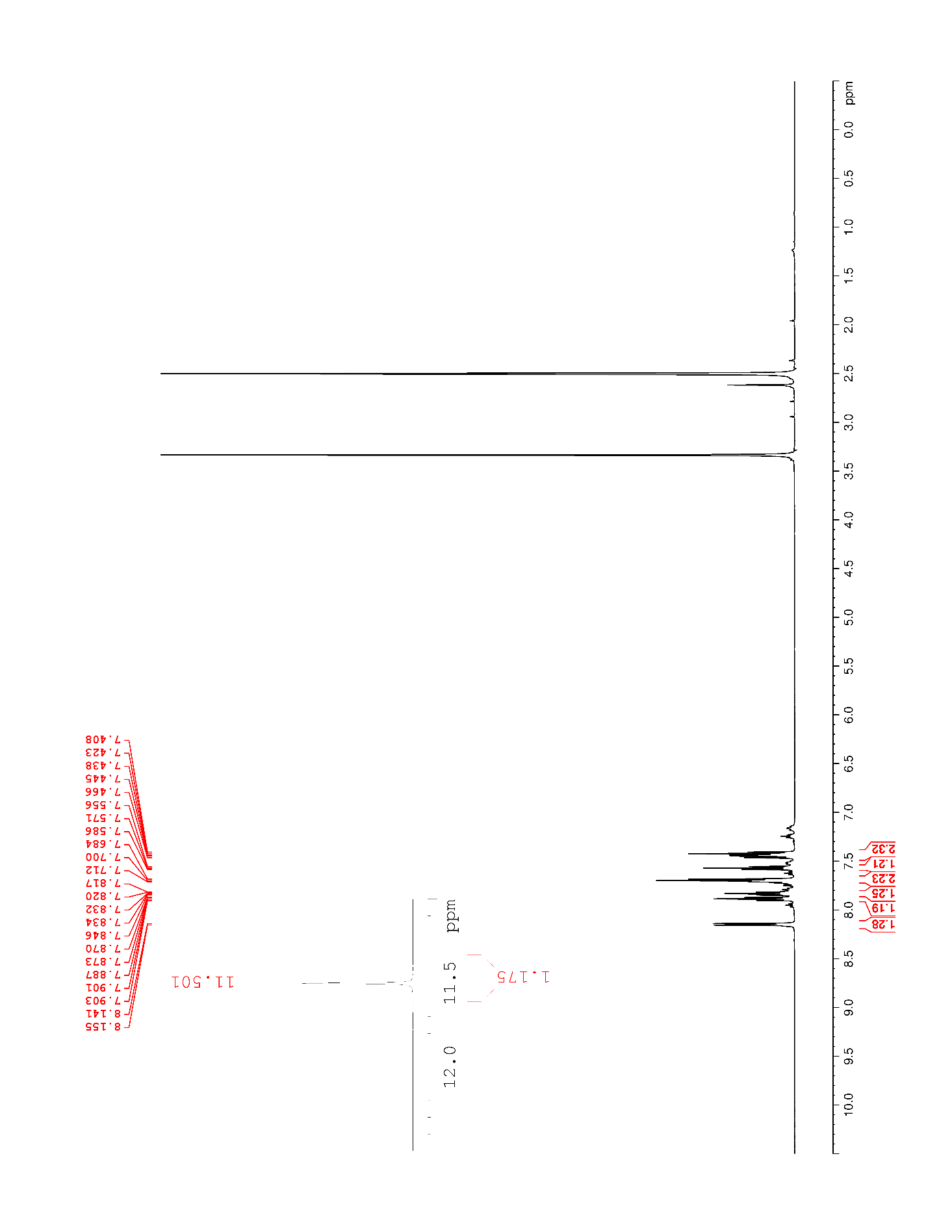

### Supplementary Figure 97. ^13^C NMR (125 MHz, DMSO-*d*_6_) of 32

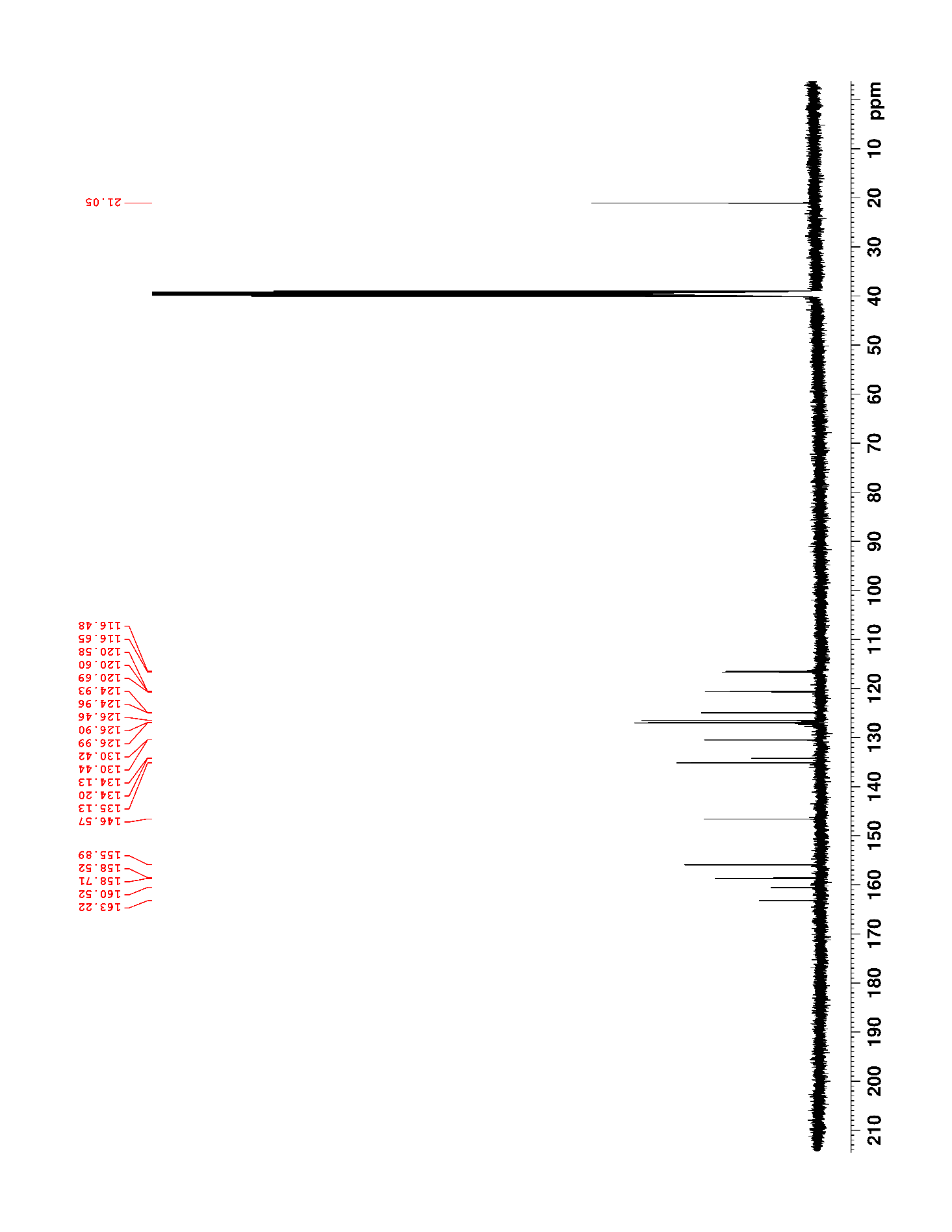

### Supplementary Figure 98. ^19^F NMR (470 MHz, DMSO-*d*_6_) of 32

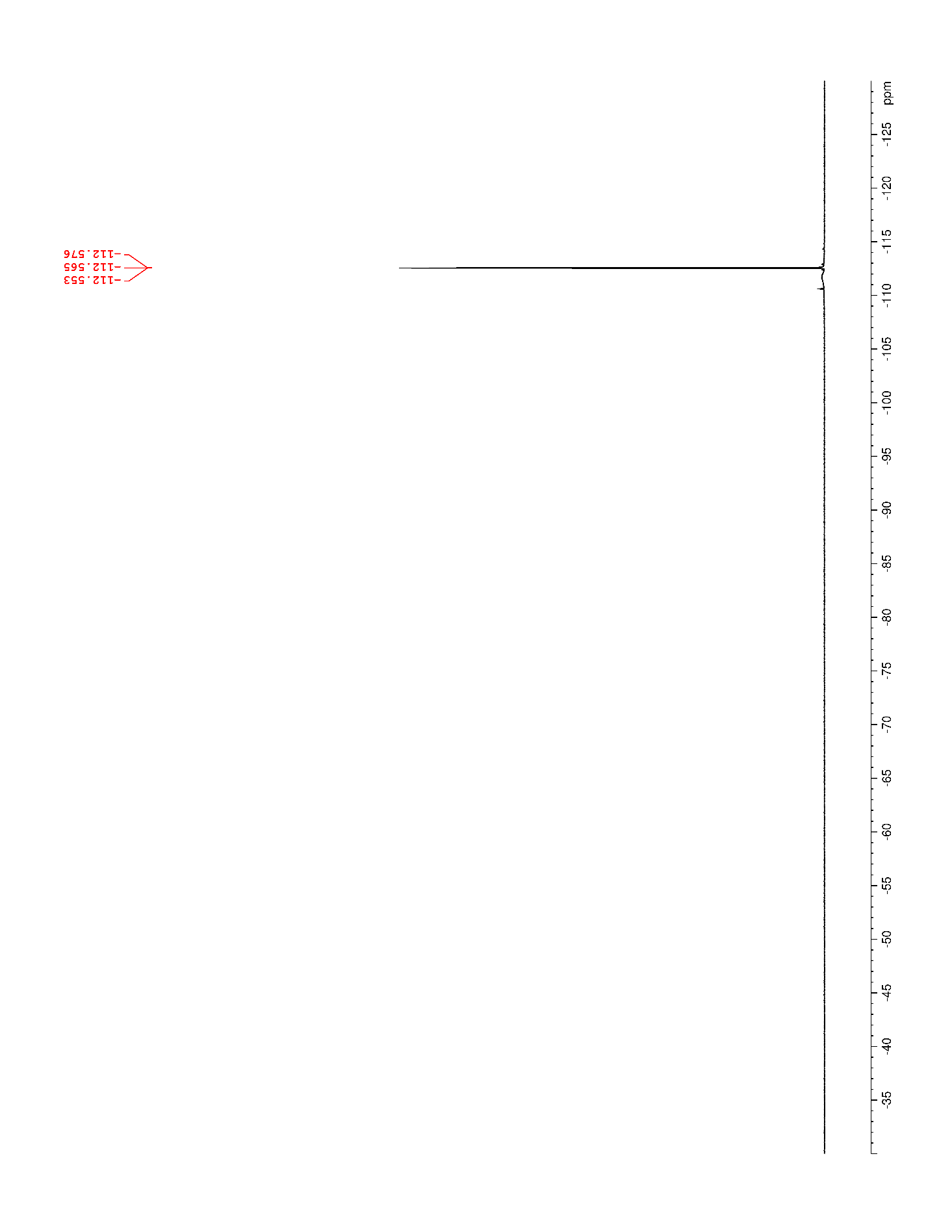

### Supplementary Figure 99. ^1^H NMR (500 MHz, DMSO-*d*_6_) of S17

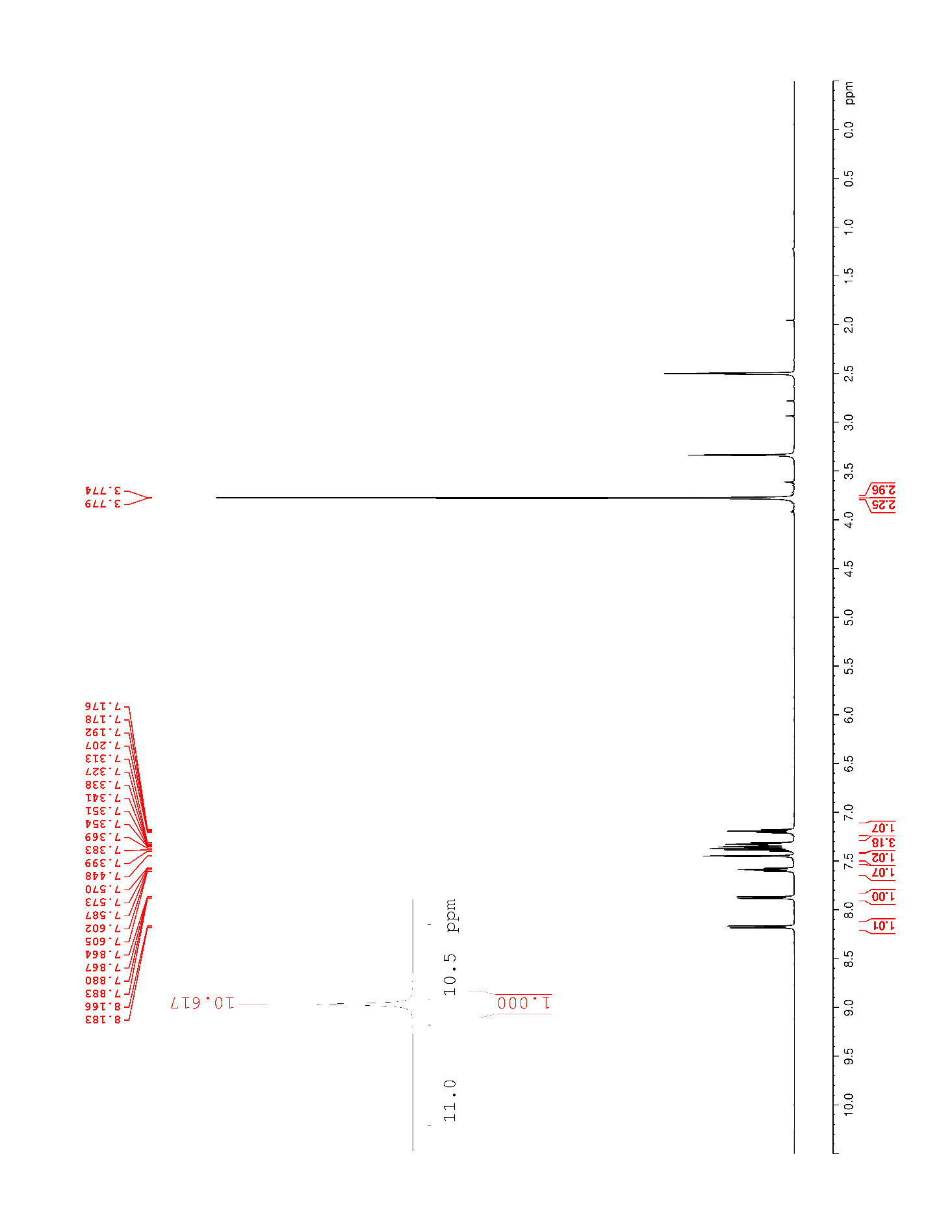

### Supplementary Figure 100. ^13^C NMR (125 MHz, DMSO-*d*_6_) of S17

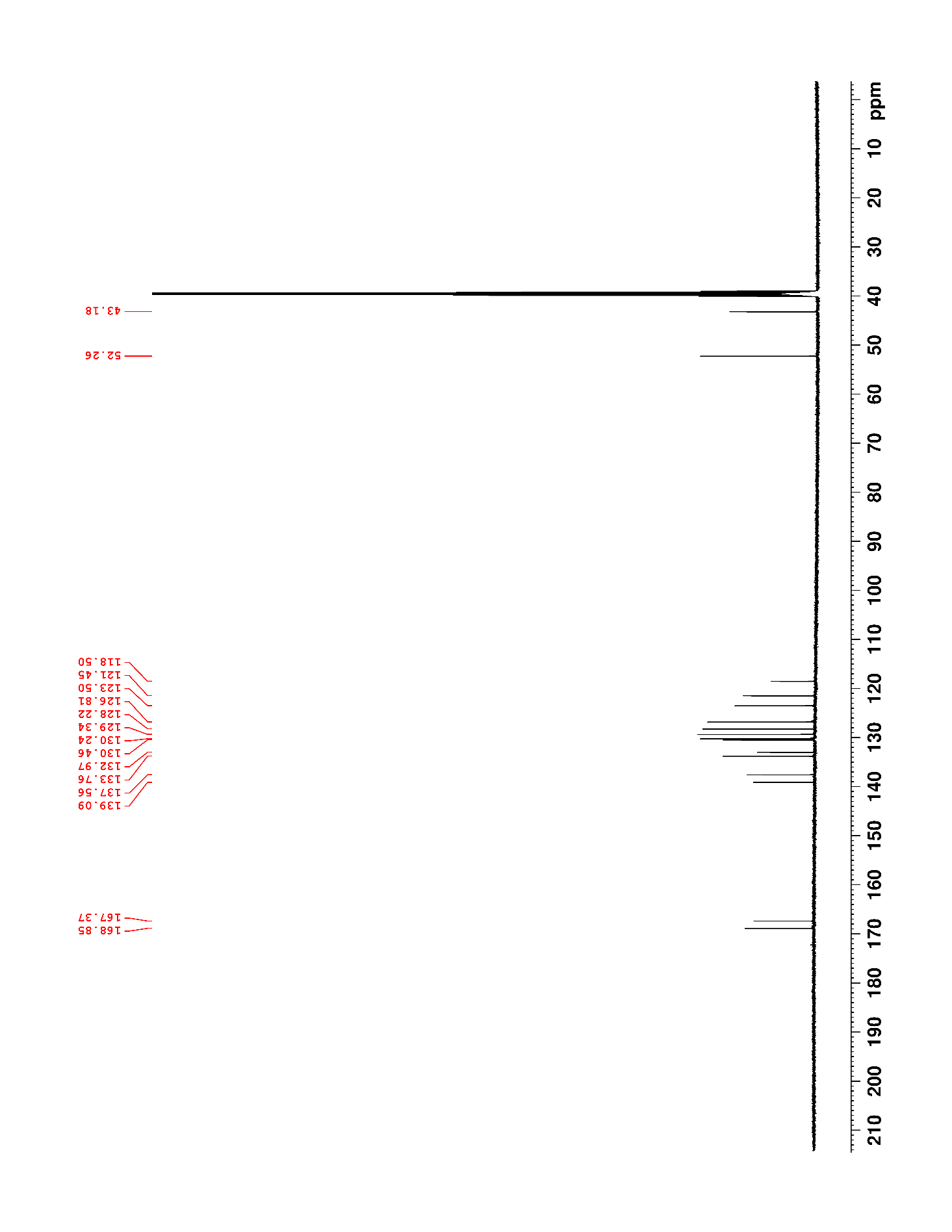

### Supplementary Figure 101. ^1^H NMR (500 MHz, DMSO-*d*_6_) of S18

### Supplementary Figure 102. ^13^C NMR/DEPT135 (125 MHz, DMSO-*d*_6_) of S18

### Supplementary Figure 103. ^1^H NMR (500 MHz, DMSO-*d*_6_) of 33

### Supplementary Figure 104. ^13^C NMR (125 MHz, DMSO-*d*_6_) of 33

### Supplementary Figure 105. ^1^H NMR (500 MHz, DMSO-*d*_6_) of S19

### Supplementary Figure 106. ^13^C NMR (125 MHz, DMSO-*d*_6_) of S19

### Supplementary Figure 107. ^1^H NMR (500 MHz, DMSO-*d*_6_) of S20

### Supplementary Figure 108. ^13^C NMR (125 MHz, DMSO-*d*_6_) of S20

### Supplementary Figure 109. ^1^H NMR (500 MHz, DMSO-*d*_6_) of 34

### Supplementary Figure 110. ^13^C NMR (125 MHz, DMSO-*d*_6_) of 34

### Supplementary Figure 111. ^1^H NMR (500 MHz, DMSO-*d*_6_) of S21

### Supplementary Figure 112. ^13^C NMR (125 MHz, DMSO-*d*_6_) of S21

### Supplementary Figure 113. ^1^H NMR (500 MHz, DMSO-*d*_6_) of S22

### Supplementary Figure 114. ^13^C NMR (125 MHz, DMSO-*d*_6_) of S22

### Supplementary Figure 115. ^1^H NMR (500 MHz, DMSO-*d*_6_) of 35

### Supplementary Figure 116. ^13^C NMR (125 MHz, DMSO-*d*_6_) of 35

### Supplementary Figure 117. ^1^H NMR (500 MHz, DMSO-*d*_6_) of S23

### Supplementary Figure 118. ^13^C NMR (125 MHz, DMSO-*d*_6_) of S23

### Supplementary Figure 119. ^1^H NMR (500 MHz, DMSO-*d*_6_) of S24

### Supplementary Figure 120. ^13^C NMR (125 MHz, DMSO-*d*_6_) of S24

### Supplementary Figure 121. ^1^H NMR (500 MHz, DMSO-*d*_6_) of 36

### Supplementary Figure 122. ^13^C NMR (125 MHz, DMSO-*d*_6_) of 36

### Supplementary Figure 123. ^1^H NMR (500 MHz, DMSO-*d*_6_) of S25

### Supplementary Figure 124. ^13^C NMR (125 MHz, DMSO-*d*_6_) of S25

### Supplementary Figure 125. ^1^H NMR (500 MHz, DMSO-*d*_6_) of S26

### Supplementary Figure 126. ^13^C NMR (125 MHz, DMSO-*d*_6_) of S26

### Supplementary Figure 127. ^1^H NMR (500 MHz, DMSO-*d*_6_) of 37

### Supplementary Figure 128. ^13^C NMR (125 MHz, DMSO-*d*_6_) of 37

### Supplementary Figure 129. ^1^H NMR (500 MHz, DMSO-*d*_6_) of 38

### Supplementary Figure 130. ^13^C NMR (125 MHz, DMSO-*d*_6_) of 38

### Supplementary Figure 131. ^1^H NMR (500 MHz, DMSO-*d*_6_) of S29

### Supplementary Figure 132. ^13^C NMR (125 MHz, DMSO-*d*_6_) of S29

### Supplementary Figure 133. ^1^H NMR (500 MHz, DMSO-*d*_6_) of S30

### Supplementary Figure 134. ^13^C NMR (125 MHz, DMSO-*d*_6_) of S30

### Supplementary Figure 135. ^1^H NMR (500 MHz, DMSO-*d*_6_) of 39

### Supplementary Figure 136. ^13^C NMR (125 MHz, DMSO-*d*_6_) of 39

### Supplementary Figure 137. ^1^H NMR (500 MHz, DMSO-*d*_6_) of S31

### Supplementary Figure 138. ^13^C NMR (125 MHz, DMSO-*d*_6_) of S31

### Supplementary Figure 139. ^1^H NMR (500 MHz, DMSO-*d*_6_) of S32

### Supplementary Figure 140. ^13^C NMR (125 MHz, DMSO-*d*_6_) of S32

### Supplementary Figure 141. ^1^H NMR (500 MHz, DMSO-*d*_6_) of 40

### Supplementary Figure 142. ^13^C NMR (125 MHz, DMSO-*d*_6_) of 40

### Supplementary Figure 143. ^1^H NMR (500 MHz, DMSO-*d*_6_) of 41

### Supplementary Figure 144. ^13^C NMR (125 MHz, DMSO-*d*_6_) of 41

### Supplementary Figure 145. ^1^H NMR (500 MHz, DMSO-*d*_6_) of S36

### Supplementary Figure 146. ^13^C NMR (125 MHz, DMSO-*d*_6_) of S36

### Supplementary Figure 147. ^1^H NMR (500 MHz, DMSO-*d*_6_) of 42

### Supplementary Figure 148. ^13^C NMR (125 MHz, DMSO-*d*_6_) of 42

1. LRMS were collected on some intermediates in place of HRMS.
2. HRMS were collected on all final benzamidoquinazolinones prior to running assays.
3. {Pan, 2022 #17}
4. At our institution, there is not the ability to decouple ^1^H coupling to ^19^F nuclei, therefore the image of the spectrum appears with splitting.
5. One carbon signal is not observed.
6. This material was contaminated with 4-methylbenzoic acid; as this sample was inactive in our assay (to 100 µM), no attempts were made to purify or characterize any further. HRMS matches product.
7. This material was contaminated with 4-methoxybenzoic acid; as this sample was inactive in our assay (to 100 µM), no attempts were made to purify or characterize any further. HRMS matches product.
8. This product was converted to the HCl salt by dissolving in CH_2_Cl_2_ and adding an excess of 2M HCl in Et_2_O, followed by concentrating *in vacuo*, triturating with hexanes, and drying under high vacuum to afford an off-white solid. Some solvent was retained in the final product despite drying. This HCl salt was what was handled for assays and for compound characterization.
9. One ^13^C signal is not visible in this spectrum.
10. Two ^13^C signals are not visible in this spectrum.
11. This molecule presents in NMR as ~5:1 mixture of rotamers. The peaks of the major rotamer are reported in the ^1^H line listing. For ^13^C NMR, please see image of spectra, as it was not possible to unambiguously assign peaks to each rotamer. The signal for the methyl carbon appears at 39.9 ppm, partially eclipsed by the residual solvent signal.
12. {Khan, 2010 #18}
13. This material was contaminated with 4-chlorobenzoic acid; as this sample was inactive in our assay (to 100 µM), no attempts were made to purify or characterize any further. HRMS matches product.
14. This material is contaminated with ring-opened hydrazide. The decision was made to telescope the material as the impurity is easier to purge at the next step.
15. This material was ~88-90% pure; because this sample was inactive in our assay (to 100 µM) no further purification was performed.
16. One ^13^C signal is not observed.
17. One ^13^C signal is not observed.
18. The methyl peak is eclipsed by the residual solvent signal ~ δ 2.50.
19. One ^13^C signal is not observed.
20. The peak for the benzylic carbon is eclipsed by the residual DMSO signal.
21. There are two peaks eclipsed here, for a total of 3H.
22. Small amount of contamination with benzoic acid.
23. This product contained impurities, and so the decision was made to telescope through the next reaction, and purge impurities there.
